# Systematic Discovery of Conservation States for Single-Nucleotide Annotation of the Human Genome

**DOI:** 10.1101/262097

**Authors:** Adriana Sperlea, Jason Ernst

## Abstract

Comparative genomics sequence data is an important source of information for interpreting genomes. Genome-wide annotations based on this data have largely focused on univariate scores or binary calls of evolutionary constraint. Here we present a complementary whole genome annotation approach, ConsHMM, which applies a multivariate hidden Markov model to learn *de novo* different ‘conservation states’ based on the combinatorial and spatial patterns of which species align to and match a reference genome in a multiple species DNA sequence alignment. We applied ConsHMM to a 100-way vertebrate sequence alignment to annotate the human genome at single nucleotide resolution into 100 different conservation states. These states have distinct enrichments for other genomic information including gene annotations, chromatin states, and repeat families, which were used to characterize their biological significance. Conservation states have greater or complementary predictive information than standard constraint based measures for a variety of genome annotations. Bases in constrained elements have distinct heritability enrichments depending on the conservation state assignment, demonstrating their relevance to analyzing phenotypic associated variation. The conservation states also highlight differences in the conservation patterns of bases prioritized by a number of scores used for variant prioritization. The ConsHMM method and conservation state annotations provide a valuable resource for interpreting genomes and genetic variation.

## Introduction

The large majority of phenotype-associated variants implicated by genome-wide association studies (GWAS) fall outside of protein coding regions.^1^ Identifying the causal variants and interpreting their biological role in these less well understood non-coding regions is a significant challenge.^2^ Large-scale mapping of epigenomic data across different cell and tissue types has been one approach for annotating and interpreting the non-coding regions of genomes.^3–5^ Using comparative genomics data to identify regions of evolutionary constraint has been a complementary approach for these purposes.^6–9^

In addition to providing evolutionary information, comparative genomics data has the advantage of providing information at single-nucleotide resolution. Furthermore, it is cell type agnostic and thus informative even when the relevant cell or tissue type has not been experimentally profiled.^10,11^ The most commonly used representations of this information are univariate scores and binary elements of evolutionary constraint, which are called based on a multiple species DNA sequence alignment and assumed models of evolution and selection.^8,9,12^– ^14^ Supporting the importance of these annotations, heritability analyses have recently implicated evolutionary constrained elements as one of the annotations most enriched for phenotype associated variants.^15^ These scores and elements have also been highly informative features to integrative methods for prioritizing pathogenic variants.^16–19^ Further improvements for predicting pathogenic variants in coding regions have been made to the integrative scores by incorporating features defined directly from a multiple sequence alignment.^20^

While highly useful, the representation of comparative genomics information into univariate scores or binary elements is limited in the amount of information it can convey about the underlying multiple sequence alignment at a specific base. This limitation has become more pronounced given the large number of species sequenced and incorporated into multi-species alignments such as a 100-way alignment to the human genome.^21^ Approaches have been developed to associate constrained elements, regions, or individual bases with specific branches in a phylogenetic tree.^22–28^ While also useful, such directed approaches are biased to only representing certain types of patterns present in the alignment. An alternative approach used for comparative genomic based annotation learned patterns of different classes of mutations between human and orangutan^29^, but this approach was only applicable at a broad region level and only incorporated information from one non-human genome.

Analogous to the many sequenced genomes available for comparative analysis, many different datasets are available for annotating the genome based on epigenomic data. Approaches that define ‘chromatin states’ based on combinatorial and spatial patterns in these datasets have effectively summarized the information in them to provide *de novo* genome annotations.^4,30–32^ Inspired by the success of these approaches, here we develop a method, ConsHMM, that extends ChromHMM^31^ to systematically annotate genomes into ‘conservation states’ at single nucleotide resolution. The conservation states assignments are based on the combinatorial and spatial patterns of which species align to and which match a reference genome at each nucleotide in a multiple species DNA sequence alignment. ConsHMM takes a relatively unbiased modeling approach that does not explicitly assume a specific phylogenetic relationship between species. The set of conservation patterns ConsHMM can infer are thus flexible and determined directly from the DNA sequence alignment.

We applied ConsHMM to assign a conservation state to each nucleotide of the human genome. These states are able to capture distinct enrichments for other genomic annotations such as gene annotations, CpG islands, repeat families, chromatin states, and genetic variation. We demonstrate that the conservation state annotations capture additional information that is not represented by scores or binary calls of constraint. We also show how the conservation states enable a deeper understanding of types of bases prioritized by a number of different scores used for variant prioritization, including those scores that integrate constraint information with a diverse set of other genomic annotations. Overall, these conservation state annotations are a resource for interpreting the genome and potential disease-associated variation, which complement both existing conservation and epigenomic-based annotations.

## Material and Methods

### Modeling conservation states with ConsHMM

ConsHMM takes as input an *N-way* multi-species sequence alignment to a designated reference genome. For each base in the reference genome, *i*, ConsHMM encodes information from the multiple species alignment into a vector, *v*_*i*_, of length *N*-1. An element of the vector, *v*_*i,j*_, corresponds to one of three possible observation for a non-reference species *j* at position *i*. The three possible observations are: (1) the non-reference species aligns with a non-indel nucleotide symbol present matching the reference nucleotide, (2) the non-reference species aligns with a non-indel nucleotide symbol present, but does not match the reference nucleotide, or (3) the non-reference species does not align with a non-indel nucleotide symbol present.

ConsHMM assumes that these observations are generated from a multivariate HMM where the emission parameters are assumed to be generated by a product of independent multinomial random variables, corresponding to each species in the alignment. Formally, the model is defined based on a fixed number of states *K*, and number of species in the multiple sequence alignment *N*. For each state *k (k =* 1,…,*K*), non-reference species *j* (*j* = 1,…,*N-1*) and possible observation *m* (*m* = 1, 2, or 3 as described above), there is an emission parameter: *p*_*k,j,m*_ corresponding to the probability in state *k* for species *j* of having observation *m*. For each possible observation *m*, let *I*_*m*_(*v*_*i,j*_) = 1 if *v*_*i,j*_ *= m*, and 0 otherwise. Let *b*_*t,u*_ be a parameter for the probability of transitioning from state *t* to state *u*. Let *c* ∈ *C* denote a chromosome, where *C* is the set of all chromosomes in the reference genome of the multiple species alignment, and let *L*_*c*_ be the number of bases on chromosome *c.* Let *a*_*k*_ (*k* = 1,…,*K*) be a parameter for the probability of the first base on a chromosome being in state *k*. Let *s*_*c*_ ∈ *S*_*c*_ be a hidden state sequence on chromosome *c* and *S*_*c*_ be the set of all such possible state sequences. Let *c*_*h*_ denote position *h* on chromosome *c*. Let 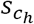 denote the hidden state at position *c*_*h*_ for state sequence *s*_*c*_.

We learn a setting of the model parameters that aims to optimize

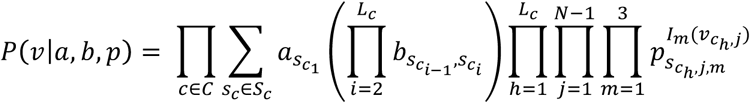

Once a model is learned, each nucleotide is assigned to the state with maximum posterior probability. To conduct the model learning and state assignments, ConsHMM calls an extended version of the ChromHMM^31^ software originally designed to solve an analogous problem of annotating a genome into chromatin states based on combinatorial and spatial patterns of the presence of different chromatin marks. The modeling in ConsHMM differs from the typical use of ChromHMM in three main respects: (1) the observation for each feature comes from a three-way multinomial distribution as opposed to a Bernoulli distribution, (2) it is applied at single nucleotide resolution as opposed to 200-bp resolution, (3) it is applied with more features than ChromHMM models have used in the past. (2) and (3) raise scalability issues in terms of time and memory, which we addressed in an updated version of ChromHMM (see below).

To apply ChromHMM in the context of three-way multinomial distributions, ConsHMM represents the three possible observations at position *i* for a species *j* with two binary variables, *y*_*ij*_ and *z*_*ij*_, corresponding to aligning and matching the reference genome respectively. *y*_*ij*_ has the value of 1 if the other species aligns to the reference with a non-indel nucleotide and 0 otherwise. *z*_*ij*_ has the value of 1 if the other species has the same nucleotide as the reference sequence and has a value of 0 if the other species has a different nucleotide present than the reference. In the case in which *y*_*ij*_=0, there is no nucleotide to compare to the reference and that value of the *z*_*ij*_ variable is considered missing (encoded with a ‘2’ for ChromHMM). If the value of an observed variable is missing, ChromHMM excludes the Bernoulli random variable corresponding to the observation from the emission distribution calculation at that position. For each state *k* and species *j*, ChromHMM thus learns two parameters, *f*_*k,j*_ and *g*_*k,j*_. *f*_*k,j*_ corresponds to the probability that at a given position in state *k*, species *j* aligns to the reference genome with a non-indel nucleotide that is P(*y*_*i,j*_=1| *s*_*i*_=*k*). *g*_*k,j*_ corresponds to the probability that at a given position in state *k*, species *j* matches the reference genome conditioned on species *j* aligning with a non-indel nucleotide that is P(*z*_*i,j*_ *=* 1| *y*_*i,j*_=1 and *s*_*i*_=*k*). This representation is equivalent to the three-way multinomial distribution, (*p*_*k,j,1*_, *p*_*k,j,2*_, *p*_*k,j,3*_) described above where *p*_*k,j,1*_ *= P(y*_*i,j*_*=1, z*_*i,j*_*=1 | s*_*i*_ *= k), p*_*k,j,2*_ *= P(y*_*i,j*_*=1, z*_*i,j*_*=0 | s*_*i*_ *= k),* and *p*_*k,j,3*_ *= P(y*_*ij*_*=0 | s*_*i*_ *= k),* since *p*_*k,j,1*_ *= f*_*k,j*_ ×*g*_*k,j,*_, *p*_*k,j,2*_ *= f*_*k,j*_ × *(1-g*_*k,j*_), and *p*_*k,j,3*_ *= 1 – f*_*k,j*_.

### Multiple species sequence alignment choice

Our method and software can be applied to any multiple species sequence alignment which is available in multiple alignment format (MAF) or which can be converted into this format. For the results presented here we applied it to the 100-way Multiz vertebrate alignment with human (hg19) as the reference genome.^21,33^

### Scaling-up ConsHMM to single base resolution with hundreds of features

Since for our application ConsHMM needs to run ChromHMM at single base resolution (‘-b 1’ flag) with 198 features after our binary encoding (2 for each non-human species in the 100-way alignment), we had to address scalability issues in terms of both memory and time. To address the memory issue we modified ChromHMM to support only loading in main memory input for chromosomes it is actively processing, as previously ChromHMM would only support loading all data into main memory upfront. This option can now be accessed in ChromHMM through the ‘-lowmem’ flag. To reduce the time required we used 12-parallel processors (‘-p 12’ flag) and we trained on a different random subset of the human genome on each iteration of the Baum-Welch algorithm. We divided each chromosome into 200kb segments (with the exception of the last segment of each chromosome which was less than this) in order to form random subsets of the human genome. We modified ChromHMM to allow training for each iteration on a randomly selected subset of 150 of these segments (‘-n 150’ flag), corresponding to 30MB per iteration. We ran this for 200 iterations by adding the ‘-d -1’ flag, which removed one of ChromHMM’s default stopping criterion based on computed likelihood change on the sampled data, since the likelihood is now expected to both increase and decrease between iterations as different sequences are sampled. These new options were included in version 1.13 of ChromHMM. The unique code to ConsHMM is written in Python. The code of ConsHMM shared with ChromHMM is written in Java and included with ConsHMM.

### Generating genome-wide annotations

After learning a 100-state model, we used it to segment and annotate the genome at base-pair resolution into one of the 100 conservation states. Each base in the human genome is classified into the state with the highest posterior probability. ConsHMM does this by running the MakeSegmentation command of ChromHMM. Due to computational constraints, the segmentation could not be generated for entire chromosomes at once. Instead, we ran MakeSegmentation on the same 200kb partitioning made for learning the model. We then merged the resulting files together using ConsHMM’s mergeSegmentation.py command with slice size parameter set to 200,000 (‘-s 200000’ flag) and the number of states parameter set to 100 (‘-n 100 flag’).

### Computing enrichments for external annotations

All overlap enrichments for external annotations were computed using the ChromHMM OverlapEnrichment command. OverlapEnrichment computes enrichments for an external annotation in each state assuming a uniform background distribution. Specifically the fold enrichment of a state for an external annotation is

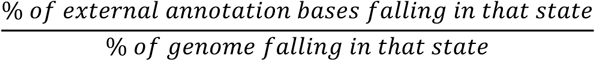

Positional enrichments of states relative to an anchor point from an external annotation were computed using the ChromHMM NeighborhoodEnrichment command at single base resolution (‘-b 1’ flag), single base spacing from the anchor point (‘-s 1’) and using the ‘-l’ and ‘-r’ flags to specify the size of the region of interest around the anchor point. The ‘-lowmem’ flag was also used for computing the enrichments for OverlapEnrichment and NeighborhoodEnrichment.

### External data sources for enrichment analyses

The external annotations of repeat elements were obtained from the UCSC genome browser RepeatMasker track.^21,34^ We generated an annotation for whether a base overlapped any repeat element, as well as separate annotations for bases falling in each class and family of repeat elements. The gene annotations were obtained from GENCODE v19 for hg19^35^. CpG Island annotations were obtained from the UCSC genome browser. Annotations of SNPs with >=1% minor allele frequency were obtained from the commonSNP147 track from the UCSC genome browser, which is based on dbSNP build 147. GWAS catalog variants were obtained from the NHGRI-EBI Catalog, accessed on Dec 5, 2016.^36^ For annotations of DNase I Hypersensitive Sites (DHS) processed by the Roadmap Epigenomics Consortium, we used Macs2 narrowPeak calls.^5^ The Fetal Brain and HepG2 DHS used were of epigenome samples E082 and E118 respectively. For the median non-exonic DHS enrichments and ranking of states in the heritability partitioning analysis we used narrowPeak calls from the ENCODE consortium.^3^ In the cases where ENCODE provided more than one replicate for a cell or tissue type, we used the first replicate.

PhyloP and PhastCons scores and constrained element calls were obtained from the UCSC genome browser. Assembly gap annotations were obtained from the Gap track from the UCSC genome browser. The context-dependent tolerance score (CDTS) used was that based on a cohort of 7784 unrelated individuals, following the analyses in Iulio et al.^37^, which focused on this version of the score. The CDTS and variants from this cohort were both lifted from hg38 to hg19 using the liftOver tool from the UCSC genome browser.^21^

### Choice of number of states

We learned models with each number of states between 2 and 100 states. We set 100 as the maximum number of states we would consider for computational tractability and maintaining a manageable number of states for analysis. The choice of a maximum of 100 also corresponds to the number of species used and allows for the possibility of each state to cover 1% of the genome. We analyzed the Bayesian Information Criterion (BIC) for models with each number of states between 2 and 100, and found that the BIC generally decreases as the number of states increases in the range considered (**Figure S1**). The BIC was calculated using the BIC_HMM function from the HMMpa R package.^38^ Analyzing the 100-state model’s internal confidence estimate of its state assignments also supported a larger number of states. Specifically, for each state in the 100-state model we computed the average posterior probability of that state at each base in the genome assigned to it, and confirmed consistently high average posterior probability values in the range [0.92,1.00] with a median of 0.97 (**Figure S2**). The posterior probabilities were computed by running the MakeSegmentation command in ChromHMM with the ‘-printposterior’ flag. We also investigated if additional states in models with larger number of states were biologically relevant. Specifically, we computed enrichments for various external annotations for models with each number of states between 2 and 100 to determine if biologically relevant enrichments were only robustly observed in models with more than a certain number of states. In the case of CpG islands, we observed that only models with at least 87 states consistently obtained >15 fold enrichment and only models with at least 95 states consistently obtained >30 fold enrichment (**Figure S3**). We saw a similar pattern of increasing enrichments for annotated transcription start sites (TSSs) for models with large number of states. We therefore decided to analyze the largest model, 100 states, that we were considering. We note that annotations based on chromatin states used fewer number of states, but were also defined on fewer features at a coarser resolution and had a less uniform genome coverage.^4,30,39^

### State clustering

We clustered the states based on the correlation of vectors containing the values *f*_*k,j*_ and *f*_*k,j*_ ×*g*_*k,j*_ for each species *j* defined above. State clustering was performed using the hclust hierarchical clustering function from the cba R package.^40^ The leaves of the resulting hierarchical tree were ordered according to the optimal leaf ordering algorithm^41^ implemented in the order.optimal R function from the cba package. We then cut the tree such that the 8 major groups of states were designated. The full tree is provided in **Figure S4**.

### Genome segmentation using uniform transition probabilities

For analyzing the effect of the transition probabilities on the genome segmentation, we created a separate model, which was the same model we used in the main analyses, except we set all transition probabilities to 0.01, corresponding to each state having an equal probability of transitioning to any state including itself. We then created a new genome segmentation by running the MakeSegmentation command in ChromHMM with this new model. For each state, we counted how many of the bases assigned to it in the original annotation were also assigned to it in the annotation created with the uniform transitions and divided this number by the number of bases in the state in the original annotation. This calculation provided a fraction from 0 to 1. We also reported the number of segments produced by each model, where a segment is defined to be one or more consecutive bases all assigned to the same state, such that any immediately adjacent bases are assigned to a different state or states.

### Gene Ontology enrichments

For each state and each protein coding gene based on GENCODE we computed the number of bases in that state that are within +/- 2kb of the gene’s TSS. In the case of genes with multiple annotated TSSs, we used the outermost TSS. We then created a ranking of genes for every state by sorting the genes in descending order of this number of bases. For each state we then created a set of 969 genes that represent the top 5% of genes in the state among the 19,397 genes we considered. We performed a Gene Ontology (GO) enrichment analysis (ontology and annotations files from Nov. 24^th^, 2016) for the top 5% genes in each state using the STEM software in batch mode with default options and the set of all genes considered as background.^42^ STEM computed an uncorrected p-value based on the hypergeometric distribution for each term displayed in the figures summarizing the analysis. STEM also reported corrected p-values for testing multiple GO terms for a single state based on randomization to three significant digits, which was less than 0.001 for all p-values mentioned in the main text.

### Transcription factor binding site motif enrichments

We computed the enrichment of the conservation states within 15 bases upstream and downstream of the center point of the POU5F1 and STAT known transcription factor-binding site motifs.^43^ The enrichment was computed relative to the background regions of the genome that were used to identify the motifs, which excluded repeat elements, coding sequence, and 3’ untranslated regions (UTRs). The *known1* version of the motifs was used for both motifs.

### Clustering of cell-type specific DNase I hypersensitive site enrichments

For the clustering of DHS analysis, we first computed the enrichments of all conservation states for DHS for 53 samples processed by the Roadmap Epigenomics consortium^5^, of which 16 were originally generated by the ENCODE project consortium.^3^ We then selected the subset of states that had a fold enrichment of at least 2 in at least one sample, leading to a subset of 21 conservation states. To more directly focus on each state’s relative enrichments across samples, we log_2_ transformed each enrichment value, and then normalized the enrichments for each state by subtracting the mean enrichment across samples and dividing by the standard deviation. We then hierarchically clustered the states based on the correlation of their enrichments across samples and hierarchically clustered the samples based on their correlations across states using the pheatmap R package.^44^ We also computed for each sample the fold enrichment of DHS bases for bases in CpG islands, as the ratio between the percent of DHS bases in CpG islands and the percent of the genome falling in CpG islands.

### Precision recall analysis for recovery of gene annotations and DHS

We randomly split the 200kb genome segments used for training the model and segmentation into two halves corresponding to training and testing data. For each target set in the precision-recall analyses we ordered the ConsHMM states in decreasing order of their enrichment for the target among the training set bases. We then used that ordering to iteratively add the testing set bases in each state to form cumulative sets of bases predicted to be of the target set and computed the precision and recall for them. For each constraint score we computed the precision-recall curve for predicting the target set in the test data using two methods. For the first method, we directly ordered bases in descending order of their assigned score. For the second method, we split the sorted scores into 400 bins such that each bin contains on average 0.25% of the genome, which was the size of the smallest state of the ConsHMM model (0.25% of the genome in state 100). Specifically, we assigned all bases in the genome where the score was not defined to one bin and then divided the remaining bases uniformly among the 399 other bins based on their score. In some cases score increments were at the boundary between two bins at their provided floating-point precision, or overlapped multiple bins. In these cases we uniformly split the target bases assigned to that score increment into multiple bins proportionally to the overall percentage of the score increment falling in each bin. We then treated the 400 bins as 400 states and followed the same procedure described for the ConsHMM states. We also computed the precision and recall of bases in each constrained element set for predicting the target set on the testing data. For the DHS analyses, we also separately evaluated recovery of DHS bases when restricting the analysis to non-exonic regions. Additionally, both genome-wide and within non-exonic regions, we evaluated the recovery of DHS bases when restricting the analysis to bases distal to a TSS, defined as more than 2kb from a TSS.

### Precision recall analysis for recovery of DHS bases aggregated across cell and tissue types

For the analysis of the recovery of DHS aggregated across cell and tissue types we concatenated DHS from 53 cell or tissue types processed by the Roadmap Epigenomics Consortium into one annotation in which each combination of chromosome and cell or tissue type effectively becomes a new chromosome. We then split the concatenated data into training and testing sets as described above. We computed the enrichments of the ConsHMM states and scores split into bins as detailed above, but multiplying the size of each state and bin by the number of DNase I hypersensitivity data sets. The precision and recall values for the ConsHMM states, constraint scores considered directly, constraint scores split into bins, and constrained element sets were then computed on the testing data.

### Enrichment analysis for constrained non-exonic elements assigned to phylogenetic branches

We lifted over the constrained non-exonic elements (CNEEs) from *Lowe et al*.^22^ from hg18 coordinates to hg19, using the liftOver tool from the UCSC genome browser with default settings.^21^ These elements were previously partitioned into subsets based on the inferred branch point of origin in a phylogenetic tree.^22^ We computed the enrichments of the conservation states for all the CNEEs and for each subset of the CNEEs separately, using the OverlapEnrichment command from ChromHMM at single nucleotide resolution (‘-b 1’ flag) and using the low memory option (‘-lowmem’). We also computed analogous enrichments for CNEEs overlapping PhastCons elements called on the same 100-way alignment that the conservation states were annotated based on. To compute the enrichments of CNEEs for bases in CpG islands we created an annotation consisting of a state for each CNEE subset and one additional state for bases not assigned to any CNEE. We then ran the same OverlapEnrichment command as above to compute enrichments of CNEE bases for non-exonic CpG islands, and non-exonic bases in general. The reported enrichment of CpG islands is the ratio of these two enrichments, effectively computing an enrichment relative to the non-exonic background. The set of non-exonic bases for the enrichment analysis was generated by excluding all bases annotated as an exon in GENCODE v19.

### Heritability partitioning analysis

The heritability partitioning was performed using the LD-score regression ldsc software. ^15^ We partitioned the PhastCons constrained elements into two halves based on a ranking of the conservation states. We focused on the PhastCons constrained elements for this analysis, since it was the only element set defined on the same alignments as our conservation states. We focused on halves since the LD-score regression estimates can be unstable for annotations covering too small of a percentage of the genome.^15^ To determine the two halves we ranked the conservation states in descending order of median fold-enrichment of non-exonic bases for DHS from 123 experiments from the University of Washington ENCODE group.^3^ We then divided bases in PhastCons elements between the top 7 ranked states (1-5, 8 and 28), which contain 51.9% of bases in PhastCons elements, and the bottom 93 states, which contain the other 48.1% of bases in PhastCons elements. We applied ldsc to these two sets for 8 traits (age at menarche, body mass index (BMI), coronary artery disease, educational attainment, height, low-density lipoprotein (LDL) levels, schizophrenia and smoking behavior), all of which were previously considered in heritability partitioning analysis.^15^ We followed the procedure for partitioning heritability as done in *Finucane et al.*^15^, including using the baseline annotation set and 500 base-pair windows around annotations to dampen the artificial inflation of heritability in neighboring regions caused by linkage disequilibrium. The baseline annotation set contains a range of annotations including DHS. For our analysis, we first removed the constrained element set already included in the baseline annotation set, then added our two halves of PhastCons elements and finally ran the ldsc software on the full set of annotations.

### Enrichment analysis for variant prioritization scores

For each variant prioritization score included in the conservation state enrichment analysis of prioritized bases, we extracted the top 1%, 5% and 10% of all the bases ranked by each score, both genome-wide and just in non-coding regions. The non-coding regions were defined as the intersection of where the LINSIGHT and FunSeq2 scores provided a value, as these two scores were only defined on non-coding regions. This intersection results in a set of bases covering 90% of the genome that excludes coding regions in addition to other regions filtered for technical reasons by either of the two methods.^19,45^ For each score we chose the score threshold that gave us a size for the top set that was as close as possible to the target percentage, which did not always exactly match the target percentage due to the precision of the scores. If a score did not provide a value for a particular base being considered, then that base was assigned to the lowest value of that score, but would still be counted when establishing the percentage thresholds. For the scores that provided separate score values for alternate alleles at a certain position, we used the maximum of the values for all alleles. The state enrichments were then computed using the OverlapEnrichment command from ChromHMM at single base resolution (‘-b 1’ flag) and with the low memory option (‘-lowmem’ flag). For the analysis restricted to non-coding regions, we also computed the enrichment of the states for this background region using the same command. The enrichment for each score in a state was then divided by the enrichment of the background region for the state. For the Eigen and Eigen-PC scores we used version 1.1, for FunSeq2 we used version 2.1.6, and for CADD we used both v1.0 and v1.4.

## Results

### Annotating the human genome into conservation states

We developed an approach, ConsHMM, to annotate a genome into different conservation states based on a multiple species DNA sequence alignment (**Figure 1A, Methods**). We model the combinatorial patterns within the alignment of which species align to and which match a reference genome, for which we used the human genome. Specifically, at each nucleotide in the human genome we encode one of three possible observations for each non-reference species in the alignment: (1) aligns with a nucleotide present that is the same as the human reference genome, (2) aligns with a nucleotide present that is different than the human reference genome, or (3) does not have a nucleotide present in the alignment for that position. We further model these observations as being generated from a multivariate hidden Markov model (HMM), which probabilistically captures both the combinatorial patterns in the observations and their spatial context. Specifically, we assume that in each state the probability of observing a specific combination of observations is determined by a product of independent multinomial random variables. The parameter values of these multinomial random variables will differ between states and are learned from the data. After the model is learned, each nucleotide in the human genome is assigned to the state that had the maximum posterior probability of generating the observations.

**Figure 1:**
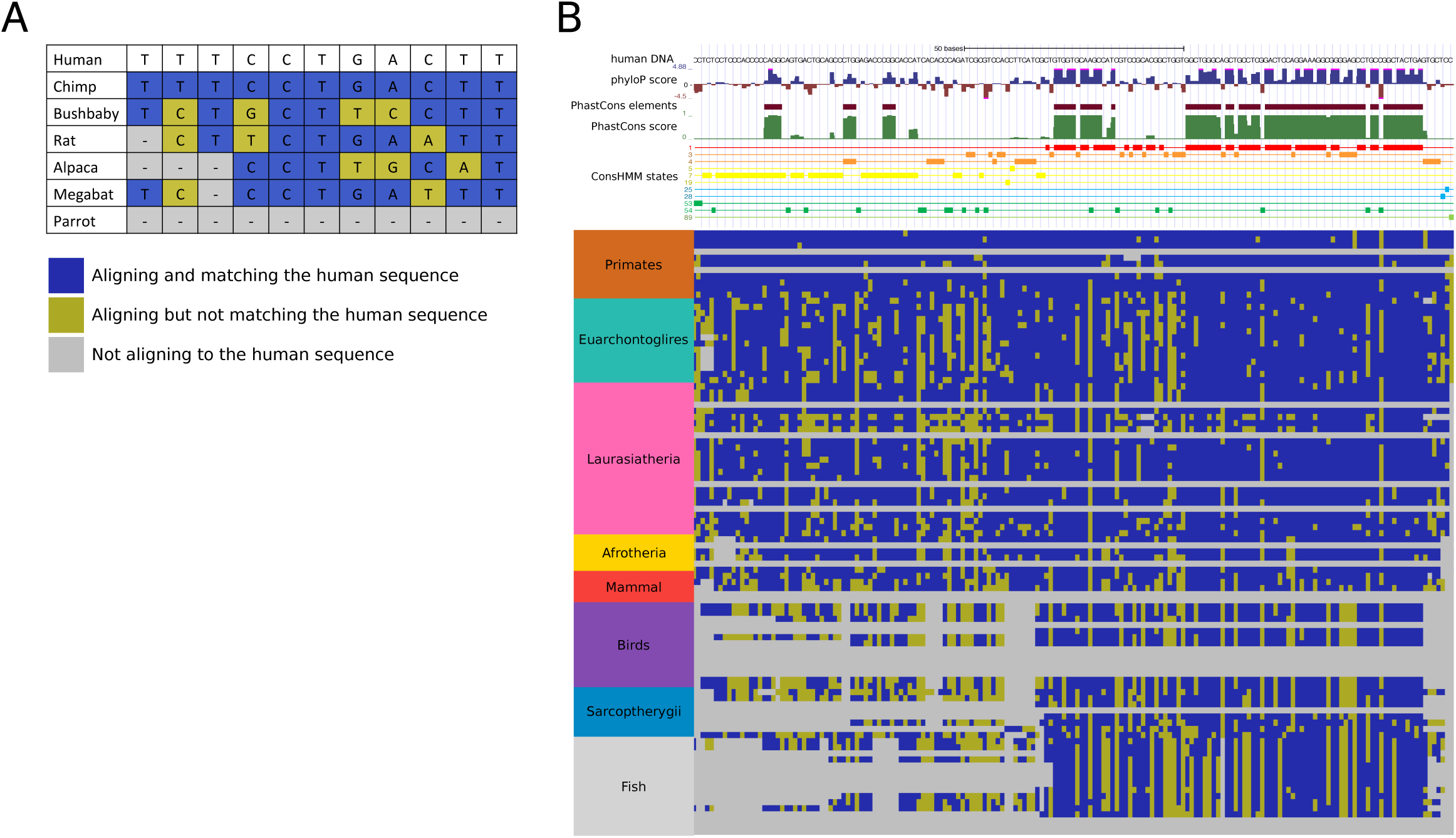
Illustration of ConsHMM modeling approach. **(A)** The input to ConsHMM is a multi-species alignment, which is illustrated for a subset of 6 species aligned to the human sequence. At each position and for each species ConsHMM represents the information as one of three observations: (1) aligns with a non-indel nucleotide matching the human sequence shown in blue, (2) aligns with a non-indel nucleotide not matching the human sequence shown in yellow, or (3) does not align with a non-indel nucleotide shown in gray. **(B)** Illustration of conservation state assignments at a locus chr22:25,024,640-25,024,812. Only states assigned to at least one nucleotide in the locus are shown. Below the conservation state assignments is a color encoding of the input multiple sequence alignment according to panel (A). The major clade of species as annotated on the UCSC genome browser^21^ are labeled and ordered based on divergence from human. Above the conservation state assignments are PhastCons constrained elements and scores and PhlyoP constraint scores. This figure and **Figure S9** together illustrate that positions of nucleotides that have the same status in terms of being in a constrained element or not or have similar constraint scores can be assigned to different conservation states depending on the patterns in the underlying multiple-species alignment.

ConsHMM builds on ChromHMM^31^, which has previously been applied to annotate genomes based on epigenomic data at 200-bp resolution^30^, to now annotate genomes at single nucleotide resolution based on a multiple species DNA sequence alignment (**Methods**). We applied ConsHMM to a 100-way Multiz vertebrate alignment with the human genome and focused our analysis here on a model learned using 100 states in order to balance recovery of additional biological features and model tractability (**Figures 2** and **S1-S8, Tables S1** and **S2, Methods**). We note that HMMs have previously been used to provide local smoothing of signal for the task of identifying constrained elements.^9,14^ We verified that the HMM had a smoothing effect in our application by comparing to a segmentation derived from a model with the same emission parameters as our learned model, but that ignored the information in the transition parameters (**Methods**). We saw an increase in the number of segments from 889 million to 1.06 billion when not using the transition information, though a large majority of the state assignments to individual bases were the same (**Figure S9**). We illustrate the ConsHMM conservation state annotations at two different loci showing that different bases that are associated with calls of evolutionary constraint from existing approaches can have very different underlying alignment patterns and conservation state assignments (**Figures 1B** and **S10**). Conservation state annotations genome-wide are available online (**Web Resources**).

**Figure 2:**
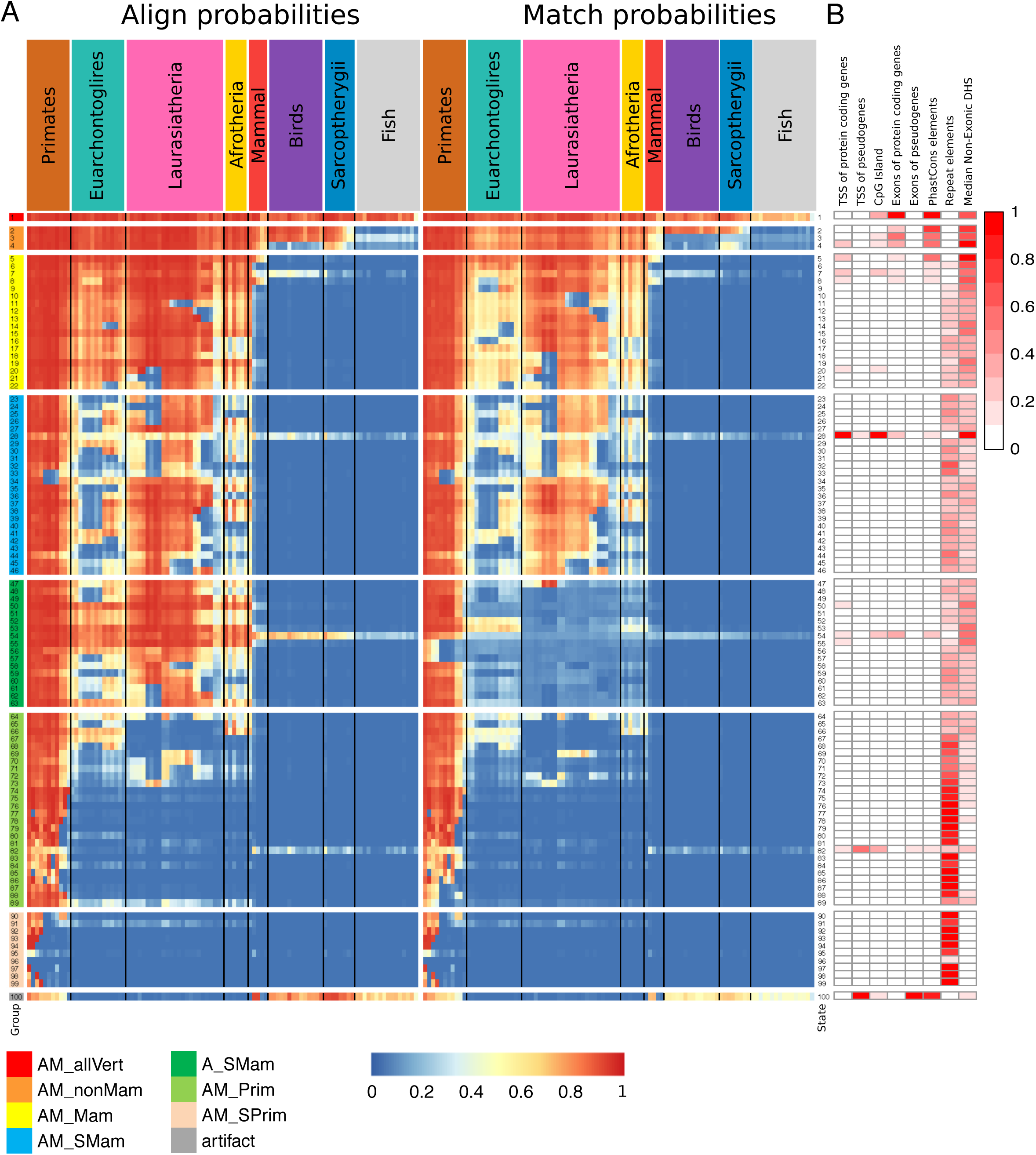
Conservation state emission parameters learned by ConsHMM and enrichments for other genomic annotations. **(A)** Each row in the heatmap corresponds to a conservation state. For each state and species, the left half of the heatmap gives the probability of aligning to the human sequence, which is one minus the probability of the not aligning emission. Analogously, the right half of the heatmap gives the probability of observing the matching emission. Each individual column corresponds to one species with the individual names displayed in **Figure S5.** For both halves, species are grouped by the major clades and ordered based on the hg19.100way.nh phylogenetic tree from the UCSC genome browser, with species that diverged more recently shown closer to the left.^21^ The conservation states are ordered based on the results of applying hierarchical clustering and optimal leaf ordering.^41^ The states are divided into eight major groups based on cutting the dendrogram of the clustering. The groups are indicated by color bars on the left hand side and a white row between them. Transition parameters between states of the model can be found in **Figure S6**. **(B)** The columns of the heatmap indicate the relative enrichments of conservation states for external genomic annotations (**Methods**). For each column, the enrichments were normalized to a [0,1] range by subtracting the minimum value of the column and dividing by the range and colored based on the indicated scale. Values for these enrichments and additional enrichments can be found in **Figure S8** and **Table S2** and enrichments for individual repeat classes and families can be found in **Figure S14.**

### Major groups of conservation states

We hierarchically clustered the conservation states based on their align and match probabilities, and then cut the resulting dendrogram to reveal eight notable groups of states or distinct individual states (**Figures 2A** and **S4, Table S3, Methods**). We named the resulting groups based on the aligning and matching properties of major subsets of species for most states in each group. We also summarized for each individual state the most distal species to human that had a majority of positions aligning and the closest one that did not, and similarly for matching (**Table S3**). The first of these subsets of states was a single state (State 1; AM_allVert) that showed high align and match probabilities through essentially all vertebrate species considered. The second subset showed relatively high align and match probabilities for all mammals and some non-mammalian vertebrates (States 2-4; AM_nonMam). The third subset showed relatively high align and match probabilities for most if not all mammals, but not non-mammalian vertebrates (States 5-22; AM_Mam). The fourth subset showed high align probabilities for many mammalian species, but had low align probabilities for notable species such as mouse and rat for many of the states in the group (States 23-46; AM_SMam). The combination of the absence of mouse and rat alignments with the presence of mammals that are assumed to have diverged earlier is consistent with the previously observed increased substitution rates for mouse and rat.^7^ The fifth subset showed high align probabilities for many mammalian species, but did not show high match probabilities (States 47-63; A_SMam). The sixth subset showed high align probabilities for most primates, but not for other species (States 64-89; AM_Prim). The seventh subset showed high align probabilities for at most a subset of primates (States 90-99; AM_SPrim). The final subset was a single state (State 100; artifact) that showed a noteworthy pattern of high align and match probabilities for most primates and non-mammalian vertebrates, but low probabilities for non-primate mammals, consistent with a previous observation that inclusion of non-mammalian vertebrates can be associated with increased presence of suspiciously aligned regions.^46^

### Conservation states exhibit distinct patterns of positional enrichments relative to gene annotations and regulatory motif instances

The conservation states showed strong and distinct positional enrichments relative to GENCODE^35^ annotated gene features including transcription start sites (TSS), transcription end sites (TES), exon start sites, and exon end sites for both protein coding genes and pseudogenes (**Figures 3A-D** and **S11**). Notable positional enrichments were also seen for regulatory motifs instances (**Figure 3E** and **3F**). Relative to starts of exons of protein coding genes seven of the states (States 1-4, 7, 28, and 54) had 13 fold or greater enrichment for some position within 20 base pairs of exon starts, both when considering all such exons and subsets of exons in specific coding phases (**Figures 3A** and **S11A-C**). These seven states were the only states that had a majority of positions aligning for at least some non-mammalian vertebrates, while still having a majority of positions aligning for all primates (**Figure 2A** and **Table S3**). Within exons we saw the strongest enrichment for states 1-4 and 54, and among these state 1 showed the strongest enrichment, as expected given its high match probabilities through all vertebrates (**Figures 2B, 3A-B,** and **S11A-E**). Interestingly, state 1 showed very strong enrichment (>80 fold) in the two nucleotides immediately upstream of the exon start, with the third upstream nucleotide also having high enrichment (46 fold) (**Figure S11C**). These three nucleotide positions correspond to the positions of the canonical 3’ splice site sequence that is highly conserved throughout vertebrates.^47^ At the ends of exons of protein coding genes (**Figure 3B**), state 1 maintained a >40 fold enrichment for six nucleotides past the end of coding sequence corresponding to positions of the known canonical 5’ splice site sequence.^47^ Downstream of the start of protein-coding exons, the enrichment profile for state 1 showed a 3-bp oscillation period, with a dip of enrichment at each 3rd base corresponding to codon wobble positions. In contrast, states 3 and 54, which were both associated with high align probabilities through many vertebrates and lower match probabilities, showed the inverse oscillation pattern to state 1 (**Figures 3A** and **S11A-C**).

**Figure 3:**
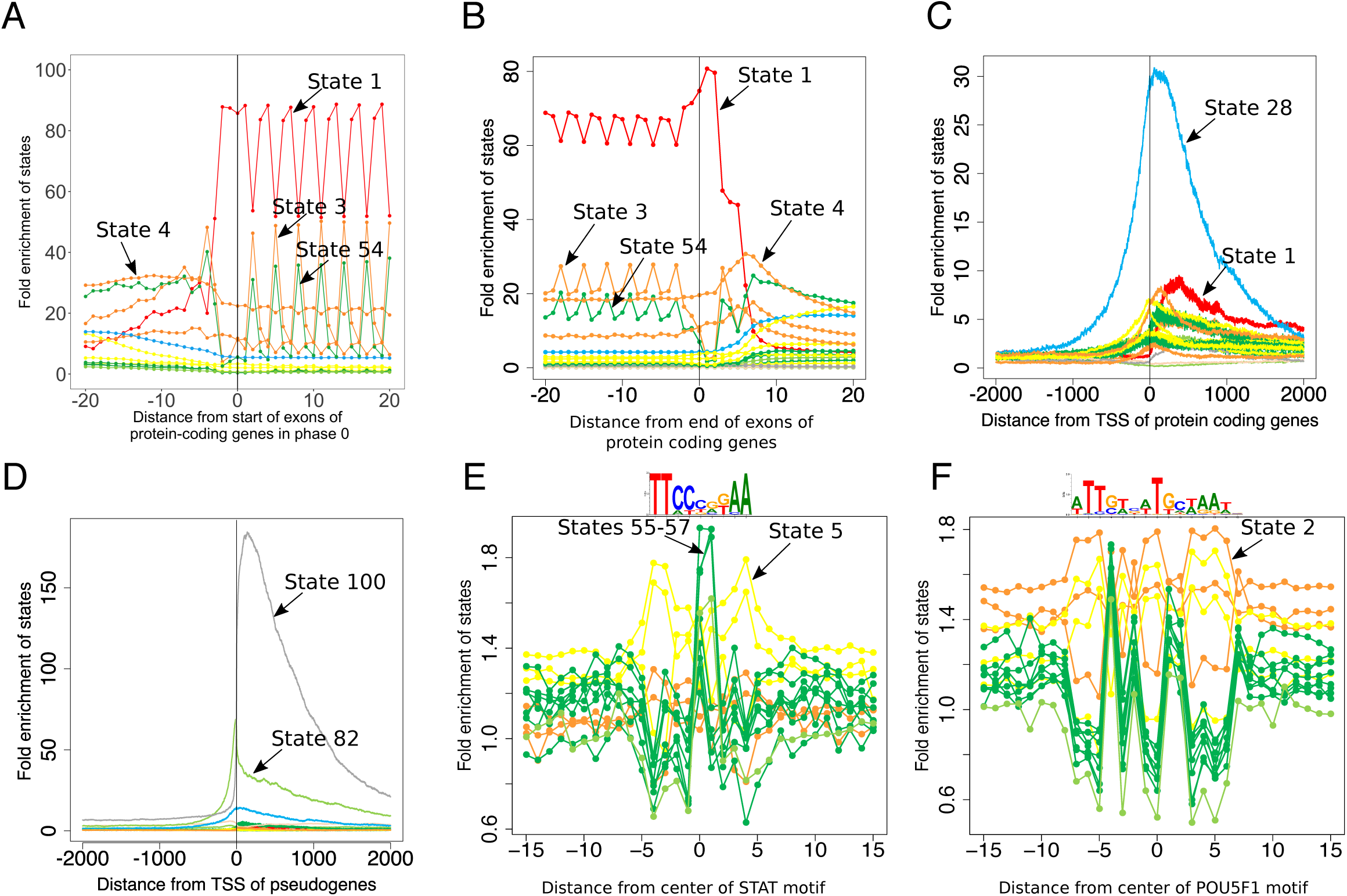
Conservation state positional enrichments. Plots of positional fold enrichments of conservation states relative to **(A)** start of exons of protein coding genes in phase 0, **(B)** end of exons of protein coding genes and **(C,D)** TSS of **(C)** protein coding, and **(D)** pseudogenes genes. Positive values represent the number of bases downstream in the 5’ to 3’ direction of transcription, while negative values represent the number of bases upstream. Enrichments relative to gene annotations are based on a genome-wide background. The subset of states included in panels (A)-(D) were the states that had at least a 3 fold enrichment at some position within +/-2kb from the anchor point. **(E,F)** Also shown are positional plots relative to the central nucleotide of a set of instances of **(E)** STAT and **(F)** POU5F1 motifs. The subset of states included in (E), (F) are the states that had an enrichment of at least 1.5 for some position within +/-15bp from the center nucleotide of either motif. Enrichments for motif instances were computed relative to the portion of the genome scanned for regulatory motifs in Kheradpour and Kellis (2014), which excludes coding, 3’UTRs, and repeat elements. Additional position enrichment plots can be found in **Figure S11**.

Relative to TSS of protein coding genes, state 28 had the strongest enrichment reaching a maximum enrichment 30 fold at the TSS (**Figure 3C**). State 28 was associated with moderate align and match probabilities for almost all the species present in the alignment. Consistent with its enrichment for TSSs state 28 also had the greatest enrichment for CpG islands (32 fold). However, state 28 also showed a 20 fold enrichment of CpG islands >2kb away from any TSS of protein coding genes and a 10 fold enrichment for TSS of protein coding genes >2kb away from a CpG island, suggesting the possibility that both of these features are making a partially independent contribution to the association, or the presence of additional unannotated TSSs that are associated with CpG islands.^48^ Relative to TES of protein coding genes we saw the enrichment peak for state 2 at almost 12 fold (**Figure S11F**), which had high align and match probabilities for almost all vertebrates except for fish.

Relative to pseudogene exon starts and ends, states 100 and 82, both associated with alignability to distal vertebrates without many mammals closer to human (**Figure 2B** and **Table S3**), had strong enrichments peaking at greater than 100 and 38 fold respectively (**Figure S11G** and **S11H**). These two states also showed the greatest enrichment relative to TSSs of pseudogenes peaking at 184 and 68 fold for states 100 or 82 respectively (**Figure 3D**) and for TESs of pseudogenes peaking at 199 and 61 fold respectively (**Figure S11I**).

Relative to instances of regulatory motifs, different conservation states showed single nucleotide enrichment variation, often associated with variation in the amount of information in the positional-weight matrix (**Figure 3E** and **3F, Methods**).^43^ For example, in the case of the POU5F1 and STAT motifs we saw state 2 from the AM_nonMam group and state 5 from the AM_Mam group respectively reach 1.8 fold enrichments, but have lower enrichments (1.4-1.5) at some nucleotides with lower information content. For the STAT motif, states 55-57, associated with high align probabilities for most mammals, but high match probabilities only for a few primates, showed enrichments that peaked at the CG dinucleotide in the center of the motif, consistent with their genome-wide enrichments for CG dinucleotides (**Figures 3E and S12**).

### Enrichment of conservation states for different gene classes

The previous analyses demonstrated that different conservation states have distinct enrichments in promoter regions of genes. We next investigated whether different conservation states also exhibit distinct enrichments for different classes of genes after controlling for the state’s relative preference for promoter regions. Specifically, for each state we determined the 5% of genes with the greatest presence of the state in its promoter region and evaluated Gene Ontology (GO) enrichments for those genes, revealing distinct enrichment patterns (**Figures 4B** and **S13, Methods**). For example, even among states 1-3, all of which had high alignability through at least birds and matching through mammals, we observed substantial differences in their gene preferences. Out of these three states, state 1 (the AM_allVert group) was the only one enriched for nucleosomes (p<10^-41^; 10.5 fold), while state 3, which had high matching only through mammals, was the only one with a significant enrichment for a set of genes related to sensory perception of smell (p<10^-300^; 15.5 fold). State 2, which had high align and match probabilities through all vertebrates except fish, was the state most enriched for cellular developmental processes (p<10^-30^; 1.8 fold), which did not show enrichment in state 3. We also observed notable enrichments for states with overall lower align or match probabilities. For example, state 89, associated with high alignability and low matching in primates as well as some alignability and low matching in non-primate mammals, was the state most enriched for antigen binding (p<10^-14^; 6.7 fold). This is consistent with antigen binding being associated with many species, but fast evolving.^49^

**Figure 4:**
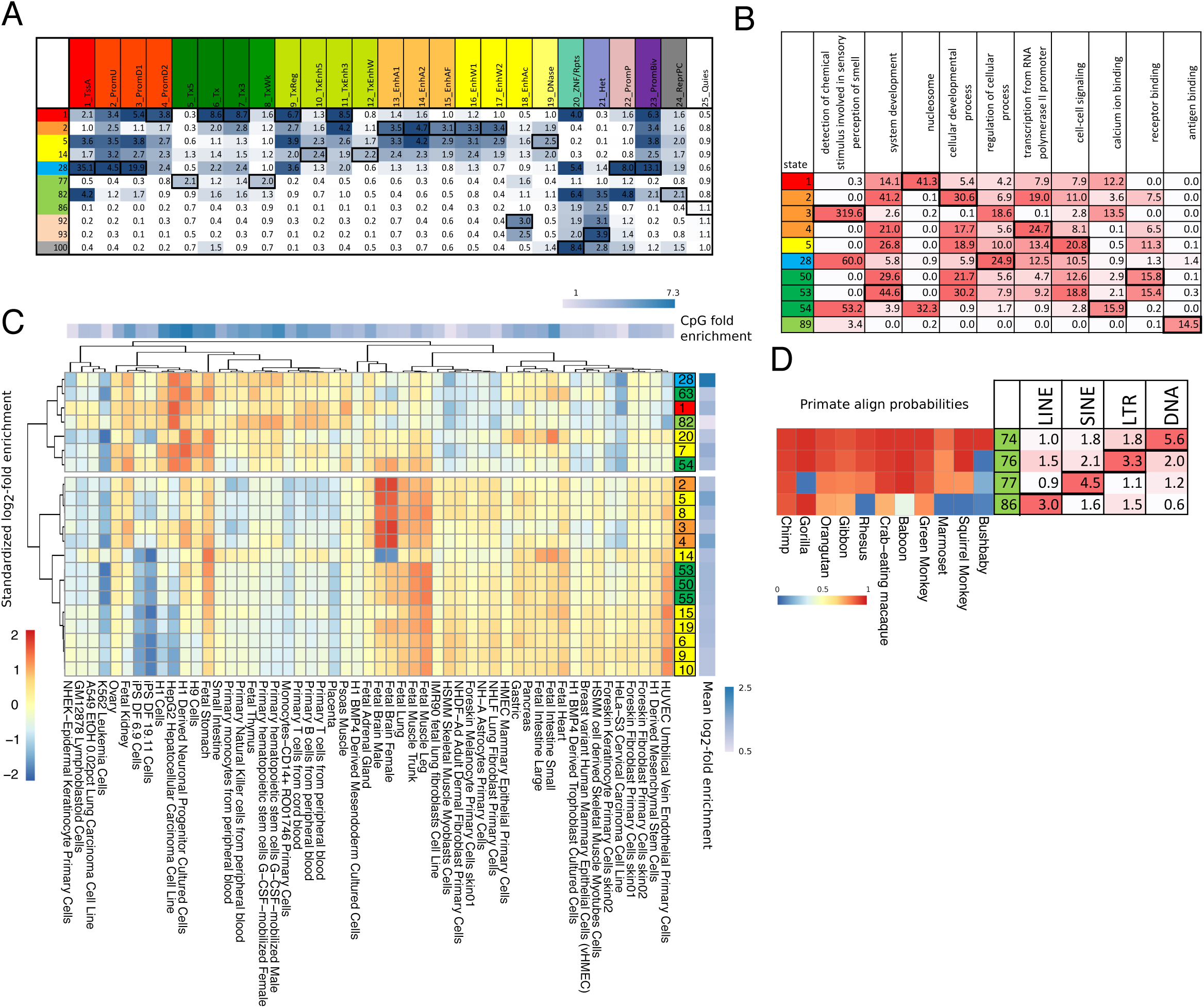
Conservation states enrichment for chromatin states, GO terms, DHS and repeat elements. **(A)** Median fold enrichment of conservation states (rows) for one of 25 chromatin states from a previously defined chromatin state model defined across 127 samples of diverse cell and tissue types (columns).^15^ Only conservation states that had the maximum value for at least one chromatin state are shown, and those values are boxed. See **Figure S15** for the enrichments of all conservation states. **(B)** –log_10_ p-value (uncorrected) of the conservation states (rows) for the GO term (columns) where each conservation state is associated with its top 5% genes based on promoter regions (**Methods**). Only GO terms which were the most enriched term for some conservation state are shown, restricted to the top 10 terms based on the significance of the enrichment. Only conservation states that had the most significant enrichment for one of the displayed GO terms are shown, with the maximal enrichments boxed. The full set of conservation states with additional GO terms are in **Figure S13**. **(C)** Relative enrichments of conservation states for DHS across cell and tissue types. Only conservation states with at least a 2 fold enrichment in one sample considered are shown. Enrichment values were log_2_ transformed and then row normalized by subtracting the mean (right heatmap) and dividing by the standard deviation. States and experiments were then hierarchically clustered and revealed two major clusters. In the top cluster conservation states showed the greatest enrichment for experiments in which the DHS also strongly enriched for CpG islands (top heatmap). In the bottom cluster conservation states generally had the strongest relative preference for a number of fetal related samples. **(D)** Enrichment of conservation states with the maximal enrichment for LINE, SINE, LTR or DNA repeats next to the state align probabilities for primates. These states all had low align probabilities outside of primates, but their differences among primates corresponded to substantial differences in repeat enrichments.

### Enrichments for repeat elements in conservation states

The conservation states showed a wide range of enrichments and depletions (from 2 fold enrichment to 133 fold depletion) for bases overlapping any repeat element (**Figures 2B** and **S8**).^21,34^ Of the 25 states that did not have any species outside of primates with a majority of positions aligning, all but two had an enrichment of 1.55 or greater for repeat elements, while the other 75 states all either had an enrichment below that or did not show enrichment (**Table S3**). The two exceptions were state 89 and state 96, neither of which showed enrichment for repeat elements. As noted above, state 89 is likely associated with fast evolving bases shared with some non-primate mammals, as opposed to bases new to primates. State 96 is associated with assembly gaps (**Figure S8**). Different repeat classes and families had distinct patterns of enrichments for different states, even though in some cases the difference in state parameters was subtle (**Figures 4D** and **S14**). For instance, among states in the AM_Prim group, which primarily differed in terms of the specific combinations of primates with high align and match probabilities, we found distinct enrichments. Notably, four different states from the group AM_Prim, 74, 86, 76, and 77, showed maximal enrichments for the DNA, LINE, LTR, and SINE repeat classes respectively (**Figure 4D**). State 74, which is characterized by high align and match probabilities for all primates, had an enrichment of 5.6 fold for DNA repeats, while the enrichment for the other three classes were between 1.0 and 1.8 fold. On the other hand, state 86, which lacked alignability of a subset of primates, had a 3.0 fold enrichment for LINE repeats, while the enrichment for the other classes were between 0.6 and 1.6 fold. States 76 and 77 had 3.3 and 4.5 fold enrichments for LTR and SINE respectively compared to 1.1 and 2.1 fold for SINE and LTR respectively. State 76 and state 77 both had high align probabilities through primates up to and including squirrel monkey, with the exception that state 77 lacked alignability to gorilla. Despite these subtle differences in the alignment probabilities, these states had substantial differences in their repeat enrichment profiles.

### Relationship of conservation states to chromatin states

We compared our conservation states to annotations of the genome based on a 25-chromatin state model defined on 127 samples of diverse cell and tissue types using imputed data (**Figures 4A** and **S15**).^5,39^ For each conservation state we determined the median enrichment of each chromatin state across the 127 samples. Eleven different conservations states were maximally enriched for at least one of the 25-chromatin states. Conservation state 28 showed the greatest enrichment for any chromatin state, with a 35 fold enrichment for a chromatin state associated with active promoters, and was maximally enriched for four other promoter associated chromatin states. Conservation state 1 was maximally enriched for five chromatin states all associated with transcribed and exonic regions^39^ (3.8-8.7 fold), which is consistent with this conservation state being most enriched for exons. Conservation state 2 had the maximal enrichment for five enhancer associated chromatin states (3.1-4.7 fold), while conservation state 5 had high enrichments for these states and also had the greatest enrichment of any conservation state for a chromatin state primarily associated with just DNase I hypersensitivity (2.5 fold). These and other distinct enrichments of the conservation states for the different chromatin states highlight that conservation states are able to capture multi-dimensional information in the genome.

### Conservation states capture enrichment patterns of DNase I hypersensitive sites across cell and tissue types

The previous analysis demonstrated that conservation states can exhibit different enrichment patterns for different chromatin states. We next investigated whether different conservation states also capture distinct enrichment patterns for a chromatin mark across cell and tissue types. For this we analyzed DNase I hypersensitive sites (DHS) from 53 of the 127 samples considered above for which maps of experimentally observed DHS were available from the Roadmap Epigenomics Consortium.^5^ We focused on the 21 conservation states that exhibited at least 2 fold enrichment in at least one sample (**Figure 4C**). We then row normalized the enrichments in order to focus on the relative enrichment patterns across cell and tissue types (**Methods**). Hierarchical clustering of the enrichment patterns revealed two major clusters of states (**Figure 4C**). One of these clusters contained 14 of the 21 states and was associated with strong enrichments for fetal related samples. Ten of the states in this cluster have maximum enrichment for a fetal sample, while the remaining four states have maximum enrichment for the cell type Human Umbilical Vein Endothelial Cells (HUVEC). The second major cluster consisted of seven states, all of which were enriched for CpG islands (**Figures 2B** and **S8**). The DHS from samples that showed the greatest enrichments in states in these clusters also had the greatest enrichment of CpG islands (**Figure 4C, Methods**), but were biologically diverse in terms of the type of cell or tissue and could potentially reflect technical experimental differences.

### Relationship of conservation states to constraint based annotations

We next investigated the relationship of our conservation state annotations with calls and univariate scores of evolutionary constraint. Specifically, we considered constrained element sets based on four methods (GERP++, SiPhy-omega, SiPhy-pi, and PhastCons) and constraint scores based on three methods (GERP++, PhastCons, and PhyloP) publicly available for hg19 and also defined on Multiz alignments. The PhastCons and PhyloP scores and elements we compared to were defined on the same 100-way vertebrate alignment. The available GERP++, SiPhy-omega, and SiPhy-pi score and elements were derived from different versions of Multiz alignments and only considered mammals.

We consistently found conservation states 1-5 to be highly enriched (>9 fold) for all constrained element sets (**Figures 2B** and **S16A**). These states were also among the top six states in terms of mean score for constraint scores considered (**Figure S16B**). Consistent with this, states 1-5 were the states that had the highest average matching probability across mammals. Two other states exhibited at least 6 fold enrichment for at least one constrained element set: states 54 and 100. State 100, associated with putative artifacts, showed high enrichments for PhastCons elements (15 fold) and high average scores for PhastCons and PhyloP. This is consistent with this state having high aligning and matching probabilities primarily in non-mammalian vertebrates and these elements and scores being defined using such species. State 54 was consistently enriched for all the constrained elements (4-7 fold), but did not show high mean base-wise scores particularly for the GERP++ and PhyloP scores. This difference of high enrichment in constrained elements but not base-wise scores is consistent with state 54 having high alignability through most vertebrates, but low matching outside primates. More generally, we found that constrained element calls did not have the resolution to exhibit biologically relevant single nucleotide variation in enrichments around regulatory motifs and exon start and ends as we saw with our conservation state annotations, with the exception of those from PhastCons (**Figures 3** and **S17**).

The objective of our conservation state annotations is different than that of binary calls and univariate scores of evolutionary constraint, which have a more specific and complementary goal. However, to better understand their relative biologically relevant information we compared their ability to recover annotated starts and ends of exons and TSS and TES of genes separately for protein coding and pseudogenes, as there are well established genome annotations of these features (**Figures 5A-C** and **S18**). In almost all cases the conservation states had greater information available for recovering annotated gene features. The only exceptions were that PhyloP scores could achieve higher precision at low recall levels for protein coding exon starts and ends, and that SiPhy-pi elements had slightly higher precision for TSS of protein coding genes at their one recall point.

**Figure 5:**
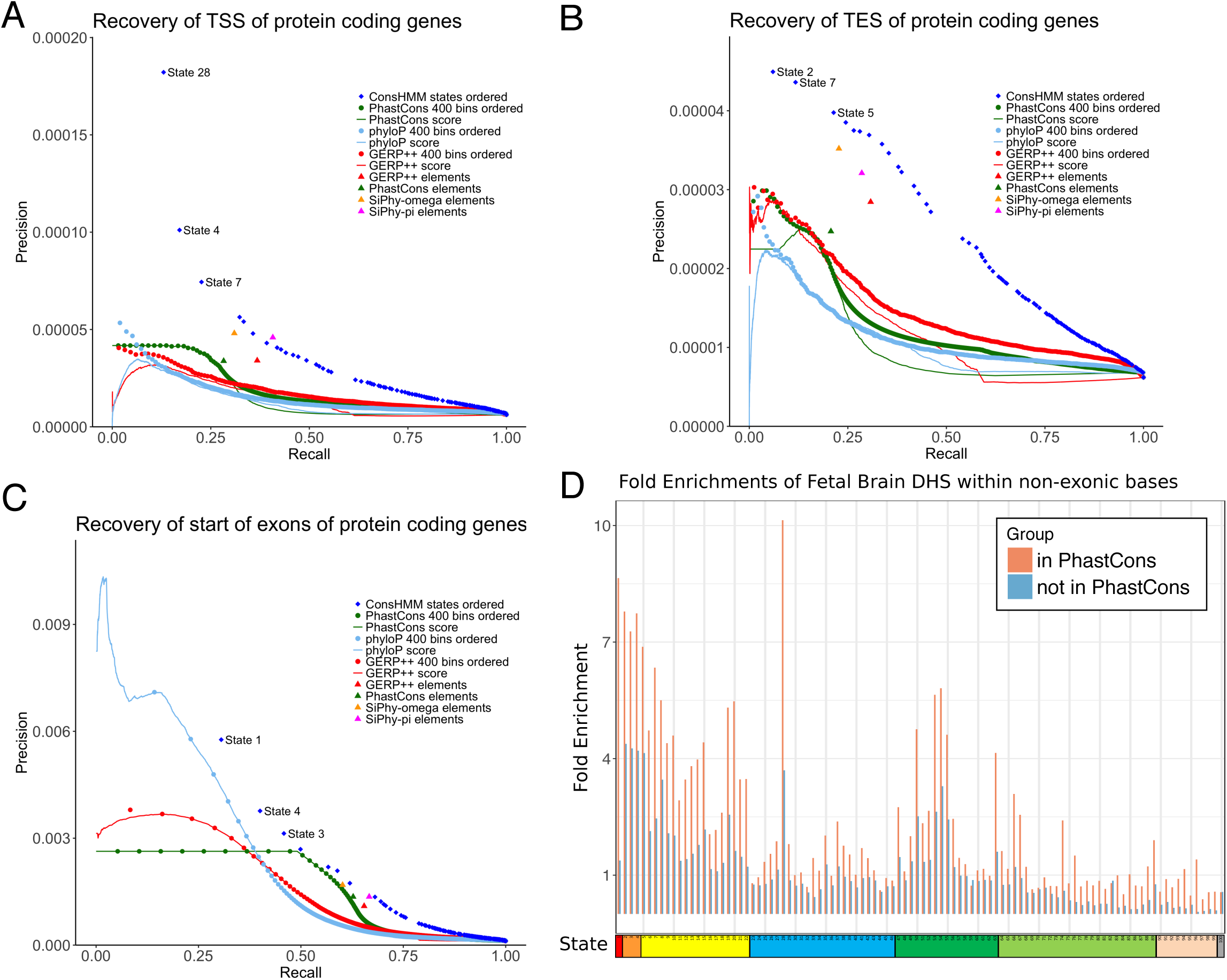
Relationship of conservation states with constrained elements and scores. Precision-recall plots for recovery of **(A)** TSS of protein coding genes, **(B)** TES of protein coding genes, and **(C)** the start of exons of protein coding genes. Recovery based on ordering ConsHMM conservation states for their enrichment for the target set in the training data, then cumulatively adding the states in that ranked order and evaluating on the test data is shown with a series of blue dots (**Methods**). The first few conservation states added are labeled with their state number. Recovery based on ranking from highest to lowest value of constraint scores is shown with continuous lines. Recovery based on score partitioning into 400 bins and subsequent ordering based on enrichment for the target set in the training data, then cumulatively adding bins in that ranked order and evaluating on the test data is shown in a series of dots of the same color as the continuous line corresponding to the score. Recovery of target test bases by a constrained element set is shown with a single dot for each constrained element set. See **Figure S18-20** for plots based on additional targets. **(D)** The graph shows the fold enrichment for Fetal Brain DHS^5^ within the non-exonic portion of each conservation state, separately for those bases in a PhastCons constrained element (pink) and bases not in such an element (blue). Enrichments within constrained elements varied substantially depending on the conservation state. For a given conservation state, bases in a constrained element had greater enrichments than bases not in a constrained element, illustrating complementary information of conservation states and constrained elements. See **Figure S21** for graphs based on different element sets or DHS data and **Figure S22** for these enrichments plotted against the size of the set.

We also compared the ability of conservation states to recover bases covered by DHS, both genome-wide and restricted to non-exonic bases, and repeated these analyses when also restricting to bases distal to a TSS (**Figures S19** and **S20, Methods**). When considering DHS bases in aggregate over 53 cell and tissue types both genome-wide and restricting to non-exonic bases, we found that at the same recall level the conservation states could identify bases in a DHS at greater precision than all constraint scores considered and PhastCons constrained elements. GERP++, SiPhy-pi and SiPhy-Omega elements did have higher precision at their single recall point (**Figure S19**). Similar results were seen when just considering distal regions, except for some of the scores in the non-exonic comparison at very low recall levels. The relative precision at the same recall levels between conservation states and the GERP++, SiPhy-pi and SiPhy-Omega constrained element sets did not hold for all cell types (**Figure S20**). The increase in precision for those constrained element sets in the aggregate evaluation over constraint scores, PhastCons elements, and ConsHMM annotations might be related to the coarser resolution at which they were defined (**Figure S17**). We note that the information about DHS in the conservation states was complementary to that in constrained element sets, as evidenced by the substantial variation in DHS enrichments of bases within constrained elements depending on their conservation state (**Figures 5D, S21** and **S22**). For example, bases in PhastCons constrained elements falling in 35 different states were depleted for Fetal Brain DHS in non-exonic regions, covering 10% of PhastCons bases, while bases in PhastCons elements in 12 other states were over 5-fold enriched, covering 37% of PhastCons bases. Additionally, we saw cases where certain states had greater enrichments for DHS for their bases not in a constrained element compared to bases in a constrained element in other states. On the other hand, constrained element calls offered additional information, as we observed that in most cases, for a given conservation state, bases that were in a constrained element call had greater enrichment for DHS than those that were not.

We also analyzed the enrichments of our conservation states for previously defined nine-subsets of PhastCons constrained non-exonic elements (CNEEs) based on a directed phylogenetic approach that assigned each element to a phylogenetic branch point of origin (**Figure S23A**).^22^ This demonstrated the heterogeneous nature of some of the resulting assignments when relying on directed phylogenetic partitioning approaches. For example, bases in elements assigned to originating at the branch point of the Tetrapod clade showed a high enrichment (37 fold) for state 2, as would be expected since state 2 is associated with aligning and matching through all vertebrates except fish, but an even greater enrichment (51 fold) for state 100, associated with putative artifacts (**Figure S23A**). We also evaluated whether within non-exonic regions any subset of CNEE assigned to a specific clade exhibits enrichments comparable to enrichments seen with the conservation states for CpG islands within non-exonic regions (**Figure S23C**). The most enriched subset of CNEE bases was only 6.7 fold enriched compared to the 37.6 fold enrichment observed for state 28 in non-exonic regions, and only covered 1.9% of non-exonic CpG island bases compared to 12.8% of such bases covered in state 28. A similar pattern of enrichments was observed when considering only the CNEEs overlapping a PhastCons element called on the same alignment as the conservation states (**Figure S23B** and **S23D**). These results highlight that the conservation states are able to capture additional biologically relevant information present in the alignment that is not captured by directed phylogenetic branch assignments of constrained elements.

### Bases prioritized by different variant prioritization scores have distinct conservation state enrichment patterns

In addition to scores defined based on just interspecies constraint, a variety of other scores have been proposed to prioritize variants, including some based on intra-species constraint or integrating inter-species constraint with other genomic annotations. A number of these scores are widely used, even though a systematic understanding of different types of bases prioritized by various scores is generally lacking. We leveraged the conservation state annotations to more systematically understand the bases prioritized by a variety of different scores in terms of their underlying pattern of conservation.

Specifically, we analyzed the conservation states’ genome-wide enrichments of bases prioritized by 12-different scores (CADD (v1.4), CDTS, DANN, Eigen, Eigen-PC, FATHMM-XF, FIRE, fitCons, GERP++, PhastCons, PhyloP, and REMM) to be in the top 1, 5, and 10% of the genome as well as the enrichment specifically in non-coding regions for those 12-scores and two additional ones defined only on non-coding regions, LINSIGHT and FunSeq2 (**Figures 6A, 6B** and **S24-S27**).^8,9,13,16,18,19,37,45,50–54^ We observed an overall strong enrichment for bases prioritized by most scores for a specific set of conservation states. For example, the top 1% CADD bases showed a 77.2 fold enrichment for state 1, amounting to 46% of the top 1% CADD bases falling in this state. This enrichment was greater than that observed for any interspecies constraint score, despite the CADD score being defined on a diverse set of genomic annotations, including many non-conservation based annotations. There was a general consistency in states with higher enrichment across the various measures. For example, when considering the top 1% bases for the genome-wide analysis, the set of states that were among the top five most enriched by at least one of the 12 scores contained only 13 states. Nine of these 13 states (states 1-5, 7, 28, 54, 100) were in the top five for at least three scores. However, there were important differences for the scores in the relative enrichment among these top states, and in several cases a single score prioritized other states.

**Figure 6:**
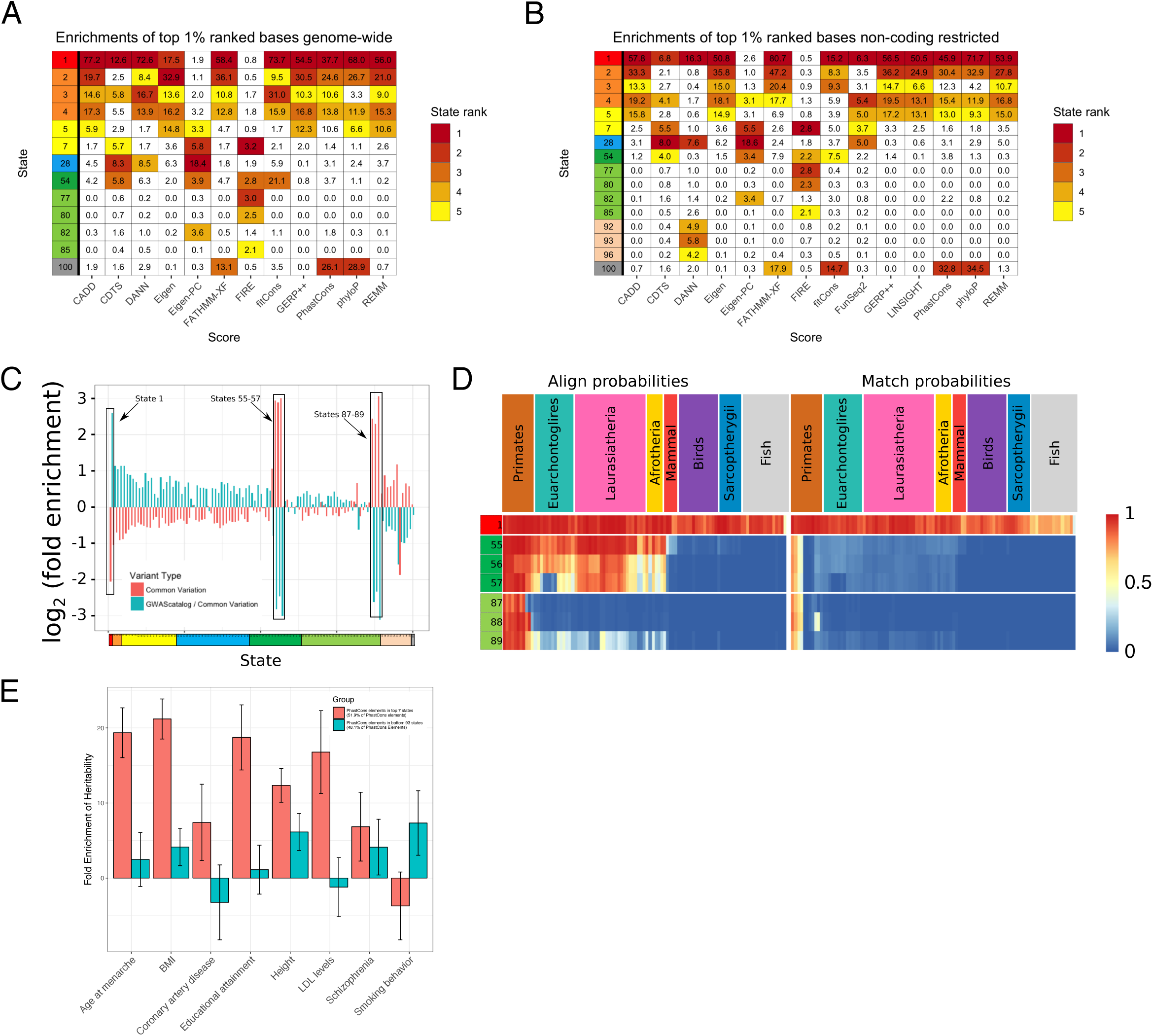
Conservation states and association with human genetic variation. **(A)** Fold enrichments of bases ranked in the top 1% genome-wide by 12 variant prioritization scores. Only states that were among the top five most enriched states for at least one score are shown. The enrichment of the top five ranking states for each score is colored according to the ranking and the color scale shown on right. **(B)** Enrichments of bases ranked in the top 1% of the non-coding genome by 14 variant prioritization scores. The criteria for selecting states to display and coloring enrichments was the same as in panel (A). Enrichments at additional thresholds and for all states both genome-wide and for the non-coding genome are in **Figure S24-S27**. The enrichments for CADD shown here are based on v1.4, while enrichments based on the original version of CADD are also shown in **Figure S25-S27**. (**C**) The panel displays the log_2_ fold enrichment of each state for common SNPs (pink) and GWAS catalog variants relative to common SNPs (blue). State 1, associated with high alignability and matching through all vertebrates, showed the greatest depletion of common SNPs and the highest enrichment for GWAS variants relative to common SNPs. States 55-57 and 87-89 exhibited the opposite pattern having the greatest enrichment for common SNPs and the greatest depletion of GWAS variants relative to this background. The second most depleted state for common SNPs, which did not show enrichment for GWAS catalog SNPs, was state 96 which captured large gaps in the assembly (**Figure S8**). (**D**) Panel shows the representation of state emission parameters from **Figure 2A** for the subset of states highlighted in panel (C). The states with the greatest depletion of GWAS variants all had relatively high alignability at least through primates, but low matching probabilities for almost all species except a few closely related primates. (**E**) Applying the heritability partitioning enrichment method of Finucane et al.^15^ on two disjoint subsets of bases in PhastCons elements, with eight phenotypes previously analyzed with heritability partitioning in the context of a baseline set of annotations (**Methods**).^15^ One set of bases are those in PhastCons elements that are also in one of the seven conservation states showing the greatest enrichment for DHS in its non-exonic portion (States 1-5, 8, and 28) covering 51.9% of PhastCons bases (pink). The other set are those bases in PhastCons elements overlapping the remaining 93 states covering 48.1% of PhastCons bases (blue).

One interesting result was the wide disagreement among the scores of the relative importance of state 2, the most enhancer enriched state, and state 28, the most promoter enriched state, particularly in non-coding regions. For example, when considering top 1% bases in non-coding regions, state 2 was the second or third most enriched state for CADD, Eigen, FATHMM-XF, GERP++, LINSIGHT, PhastCons, PhyloP, and REMM prioritized variants, with fold enrichments in the range of 24.9-47.2. On the other hand, state 28 was not one of the top five most enriched states for any of those scores and its enrichments ranged between 0.3-6.2. In contrast, for CDTS, DANN, and Eigen-PC state 2 only had enrichments between 0.8-2.1, while state 28 was the first or second most enriched state for each score, with enrichments ranging from 7.6-18.6. Also surprising was that DANN showed a depletion for state 2, which showed high matching through all vertebrates except fish, but enriched for states that were only associated with subsets of primates. For example, states 92, 93, and 96 did not have an alignment frequency greater than 0.05 for any species past Gibbon, but were among the top five states with the greatest enrichments for DANN prioritized variants, with enrichments in the range 4.2-5.8. None of the other 13 scores considered showed enrichment for these states. This is despite DANN using the same overall framework as CADD except using a deep neural network, and previously reporting to be better able to predict the simulated variants used to train CADD^51^. We verified that this difference with CADD also held for the original version of the CADD score that used the same features as DANN (**Figure S25**). FitCons and FunSeq2 had more balanced and relatively lower maximum enrichments for states 2 and 28. The FIRE score was an outlier in that the maximum enrichment it had for any conservation state was only 2.8. States for which FIRE prioritized bases showed the greatest enrichment included states such as 77, 80, and 85, which only showed substantial alignments among primates. The FIRE score was trained based on predicting expression quantitative trait loci (eQTL) in lymphoblastoid cell lines, which is a very different training objective than the other scores considered. It was previously noted that this led to background selection being the most important feature to this score.^50^

There were also strong disagreements about the relative importance of other states across scores. State 100, associated with likely alignment artifacts, was one such state. For example, at the top 1% threshold for the non-coding genome analysis, the state was among the most enriched states for FATHMM-XF, FitCons, PhastCons and PhyloP with enrichments in the range 14.7-34.5, while other scores showed more modest enrichments or even depletions, highlighting differences in the vulnerability of each score to likely alignment artifacts. State 54, which associated with high aligning through most vertebrates but not matching, was another state with wide disagreement among scores in the importance of those bases, particularly in the genome-wide analysis. At the top 1% threshold in the genome-wide analyses, this was the third most enriched state based on CDTS, Eigen-PC, FIRE, and fitCons, with the enrichment for fitCons reaching 21.1. In contrast, state 54 was depleted for the top GERP++ and REMM bases. The high enrichment of fitCons for these bases is expected, as the features it considers lack the resolution to differentiate the third codon position from the more conserved first and second codon positions when scoring coding regions. There were also differences in the relative importance given to state 1. For example, when considering variants genome-wide at the top 1% threshold, nine scores had the strongest enrichment for this state, but EIGEN, EIGEN-PC, and FIRE did not. EIGEN-PC did show the strongest enrichment for state 1 at other thresholds and EIGEN did when restricting to non-coding genome at all thresholds. However, their inconsistent ranking of state 1 is likely reflective of the unsupervised prioritization scheme used by these scores. Overall, these results show that the ConsHMM state annotations provides insights into key differences in variants prioritized by various scores by systematically and in an unbiased way capturing biologically diverse classes of nucleotides at single nucleotide resolution.

### Enrichment of conservation states for human genetic variation

Previous analyses have found a depletion of human genetic variation in evolutionarily constrained elements.^7^ Consistent with that, the greatest depletion (3.3 fold depletion) of common SNPs from dbSNP is in state 1, the state most enriched for constrained elements. Interestingly, six states, A_SMam states 55-57 and AM_Prim states 87-89, had enrichments in the range 5 to 8 fold for common SNPs. These were also the six states with greatest enrichment of CG dinucleotides (**Figure S12**). These six states have in common that they show high align probabilities for most primates, but low match probabilities for some of those same primates. These states are thus associated with substantial variation both among primates and among humans. We observed similar patterns of enrichment for variants identified from whole genome sequencing (WGS) of a cohort of 7784 unrelated individuals^37^, with the levels of state enrichments and depletions increasing with the minor allele frequency (**Figure S28**).

When analyzing the enrichment of GWAS catalog variants^36^ relative to the background of common SNPs we saw opposite enrichment patterns for these states (**Figure 6C** and **6D**). For example, relative to this background, state 1 was most enriched for GWAS catalog variants, which is consistent with previous observations of constrained elements enriching for GWAS variants.^7^ On the other hand, states 55-57 and 87-89 showed the greatest depletion. These results suggest that common variants are less likely to be phenotypically significant if they fall in conservation states most enriched for common genetic variation.

### Constrained element enrichment for partitioned heritability of complex traits depends on conservation state

Previous analyses have suggested a strong enrichment of constrained elements and DHS for phenotype heritability.^15,55^ As we saw large differences in DHS enrichments of constrained elements depending on the conservation state, we investigated the extent to which constrained elements in conservation states most enriched for DHS enriched for phenotype heritability compared to the remaining states. Specifically, we ranked the conservation states in descending order of their median enrichment for DHS from a compendium of 123 experiments from the ENCODE consortium, within the non-exonic portion of the state (**Figure 2B, Methods**).^3^ We then partitioned bases in PhastCons constrained elements into two almost equal size sets based on whether they overlapped one of the top seven ranked conservation states (states 1-5, 8, 28) or not. We then computed the heritability for these two sets for eight phenotypes in the context of a set of baseline annotations that include DHS annotations (**Methods**).^15^ For seven of the phenotypes, we found that bases in constrained elements overlapping the top seven states had greater enrichment than those in the remaining 93 states, often substantially so (**Figure 6E**). These results suggest additional value in the conservation state annotations for isolating more likely disease relevant variants.

## Discussion

We presented a framework for genome annotation based on comparative genomics sequence data. Our approach learns a set of conservation states *de novo* using a multivariate HMM based on the combinatorial and spatial patterns of which species align and match a reference genome in a multi-species DNA sequence alignment. We applied this approach to annotate the human genome at single nucleotide resolution into one of 100 conservation states. Conservation state annotations exhibited substantial enrichments for a wide range of other genomic annotations that were not provided to the model in training, thus supporting their biological significance. Specific conservation states exhibited strong enrichments for various gene annotations including exons, TSS and TES of genes, while others showed strong enrichments for specific types of repeat elements. Conservation states showed differential enrichment patterns for various classes of genes and DHS from multiple cell types, even though they were defined independently of any functional genomics data. Specific conservation states exhibited enrichments for common human variants, while a different set of states exhibited enrichments for variants identified by GWAS relative to common variation.

ConsHMM differs from other comparative genomics based annotation approaches in several respects. One difference is that it takes an unsupervised approach that does not explicitly use a phylogenetic tree, except to the extent to which a phylogenetic tree was used to generate the input multi-species sequence alignment. This leads to relatively unbiased, simple and interpretable models. However, many state patterns discovered are consistent with expected observations from commonly assumed phylogenetic relationships of the species. While states’ parameters often decreased with divergence time from human, there were a number of exceptions. Some of these exceptions corresponded to missing specific sub-clades of species, particularly those with long branch lengths. For example, a number of states were not represented by mouse and rat, while being represented by more distally diverged mammals. Other exceptions isolated putative artifacts in the alignments that might otherwise confound analyses, as we saw for two states heavily enriched for pseudogenes. ConsHMM also differs from other commonly used modeling approaches in how it explicitly differentiates non-aligning bases from aligning non-matching bases, which allowed it, for example, to identify states particularly associated with third codon positions. Another difference between the ConsHMM annotations and standard constraint measures is that the ConsHMM annotations are defined directly relative to the variant present in the genome being annotated. When applying ConsHMM to annotate the human genome, a mutation unique to human would be expected to have a much larger effect on the ConsHMM annotations than a mutation unique to a single other species. This would not in general be expected to be the case for constraint measures that do not differentiate the target genome for annotation from other genomes in an alignment. An interesting future direction would be to produce and analyze individual specific ConsHMM annotations.

Our conservation state annotation is complementary to existing binary calls and scores of evolutionary constraint based on phylogenetic modeling. Both locations called as constrained and those called as non-constrained are heterogeneous in their assigned conservation state. Our annotations thus provide additional descriptive information about the conservation patterns at each base. In terms of information for predicting external annotations, we found that in many cases the conservation states had greater information than constraint scores or elements. Notably, our modeling approach identified a conservation state, state 28, associated with a pattern of aligning and matching some mammalian and non-mammalian vertebrates, but not with high probability for any one species. This conservation state strongly enriched for transcription start sites and CpG islands, and was not well captured by phylogenetic modeling approaches. For other cases, such as DHS, the relative information depended on the constrained element set or score being compared. Importantly, we observed that DHS information provided by the states was complementary to information in the constrained element calls. We also used the conservation state annotations to systematically understand key similarities and differences in the patterns of conservation in bases prioritized by a large number of different variant prioritization scores, including scores based on integrating diverse features (**Figure 6A** and **6B**). The conservation state annotations provide a powerful framework for gaining such an understanding, since the corresponding conservation patterns are defined systematically in an unbiased way, at single nucleotide resolution and capture a diverse set of biological features. Furthermore, we observed that bases in constrained elements showed substantially different enrichments for phenotype-associated heritability, depending on their conservation state.

The conservation states are both inspired by, and provide complementary information to, existing chromatin state annotation approaches. While the states from the two approaches are based on very different data and have fundamental differences, they also exhibited substantial cross-enrichments. In general, conservation states have the advantage of providing information at single nucleotide resolution, which we demonstrated by showing enrichments patterns in and around coding exons and regulatory motifs. Conservation states can also provide information about bases in the genome even if the relevant cell type has not been experimentally profiled, while chromatin states have the advantage of directly providing cell type specific information.

We expect many applications for the methodology and annotations we have presented here. While we applied ConsHMM here to one multiple species alignment, a 100-way Multiz human alignment, the methodology is general, and thus can be readily applied to alignments to other species or alignments generated by other methods.^26^ The annotations we produced serve as a resource to directly interpret other genomic datasets or variant prioritization scores. They could also potentially be integrated with other complementary genomic annotations in methods that produce integrated variant prioritization scores. This work represents a step in the direction of improving whole genome annotations, which will continue to be of increasing importance towards understanding health and disease as the availability of whole genome sequencing data increases.

## Supporting information

## Supplemental Data

Supplemental Data include twenty-eight figures and three tables.

## Acknowledgements

We thank Ewan Birney and members of the Ernst lab for useful discussions. We acknowledge funding from US National Institutes of Health grants DP1DA044371, R01ES024995, U01HG007912 and U01MH105578 (J.E.), and T32CA201160 (A.S.), US National Science Foundation CAREER Award #1254200 (J.E.), a Kure-IT award and an Alfred P. Sloan Fellowship (J.E.).

## Web Resources

25-state chromatin state annotations:

http://compbio.mit.edu/roadmapCADD score v1.0:

http://krishna.gs.washington.edu/download/CADD/v1.0/whole_genome_SNVs.tsv.gzCADD score v1.4:

http://krishna.gs.washington.edu/download/CADD/v1.4/GRCh37/whole_genome_SNVs.tsv.gz

CDTS score:

http://www.hli-opendata.com/noncoding/coord_CDTS_percentile_N7794unrelated.txt.gz

http://www.hli-opendata.com/noncoding/SNVusedForCDTScomputation_N7794unrelated_allelicFrequency0.001truncated.txt.gz

CNEEs from Lowe et al.^22^:

http://www.stanford.edu/∼lowec/data/threePeriods/hg19cnee.bed.gz

ConsHMM v1.0 software and ConsHMM state annotations:

https://github.com/ernstlab/ConsHMMDANN score:

https://cbcl.ics.uci.edu/public_data/DANN/data/EIGENandEigen-PC score:

https://xioniti01.u.hpc.mssm.edu/v1.1/

ENCODE DHS:

http://hgdownload.cse.ucsc.edu/goldenPath/hg19/encodeDCC/wgEncodeUwDnase/

FATHMM-XF score:

http://fathmm.biocompute.org.uk/fathmm-xf/

<u>FIRE score:</u>

https://sites.google.com/site/fireregulatoryvariation/

<u>fitCons score:</u>

http://compgen.cshl.edu/fitCons/0downloads/tracks/i6/scores/

<u>FunSeq2 score:</u>

http://org.gersteinlab.funseq.s3-website-us-east-1.amazonaws.com/funseq2.1.2/hg19_NCscore_funseq216.tsv.bgz

GENCODE v19:

https://www.gencodegenes.org/releases/19.html

GERP++ scores and constrained element calls: http://mendel.stanford.edu/SidowLab/downloads/gerp/

GWAS catalog variants:

<u>LINSIGHT score:</u>

http://compgen.cshl.edu/∼yihuang/tracks/LINSIGHT.bw

Motif instances and background: http://compbio.mit.edu/encode-motifs/

https://www.ebi.ac.uk/gwas/

Multiz 100-way alignment to hg19 reference:

http://hgdownload.soe.ucsc.edu/goldenPath/hg19/multiz100way/

<u>REMM score:</u>

https://zenodo.org/record/1197579/files/ReMM.v0.3.1.tsv.gz

Roadmap Epigenomics DHS: http://egg2.wustl.edu/roadmap/data/byFileType/peaks/consolidated/narrowPeak/

SiPhy-omega and SiPhy-pi constrained element calls (hg19 liftOver): https://www.broadinstitute.org/mammals-models/29-mammals-project-supplementary-info

