## Supplementary material for "Systematic Discovery of Conservation States for Single-Nucleotide Annotation of the Human Genome"

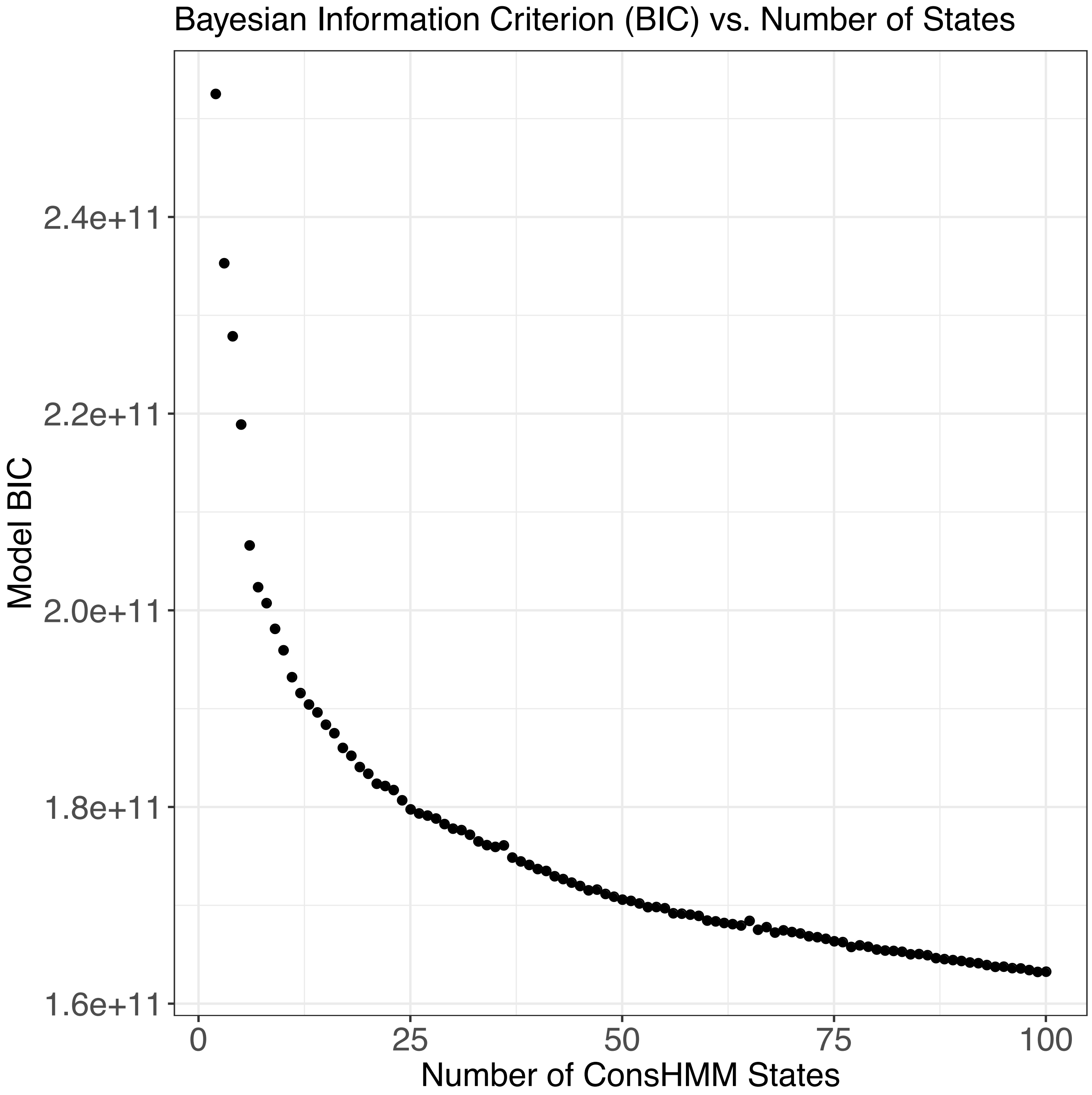


**Figure S1: BIC as a function of number of states in the model.** The BIC criterion computed for models with each number of states from 2 to 100. For this criterion lower values correspond to preferred models.

| State | Average posterior probability of state at positions assigned to state |
|---|---|
| 1 | 1.00 |
| 2 | 0.99 |
| 3 | 0.99 |
| 4 | 0.98 |
| 5 | 0.96 |
| 6 | 0.96 |
| 7 | 0.97 |
| 8 | 0.92 |
| 9 | 0.93 |
| 10 | 0.94 |
| 11 | 0.98 |
| 12 | 0.97 |
| 13 | 0.97 |
| 14 | 0.95 |
| 15 | 0.92 |
| 16 | 0.92 |
| 17 | 0.95 |
| 18 | 0.94 |
| 19 | 0.97 |
| 20 | 0.98 |
| 21 | 0.98 |
| 22 | 0.96 |
| 23 | 0.97 |
| 24 | 0.98 |
| 25 | 0.97 |
| 26 | 0.97 |
| 27 | 0.96 |
| 28 | 0.98 |
| 29 | 0.97 |
| 30 | 0.98 |
| 31 | 0.97 |
| 32 | 0.97 |
| 33 | 0.98 |
| 34 | 0.98 |
| 35 | 0.97 |
| 36 | 0.96 |
| 37 | 0.97 |
| 38 | 0.97 |
| 39 | 0.97 |
| 40 | 0.97 |
| 41 | 0.96 |
| 42 | 0.95 |
| 43 | 0.97 |
| 44 | 0.96 |
| 45 | 0.97 |
| 46 | 0.97 |
| 47 | 0.93 |
| 48 | 0.95 |
| 49 | 0.96 |
| 50 | 0.96 |
| 51 | 0.95 |
| 52 | 0.93 |
| 53 | 0.95 |
| 54 | 0.98 |
| 55 | 0.97 |
| 56 | 0.96 |
| 57 | 0.96 |
| 58 | 0.96 |
| 59 | 0.95 |
| 60 | 0.95 |
| 61 | 0.96 |
| 62 | 0.96 |
| 63 | 0.96 |
| 64 | 0.97 |
| 65 | 0.98 |
| 66 | 0.99 |
| 67 | 0.98 |
| 68 | 0.98 |
| 69 | 0.97 |
| 70 | 0.95 |
| 71 | 0.94 |
| 72 | 0.96 |
| 73 | 0.98 |
| 74 | 0.99 |
| 75 | 0.99 |
| 76 | 0.99 |
| 77 | 1.00 |
| 78 | 1.00 |
| 79 | 0.99 |
| 80 | 0.98 |
| 81 | 0.98 |
| 82 | 0.99 |
| 83 | 0.99 |
| 84 | 0.99 |
| 85 | 0.99 |
| 86 | 1.00 |
| 87 | 0.97 |
| 88 | 0.98 |
| 89 | 0.95 |
| 90 | 0.99 |
| 91 | 0.98 |
| 92 | 1.00 |
| 93 | 1.00 |
| 94 | 1.00 |
| 95 | 0.99 |
| 96 | 1.00 |
| 97 | 1.00 |
| 98 | 1.00 |
| 99 | 1.00 |
| 100 | 1.00 |

**Figure S2: Average posterior probability of ConsHMM state assignments**. The value listed for each state is the average posterior probability of that state for all bases in the genome assigned to that state.

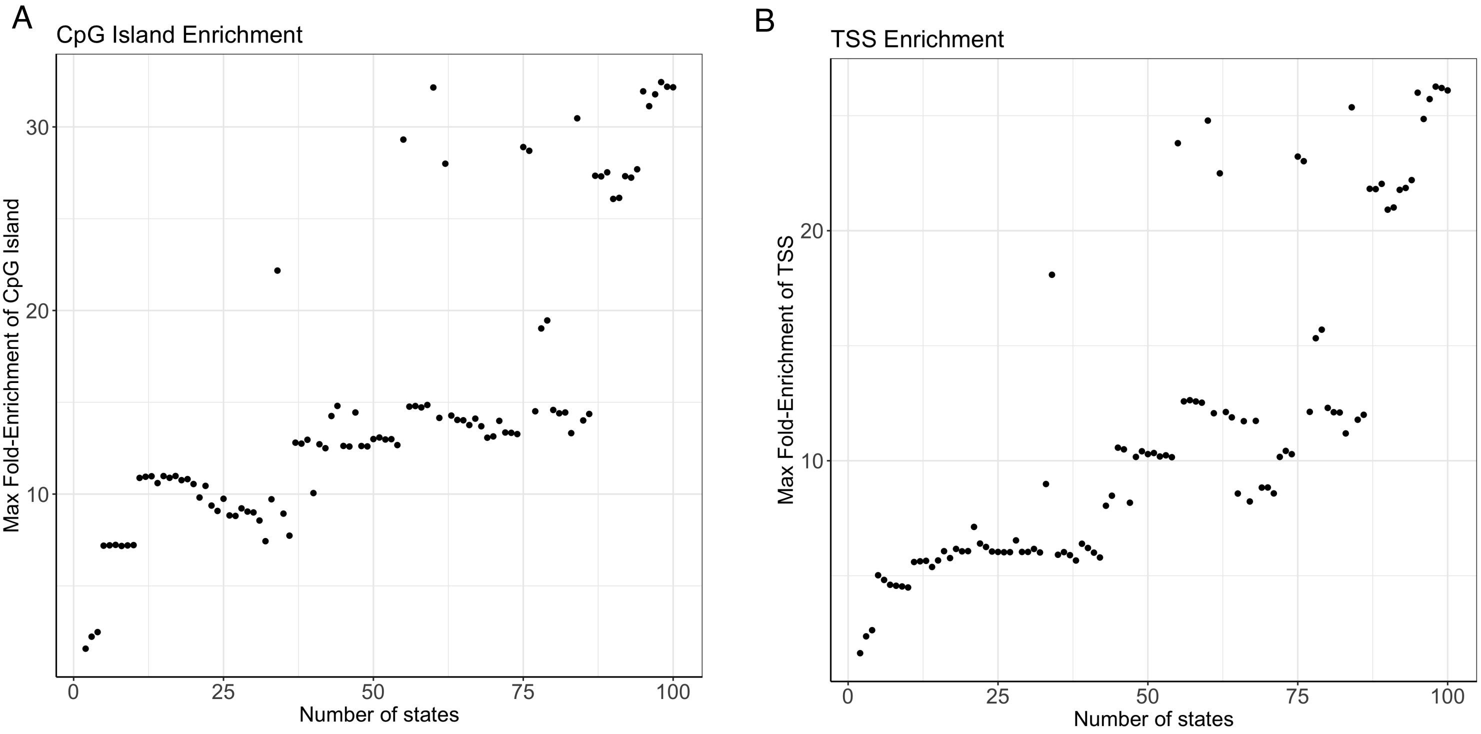


**Figure S3: Maximum CpG and TSS state enrichments as a function of number of states in the model.** The figures show the maximum fold enrichment for **(A)** CpG islands and **(B)** TSS of any state in a model as a function of the number of states in the model. The figure shows that states with substantially higher enrichment for these annotations are only found consistently in models with a large number of states. There were isolated cases of models with a moderate number of states also exhibiting high enrichment. However, since similar enrichment levels were not captured in models with similar numbers of states this suggest the possibility that other biologically relevant states might be missing from these models.

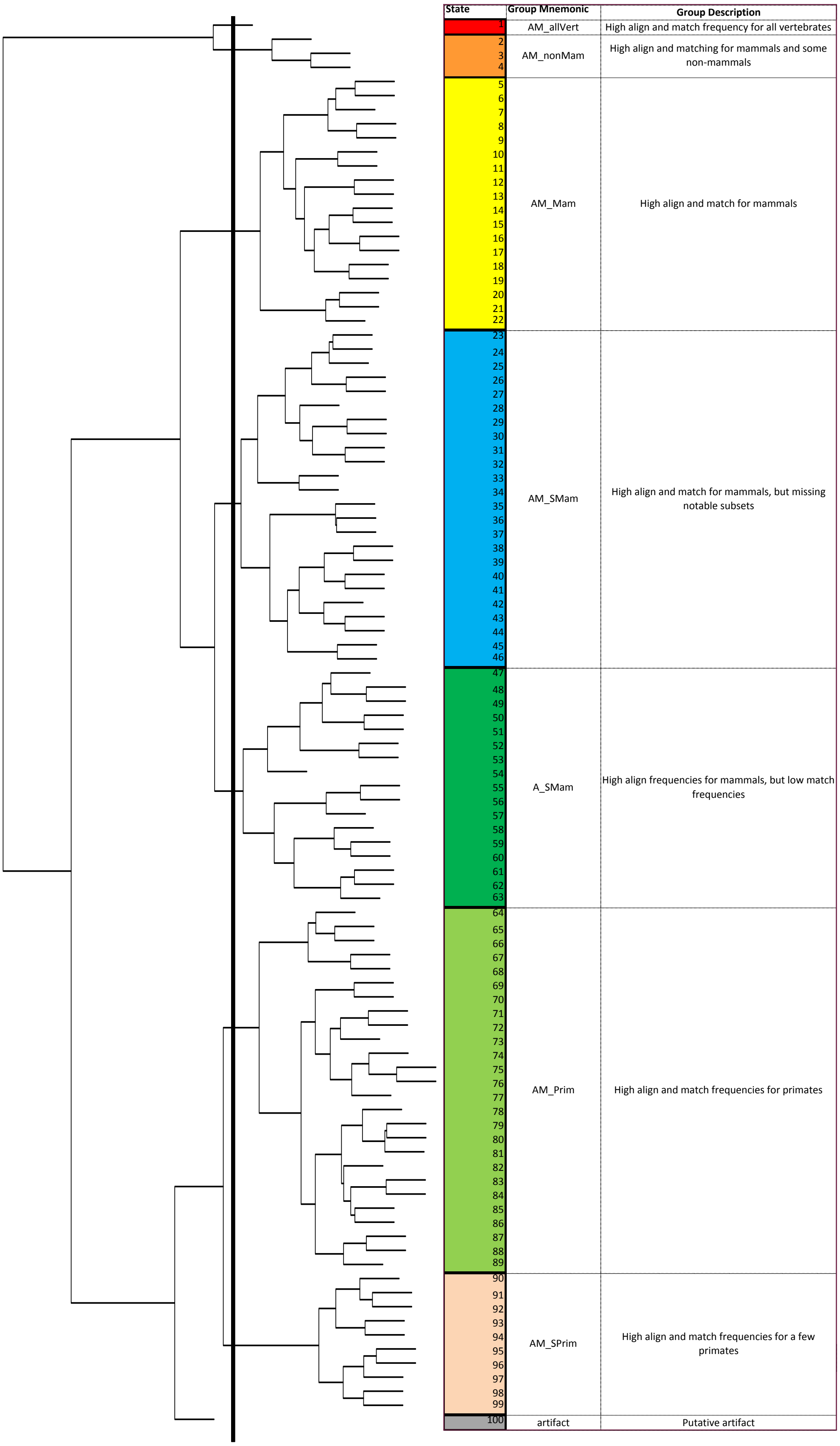


**Figure S4: Hierarchical clustering and grouping of conservation states.** The dendrogram on left displays a hierarchical clustering of the states based on the values in **Figure 2A**, with the leaves ordered based on optimal leaf ordering.[41] The thick black line indicates where the dendrogram was cut to form the eight major groups of conservation states, each receiving a different color shown on right. To the right of the state numbers are state group abbreviations from **Figure 2A**. To the right of the state labels is a high level description of the general patterns of the parameters of the state groups. Notable enrichments associated with specific states are summarized in **Table S3**.

A

### Align Probabilities

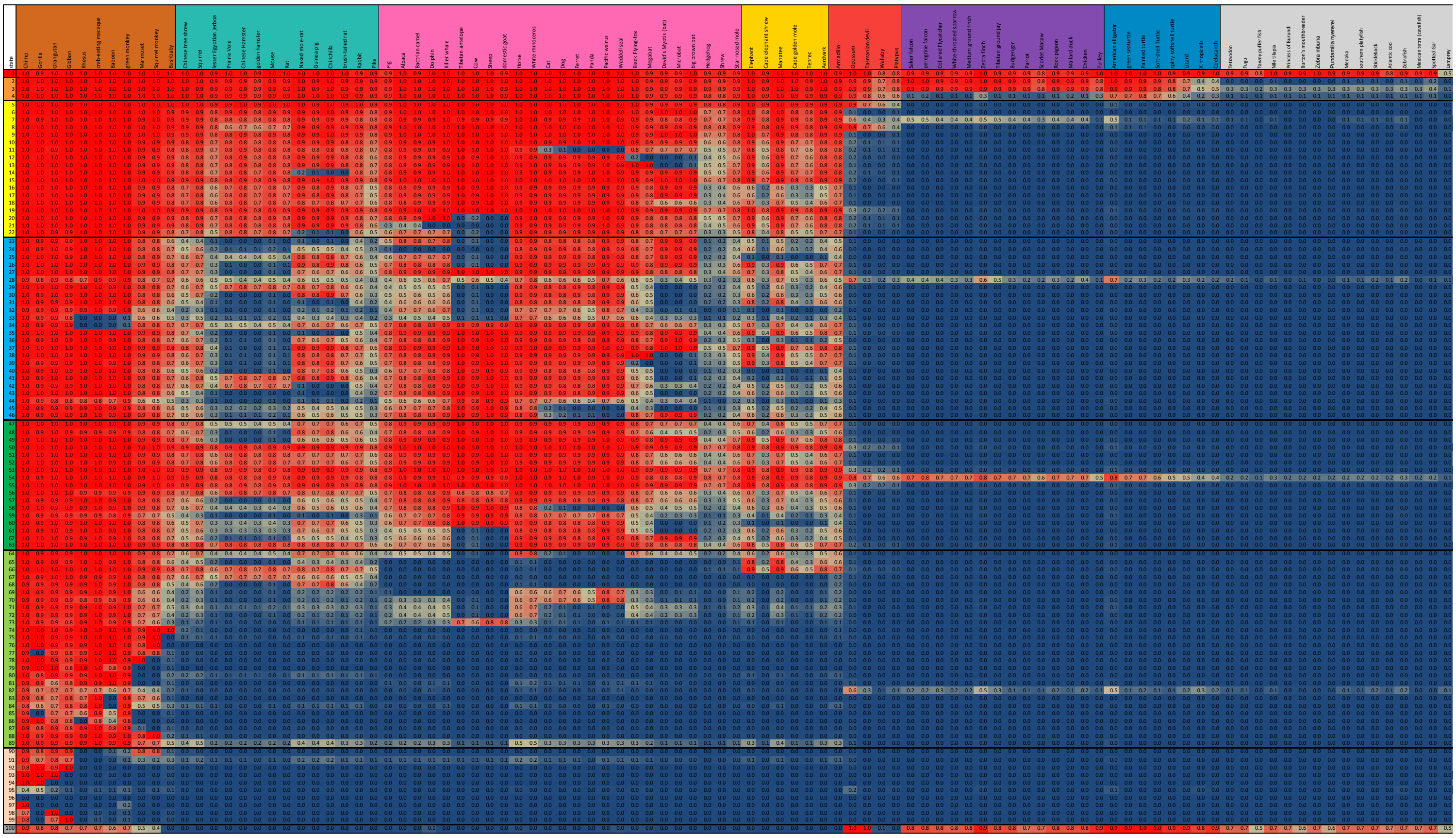

B

### Match Probabilities

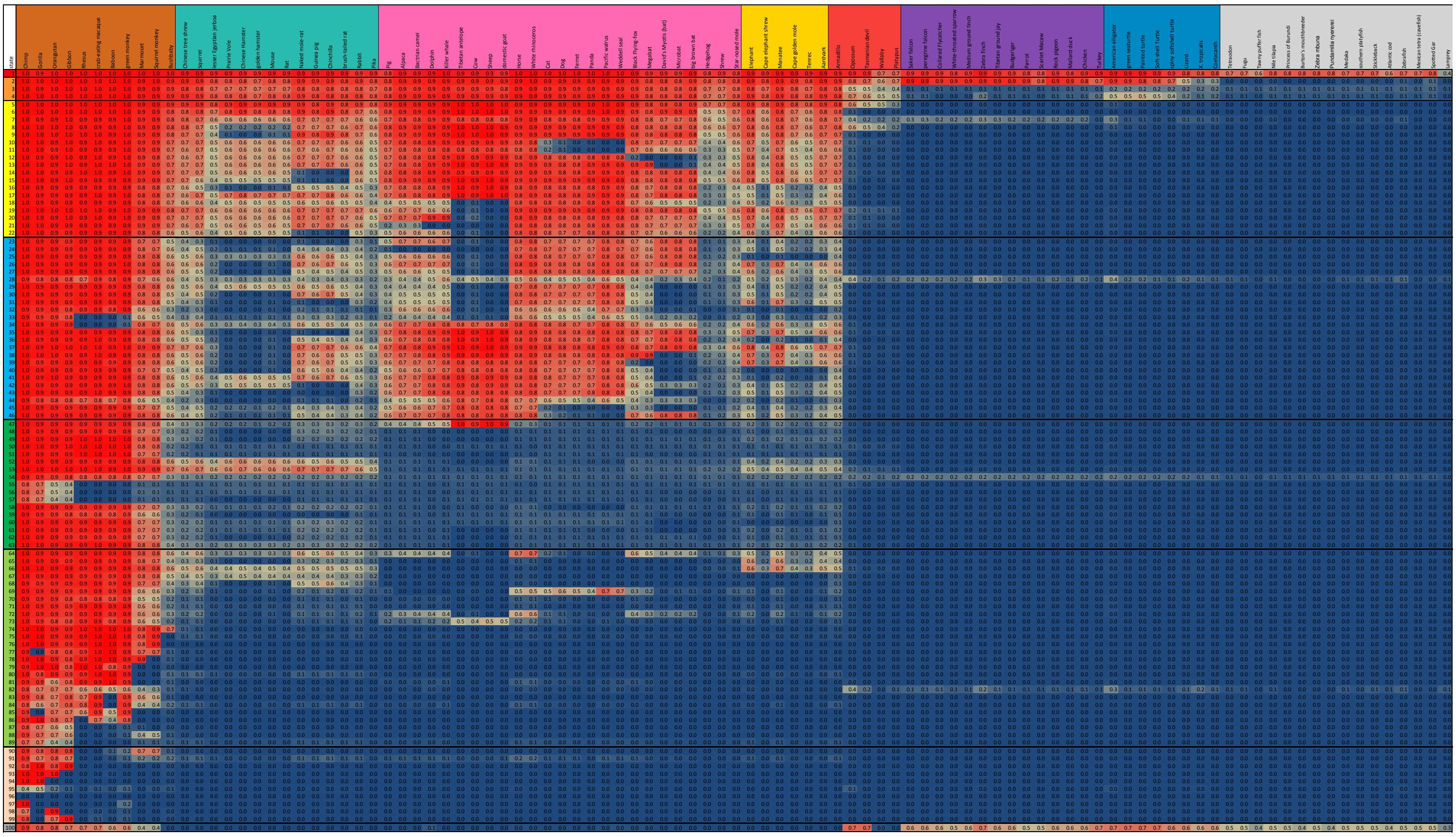

**Figure S5: Representation of the emission parameters.** This is a more detailed view of the representation of the emission parameters shown in the heatmap in **Figure 2A**. In this figure the actual probabilities values and the individual species names are also displayed. The values in this figure are also available in **Table S1**.

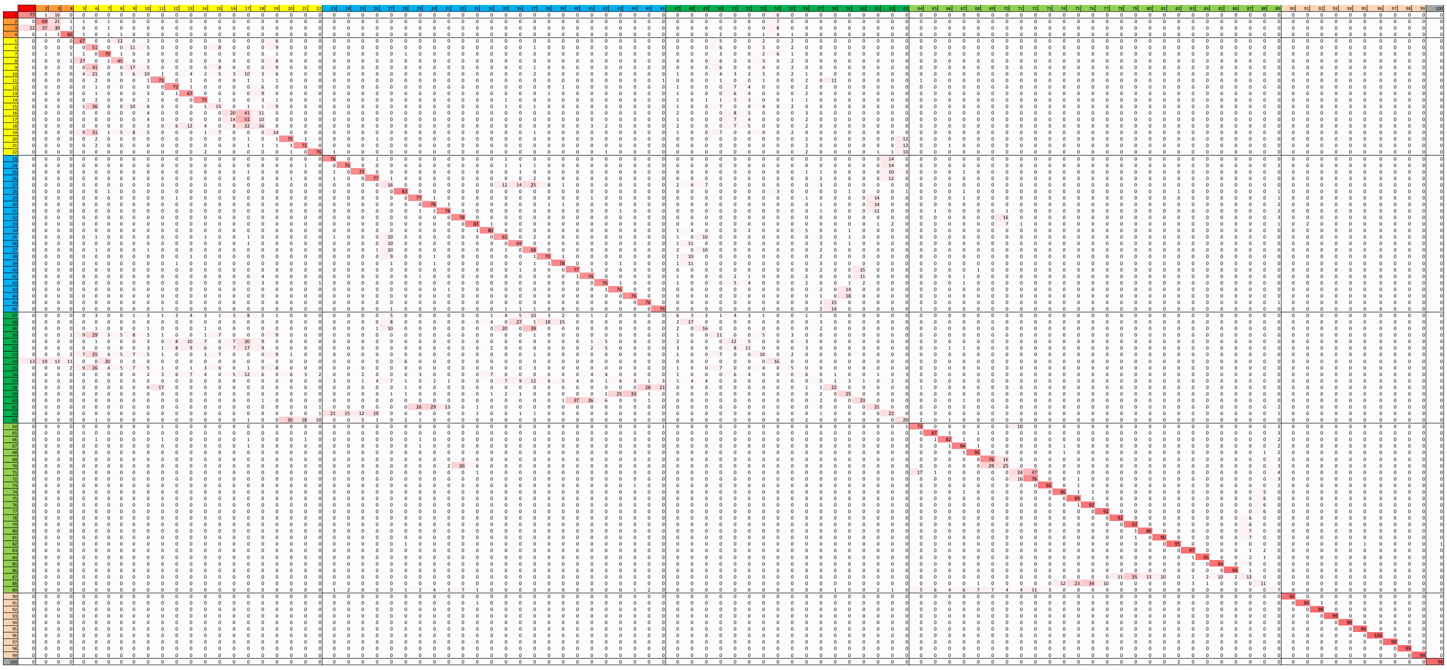

**Figure S6: Conservation state model transition probabilities.** The figure displays the conservation state model transition probabilities. The values correspond to the probability when in the state of the row to transition to the state of the column at the next base. Probabilities are displayed multiplied by 100, rounded to the nearest integer and shaded based on their value, with darker red corresponding to greater transition probabilities. Transition probabilities along the diagonal show the probability of remaining in the state at a neighboring position, which were often the highest values for some states. For states associated with low matching probabilities relative to the alignment probabilities such as the A_SMam subgroup (states 47-63) the probability of remaining in the same state was low. Transition probabilities to stay in the same state were highest in some states only showing substantial alignability at most within primates, thus the model can use spatial information through these transition probabilities to better differentiate instances of states with relatively similar emission probabilities.

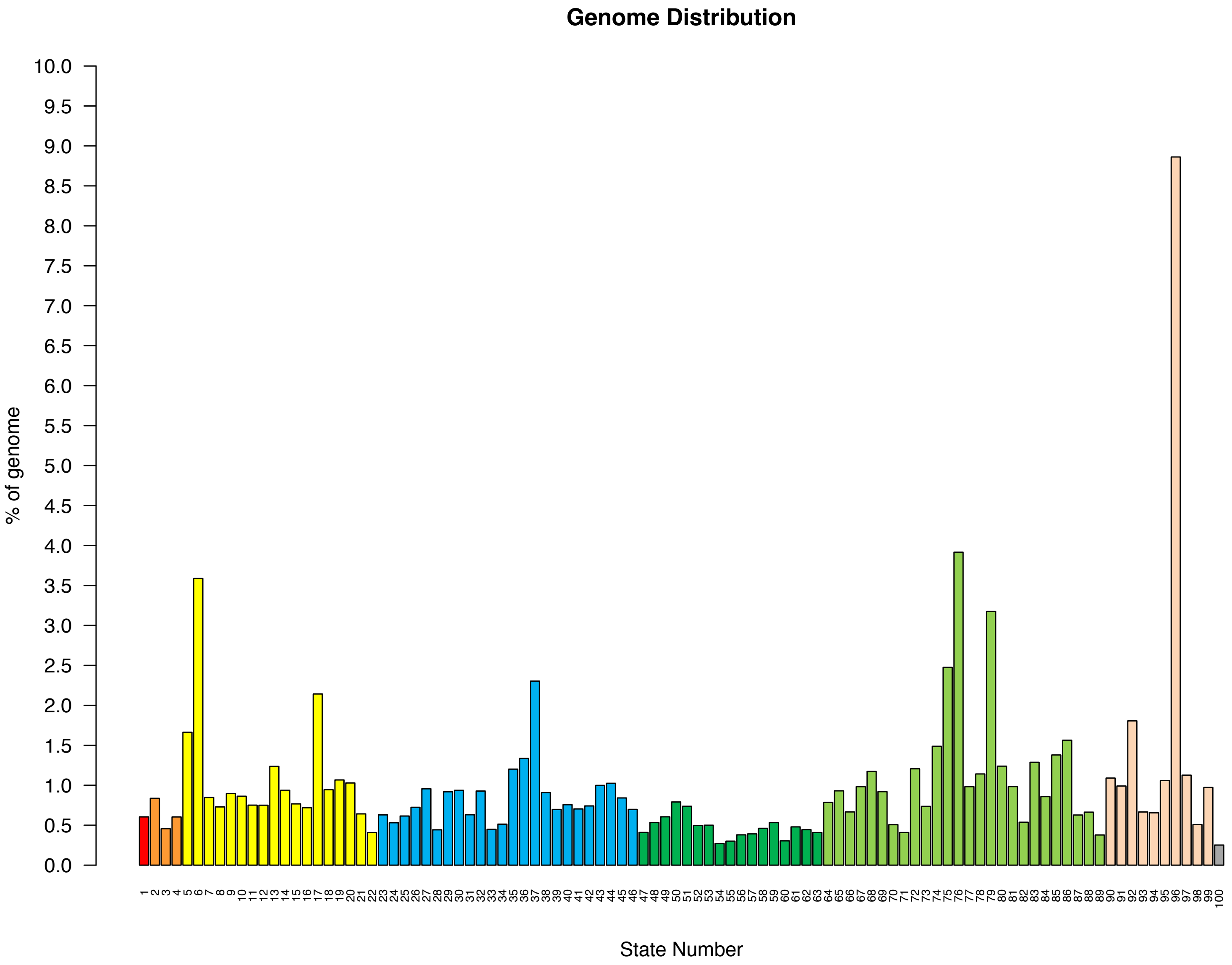


**Figure S7: Distribution of the genome in each state.** The graph displays the percent of the genome assigned to each conservation state. The median state coverage was 0.76% of the genome. All states except state 96 were in the range of 0.25% to 3.92% of the genome. State 96 was the largest state covering 8.86% of the genome and was associated with assembly gaps (**Figure S8**).

| | Genome % | TSS of protein coding genes | TSS of pseudogenes | TES of protein coding genes | TES of pseudogenes | CDS | Exons of protein coding genes | Exons of pseudogenes | UTR | CpG Island | PhastCons elements | Repeat Elements | Median Non-Exonic DHS | Assembly Gaps |
|---|---|---|---|---|---|---|---|---|---|---|---|---|---|---|
| 1 | 0.60 | 1.78 | 2.37 | 1.98 | 2.06 | 71.49 | 28.14 | 7.36 | 12.34 | 12.54 | 17.94 | 0.01 | 2.84 | 0.00 |
| 2 | 0.84 | 2.37 | 0.77 | 7.00 | 0.61 | 9.15 | 7.85 | 1.23 | 9.00 | 1.53 | 14.19 | 0.02 | 3.07 | 0.00 |
| 3 | 0.46 | 1.85 | 0.80 | 5.10 | 0.94 | 30.08 | 14.81 | 4.70 | 9.98 | 5.45 | 9.39 | 0.02 | 2.62 | 0.00 |
| 4 | 0.60 | 6.88 | 1.95 | 5.30 | 1.79 | 16.20 | 8.98 | 2.92 | 7.06 | 5.46 | 10.45 | 0.03 | 3.97 | 0.00 |
| 5 | 1.66 | 5.99 | 0.30 | 5.93 | 0.23 | 1.83 | 3.28 | 0.27 | 4.54 | 2.75 | 9.08 | 0.06 | 4.09 | 0.00 |
| 6 | 3.59 | 1.73 | 0.11 | 2.19 | 0.10 | 0.25 | 1.19 | 0.09 | 1.62 | 0.94 | 2.92 | 0.36 | 2.49 | 0.00 |
| 7 | 0.85 | 6.76 | 0.91 | 6.72 | 0.96 | 2.30 | 4.76 | 0.69 | 5.39 | 7.10 | 2.09 | 0.10 | 2.14 | 0.00 |
| 8 | 0.73 | 3.30 | 0.26 | 4.55 | 0.36 | 1.19 | 2.88 | 0.34 | 3.84 | 2.16 | 2.75 | 0.05 | 2.83 | 0.00 |
| 9 | 0.90 | 1.28 | 0.06 | 1.95 | 0.11 | 0.19 | 1.14 | 0.08 | 1.53 | 0.91 | 1.01 | 0.37 | 2.13 | 0.00 |
| 10 | 0.86 | 1.72 | 0.08 | 2.25 | 0.14 | 0.30 | 1.30 | 0.12 | 1.69 | 1.11 | 0.65 | 0.40 | 2.02 | 0.00 |
| 11 | 0.75 | 1.23 | 0.37 | 1.20 | 0.31 | 0.21 | 1.05 | 0.31 | 1.29 | 0.93 | 0.64 | 0.49 | 1.50 | 0.00 |
| 12 | 0.75 | 0.91 | 0.22 | 1.00 | 0.20 | 0.15 | 0.73 | 0.25 | 0.92 | 0.73 | 0.84 | 0.46 | 1.42 | 0.00 |
| 13 | 1.24 | 1.03 | 0.11 | 1.03 | 0.06 | 0.13 | 0.74 | 0.09 | 0.94 | 0.83 | 1.05 | 0.46 | 1.57 | 0.00 |
| 14 | 0.94 | 1.65 | 0.20 | 1.45 | 0.11 | 0.19 | 1.22 | 0.15 | 1.57 | 1.43 | 0.94 | 0.43 | 2.16 | 0.00 |
| 15 | 0.77 | 1.75 | 0.13 | 1.99 | 0.09 | 0.24 | 1.30 | 0.10 | 1.72 | 1.25 | 0.61 | 0.36 | 2.20 | 0.00 |
| 16 | 0.72 | 0.77 | 0.15 | 0.83 | 0.09 | 0.09 | 0.61 | 0.14 | 0.71 | 0.60 | 0.19 | 0.64 | 1.28 | 0.00 |
| 17 | 2.14 | 0.62 | 0.21 | 0.83 | 0.15 | 0.09 | 0.58 | 0.15 | 0.69 | 0.44 | 0.77 | 0.64 | 1.28 | 0.00 |
| 18 | 0.94 | 0.71 | 0.17 | 0.89 | 0.14 | 0.11 | 0.69 | 0.14 | 0.82 | 0.70 | 0.18 | 0.59 | 1.41 | 0.00 |
| 19 | 1.07 | 2.24 | 0.14 | 2.88 | 0.08 | 0.54 | 1.71 | 0.15 | 2.26 | 1.35 | 1.22 | 0.29 | 2.42 | 0.00 |
| 20 | 1.03 | 3.31 | 0.16 | 1.14 | 0.14 | 0.26 | 1.12 | 0.16 | 1.48 | 3.29 | 1.00 | 0.48 | 1.73 | 0.00 |
| 21 | 0.64 | 1.00 | 0.22 | 1.21 | 0.17 | 0.22 | 1.05 | 0.16 | 1.30 | 0.96 | 0.72 | 0.46 | 1.51 | 0.00 |
| 22 | 0.41 | 1.79 | 0.21 | 0.75 | 0.23 | 0.15 | 0.82 | 0.19 | 1.00 | 2.25 | 0.40 | 0.63 | 1.47 | 0.00 |
| 23 | 0.63 | 0.48 | 0.31 | 0.68 | 0.22 | 0.07 | 0.45 | 0.18 | 0.51 | 0.66 | 0.37 | 0.92 | 1.00 | 0.00 |
| 24 | 0.53 | 0.45 | 0.10 | 0.58 | 0.14 | 0.06 | 0.42 | 0.19 | 0.48 | 0.40 | 0.28 | 0.83 | 0.96 | 0.00 |
| 25 | 0.61 | 0.49 | 0.20 | 0.46 | 0.11 | 0.05 | 0.36 | 0.12 | 0.39 | 0.40 | 0.31 | 1.01 | 0.91 | 0.00 |
| 26 | 0.72 | 0.63 | 0.14 | 0.67 | 0.14 | 0.06 | 0.49 | 0.18 | 0.57 | 0.55 | 0.44 | 0.68 | 1.06 | 0.00 |
| 27 | 0.96 | 0.71 | 0.09 | 0.96 | 0.13 | 0.09 | 0.67 | 0.15 | 0.79 | 0.59 | 0.20 | 0.62 | 1.34 | 0.00 |
| 28 | 0.44 | 29.03 | 14.52 | 5.03 | 14.52 | 7.34 | 6.79 | 14.19 | 6.59 | 32.16 | 3.27 | 0.20 | 4.13 | 0.00 |
| 29 | 0.92 | 0.63 | 0.20 | 0.39 | 0.13 | 0.05 | 0.35 | 0.12 | 0.40 | 0.55 | 0.28 | 0.81 | 0.91 | 0.00 |
| 30 | 0.94 | 0.21 | 0.20 | 0.34 | 0.09 | 0.03 | 0.26 | 0.12 | 0.30 | 0.19 | 0.29 | 0.86 | 0.79 | 0.00 |
| 31 | 0.63 | 0.51 | 0.21 | 0.55 | 0.17 | 0.08 | 0.37 | 0.22 | 0.42 | 0.62 | 0.31 | 0.80 | 0.90 | 0.00 |
| 32 | 0.93 | 0.37 | 0.24 | 0.43 | 0.24 | 0.06 | 0.28 | 0.26 | 0.33 | 0.48 | 0.36 | 1.33 | 0.67 | 0.00 |
| 33 | 0.45 | 0.92 | 0.83 | 0.52 | 0.60 | 0.09 | 0.29 | 0.63 | 0.33 | 0.78 | 0.41 | 1.05 | 0.63 | 0.00 |
| 34 | 0.51 | 0.71 | 0.56 | 0.65 | 0.67 | 0.13 | 0.48 | 0.57 | 0.55 | 0.50 | 0.65 | 0.62 | 0.88 | 0.00 |
| 35 | 1.20 | 0.99 | 0.26 | 1.22 | 0.15 | 0.14 | 0.92 | 0.19 | 1.11 | 1.01 | 0.64 | 0.56 | 1.55 | 0.00 |
| 36 | 1.34 | 0.34 | 0.19 | 0.58 | 0.13 | 0.04 | 0.35 | 0.13 | 0.40 | 0.20 | 0.47 | 0.89 | 0.95 | 0.00 |
| 37 | 2.30 | 0.61 | 0.13 | 1.11 | 0.11 | 0.09 | 0.72 | 0.12 | 0.89 | 0.47 | 0.89 | 0.51 | 1.46 | 0.00 |
| 38 | 0.91 | 0.58 | 0.15 | 0.60 | 0.13 | 0.08 | 0.47 | 0.11 | 0.57 | 0.36 | 0.56 | 0.63 | 1.12 | 0.00 |
| 39 | 0.70 | 0.41 | 0.24 | 0.58 | 0.31 | 0.08 | 0.40 | 0.27 | 0.47 | 0.29 | 0.45 | 0.65 | 0.96 | 0.00 |
| 40 | 0.76 | 0.31 | 0.16 | 0.27 | 0.12 | 0.03 | 0.21 | 0.14 | 0.24 | 0.14 | 0.33 | 1.03 | 0.76 | 0.00 |
| 41 | 0.70 | 0.39 | 0.10 | 0.57 | 0.19 | 0.05 | 0.36 | 0.14 | 0.41 | 0.32 | 0.38 | 0.84 | 0.96 | 0.00 |
| 42 | 0.74 | 0.82 | 0.26 | 0.66 | 0.18 | 0.12 | 0.54 | 0.19 | 0.60 | 0.67 | 0.36 | 0.75 | 1.20 | 0.00 |
| 43 | 1.00 | 0.27 | 0.17 | 0.51 | 0.11 | 0.07 | 0.39 | 0.14 | 0.45 | 0.40 | 0.39 | 0.77 | 1.07 | 0.00 |
| 44 | 1.02 | 0.68 | 0.66 | 0.65 | 0.55 | 0.13 | 0.39 | 0.64 | 0.44 | 0.64 | 0.46 | 1.11 | 0.78 | 0.00 |
| 45 | 0.84 | 0.48 | 0.37 | 0.57 | 0.22 | 0.10 | 0.38 | 0.29 | 0.42 | 0.34 | 0.30 | 0.87 | 0.86 | 0.00 |
| 46 | 0.70 | 0.45 | 0.26 | 0.63 | 0.26 | 0.10 | 0.48 | 0.33 | 0.51 | 0.33 | 0.38 | 0.80 | 0.98 | 0.00 |
| 47 | 0.41 | 1.28 | 0.29 | 1.29 | 0.27 | 0.18 | 0.93 | 0.19 | 1.10 | 1.18 | 0.05 | 0.53 | 1.59 | 0.00 |
| 48 | 0.53 | 0.43 | 0.11 | 0.64 | 0.16 | 0.08 | 0.43 | 0.20 | 0.49 | 0.42 | 0.14 | 0.71 | 1.03 | 0.00 |
| 49 | 0.61 | 0.92 | 0.16 | 1.13 | 0.18 | 0.13 | 0.90 | 0.18 | 1.08 | 0.99 | 0.23 | 0.48 | 1.54 | 0.00 |
| 50 | 0.79 | 3.11 | 0.16 | 3.03 | 0.17 | 0.74 | 1.92 | 0.21 | 2.48 | 1.89 | 1.22 | 0.26 | 2.38 | 0.00 |
| 51 | 0.74 | 1.05 | 0.26 | 0.89 | 0.14 | 0.13 | 0.75 | 0.17 | 0.90 | 0.96 | 0.21 | 0.54 | 1.45 | 0.00 |
| 52 | 0.50 | 1.14 | 0.18 | 1.00 | 0.22 | 0.13 | 0.81 | 0.17 | 0.97 | 1.06 | 0.14 | 0.55 | 1.49 | 0.00 |
| 53 | 0.50 | 2.64 | 0.20 | 3.48 | 0.19 | 0.83 | 2.09 | 0.21 | 2.62 | 2.05 | 0.75 | 0.27 | 2.41 | 0.00 |
| 54 | 0.27 | 3.88 | 1.62 | 5.53 | 3.13 | 20.62 | 11.07 | 6.63 | 7.67 | 6.87 | 4.24 | 0.04 | 2.15 | 0.00 |
| 55 | 0.30 | 3.12 | 0.15 | 2.82 | 0.29 | 0.80 | 1.87 | 0.31 | 2.37 | 2.03 | 1.44 | 0.26 | 2.31 | 0.00 |
| 56 | 0.38 | 1.55 | 0.21 | 1.10 | 0.27 | 0.15 | 0.72 | 0.27 | 0.86 | 1.27 | 0.22 | 0.56 | 1.33 | 0.00 |
| 57 | 0.39 | 0.80 | 0.39 | 0.84 | 0.17 | 0.12 | 0.57 | 0.27 | 0.66 | 0.83 | 0.17 | 0.67 | 1.17 | 0.00 |
| 58 | 0.46 | 1.08 | 0.27 | 0.95 | 0.29 | 0.18 | 0.68 | 0.37 | 0.75 | 0.87 | 0.10 | 0.69 | 1.15 | 0.00 |
| 59 | 0.53 | 0.85 | 0.62 | 0.80 | 0.51 | 0.16 | 0.56 | 0.51 | 0.62 | 1.20 | 0.11 | 0.85 | 1.08 | 0.00 |
| 60 | 0.30 | 0.32 | 0.14 | 0.56 | 0.22 | 0.06 | 0.34 | 0.17 | 0.37 | 0.41 | 0.08 | 0.90 | 0.90 | 0.00 |
| 61 | 0.48 | 0.52 | 0.14 | 0.40 | 0.15 | 0.06 | 0.37 | 0.15 | 0.42 | 0.61 | 0.07 | 0.76 | 0.92 | 0.00 |
| 62 | 0.44 | 0.70 | 0.20 | 0.52 | 0.13 | 0.08 | 0.50 | 0.20 | 0.55 | 0.80 | 0.10 | 0.79 | 1.08 | 0.00 |
| 63 | 0.41 | 2.79 | 0.30 | 1.18 | 0.11 | 0.29 | 1.19 | 0.25 | 1.45 | 3.15 | 0.19 | 0.49 | 1.63 | 0.00 |
| 64 | 0.79 | 0.79 | 0.23 | 0.56 | 0.23 | 0.07 | 0.42 | 0.18 | 0.47 | 0.61 | 0.30 | 0.82 | 0.91 | 0.00 |
| 65 | 0.93 | 0.49 | 0.19 | 0.54 | 0.17 | 0.07 | 0.48 | 0.12 | 0.54 | 0.75 | 0.16 | 0.81 | 0.91 | 0.00 |
| 66 | 0.67 | 1.21 | 0.19 | 1.07 | 0.13 | 0.15 | 0.93 | 0.16 | 1.11 | 1.28 | 0.29 | 0.54 | 1.33 | 0.00 |
| 67 | 0.98 | 0.85 | 0.62 | 0.77 | 0.46 | 0.13 | 0.73 | 0.25 | 0.81 | 1.22 | 0.16 | 1.17 | 0.97 | 0.00 |
| 68 | 1.17 | 0.30 | 0.31 | 0.38 | 0.16 | 0.04 | 0.33 | 0.18 | 0.35 | 0.29 | 0.11 | 1.46 | 0.60 | 0.00 |
| 69 | 0.92 | 0.25 | 0.36 | 0.34 | 0.23 | 0.05 | 0.26 | 0.21 | 0.29 | 0.34 | 0.26 | 1.30 | 0.63 | 0.00 |
| 70 | 0.51 | 0.46 | 0.29 | 0.50 | 0.32 | 0.08 | 0.36 | 0.31 | 0.40 | 0.89 | 0.09 | 1.20 | 0.74 | 0.00 |
| 71 | 0.41 | 0.78 | 0.54 | 0.56 | 0.38 | 0.12 | 0.48 | 0.33 | 0.51 | 1.02 | 0.08 | 1.08 | 0.84 | 0.00 |
| 72 | 1.21 | 0.43 | 0.36 | 0.58 | 0.35 | 0.08 | 0.34 | 0.32 | 0.37 | 0.52 | 0.30 | 1.27 | 0.68 | 0.00 |
| 73 | 0.74 | 0.48 | 0.65 | 0.62 | 0.69 | 0.18 | 0.49 | 0.58 | 0.52 | 0.42 | 0.18 | 1.53 | 0.56 | 0.00 |
| 74 | 1.49 | 0.37 | 0.40 | 0.56 | 0.28 | 0.06 | 0.48 | 0.15 | 0.55 | 0.34 | 0.12 | 1.67 | 0.78 | 0.00 |
| 75 | 2.47 | 0.22 | 1.34 | 0.31 | 0.76 | 0.05 | 0.46 | 0.40 | 0.51 | 0.15 | 0.19 | 1.80 | 0.45 | 0.00 |
| 76 | 3.92 | 0.15 | 0.37 | 0.22 | 0.27 | 0.03 | 0.22 | 0.18 | 0.25 | 0.03 | 0.14 | 2.00 | 0.48 | 0.00 |
| 77 | 0.98 | 0.27 | 0.32 | 0.35 | 0.21 | 0.04 | 0.37 | 0.17 | 0.42 | 0.18 | 0.11 | 1.95 | 0.36 | 0.00 |
| 78 | 1.14 | 0.24 | 0.51 | 0.24 | 0.38 | 0.03 | 0.24 | 0.25 | 0.28 | 0.21 | 0.14 | 1.93 | 0.49 | 0.00 |
| 79 | 3.18 | 0.14 | 0.29 | 0.22 | 0.25 | 0.02 | 0.17 | 0.22 | 0.21 | 0.06 | 0.11 | 2.02 | 0.39 | 0.00 |
| 80 | 1.24 | 0.22 | 0.31 | 0.39 | 0.20 | 0.04 | 0.39 | 0.17 | 0.44 | 0.19 | 0.16 | 1.81 | 0.37 | 0.00 |
| 81 | 0.98 | 0.29 | 1.91 | 0.42 | 1.13 | 0.04 | 0.26 | 0.76 | 0.30 | 0.64 | 0.39 | 1.82 | 0.40 | 0.00 |
| 82 | 0.54 | 3.07 | 57.15 | 2.36 | 48.00 | 1.24 | 1.71 | 26.12 | 1.69 | 10.08 | 3.57 | 0.50 | 0.86 | 0.00 |
| 83 | 1.29 | 0.18 | 0.37 | 0.23 | 0.33 | 0.03 | 0.15 | 0.45 | 0.17 | 0.12 | 0.09 | 2.00 | 0.27 | 0.00 |
| 84 | 0.86 | 0.43 | 1.06 | 0.58 | 0.94 | 0.16 | 0.34 | 1.08 | 0.36 | 0.72 | 0.26 | 1.64 | 0.29 | 0.00 |
| 85 | 1.38 | 0.13 | 0.27 | 0.21 | 0.24 | 0.03 | 0.15 | 0.27 | 0.19 | 0.15 | 0.06 | 2.02 | 0.23 | 0.00 |
| 86 | 1.56 | 0.08 | 0.25 | 0.11 | 0.23 | 0.01 | 0.10 | 0.20 | 0.11 | 0.06 | 0.08 | 2.07 | 0.21 | 0.00 |
| 87 | 0.63 | 0.28 | 0.61 | 0.37 | 0.38 | 0.04 | 0.26 | 0.34 | 0.31 | 0.31 | 0.06 | 1.92 | 0.38 | 0.00 |
| 88 | 0.66 | 0.32 | 0.85 | 0.26 | 0.47 | 0.04 | 0.36 | 0.26 | 0.41 | 0.20 | 0.06 | 1.85 | 0.50 | 0.00 |
| 89 | 0.38 | 1.14 | 0.52 | 0.63 | 0.43 | 0.15 | 0.53 | 0.40 | 0.58 | 1.31 | 0.09 | 1.00 | 0.88 | 0.00 |
| 90 | 1.09 | 0.17 | 0.46 | 0.27 | 0.41 | 0.03 | 0.17 | 0.29 | 0.19 | 0.14 | 0.13 | 1.92 | 0.34 | 0.00 |
| 91 | 0.99 | 0.54 | 0.93 | 0.41 | 0.62 | 0.08 | 0.31 | 0.66 | 0.35 | 0.50 | 0.24 | 1.55 | 0.40 | 0.00 |
| 92 | 1.81 | 0.13 | 0.36 | 0.19 | 0.32 | 0.02 | 0.13 | 0.24 | 0.14 | 0.09 | 0.07 | 2.02 | 0.25 | 0.00 |
| 93 | 0.67 | 0.21 | 0.38 | 0.14 | 0.33 | 0.02 | 0.14 | 0.25 | 0.17 | 0.08 | 0.04 | 2.02 | 0.29 | 0.00 |
| 94 | 0.65 | 0.23 | 0.33 | 0.22 | 0.32 | 0.03 | 0.12 | 0.34 | 0.15 | 0.10 | 0.05 | 2.00 | 0.21 | 0.00 |
| 95 | 1.06 | 0.51 | 3.78 | 0.46 | 3.02 | 0.16 | 0.29 | 2.41 | 0.32 | 2.04 | 1.52 | 1.64 | 0.30 | 0.00 |
| 96 | 8.86 | 0.02 | 0.03 | 0.02 | 0.03 | 0.00 | 0.01 | 0.03 | 0.01 | 0.01 | 0.00 | 0.30 | 0.02 | 11.28 |
| 97 | 1.13 | 0.17 | 0.42 | 0.14 | 0.27 | 0.02 | 0.07 | 0.38 | 0.08 | 0.08 | 0.03 | 1.99 | 0.11 | 0.00 |
| 98 | 0.51 | 0.07 | 0.13 | 0.12 | 0.14 | 0.01 | 0.07 | 0.26 | 0.09 | 0.07 | 0.02 | 2.06 | 0.22 | 0.00 |
| 99 | 0.97 | 0.07 | 0.20 | 0.13 | 0.21 | 0.01 | 0.10 | 0.20 | 0.13 | 0.11 | 0.04 | 2.04 | 0.17 | 0.00 |
| 100 | 0.25 | 0.78 | 98.15 | 1.21 | 147.04 | 4.02 | 2.09 | 176.81 | 1.32 | 4.24 | 15.51 | 0.02 | 0.63 | 0.00 |
| Base | 100 | 6.30E-04 | 4.40E-04 | 6.30E-04 | 4.40E-04 | 1.13 | 2.98 | 0.38 | 1.58 | 0.68 | 5.17 | 46.72 | 0.72 | 7.57 |

**Figure S8: Conservation state enrichment values.** The first column displays the state number followed by the percentage of genome in each state, and then the values of the fold enrichments represented in the heatmap in **Figure 2B** along with additional columns for: enrichments for TES of protein coding genes, TES of pseudogenes, coding sequence (CDS), untranslated regions (UTRs), and assembly gaps. The last row gives the % of the genome covered by the annotation in each column. In the case of median non-exonic DHS, the last row is the median genome coverage across the DNase I experiments. The values in this figure are also available in **Table S2**.

| State assignment using original model | Probability of assignment to same state using uniform transition probabilities |
|---|---|
| 1 | 0.99 |
| 2 | 0.98 |
| 3 | 0.98 |
| 4 | 0.91 |
| 5 | 0.91 |
| 6 | 0.87 |
| 7 | 0.86 |
| 8 | 0.84 |
| 9 | 0.93 |
| 10 | 0.97 |
| 11 | 0.92 |
| 12 | 0.88 |
| 13 | 0.88 |
| 14 | 0.85 |
| 15 | 0.95 |
| 16 | 0.85 |
| 17 | 0.85 |
| 18 | 0.91 |
| 19 | 0.93 |
| 20 | 0.92 |
| 21 | 0.93 |
| 22 | 0.91 |
| 23 | 0.92 |
| 24 | 0.94 |
| 25 | 0.92 |
| 26 | 0.92 |
| 27 | 0.94 |
| 28 | 0.90 |
| 29 | 0.91 |
| 30 | 0.93 |
| 31 | 0.93 |
| 32 | 0.90 |
| 33 | 0.93 |
| 34 | 0.94 |
| 35 | 0.92 |
| 36 | 0.89 |
| 37 | 0.88 |
| 38 | 0.92 |
| 39 | 0.89 |
| 40 | 0.88 |
| 41 | 0.86 |
| 42 | 0.84 |
| 43 | 0.91 |
| 44 | 0.84 |
| 45 | 0.92 |
| 46 | 0.91 |
| 47 | 0.97 |
| 48 | 0.78 |
| 49 | 0.84 |
| 50 | 0.91 |
| 51 | 0.90 |
| 52 | 0.91 |
| 53 | 0.93 |
| 54 | 0.95 |
| 55 | 0.96 |
| 56 | 0.87 |
| 57 | 0.93 |
| 58 | 0.85 |
| 59 | 0.87 |
| 60 | 0.78 |
| 61 | 0.88 |
| 62 | 0.86 |
| 63 | 0.90 |
| 64 | 0.92 |
| 65 | 0.92 |
| 66 | 0.96 |
| 67 | 0.95 |
| 68 | 0.92 |
| 69 | 0.92 |
| 70 | 0.90 |
| 71 | 0.92 |
| 72 | 0.87 |
| 73 | 0.87 |
| 74 | 0.95 |
| 75 | 0.92 |
| 76 | 0.99 |
| 77 | 0.97 |
| 78 | 0.97 |
| 79 | 0.99 |
| 80 | 0.91 |
| 81 | 0.93 |
| 82 | 0.94 |
| 83 | 0.96 |
| 84 | 0.90 |
| 85 | 0.97 |
| 86 | 0.95 |
| 87 | 0.99 |
| 88 | 0.96 |
| 89 | 0.93 |
| 90 | 0.97 |
| 91 | 0.91 |
| 92 | 0.99 |
| 93 | 0.99 |
| 94 | 1.00 |
| 95 | 0.93 |
| 96 | 1.00 |
| 97 | 1.00 |
| 98 | 1.00 |
| 99 | 0.99 |
| 100 | 0.98 |

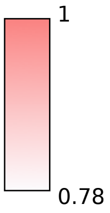


**Figure S9: Comparison of state annotations from the learned model versus the learned model except with uniform probabilities.** The first column shows the state IDs. The second column displays for each state the fraction of bases annotated to the state using the learned model that were also assigned when using the learned model except with uniform transition probabilities. For all states the majority of bases were assigned to the same state when using the model with uniform transition probabilities with the fraction ranging between 0.78 and 1.00.

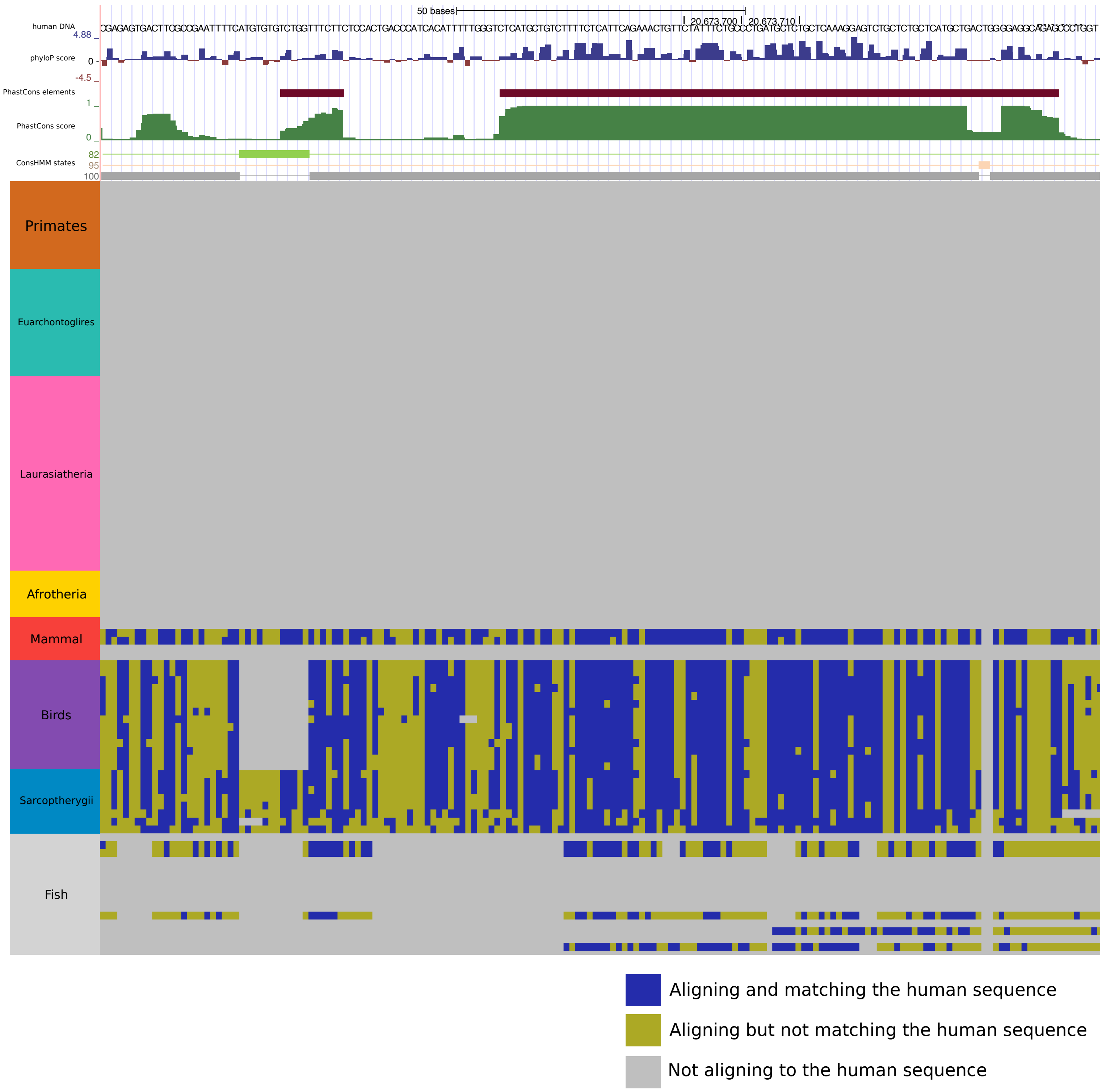


**Figure S10: Illustration of conservation state assignments at an additional locus.** Similar to **Figure 1B**, but illustrating the conservation state assignment at a different locus: chr22:20,673,589-20,673,761. The top tracks are the DNA sequence, the PhyloP score, PhastCons elements, and then PhastCons score. Below this set of tracks are the conservation state assignments with only the states assigned to at least one nucleotide in the locus shown. Below the conservation state assignments is a color encoding of the input multiple sequence alignment. The major clade of species as annotated on the UCSC genome browser[21] are labeled and ordered based on divergence from human. The figure is an example of positions with high constraint scores from PhyloP and PhastCons, while the multiple sequence alignment lacks alignment to most mammals, which is suggestive of alignment artifacts. ConsHMM states 82 and 100 capture the pattern of non-mammalian vertebrates aligning and/or matching the human genome, without most mammals. State 95 captures the pattern of all species having low alignment and matching probabilities and relatively proximal to states with higher probabilities of alignment and matching (**Figures S5** and **S6**).

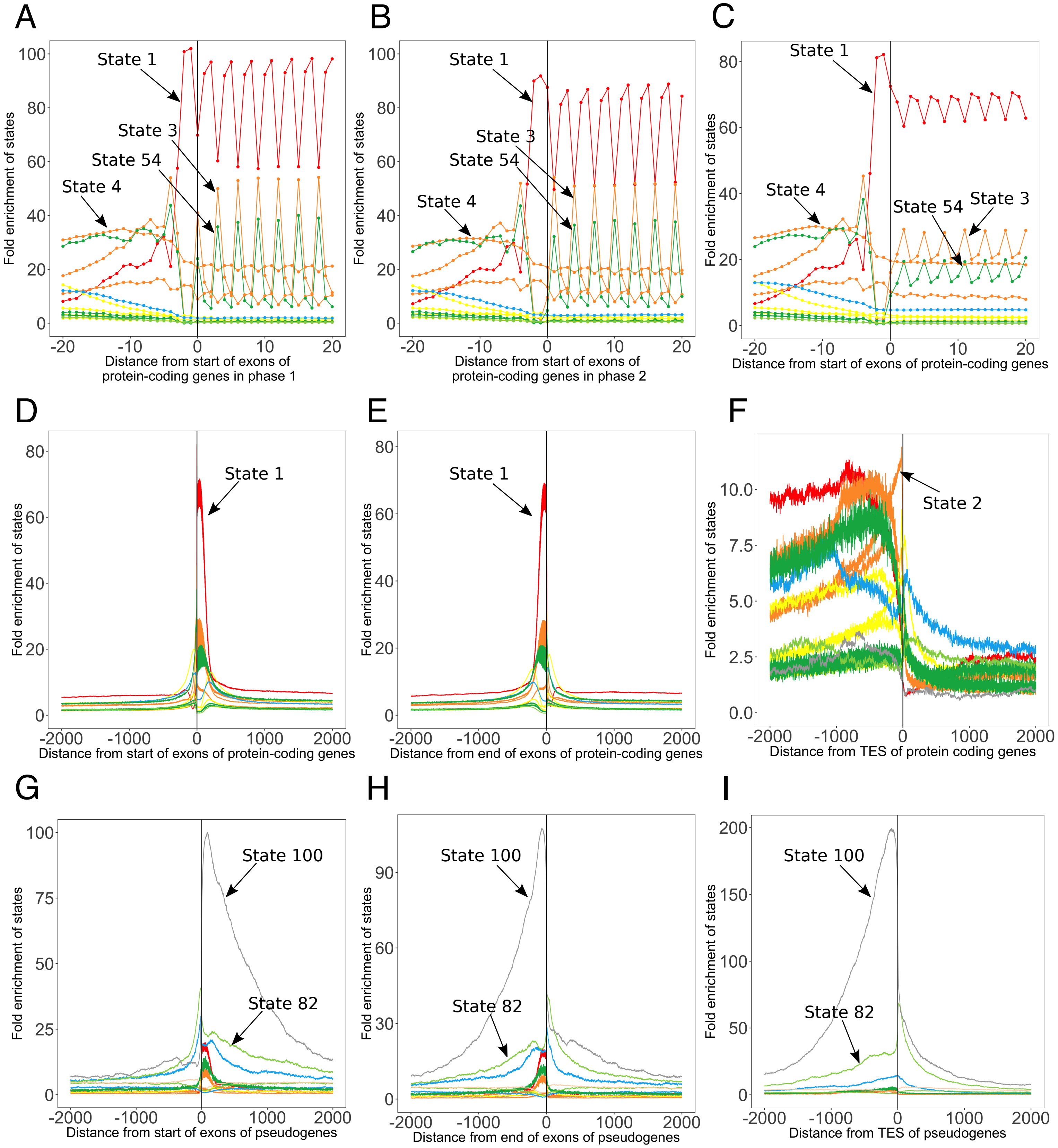


**Figure S11: Additional conservation state positional enrichments.** The figure shows additional positional enrichment plots similar to what was shown in **Figure 3**. These additional enrichment plots include enrichments relative to start of exons of protein coding genes for **(A)** phase 1 and **(B)** phase 2 exons, **(C)** all exons, and a zoomed out view of enrichments relative to **(D)** the start and **(E)** end of all exons of protein coding genes. Also shown are enrichment plots relative to **(F)** TES of protein coding genes, **(G)** start and **(H)** end of exons of pseudogenes as well as **(I)** TES of pseudogenes. Enrichments were computed relative to a genome-wide background. The subset of states included in the figure was composed of the states that had at least a 3 fold enrichment at some position within +/-2kb from the anchor point.

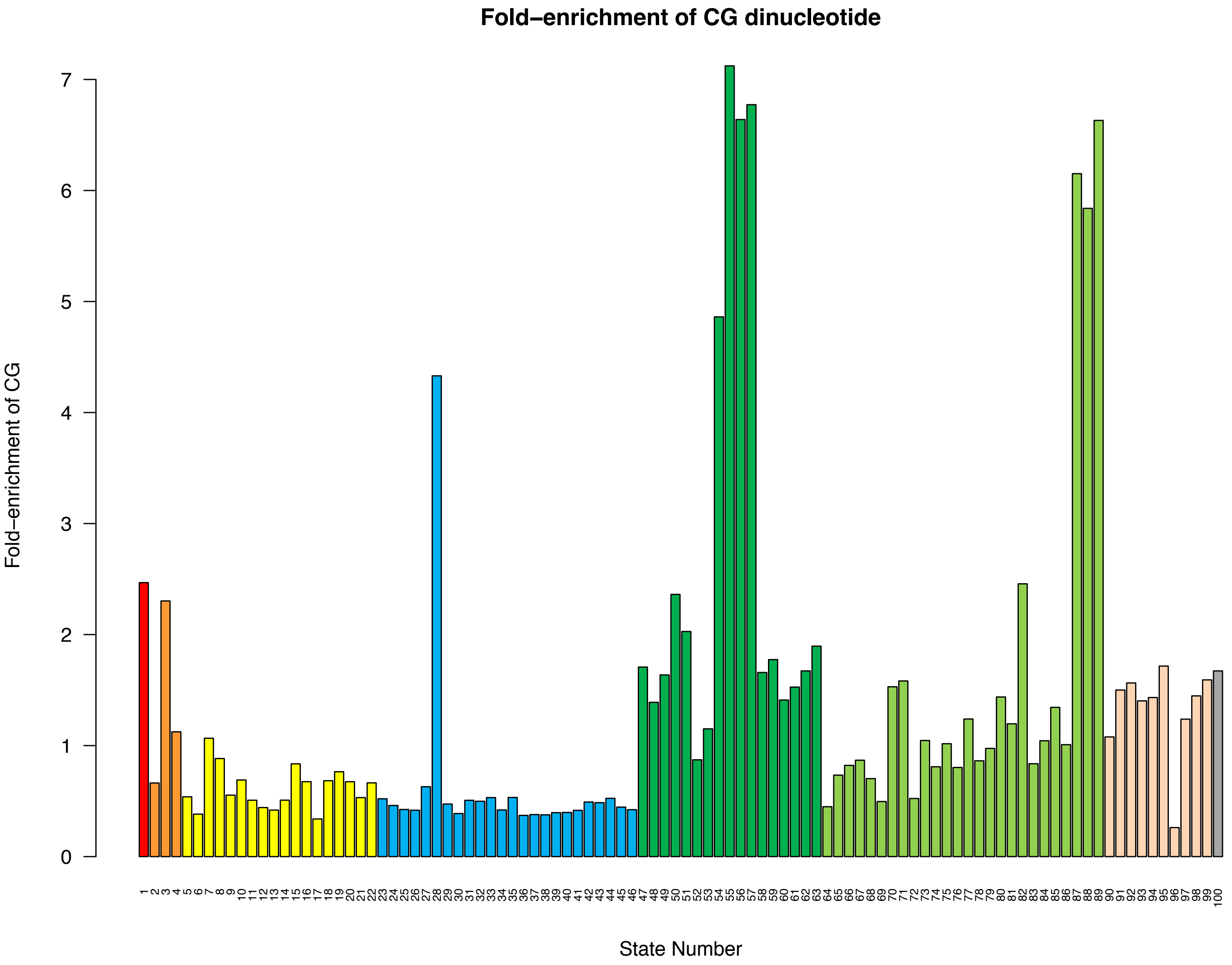


**Figure S12: Enrichment of CG dinucleotides in the states.** The bar graph shows for each state its fold enrichment for CG dinucleotides. States 55-57 and 87-89 had the highest enrichments followed by states 28 and 54.

| state | detection of chemical stimulus involved in sensory perception of smell | system development | nucleosome | cellular developmental process | regulation of cellular process | transcription from RNA polymerase II promoter | cell-cell signaling | calcium ion binding | receptor binding | antigen binding | protein-DNA complex assembly | cell adhesion | mesonephros development | neuromuscular process |
|---|---|---|---|---|---|---|---|---|---|---|---|---|---|---|
| 1 | 0.3 | 14.1 | 41.3 | 5.4 | 4.2 | 7.9 | 7.9 | 12.2 | 0.0 | 0.0 | 13.2 | 8.8 | 1.7 | 2.5 |
| 2 | 0.0 | 41.2 | 0.1 | 30.6 | 6.9 | 19.0 | 11.0 | 3.6 | 7.5 | 0.0 | 0.0 | 3.7 | 5.3 | 1.7 |
| 3 | 319.6 | 2.6 | 0.2 | 0.1 | 18.6 | 0.1 | 2.8 | 13.5 | 0.0 | 0.0 | 0.0 | 4.3 | 0.8 | 0.1 |
| 4 | 0.0 | 21.0 | 0.0 | 17.7 | 5.6 | 24.7 | 8.1 | 0.1 | 6.5 | 0.0 | 0.1 | 2.2 | 2.2 | 4.0 |
| 5 | 0.0 | 26.8 | 0.0 | 18.9 | 10.0 | 13.4 | 20.8 | 0.5 | 11.3 | 0.1 | 0.0 | 8.6 | 3.6 | 3.4 |
| 6 | 0.0 | 5.6 | 0.0 | 5.3 | 3.7 | 2.4 | 3.4 | 1.5 | 1.1 | 0.0 | 0.0 | 2.7 | 2.2 | 0.1 |
| 7 | 0.2 | 21.6 | 0.1 | 12.7 | 11.6 | 9.2 | 12.0 | 1.4 | 5.1 | 0.0 | 0.0 | 1.5 | 3.6 | 9.4 |
| 8 | 0.0 | 19.8 | 0.0 | 21.0 | 2.9 | 8.2 | 9.0 | 1.1 | 6.5 | 0.7 | 0.0 | 11.7 | 5.3 | 2.9 |
| 9 | 0.0 | 10.2 | 0.0 | 6.4 | 3.6 | 1.7 | 4.2 | 4.4 | 4.1 | 0.0 | 0.0 | 4.6 | 4.1 | 0.4 |
| 10 | 0.0 | 8.6 | 0.0 | 6.2 | 1.0 | 0.7 | 2.9 | 3.3 | 5.1 | 0.0 | 0.0 | 7.5 | 2.6 | 0.7 |
| 11 | 0.1 | 0.6 | 0.0 | 0.1 | 0.1 | 0.0 | 0.5 | 4.4 | 1.0 | 0.5 | 0.0 | 0.1 | 0.0 | 0.0 |
| 12 | 1.7 | 0.3 | 0.1 | 0.2 | 0.6 | 0.0 | 0.8 | 0.7 | 1.2 | 0.1 | 0.0 | 0.4 | 0.0 | 0.1 |
| 13 | 0.0 | 0.7 | 0.0 | 0.6 | 0.0 | 0.0 | 0.7 | 0.5 | 0.1 | 0.1 | 0.0 | 0.7 | 0.0 | 0.1 |
| 14 | 0.0 | 2.0 | 0.0 | 2.6 | 2.9 | 0.4 | 5.2 | 1.4 | 2.6 | 0.3 | 0.0 | 0.2 | 0.4 | 0.0 |
| 15 | 0.0 | 15.4 | 0.0 | 10.8 | 2.9 | 2.0 | 4.6 | 2.5 | 6.0 | 0.0 | 0.0 | 5.2 | 3.1 | 0.3 |
| 16 | 0.1 | 0.0 | 0.0 | 0.0 | 0.0 | 0.0 | 0.0 | 0.5 | 0.1 | 1.4 | 0.0 | 0.4 | 0.0 | 0.1 |
| 17 | 0.3 | 0.0 | 0.0 | 0.0 | 0.0 | 0.0 | 0.0 | 0.3 | 0.0 | 0.7 | 0.0 | 0.2 | 0.0 | 0.0 |
| 18 | 0.0 | 0.1 | 0.0 | 0.0 | 0.0 | 0.0 | 0.2 | 0.1 | 0.1 | 1.0 | 0.0 | 0.5 | 0.0 | 0.1 |
| 19 | 0.0 | 19.8 | 0.0 | 15.5 | 2.6 | 1.1 | 8.1 | 3.1 | 9.5 | 0.1 | 0.0 | 10.5 | 3.1 | 0.9 |
| 20 | 0.0 | 5.1 | 0.1 | 2.1 | 2.0 | 1.4 | 1.1 | 0.4 | 0.1 | 0.1 | 0.3 | 0.7 | 0.1 | 1.7 |
| 21 | 1.2 | 0.4 | 0.1 | 0.1 | 0.0 | 0.0 | 0.1 | 0.5 | 0.1 | 0.0 | 0.4 | 1.4 | 0.0 | 0.1 |
| 22 | 0.0 | 0.1 | 0.2 | 0.1 | 0.1 | 0.1 | 0.9 | 0.5 | 0.7 | 0.0 | 0.3 | 1.0 | 0.1 | 0.1 |
| 23 | 0.0 | 0.0 | 0.1 | 0.0 | 0.0 | 0.0 | 0.0 | 0.0 | 0.0 | 0.5 | 0.3 | 0.0 | 0.0 | 0.1 |
| 24 | 0.0 | 0.0 | 2.3 | 0.0 | 0.0 | 0.0 | 0.0 | 0.0 | 0.0 | 0.1 | 3.2 | 0.1 | 0.0 | 0.0 |
| 25 | 0.0 | 0.0 | 0.0 | 0.0 | 0.0 | 0.0 | 0.0 | 0.0 | 0.0 | 0.1 | 0.3 | 0.0 | 0.0 | 0.1 |
| 26 | 0.0 | 0.0 | 0.0 | 0.0 | 0.0 | 0.0 | 0.0 | 0.0 | 0.0 | 0.5 | 0.7 | 0.1 | 0.0 | 0.0 |
| 27 | 0.0 | 0.0 | 0.4 | 0.0 | 0.0 | 0.0 | 0.0 | 0.0 | 0.0 | 0.1 | 0.9 | 0.0 | 0.0 | 0.3 |
| 28 | 60.0 | 5.8 | 0.9 | 5.9 | 24.9 | 12.5 | 10.5 | 0.9 | 1.3 | 1.4 | 0.1 | 0.9 | 1.1 | 0.7 |
| 29 | 0.5 | 0.0 | 0.1 | 0.0 | 0.0 | 0.0 | 0.0 | 0.0 | 0.0 | 0.5 | 0.1 | 0.0 | 0.0 | 0.7 |
| 30 | 0.2 | 0.0 | 3.9 | 0.0 | 0.0 | 0.0 | 0.0 | 0.0 | 0.0 | 0.5 | 2.0 | 0.0 | 0.0 | 0.3 |
| 31 | 1.5 | 0.0 | 1.5 | 0.0 | 0.0 | 0.0 | 0.0 | 0.0 | 0.0 | 0.0 | 1.4 | 0.0 | 0.0 | 0.1 |
| 32 | 0.0 | 0.0 | 2.3 | 0.0 | 0.3 | 0.0 | 0.0 | 0.0 | 0.0 | 1.0 | 1.7 | 0.0 | 0.0 | 0.0 |
| 33 | 15.3 | 0.0 | 2.8 | 0.0 | 1.0 | 0.1 | 0.0 | 0.0 | 0.0 | 1.7 | 1.4 | 0.0 | 0.6 | 0.4 |
| 34 | 16.5 | 0.0 | 1.5 | 0.0 | 0.0 | 0.0 | 0.0 | 0.0 | 0.0 | 0.1 | 1.2 | 0.1 | 0.0 | 0.1 |
| 35 | 0.0 | 0.0 | 0.2 | 0.0 | 0.0 | 0.1 | 0.1 | 0.4 | 0.0 | 0.0 | 1.2 | 0.0 | 0.0 | 0.0 |
| 36 | 0.0 | 0.0 | 0.1 | 0.0 | 0.0 | 0.0 | 0.0 | 0.0 | 0.0 | 0.1 | 0.3 | 0.0 | 0.0 | 0.1 |
| 37 | 0.0 | 0.0 | 0.1 | 0.0 | 0.0 | 0.0 | 0.0 | 0.0 | 0.0 | 0.1 | 0.1 | 0.0 | 0.0 | 0.4 |
| 38 | 0.1 | 0.0 | 0.9 | 0.0 | 0.1 | 0.1 | 0.0 | 0.0 | 0.0 | 1.0 | 4.0 | 0.1 | 0.0 | 0.0 |
| 39 | 4.8 | 0.0 | 10.7 | 0.0 | 0.0 | 0.0 | 0.0 | 0.1 | 0.0 | 0.5 | 3.6 | 0.5 | 0.0 | 0.0 |
| 40 | 0.9 | 0.0 | 0.1 | 0.0 | 0.0 | 0.0 | 0.0 | 0.0 | 0.0 | 0.3 | 1.4 | 0.3 | 0.0 | 0.3 |
| 41 | 0.9 | 0.0 | 0.2 | 0.0 | 0.0 | 0.0 | 0.0 | 0.5 | 0.0 | 0.0 | 0.1 | 0.2 | 0.0 | 0.0 |
| 42 | 0.5 | 0.0 | 0.0 | 0.0 | 0.0 | 0.0 | 0.0 | 0.4 | 0.0 | 0.0 | 0.0 | 0.0 | 0.0 | 0.1 |
| 43 | 0.0 | 0.0 | 5.0 | 0.0 | 0.1 | 0.0 | 0.0 | 0.0 | 0.0 | 0.1 | 2.5 | 0.0 | 0.0 | 0.0 |
| 44 | 28.0 | 0.0 | 3.9 | 0.0 | 0.1 | 0.0 | 0.0 | 0.0 | 0.0 | 1.7 | 0.9 | 0.0 | 0.0 | 0.0 |
| 45 | 6.7 | 0.0 | 4.4 | 0.0 | 0.1 | 0.0 | 0.0 | 0.1 | 0.3 | 1.7 | 2.9 | 0.0 | 0.0 | 0.0 |
| 46 | 2.6 | 0.0 | 0.1 | 0.0 | 0.0 | 0.0 | 0.0 | 0.4 | 0.1 | 0.3 | 0.2 | 0.1 | 0.0 | 0.0 |
| 47 | 0.0 | 0.1 | 0.1 | 0.1 | 0.0 | 0.0 | 0.0 | 0.6 | 1.3 | 0.7 | 0.0 | 1.0 | 0.1 | 0.3 |
| 48 | 3.1 | 0.0 | 2.8 | 0.0 | 0.0 | 0.0 | 0.0 | 0.0 | 0.0 | 1.0 | 1.2 | 0.0 | 0.0 | 0.4 |
| 49 | 0.0 | 0.0 | 0.2 | 0.0 | 0.0 | 0.0 | 0.1 | 0.0 | 0.0 | 0.5 | 0.3 | 0.0 | 0.0 | 0.4 |
| 50 | 0.0 | 29.6 | 0.0 | 21.7 | 5.6 | 4.7 | 12.6 | 2.9 | 15.8 | 0.1 | 0.0 | 11.1 | 8.0 | 0.4 |
| 51 | 0.0 | 0.5 | 0.0 | 0.0 | 0.1 | 0.0 | 0.2 | 0.7 | 0.2 | 0.5 | 0.0 | 0.8 | 0.0 | 0.1 |
| 52 | 0.0 | 0.7 | 0.0 | 0.2 | 0.4 | 0.0 | 0.6 | 0.9 | 0.8 | 0.0 | 0.0 | 0.8 | 0.0 | 0.1 |
| 53 | 0.0 | 44.6 | 0.0 | 30.2 | 7.9 | 9.2 | 18.8 | 2.1 | 15.4 | 0.3 | 0.0 | 9.4 | 9.5 | 2.0 |
| 54 | 53.2 | 3.9 | 32.3 | 0.9 | 1.7 | 0.9 | 2.8 | 15.9 | 0.2 | 0.0 | 8.9 | 8.6 | 1.7 | 1.2 |
| 55 | 0.0 | 22.2 | 0.0 | 18.3 | 2.6 | 4.0 | 12.0 | 5.0 | 13.3 | 0.1 | 0.0 | 6.3 | 11.0 | 2.5 |
| 56 | 6.3 | 0.1 | 0.1 | 0.0 | 0.0 | 0.0 | 0.2 | 0.5 | 0.3 | 0.3 | 0.0 | 0.6 | 0.0 | 0.1 |
| 57 | 3.1 | 0.0 | 1.2 | 0.0 | 0.0 | 0.0 | 0.0 | 0.0 | 0.0 | 1.4 | 1.4 | 0.0 | 0.0 | 0.3 |
| 58 | 5.9 | 0.0 | 1.2 | 0.0 | 0.0 | 0.0 | 0.0 | 0.5 | 0.0 | 0.5 | 0.6 | 0.0 | 0.0 | 0.1 |
| 59 | 17.7 | 0.0 | 7.7 | 0.0 | 0.2 | 0.0 | 0.0 | 0.0 | 0.0 | 0.3 | 0.9 | 0.0 | 0.0 | 0.0 |
| 60 | 3.7 | 0.0 | 0.1 | 0.0 | 0.0 | 0.0 | 0.0 | 0.2 | 0.0 | 0.3 | 0.1 | 0.1 | 0.0 | 0.1 |
| 61 | 2.6 | 0.0 | 0.9 | 0.0 | 0.0 | 0.0 | 0.0 | 0.0 | 0.1 | 0.5 | 0.7 | 0.0 | 0.0 | 0.7 |
| 62 | 0.0 | 0.0 | 0.1 | 0.0 | 0.0 | 0.0 | 0.0 | 0.0 | 0.1 | 1.0 | 0.9 | 0.0 | 0.0 | 0.1 |
| 63 | 0.0 | 1.4 | 0.0 | 0.2 | 0.2 | 0.0 | 0.2 | 1.2 | 0.6 | 0.5 | 0.1 | 2.6 | 0.1 | 0.1 |
| 64 | 1.7 | 0.0 | 0.1 | 0.0 | 0.0 | 0.0 | 0.0 | 0.7 | 0.2 | 1.4 | 0.4 | 0.0 | 0.1 | 0.1 |
| 65 | 0.3 | 0.0 | 6.3 | 0.0 | 0.0 | 0.0 | 0.0 | 0.1 | 0.0 | 0.3 | 3.6 | 0.0 | 0.0 | 0.0 |
| 66 | 0.1 | 0.9 | 0.0 | 0.4 | 0.0 | 0.0 | 0.1 | 2.1 | 0.3 | 0.3 | 0.0 | 0.7 | 0.0 | 0.1 |
| 67 | 0.0 | 0.0 | 0.0 | 0.0 | 0.0 | 0.0 | 0.2 | 0.5 | 0.2 | 0.5 | 0.0 | 0.3 | 0.0 | 0.1 |
| 68 | 0.0 | 0.0 | 0.0 | 0.0 | 0.0 | 0.0 | 0.0 | 0.7 | 0.0 | 0.5 | 0.7 | 0.0 | 0.0 | 0.1 |
| 69 | 0.5 | 0.0 | 0.6 | 0.0 | 0.0 | 0.0 | 0.0 | 0.0 | 0.0 | 1.0 | 0.9 | 0.0 | 0.0 | 0.3 |
| 70 | 0.5 | 0.0 | 4.4 | 0.0 | 0.0 | 0.0 | 0.0 | 0.0 | 0.0 | 2.3 | 1.4 | 0.0 | 0.0 | 0.0 |
| 71 | 0.1 | 0.0 | 0.2 | 0.0 | 0.0 | 0.0 | 0.0 | 0.0 | 0.0 | 8.2 | 0.2 | 0.0 | 0.0 | 0.1 |
| 72 | 0.1 | 0.0 | 0.6 | 0.0 | 0.0 | 0.0 | 0.0 | 0.0 | 0.0 | 7.4 | 0.3 | 0.0 | 0.0 | 0.0 |
| 73 | 1.0 | 0.0 | 0.4 | 0.0 | 0.0 | 0.0 | 0.0 | 0.0 | 0.1 | 4.6 | 0.3 | 0.0 | 0.0 | 0.0 |
| 74 | 0.0 | 0.0 | 0.4 | 0.0 | 0.0 | 0.0 | 0.0 | 0.0 | 0.0 | 0.3 | 0.7 | 0.0 | 0.0 | 0.1 |
| 75 | 0.0 | 0.0 | 0.4 | 0.0 | 0.0 | 0.0 | 0.0 | 0.0 | 0.0 | 1.4 | 0.3 | 0.0 | 0.0 | 0.0 |
| 76 | 0.0 | 0.0 | 0.4 | 0.0 | 0.0 | 0.0 | 0.0 | 0.0 | 0.0 | 0.7 | 0.0 | 0.0 | 0.0 | 0.0 |
| 77 | 0.0 | 0.0 | 0.2 | 0.0 | 0.0 | 0.1 | 0.0 | 0.0 | 0.0 | 0.3 | 0.7 | 0.1 | 0.0 | 0.0 |
| 78 | 0.1 | 0.0 | 0.4 | 0.0 | 0.0 | 0.0 | 0.0 | 0.0 | 0.1 | 0.3 | 0.2 | 0.0 | 0.0 | 0.0 |
| 79 | 0.0 | 0.0 | 0.4 | 0.0 | 0.2 | 0.0 | 0.0 | 0.0 | 0.0 | 0.5 | 0.3 | 0.0 | 0.0 | 0.1 |
| 80 | 0.0 | 0.0 | 0.9 | 0.0 | 0.0 | 0.0 | 0.0 | 0.0 | 0.1 | 0.3 | 2.2 | 0.5 | 0.0 | 0.0 |
| 81 | 2.8 | 0.0 | 1.2 | 0.0 | 0.0 | 0.0 | 0.0 | 0.0 | 0.0 | 0.7 | 0.6 | 0.0 | 0.2 | 0.0 |
| 82 | 58.0 | 0.0 | 7.7 | 0.0 | 0.6 | 0.0 | 0.0 | 0.0 | 0.0 | 0.3 | 3.2 | 0.0 | 0.0 | 0.1 |
| 83 | 0.0 | 0.0 | 0.0 | 0.0 | 0.0 | 0.0 | 0.0 | 0.0 | 0.0 | 1.0 | 0.0 | 0.0 | 0.0 | 0.0 |
| 84 | 0.1 | 0.0 | 0.1 | 0.0 | 0.0 | 0.0 | 0.0 | 0.0 | 0.0 | 0.5 | 0.1 | 0.0 | 0.0 | 0.0 |
| 85 | 0.0 | 0.0 | 0.4 | 0.0 | 0.0 | 0.0 | 0.0 | 0.1 | 0.0 | 0.1 | 0.3 | 0.0 | 0.0 | 0.0 |
| 86 | 0.0 | 0.0 | 0.0 | 0.0 | 0.0 | 0.0 | 0.0 | 0.1 | 0.0 | 1.0 | 0.2 | 0.0 | 0.0 | 0.0 |
| 87 | 0.0 | 0.0 | 0.9 | 0.0 | 0.0 | 0.0 | 0.0 | 0.0 | 0.0 | 0.1 | 0.3 | 0.0 | 0.0 | 0.1 |
| 88 | 0.0 | 0.0 | 0.6 | 0.0 | 0.0 | 0.0 | 0.0 | 0.0 | 0.0 | 2.8 | 0.0 | 0.0 | 0.0 | 0.0 |
| 89 | 3.4 | 0.0 | 0.2 | 0.0 | 0.0 | 0.0 | 0.0 | 0.0 | 0.1 | 14.5 | 0.0 | 0.0 | 0.0 | 0.1 |
| 90 | 0.1 | 0.0 | 0.2 | 0.0 | 0.0 | 0.0 | 0.0 | 0.0 | 0.0 | 0.5 | 0.3 | 0.0 | 0.0 | 0.0 |
| 91 | 9.4 | 0.0 | 1.2 | 0.0 | 0.0 | 0.0 | 0.0 | 0.0 | 0.0 | 1.0 | 0.7 | 0.0 | 0.0 | 0.1 |
| 92 | 0.0 | 0.0 | 0.0 | 0.0 | 0.0 | 0.0 | 0.0 | 0.0 | 0.0 | 0.5 | 0.4 | 0.0 | 0.0 | 0.0 |
| 93 | 1.0 | 0.0 | 0.0 | 0.0 | 0.0 | 0.0 | 0.0 | 0.0 | 0.1 | 2.8 | 0.0 | 0.0 | 0.0 | 0.0 |
| 94 | 0.1 | 0.0 | 0.1 | 0.0 | 0.0 | 0.0 | 0.0 | 0.2 | 0.1 | 1.0 | 0.2 | 0.0 | 0.1 | 0.1 |
| 95 | 17.1 | 0.0 | 0.1 | 0.0 | 0.0 | 0.0 | 0.0 | 0.0 | 0.0 | 1.0 | 0.9 | 0.0 | 0.0 | 0.1 |
| 96 | 0.1 | 0.0 | 0.1 | 0.0 | 0.0 | 0.0 | 0.0 | 0.0 | 0.2 | 0.3 | 0.3 | 0.1 | 0.0 | 0.0 |
| 97 | 0.0 | 0.0 | 0.1 | 0.0 | 0.0 | 0.0 | 0.0 | 0.0 | 0.0 | 0.7 | 0.3 | 0.0 | 0.1 | 0.4 |
| 98 | 0.3 | 0.0 | 0.1 | 0.0 | 0.0 | 0.0 | 0.0 | 0.0 | 0.0 | 0.5 | 0.3 | 0.0 | 0.0 | 0.1 |
| 99 | 0.0 | 0.0 | 0.0 | 0.0 | 0.0 | 0.0 | 0.0 | 0.0 | 0.0 | 0.7 | 0.1 | 0.0 | 0.0 | 0.1 |
| 100 | 9.7 | 0.1 | 26.1 | 0.1 | 0.3 | 1.1 | 0.1 | 0.1 | 0.1 | 0.7 | 14.0 | 0.0 | 0.3 | 0.2 |

**Figure S13: GO term enrichment p-values.** This figure is an extended version of **Figure 4B**, now showing $-\log_{10}$ p-values for GO term enrichments for all states. The criteria for including a GO term were the same as in **Figure 4B**, except the top 10 term criterion was relaxed to include four additional terms. Values that are boxed mark the top most significant enrichment for the term of the column.

| | DNA class | LINE class | LTR class | Low complexity class | Other class | RC class | RNA class | SINE class | Satellite class | Simple class | Unknown class | rRNA class | scRNA class | snRNA class | srpRNA class | tRNA class | Alu family | CR1 family | DNA family | Deu family |
|---|---|---|---|---|---|---|---|---|---|---|---|---|---|---|---|---|---|---|---|---|
| 1 | 0.00 | 0.00 | 0.00 | 0.35 | 0.00 | 0.00 | 0.00 | 0.00 | 0.00 | 0.12 | 0.00 | 0.09 | 0.00 | 0.04 | 0.00 | 0.16 | 0.00 | 0.05 | 0.00 | 0.00 |
| 2 | 0.05 | 0.00 | 0.00 | 0.90 | 0.00 | 0.01 | 0.01 | 0.01 | 0.00 | 0.16 | 1.38 | 0.04 | 0.00 | 0.05 | 0.00 | 0.05 | 0.00 | 0.16 | 0.05 | 0.68 |
| 3 | 0.02 | 0.00 | 0.00 | 0.77 | 0.00 | 0.01 | 0.01 | 0.01 | 0.00 | 0.15 | 0.73 | 0.07 | 0.00 | 0.02 | 0.00 | 0.03 | 0.00 | 0.11 | 0.02 | 0.48 |
| 4 | 0.05 | 0.01 | 0.00 | 0.92 | 0.00 | 0.00 | 0.04 | 0.02 | 0.01 | 0.19 | 1.76 | 0.00 | 0.00 | 0.05 | 0.08 | 0.11 | 0.00 | 0.22 | 0.05 | 1.09 |
| 5 | 0.17 | 0.03 | 0.00 | 0.80 | 0.00 | 0.24 | 0.17 | 0.07 | 0.00 | 0.15 | 4.82 | 0.09 | 0.02 | 0.05 | 0.00 | 2.51 | 0.00 | 1.02 | 0.17 | 20.37 |
| 6 | 0.81 | 0.31 | 0.18 | 0.67 | 0.00 | 1.84 | 1.64 | 0.45 | 0.00 | 0.17 | 2.81 | 0.25 | 0.00 | 0.17 | 0.05 | 4.33 | 0.00 | 2.51 | 0.81 | 4.35 |
| 7 | 0.17 | 0.07 | 0.01 | 1.58 | 0.00 | 0.55 | 0.11 | 0.12 | 0.01 | 0.31 | 0.68 | 0.05 | 0.00 | 0.08 | 0.08 | 0.46 | 0.00 | 0.40 | 0.17 | 0.58 |
| 8 | 0.10 | 0.03 | 0.00 | 0.93 | 0.00 | 0.21 | 0.39 | 0.07 | 0.01 | 0.17 | 1.96 | 0.02 | 0.01 | 0.02 | 0.01 | 1.71 | 0.00 | 0.69 | 0.10 | 9.14 |
| 9 | 0.81 | 0.32 | 0.20 | 0.60 | 0.00 | 1.63 | 1.53 | 0.47 | 0.00 | 0.16 | 2.12 | 0.24 | 0.00 | 0.09 | 0.02 | 3.13 | 0.00 | 2.23 | 0.81 | 2.89 |
| 10 | 0.83 | 0.36 | 0.28 | 0.72 | 0.00 | 1.53 | 1.27 | 0.45 | 0.01 | 0.26 | 1.73 | 0.23 | 0.00 | 0.09 | 0.02 | 2.12 | 0.00 | 1.91 | 0.83 | 2.45 |
| 11 | 0.93 | 0.47 | 0.41 | 0.99 | 0.00 | 1.59 | 0.96 | 0.46 | 0.01 | 0.46 | 1.70 | 0.14 | 0.00 | 0.08 | 0.02 | 1.49 | 0.00 | 1.91 | 0.93 | 1.32 |
| 12 | 0.91 | 0.46 | 0.39 | 0.84 | 0.00 | 1.50 | 1.27 | 0.43 | 0.01 | 0.30 | 2.01 | 0.22 | 0.00 | 0.09 | 0.02 | 1.86 | 0.00 | 2.04 | 0.91 | 1.24 |
| 13 | 0.97 | 0.44 | 0.35 | 0.77 | 0.00 | 2.19 | 1.26 | 0.45 | 0.01 | 0.21 | 1.99 | 0.32 | 0.00 | 0.07 | 0.00 | 1.86 | 0.00 | 2.18 | 0.97 | 1.32 |
| 14 | 0.87 | 0.40 | 0.29 | 0.76 | 0.00 | 1.53 | 1.14 | 0.47 | 0.01 | 0.26 | 1.51 | 0.22 | 0.00 | 0.13 | 0.02 | 1.88 | 0.00 | 1.95 | 0.87 | 2.21 |
| 15 | 0.78 | 0.31 | 0.21 | 0.64 | 0.00 | 1.48 | 1.46 | 0.46 | 0.01 | 0.20 | 1.80 | 0.23 | 0.00 | 0.07 | 0.02 | 2.74 | 0.00 | 2.01 | 0.78 | 2.24 |
| 16 | 1.24 | 0.66 | 0.64 | 0.90 | 0.00 | 2.13 | 1.17 | 0.50 | 0.00 | 0.40 | 1.76 | 0.38 | 0.00 | 0.17 | 0.02 | 1.20 | 0.00 | 2.15 | 1.24 | 0.47 |
| 17 | 1.28 | 0.66 | 0.61 | 0.99 | 0.00 | 2.44 | 1.15 | 0.51 | 0.00 | 0.39 | 2.00 | 0.40 | 0.00 | 0.17 | 0.03 | 1.58 | 0.00 | 2.32 | 1.28 | 0.67 |
| 18 | 1.15 | 0.59 | 0.57 | 0.84 | 0.00 | 1.99 | 1.22 | 0.49 | 0.01 | 0.42 | 1.70 | 0.33 | 0.00 | 0.13 | 0.02 | 1.25 | 0.00 | 2.04 | 1.15 | 0.59 |
| 19 | 0.63 | 0.24 | 0.15 | 0.65 | 0.00 | 1.23 | 1.27 | 0.37 | 0.01 | 0.18 | 2.00 | 0.16 | 0.01 | 0.06 | 0.02 | 2.81 | 0.00 | 1.75 | 0.63 | 4.53 |
| 20 | 0.98 | 0.46 | 0.37 | 1.00 | 0.00 | 1.99 | 0.87 | 0.47 | 0.01 | 0.31 | 1.79 | 0.24 | 0.00 | 0.05 | 0.00 | 1.81 | 0.00 | 2.09 | 0.98 | 1.89 |
| 21 | 0.89 | 0.43 | 0.38 | 0.98 | 0.00 | 1.89 | 1.14 | 0.42 | 0.01 | 0.42 | 2.15 | 0.25 | 0.01 | 0.04 | 0.01 | 1.86 | 0.00 | 1.93 | 0.89 | 1.84 |
| 22 | 1.15 | 0.66 | 0.59 | 1.02 | 0.00 | 2.07 | 0.93 | 0.47 | 0.02 | 0.53 | 1.81 | 0.17 | 0.00 | 0.17 | 0.01 | 1.13 | 0.01 | 1.90 | 1.15 | 0.46 |
| 23 | 1.45 | 1.16 | 0.94 | 1.19 | 0.00 | 2.21 | 0.70 | 0.42 | 0.00 | 0.61 | 1.50 | 0.50 | 0.08 | 0.25 | 0.05 | 0.63 | 0.04 | 1.72 | 1.45 | 0.61 |
| 24 | 1.33 | 1.00 | 0.92 | 1.25 | 0.00 | 1.70 | 0.85 | 0.40 | 0.00 | 0.71 | 1.49 | 0.44 | 0.08 | 0.11 | 0.04 | 0.51 | 0.03 | 1.64 | 1.33 | 0.22 |
| 25 | 1.48 | 1.19 | 1.37 | 1.11 | 0.00 | 1.92 | 0.93 | 0.40 | 0.00 | 0.71 | 1.40 | 0.32 | 0.02 | 0.09 | 0.01 | 0.69 | 0.01 | 1.87 | 1.48 | 0.30 |
| 26 | 1.36 | 0.76 | 0.60 | 0.96 | 0.00 | 2.23 | 1.40 | 0.48 | 0.00 | 0.29 | 1.88 | 0.49 | 0.00 | 0.09 | 0.02 | 0.99 | 0.01 | 2.19 | 1.36 | 0.70 |
| 27 | 1.21 | 0.66 | 0.60 | 0.87 | 0.00 | 2.03 | 1.13 | 0.46 | 0.00 | 0.35 | 1.73 | 0.42 | 0.01 | 0.11 | 0.02 | 1.20 | 0.00 | 1.93 | 1.21 | 0.55 |
| 28 | 0.13 | 0.08 | 0.04 | 7.05 | 0.00 | 0.07 | 0.00 | 0.10 | 0.23 | 2.45 | 0.13 | 0.15 | 0.08 | 0.21 | 0.05 | 0.40 | 0.04 | 0.15 | 0.13 | 0.40 |
| 29 | 1.27 | 0.95 | 0.96 | 1.06 | 0.00 | 1.96 | 0.69 | 0.41 | 0.01 | 0.65 | 1.41 | 0.17 | 0.02 | 0.13 | 0.05 | 0.95 | 0.01 | 1.91 | 1.27 | 0.22 |
| 30 | 1.35 | 1.07 | 0.94 | 1.05 | 0.00 | 2.45 | 0.59 | 0.38 | 0.00 | 0.38 | 1.69 | 0.30 | 0.02 | 0.09 | 0.04 | 0.69 | 0.01 | 1.70 | 1.35 | 0.32 |
| 31 | 1.30 | 1.00 | 0.79 | 1.11 | 0.00 | 2.08 | 0.38 | 0.39 | 0.01 | 0.49 | 1.16 | 0.41 | 0.06 | 0.11 | 0.05 | 0.87 | 0.03 | 1.62 | 1.30 | 0.10 |
| 32 | 1.37 | 2.01 | 1.26 | 1.09 | 0.00 | 1.22 | 0.38 | 0.38 | 0.01 | 0.88 | 0.60 | 0.28 | 0.38 | 0.46 | 0.19 | 0.95 | 0.24 | 0.93 | 1.37 | 0.06 |
| 33 | 1.22 | 1.46 | 0.88 | 1.97 | 0.03 | 1.87 | 0.28 | 0.37 | 0.04 | 2.28 | 1.01 | 0.80 | 0.10 | 0.27 | 0.16 | 0.28 | 0.15 | 1.28 | 1.22 | 0.14 |
| 34 | 1.05 | 0.69 | 0.56 | 1.41 | 0.00 | 1.59 | 1.14 | 0.42 | 0.01 | 0.99 | 1.50 | 0.38 | 0.01 | 0.32 | 0.03 | 1.13 | 0.02 | 1.86 | 1.05 | 0.41 |
| 35 | 1.10 | 0.58 | 0.45 | 1.05 | 0.00 | 1.76 | 1.02 | 0.46 | 0.01 | 0.39 | 1.54 | 0.27 | 0.01 | 0.14 | 0.04 | 1.55 | 0.01 | 1.83 | 1.10 | 0.73 |
| 36 | 1.49 | 1.05 | 1.07 | 1.11 | 0.00 | 2.55 | 0.64 | 0.43 | 0.00 | 0.46 | 1.77 | 0.70 | 0.15 | 0.24 | 0.02 | 0.79 | 0.01 | 1.90 | 1.49 | 0.30 |
| 37 | 1.10 | 0.50 | 0.38 | 0.81 | 0.00 | 2.23 | 1.11 | 0.49 | 0.00 | 0.20 | 2.21 | 0.31 | 0.03 | 0.12 | 0.02 | 1.95 | 0.00 | 2.21 | 1.10 | 1.24 |
| 38 | 1.25 | 0.70 | 0.54 | 0.94 | 0.00 | 2.25 | 1.17 | 0.45 | 0.00 | 0.24 | 1.97 | 0.30 | 0.02 | 0.08 | 0.13 | 1.26 | 0.00 | 2.12 | 1.25 | 0.52 |
| 39 | 1.21 | 0.76 | 0.60 | 0.99 | 0.00 | 2.26 | 0.83 | 0.40 | 0.00 | 0.31 | 1.88 | 0.47 | 0.01 | 0.15 | 0.04 | 0.96 | 0.01 | 2.02 | 1.21 | 0.69 |
| 40 | 1.41 | 1.35 | 1.29 | 1.06 | 0.00 | 2.18 | 1.52 | 0.33 | 0.00 | 0.48 | 1.34 | 0.56 | 0.06 | 0.25 | 0.03 | 0.51 | 0.02 | 1.60 | 1.41 | 0.34 |
| 41 | 1.32 | 0.95 | 1.08 | 1.04 | 0.00 | 2.10 | 1.03 | 0.42 | 0.00 | 0.59 | 1.63 | 0.21 | 0.03 | 0.21 | 0.04 | 0.75 | 0.01 | 1.91 | 1.32 | 0.15 |
| 42 | 1.27 | 0.86 | 0.80 | 1.00 | 0.00 | 1.94 | 0.93 | 0.45 | 0.02 | 0.61 | 1.66 | 0.29 | 0.02 | 0.21 | 0.04 | 0.89 | 0.01 | 1.89 | 1.27 | 0.28 |
| 43 | 1.26 | 0.93 | 0.82 | 1.02 | 0.00 | 1.91 | 1.02 | 0.40 | 0.00 | 0.42 | 1.43 | 0.49 | 0.03 | 0.17 | 0.03 | 1.87 | 0.01 | 1.65 | 1.26 | 0.13 |
| 44 | 1.40 | 1.60 | 0.98 | 1.47 | 0.01 | 1.39 | 0.36 | 0.37 | 0.01 | 1.31 | 1.04 | 0.67 | 0.32 | 0.53 | 0.20 | 1.05 | 0.15 | 1.13 | 1.40 | 0.16 |
| 45 | 1.27 | 1.09 | 0.93 | 1.34 | 0.00 | 1.63 | 0.83 | 0.39 | 0.01 | 0.89 | 1.26 | 0.34 | 0.10 | 0.33 | 0.24 | 0.79 | 0.06 | 1.66 | 1.27 | 0.06 |
| 46 | 1.36 | 0.95 | 0.80 | 1.17 | 0.00 | 1.67 | 0.58 | 0.45 | 0.00 | 0.62 | 1.46 | 0.42 | 0.08 | 0.20 | 0.02 | 0.72 | 0.03 | 1.89 | 1.36 | 0.30 |
| 47 | 1.03 | 0.53 | 0.50 | 1.18 | 0.00 | 1.57 | 0.99 | 0.42 | 0.01 | 0.57 | 1.32 | 0.30 | 0.01 | 0.13 | 0.01 | 1.05 | 0.00 | 1.67 | 1.03 | 0.60 |
| 48 | 1.24 | 0.81 | 0.79 | 0.98 | 0.00 | 2.03 | 0.69 | 0.39 | 0.00 | 0.40 | 1.52 | 0.54 | 0.04 | 0.20 | 0.03 | 0.63 | 0.01 | 1.71 | 1.24 | 0.29 |
| 49 | 1.00 | 0.48 | 0.40 | 0.84 | 0.00 | 1.76 | 0.98 | 0.44 | 0.00 | 0.26 | 1.52 | 0.25 | 0.02 | 0.07 | 0.03 | 1.23 | 0.00 | 1.78 | 1.00 | 0.72 |
| 50 | 0.55 | 0.21 | 0.14 | 0.67 | 0.00 | 1.06 | 0.95 | 0.33 | 0.01 | 0.16 | 1.66 | 0.14 | 0.01 | 0.06 | 0.02 | 2.10 | 0.00 | 1.56 | 0.55 | 3.65 |
| 51 | 1.03 | 0.54 | 0.52 | 0.75 | 0.00 | 1.77 | 1.04 | 0.45 | 0.00 | 0.30 | 1.52 | 0.30 | 0.02 | 0.11 | 0.01 | 1.06 | 0.00 | 1.88 | 1.03 | 0.56 |
| 52 | 1.06 | 0.54 | 0.53 | 1.10 | 0.00 | 1.66 | 1.05 | 0.45 | 0.01 | 0.53 | 1.45 | 0.28 | 0.00 | 0.11 | 0.01 | 1.06 | 0.00 | 1.82 | 1.06 | 0.53 |
| 53 | 0.58 | 0.22 | 0.14 | 0.93 | 0.00 | 1.01 | 1.06 | 0.34 | 0.01 | 0.27 | 1.53 | 0.12 | 0.00 | 0.07 | 0.01 | 2.17 | 0.00 | 1.48 | 0.58 | 3.13 |
| 54 | 0.04 | 0.02 | 0.00 | 1.10 | 0.00 | 0.08 | 0.02 | 0.03 | 0.01 | 0.26 | 0.47 | 0.04 | 0.00 | 0.03 | 0.01 | 0.22 | 0.00 | 0.14 | 0.04 | 0.40 |
| 55 | 0.56 | 0.21 | 0.14 | 0.60 | 0.00 | 1.12 | 0.89 | 0.33 | 0.01 | 0.18 | 1.83 | 0.13 | 0.00 | 0.06 | 0.02 | 2.25 | 0.00 | 1.54 | 0.56 | 4.02 |
| 56 | 1.04 | 0.57 | 0.57 | 0.76 | 0.00 | 1.80 | 1.02 | 0.44 | 0.01 | 0.47 | 1.54 | 0.23 | 0.00 | 0.12 | 0.02 | 1.03 | 0.00 | 1.87 | 1.04 | 0.57 |
| 57 | 1.18 | 0.76 | 0.70 | 0.77 | 0.00 | 1.94 | 0.84 | 0.40 | 0.00 | 0.39 | 1.44 | 0.40 | 0.01 | 0.17 | 0.03 | 0.82 | 0.01 | 1.68 | 1.18 | 0.37 |
| 58 | 1.10 | 0.79 | 0.74 | 1.26 | 0.00 | 1.31 | 0.75 | 0.39 | 0.01 | 0.80 | 1.17 | 0.31 | 0.08 | 0.24 | 0.08 | 0.69 | 0.04 | 1.55 | 1.10 | 0.30 |
| 59 | 1.17 | 1.10 | 0.87 | 1.38 | 0.01 | 1.34 | 0.56 | 0.36 | 0.01 | 1.12 | 0.93 | 0.67 | 0.43 | 0.43 | 0.15 | 0.61 | 0.10 | 1.14 | 1.17 | 0.10 |
| 60 | 1.24 | 1.08 | 1.22 | 1.08 | 0.00 | 1.80 | 0.90 | 0.33 | 0.00 | 0.70 | 1.11 | 0.32 | 0.12 | 0.28 | 0.06 | 0.38 | 0.02 | 1.43 | 1.24 | 0.18 |
| 61 | 1.18 | 0.91 | 0.90 | 1.01 | 0.00 | 1.69 | 0.53 | 0.36 | 0.00 | 0.54 | 1.22 | 0.30 | 0.02 | 0.12 | 0.06 | 0.57 | 0.02 | 1.49 | 1.18 | 0.15 |
| 62 | 1.27 | 0.93 | 0.91 | 1.12 | 0.00 | 1.83 | 0.78 | 0.38 | 0.00 | 0.60 | 1.28 | 0.44 | 0.13 | 0.16 | 0.05 | 0.50 | 0.03 | 1.57 | 1.27 | 0.36 |
| 63 | 0.92 | 0.47 | 0.47 | 1.07 | 0.00 | 1.62 | 0.89 | 0.41 | 0.01 | 0.45 | 1.43 | 0.16 | 0.01 | 0.08 | 0.00 | 1.15 | 0.00 | 1.69 | 0.92 | 0.88 |
| 64 | 1.26 | 0.97 | 0.91 | 1.32 | 0.00 | 1.40 | 0.62 | 0.42 | 0.01 | 0.88 | 1.34 | 0.54 | 0.09 | 0.17 | 0.07 | 0.63 | 0.05 | 1.74 | 1.26 | 0.14 |
| 65 | 1.16 | 0.91 | 0.75 | 1.90 | 0.00 | 1.17 | 0.42 | 0.55 | 0.02 | 1.44 | 1.14 | 1.01 | 0.88 | 1.07 | 0.31 | 0.67 | 0.31 | 1.25 | 1.16 | 0.13 |
| 66 | 0.93 | 0.54 | 0.53 | 1.60 | 0.00 | 1.66 | 0.77 | 0.39 | 0.03 | 1.16 | 1.32 | 0.22 | 0.10 | 0.17 | 0.04 | 0.75 | 0.02 | 1.52 | 0.93 | 0.34 |
| 67 | 0.70 | 1.08 | 2.33 | 2.33 | 0.00 | 0.98 | 0.84 | 0.55 | 0.02 | 2.64 | 0.75 | 1.50 | 0.94 | 1.38 | 0.37 | 0.70 | 0.48 | 0.74 | 0.70 | 0.15 |
| 68 | 0.84 | 1.53 | 2.24 | 1.46 | 0.00 | 0.71 | 1.56 | 1.04 | 0.00 | 1.39 | 0.42 | 0.90 | 1.21 | 1.45 | 0.82 | 0.38 | 1.16 | 0.62 | 0.84 | 0.04 |
| 69 | 1.28 | 1.80 | 1.20 | 1.40 | 0.00 | 1.54 | 0.38 | 0.62 | 0.02 | 1.33 | 0.77 | 0.88 | 0.99 | 0.84 | 0.58 | 0.49 | 0.55 | 0.87 | 1.28 | 0.06 |
| 70 | 1.19 | 1.61 | 1.16 | 1.33 | 0.00 | 1.11 | 0.44 | 0.60 | 0.01 | 1.34 | 0.53 | 0.90 | 1.31 | 0.98 | 0.54 | 0.38 | 0.54 | 0.73 | 1.19 | 0.07 |
| 71 | 1.23 | 1.24 | 1.00 | 1.46 | 0.00 | 1.08 | 0.55 | 0.85 | 0.01 | 1.27 | 0.70 | 1.25 | 1.39 | 1.21 | 0.98 | 0.42 | 0.81 | 0.92 | 1.23 | 0.08 |
| 72 | 1.36 | 1.67 | 1.06 | 1.44 | 0.00 | 1.29 | 0.33 | 0.78 | 0.01 | 1.23 | 0.78 | 0.91 | 0.92 | 0.93 | 0.81 | 0.54 | 0.74 | 0.92 | 1.36 | 0.05 |
| 73 | 1.26 | 1.21 | 0.98 | 1.54 | 0.00 | 0.93 | 1.51 | 2.52 | 0.01 | 1.94 | 0.31 | 2.54 | 3.49 | 3.75 | 3.16 | 0.97 | 3.07 | 0.55 | 1.26 | 0.01 |
| 74 | 5.57 | 1.02 | 1.78 | 1.02 | 0.00 | 0.55 | 0.83 | 1.80 | 0.00 | 0.95 | 0.23 | 2.47 | 3.60 | 3.66 | 5.27 | 0.69 | 2.20 | 0.30 | 5.57 | 0.11 |
| 75 | 1.05 | 0.97 | 1.44 | 1.01 | 0.00 | 0.42 | 4.47 | 3.71 | 0.01 | 1.01 | 0.18 | 4.07 | 8.12 | 6.29 | 6.21 | 0.86 | 4.67 | 0.23 | 1.05 | 0.04 |
| 76 | 1.99 | 1.53 | 3.31 | 0.39 | 0.00 | 0.08 | 3.02 | 2.14 | 0.02 | 0.45 | 0.07 | 2.78 | 3.90 | 3.74 | 5.13 | 0.57 | 2.71 | 0.08 | 1.99 | 0.02 |
| 77 | 1.15 | 0.93 | 1.15 | 0.80 | 0.02 | 0.17 | 1.20 | 4.52 | 0.05 | 1.29 | 0.07 | 2.15 | 3.21 | 2.49 | 4.19 | 0.38 | 5.73 | 0.12 | 1.15 | 0.08 |
| 78 | 1.88 | 1.62 | 2.20 | 0.81 | 0.00 | 0.17 | 2.71 | 2.47 | 0.03 | 0.98 | 0.09 | 2.66 | 5.20 | 4.21 | 4.92 | 0.67 | 3.12 | 0.10 | 1.88 | 0.04 |
| 79 | 1.50 | 1.86 | 2.65 | 0.52 | 0.05 | 0.12 | 1.15 | 2.20 | 0.49 | 0.74 | 0.08 | 1.77 | 2.19 | 2.22 | 2.56 | 0.48 | 2.80 | 0.07 | 1.50 | 0.05 |
| 80 | 0.66 | 1.14 | 0.88 | 1.61 | 0.02 | 0.39 | 1.42 | 3.86 | 0.02 | 2.44 | 0.19 | 1.95 | 2.52 | 2.09 | 1.34 | 0.40 | 4.86 | 0.24 | 0.66 | 0.03 |
| 81 | 0.90 | 1.60 | 1.31 | 1.38 | 0.02 | 0.29 | 1.38 | 2.79 | 0.34 | 2.28 | 0.14 | 3.05 | 2.68 | 2.50 | 1.74 | 0.77 | 3.52 | 0.17 | 0.90 | 0.01 |
| 82 | 0.37 | 0.23 | 0.30 | 4.31 | 0.13 | 0.04 | 0.05 | 0.66 | 0.57 | 4.38 | 0.04 | 3.16 | 2.85 | 2.51 | 2.58 | 1.14 | 0.82 | 0.06 | 0.37 | 0.02 |
| 83 | 1.51 | 2.31 | 2.17 | 0.46 | 0.01 | 0.10 | 0.86 | 1.74 | 0.28 | 0.69 | 0.03 | 2.12 | 1.76 | 2.64 | 2.75 | 0.30 | 2.20 | 0.05 | 1.51 | 0.00 |
| 84 | 0.80 | 1.71 | 0.83 | 1.56 | 0.01 | 0.36 | 1.20 | 2.30 | 0.08 | 2.22 | 0.13 | 1.16 | 2.27 | 2.80 | 2.46 | 1.24 | 2.85 | 0.25 | 0.80 | 0.00 |
| 85 | 0.78 | 1.69 | 1.50 | 0.97 | 0.07 | 0.01 | 0.32 | 3.33 | 1.24 | 1.70 | 0.06 | 1.36 | 2.62 | 1.62 | 2.61 | 0.31 | 4.23 | 0.08 | 0.78 | 0.00 |
| 86 | 0.56 | 2.95 | 1.52 | 0.71 | 0.10 | 0.10 | 0.47 | 1.61 | 0.66 | 1.15 | 0.01 | 0.51 | 1.23 | 0.91 | 1.02 | 0.18 | 2.05 | 0.02 | 0.56 | 0.01 |
| 87 | 1.01 | 1.33 | 1.69 | 1.33 | 0.06 | 0.14 | 1.38 | 3.34 | 0.38 | 2.20 | 0.08 | 2.05 | 2.25 | 2.22 | 2.27 | 0.50 | 4.24 | 0.10 | 1.01 | 0.03 |
| 88 | 1.94 | 1.08 | 2.09 | 0.81 | 0.00 | 0.24 | 3.15 | 3.09 | 0.02 | 1.10 | 0.12 | 3.30 | 3.95 | 3.86 | 4.88 | 0.60 | 3.89 | 0.16 | 1.94 | 0.04 |
| 89 | 1.07 | 1.16 | 1.09 | 1.42 | 0.00 | 1.27 | 0.57 | 0.63 | 0.03 | 1.59 | 0.73 | 0.72 | 0.59 | 0.69 | 0.41 | 0.53 | 0.52 | 0.95 | 1.07 | 0.09 |
| 90 | 1.53 | 1.91 | 2.11 | 0.96 | 0.00 | 0.09 | 1.48 | 2.04 | 0.23 | 1.79 | 0.07 | 1.42 | 2.48 | 2.80 | 2.55 | 0.46 | 2.57 | 0.11 | 1.53 | 0.08 |
| 91 | 0.70 | 1.53 | 0.70 | 2.80 | 0.20 | 0.38 | 0.32 | 2.09 | 0.07 | 5.87 | 0.24 | 1.08 | 1.69 | 1.68 | 0.74 | 0.71 | 2.55 | 0.39 | 0.70 | 0.06 |
| 92 | 0.57 | 2.41 | 2.21 | 0.85 | 0.07 | 0.08 | 0.90 | 1.78 | 0.94 | 1.62 | 0.04 | 1.30 | 0.94 | 1.27 | 1.04 | 0.37 | 2.25 | 0.06 | 0.57 | 0.02 |
| 93 | 0.54 | 2.46 | 2.53 | 0.82 | 0.15 | 0.09 | 0.14 | 1.44 | 2.10 | 1.70 | 0.08 | 3.35 | 0.72 | 1.51 | 1.11 | 0.46 | 1.82 | 0.04 | 0.54 | 0.00 |
| 94 | 0.34 | 2.92 | 1.99 | 0.84 | 16.38 | 0.05 | 1.05 | 0.78 | 4.07 | 2.34 | 0.20 | 3.88 | 0.67 | 0.81 | 0.51 | 0.42 | 0.97 | 0.08 | 0.34 | 0.01 |
| 95 | 0.35 | 1.65 | 1.30 | 3.77 | 4.96 | 0.20 | 0.42 | 1.44 | 4.54 | 9.49 | 0.06 | 6.24 | 0.83 | 1.29 | 1.38 | 0.38 | 1.80 | 0.13 | 0.35 | 0.07 |
| 96 | 0.02 | 0.30 | 0.23 | 0.16 | 7.20 | 0.00 | 0.04 | 0.10 | 7.52 | 0.77 | 0.01 | 0.22 | 0.06 | 0.09 | 0.02 | 0.02 | 0.13 | 0.01 | 0.02 | 0.00 |
| 97 | 0.24 | 2.78 | 1.83 | 0.79 | 16.75 | 0.04 | 0.09 | 0.84 | 10.79 | 3.26 | 0.16 | 2.20 | 0.86 | 0.43 | 0.21 | 0.11 | 1.06 | 0.08 | 0.24 | 0.00 |
| 98 | 0.38 | 2.37 | 2.04 | 1.17 | 0.37 | 0.07 | 0.00 | 1.95 | 4.32 | 2.81 | 0.00 | 2.60 | 0.89 | 0.99 | 1.57 | 0.14 | 2.47 | 0.05 | 0.38 | 0.00 |
| 99 | 0.41 | 2.39 | 1.54 | 1.06 | 0.07 | 0.00 | 0.30 | 2.26 | 1.82 | 2.55 | 0.04 | 0.50 | 1.01 | 0.83 | 1.04 | 0.22 | 2.87 | 0.03 | 0.41 | 0.00 |
| 100 | 0.00 | 0.01 | 0.01 | 0.74 | 0.00 | 0.00 | 0.00 | 0.01 | 0.00 | 0.44 | 0.00 | 2.51 | 0.00 | 0.14 | 0.00 | 5.39 | 0.01 | 0.03 | 0.00 | 0.00 |
| Base | 3.18 | 20.37 | 8.52 | 0.54 | 0.13 | 0.01 | 0.00 | 12.65 | 0.40 | 0.84 | 0.04 | 0.01 | 0.00 | 0.01 | 0.01 | 0.01 | 9.91 | 0.35 | 3.18 | 0.01 |

| | Dong-R4 family | ERV1 family | ERVK family | ERVL-MaLR family | ERVL family | ERV family | Gypsy family | Helitron family | L1 family | L2 family | LTR family | Low complexity family | MIR family | Merlin family | MuDR family | Other family | PiggyBac family | RNA family | RTE-BovB family | RTE family |
|---|---|---|---|---|---|---|---|---|---|---|---|---|---|---|---|---|---|---|---|---|
| 1 | 0.00 | 0.00 | 0.00 | 0.00 | 0.00 | 0.00 | 0.00 | 0.00 | 0.00 | 0.00 | 0.00 | 0.35 | 0.00 | 0.61 | 0.00 | 0.00 | 0.00 | 0.00 | 0.14 | 0.07 |
| 2 | 0.00 | 0.00 | 0.00 | 0.00 | 0.00 | 0.00 | 0.00 | 0.01 | 0.00 | 0.00 | 0.00 | 0.90 | 0.01 | 0.43 | 0.00 | 0.00 | 0.10 | 0.01 | 0.42 | 0.08 |
| 3 | 0.00 | 0.00 | 0.00 | 0.00 | 0.00 | 0.00 | 0.00 | 0.01 | 0.00 | 0.01 | 0.00 | 0.77 | 0.01 | 0.20 | 0.00 | 0.00 | 0.08 | 0.01 | 0.30 | 0.07 |
| 4 | 0.07 | 0.00 | 0.00 | 0.00 | 0.00 | 0.00 | 0.00 | 0.00 | 0.00 | 0.02 | 0.00 | 0.92 | 0.04 | 0.00 | 0.00 | 0.00 | 0.11 | 0.04 | 0.60 | 0.08 |
| 5 | 2.28 | 0.00 | 0.00 | 0.00 | 0.00 | 0.00 | 0.01 | 0.24 | 0.00 | 0.05 | 0.00 | 0.80 | 0.24 | 0.00 | 0.00 | 0.00 | 1.58 | 0.17 | 4.61 | 0.27 |
| 6 | 7.21 | 0.03 | 0.00 | 0.24 | 0.24 | 0.16 | 1.25 | 1.84 | 0.10 | 1.06 | 0.18 | 0.67 | 2.04 | 0.01 | 0.00 | 0.00 | 0.39 | 1.64 | 5.11 | 2.15 |
| 7 | 0.40 | 0.00 | 0.00 | 0.02 | 0.02 | 0.01 | 0.05 | 0.55 | 0.04 | 0.19 | 0.01 | 1.58 | 0.52 | 0.00 | 0.00 | 0.00 | 0.15 | 0.11 | 0.49 | 0.34 |
| 8 | 1.07 | 0.00 | 0.00 | 0.00 | 0.00 | 0.00 | 0.01 | 0.21 | 0.01 | 0.06 | 0.00 | 0.93 | 0.27 | 0.00 | 0.00 | 0.00 | 0.91 | 0.39 | 3.13 | 0.23 |
| 9 | 4.60 | 0.03 | 0.00 | 0.27 | 0.27 | 0.19 | 1.24 | 1.63 | 0.10 | 1.15 | 0.20 | 0.60 | 2.17 | 0.03 | 0.00 | 0.00 | 0.27 | 1.53 | 4.02 | 1.89 |
| 10 | 2.60 | 0.05 | 0.00 | 0.38 | 0.38 | 0.26 | 1.37 | 1.53 | 0.13 | 1.27 | 0.28 | 0.72 | 2.07 | 0.02 | 0.00 | 0.00 | 0.31 | 1.27 | 2.90 | 1.65 |
| 11 | 0.97 | 0.07 | 0.00 | 0.55 | 0.57 | 0.39 | 1.62 | 1.59 | 0.20 | 1.59 | 0.41 | 0.99 | 2.12 | 0.11 | 0.01 | 0.00 | 0.22 | 0.96 | 2.17 | 1.63 |
| 12 | 1.92 | 0.08 | 0.00 | 0.52 | 0.52 | 0.36 | 1.94 | 1.50 | 0.20 | 1.49 | 0.39 | 0.84 | 1.97 | 0.29 | 0.00 | 0.00 | 0.25 | 1.27 | 1.81 | 2.13 |
| 13 | 2.61 | 0.07 | 0.00 | 0.47 | 0.48 | 0.33 | 1.87 | 2.19 | 0.18 | 1.45 | 0.35 | 0.77 | 2.09 | 0.00 | 0.00 | 0.00 | 0.32 | 1.26 | 1.75 | 2.19 |
| 14 | 1.45 | 0.04 | 0.00 | 0.40 | 0.40 | 0.27 | 1.50 | 1.53 | 0.14 | 1.47 | 0.29 | 0.76 | 2.19 | 0.00 | 0.00 | 0.00 | 0.18 | 1.14 | 2.48 | 1.65 |
| 15 | 3.43 | 0.03 | 0.00 | 0.28 | 0.28 | 0.19 | 1.23 | 1.48 | 0.09 | 1.17 | 0.21 | 0.64 | 2.11 | 0.01 | 0.00 | 0.00 | 0.19 | 1.46 | 3.26 | 1.63 |
| 16 | 1.61 | 0.14 | 0.00 | 0.89 | 0.89 | 0.62 | 2.21 | 2.13 | 0.33 | 2.05 | 0.64 | 0.90 | 2.33 | 0.01 | 0.01 | 0.00 | 0.34 | 1.17 | 1.78 | 2.20 |
| 17 | 2.23 | 0.13 | 0.00 | 0.85 | 0.84 | 0.58 | 2.21 | 2.44 | 0.35 | 1.95 | 0.61 | 0.99 | 2.34 | 0.00 | 0.01 | 0.00 | 0.40 | 1.15 | 1.84 | 2.49 |
| 18 | 1.52 | 0.12 | 0.00 | 0.78 | 0.79 | 0.55 | 2.16 | 1.99 | 0.28 | 1.90 | 0.57 | 0.84 | 2.25 | 0.04 | 0.01 | 0.00 | 0.28 | 1.22 | 1.67 | 2.04 |
| 19 | 3.39 | 0.02 | 0.00 | 0.21 | 0.20 | 0.14 | 0.91 | 1.23 | 0.07 | 0.88 | 0.15 | 0.65 | 1.69 | 0.01 | 0.00 | 0.00 | 0.43 | 1.27 | 3.93 | 1.38 |
| 20 | 2.05 | 0.07 | 0.00 | 0.49 | 0.51 | 0.35 | 1.67 | 1.99 | 0.19 | 1.56 | 0.37 | 1.00 | 2.15 | 0.00 | 0.00 | 0.00 | 0.30 | 0.87 | 1.68 | 1.94 |
| 21 | 1.64 | 0.07 | 0.00 | 0.51 | 0.53 | 0.36 | 1.80 | 1.89 | 0.18 | 1.47 | 0.38 | 0.98 | 1.95 | 0.00 | 0.00 | 0.00 | 0.22 | 1.14 | 1.48 | 1.74 |
| 22 | 2.08 | 0.10 | 0.00 | 0.77 | 0.82 | 0.56 | 2.20 | 2.07 | 0.34 | 2.08 | 0.59 | 1.02 | 2.17 | 0.00 | 0.00 | 0.00 | 0.35 | 0.93 | 0.61 | 1.74 |
| 23 | 0.81 | 0.22 | 0.00 | 1.20 | 1.31 | 0.92 | 2.23 | 2.21 | 0.89 | 2.43 | 0.94 | 1.19 | 1.78 | 0.00 | 0.04 | 0.00 | 0.89 | 0.70 | 0.62 | 1.98 |
| 24 | 0.49 | 0.20 | 0.00 | 1.16 | 1.29 | 0.89 | 2.45 | 1.70 | 0.72 | 2.26 | 0.92 | 1.25 | 1.73 | 0.00 | 0.04 | 0.00 | 1.01 | 0.85 | 0.38 | 2.00 |
| 25 | 0.55 | 0.34 | 0.00 | 1.82 | 1.92 | 1.35 | 2.41 | 1.92 | 0.93 | 2.38 | 1.37 | 1.11 | 1.82 | 0.00 | 0.03 | 0.00 | 0.61 | 0.93 | 1.20 | 2.18 |
| 26 | 1.56 | 0.11 | 0.00 | 0.81 | 0.83 | 0.57 | 2.33 | 2.23 | 0.44 | 2.18 | 0.60 | 0.96 | 2.22 | 0.00 | 0.00 | 0.00 | 0.41 | 1.40 | 1.18 | 2.22 |
| 27 | 1.55 | 0.12 | 0.00 | 0.81 | 0.83 | 0.58 | 2.07 | 2.03 | 0.37 | 1.94 | 0.60 | 0.87 | 2.13 | 0.00 | 0.01 | 0.00 | 0.39 | 1.13 | 1.63 | 1.95 |
| 28 | 0.32 | 0.02 | 0.00 | 0.05 | 0.05 | 0.04 | 0.11 | 0.07 | 0.06 | 0.15 | 0.04 | 7.05 | 0.32 | 0.00 | 0.00 | 0.00 | 0.09 | 0.00 | 0.12 | 0.16 |
| 29 | 0.69 | 0.21 | 0.00 | 1.24 | 1.34 | 0.93 | 2.53 | 1.96 | 0.67 | 2.19 | 0.96 | 1.06 | 1.84 | 0.13 | 0.04 | 0.00 | 0.57 | 0.69 | 1.33 | 2.04 |
| 30 | 0.69 | 0.21 | 0.00 | 1.20 | 1.32 | 0.92 | 2.46 | 2.45 | 0.82 | 2.22 | 0.94 | 1.05 | 1.71 | 0.00 | 0.03 | 0.00 | 0.63 | 0.59 | 1.34 | 2.00 |
| 31 | 1.53 | 0.16 | 0.00 | 0.99 | 1.11 | 0.77 | 2.48 | 2.08 | 0.73 | 2.24 | 0.79 | 1.11 | 1.70 | 0.37 | 0.03 | 0.00 | 0.90 | 0.38 | 0.71 | 1.87 |
| 32 | 0.25 | 0.31 | 0.01 | 1.51 | 1.79 | 1.25 | 1.26 | 1.22 | 2.06 | 1.94 | 1.26 | 1.09 | 0.90 | 0.04 | 0.14 | 0.00 | 1.21 | 0.38 | 0.30 | 1.15 |
| 33 | 0.29 | 0.22 | 0.02 | 1.06 | 1.24 | 0.87 | 1.66 | 1.87 | 1.39 | 1.82 | 0.88 | 1.97 | 1.18 | 0.00 | 0.03 | 0.03 | 0.73 | 0.28 | 0.07 | 1.42 |
| 34 | 1.34 | 0.12 | 0.00 | 0.75 | 0.78 | 0.54 | 1.92 | 1.59 | 0.44 | 1.72 | 0.56 | 1.41 | 1.88 | 0.00 | 0.02 | 0.00 | 0.73 | 1.14 | 1.15 | 1.92 |
| 35 | 0.97 | 0.08 | 0.00 | 0.60 | 0.62 | 0.43 | 1.85 | 1.76 | 0.29 | 1.84 | 0.45 | 1.05 | 2.12 | 0.00 | 0.01 | 0.00 | 0.27 | 1.02 | 1.39 | 1.76 |
| 36 | 1.14 | 0.26 | 0.00 | 1.45 | 1.49 | 1.05 | 2.37 | 2.55 | 0.77 | 2.30 | 1.07 | 1.11 | 1.96 | 0.00 | 0.04 | 0.00 | 0.86 | 0.64 | 0.91 | 2.29 |
| 37 | 3.04 | 0.07 | 0.00 | 0.52 | 0.52 | 0.36 | 1.81 | 2.23 | 0.22 | 1.60 | 0.38 | 0.81 | 2.24 | 0.00 | 0.00 | 0.00 | 0.32 | 1.11 | 2.70 | 2.23 |
| 38 | 1.76 | 0.12 | 0.00 | 0.72 | 0.73 | 0.51 | 2.31 | 2.25 | 0.41 | 1.89 | 0.54 | 0.94 | 2.04 | 0.01 | 0.01 | 0.00 | 0.49 | 1.17 | 1.88 | 2.34 |
| 39 | 0.84 | 0.13 | 0.00 | 0.77 | 0.83 | 0.57 | 2.17 | 2.26 | 0.47 | 1.99 | 0.60 | 0.99 | 1.82 | 0.02 | 0.00 | 0.00 | 0.51 | 0.83 | 1.12 | 2.18 |
| 40 | 0.52 | 0.34 | 0.00 | 1.65 | 1.80 | 1.27 | 2.12 | 2.18 | 1.17 | 2.20 | 1.29 | 1.06 | 1.48 | 0.00 | 0.07 | 0.00 | 1.06 | 1.52 | 0.25 | 2.05 |
| 41 | 0.42 | 0.26 | 0.00 | 1.43 | 1.50 | 1.05 | 2.73 | 2.10 | 0.68 | 2.14 | 1.08 | 1.04 | 1.92 | 0.00 | 0.04 | 0.00 | 0.32 | 1.03 | 1.08 | 2.17 |
| 42 | 1.13 | 0.18 | 0.00 | 1.06 | 1.12 | 0.78 | 2.53 | 1.94 | 0.54 | 2.28 | 0.80 | 1.00 | 2.04 | 0.02 | 0.02 | 0.00 | 0.82 | 0.93 | 1.18 | 1.91 |
| 43 | 0.83 | 0.19 | 0.00 | 1.06 | 1.14 | 0.80 | 2.40 | 1.91 | 0.65 | 2.19 | 0.82 | 1.02 | 1.80 | 0.16 | 0.02 | 0.00 | 0.95 | 1.02 | 0.83 | 1.91 |
| 44 | 0.37 | 0.25 | 0.01 | 1.23 | 1.38 | 0.97 | 1.47 | 1.39 | 1.53 | 2.03 | 0.98 | 1.47 | 1.15 | 0.03 | 0.06 | 0.01 | 1.65 | 0.36 | 0.34 | 1.45 |
| 45 | 0.58 | 0.22 | 0.00 | 1.18 | 1.30 | 0.91 | 2.17 | 1.63 | 0.86 | 2.11 | 0.93 | 1.34 | 1.58 | 0.03 | 0.05 | 0.00 | 0.98 | 0.83 | 0.31 | 1.68 |
| 46 | 0.49 | 0.17 | 0.00 | 1.02 | 1.11 | 0.77 | 2.30 | 1.67 | 0.67 | 2.22 | 0.80 | 1.17 | 1.96 | 0.02 | 0.03 | 0.00 | 1.09 | 0.58 | 0.61 | 1.84 |
| 47 | 1.19 | 0.10 | 0.00 | 0.67 | 0.69 | 0.48 | 1.76 | 1.57 | 0.26 | 1.70 | 0.50 | 1.18 | 1.95 | 0.00 | 0.01 | 0.00 | 0.26 | 0.99 | 1.37 | 1.58 |
| 48 | 0.95 | 0.18 | 0.00 | 1.06 | 1.10 | 0.77 | 2.14 | 2.03 | 0.53 | 2.04 | 0.79 | 0.98 | 1.80 | 0.01 | 0.01 | 0.00 | 0.61 | 0.69 | 0.98 | 1.92 |
| 49 | 1.52 | 0.07 | 0.00 | 0.54 | 0.55 | 0.38 | 1.68 | 1.76 | 0.21 | 1.62 | 0.40 | 0.84 | 2.04 | 0.00 | 0.00 | 0.00 | 0.24 | 0.98 | 1.65 | 1.69 |
| 50 | 2.88 | 0.02 | 0.00 | 0.19 | 0.19 | 0.13 | 0.78 | 1.06 | 0.06 | 0.78 | 0.14 | 0.67 | 1.52 | 0.01 | 0.00 | 0.00 | 0.33 | 0.95 | 3.01 | 1.20 |
| 51 | 1.31 | 0.10 | 0.00 | 0.71 | 0.72 | 0.50 | 1.92 | 1.77 | 0.25 | 1.75 | 0.52 | 0.75 | 2.09 | 0.03 | 0.01 | 0.00 | 0.28 | 1.04 | 1.35 | 1.85 |
| 52 | 1.16 | 0.11 | 0.00 | 0.72 | 0.74 | 0.51 | 1.95 | 1.66 | 0.25 | 1.77 | 0.53 | 1.10 | 2.06 | 0.06 | 0.00 | 0.00 | 0.26 | 1.05 | 1.38 | 1.77 |
| 53 | 2.29 | 0.02 | 0.00 | 0.19 | 0.19 | 0.13 | 0.82 | 1.01 | 0.06 | 0.81 | 0.14 | 0.93 | 1.55 | 0.02 | 0.00 | 0.00 | 0.30 | 1.06 | 2.71 | 1.15 |
| 54 | 0.10 | 0.00 | 0.00 | 0.00 | 0.00 | 0.00 | 0.01 | 0.08 | 0.01 | 0.04 | 0.00 | 1.10 | 0.11 | 0.04 | 0.00 | 0.00 | 0.06 | 0.02 | 0.34 | 0.11 |
| 55 | 3.10 | 0.02 | 0.00 | 0.19 | 0.19 | 0.13 | 0.80 | 1.12 | 0.06 | 0.77 | 0.14 | 0.60 | 1.50 | 0.00 | 0.00 | 0.00 | 0.36 | 0.89 | 3.04 | 1.23 |
| 56 | 1.22 | 0.11 | 0.00 | 0.77 | 0.79 | 0.54 | 1.91 | 1.80 | 0.29 | 1.76 | 0.57 | 0.76 | 2.02 | 0.00 | 0.01 | 0.00 | 0.31 | 1.02 | 1.36 | 1.83 |
| 57 | 1.03 | 0.15 | 0.00 | 0.93 | 0.98 | 0.68 | 1.96 | 1.94 | 0.50 | 1.94 | 0.70 | 0.77 | 1.83 | 0.00 | 0.02 | 0.00 | 0.54 | 0.84 | 1.09 | 1.80 |
| 58 | 0.41 | 0.16 | 0.00 | 0.95 | 1.04 | 0.72 | 1.88 | 1.31 | 0.53 | 1.92 | 0.74 | 1.26 | 1.68 | 0.07 | 0.03 | 0.00 | 0.63 | 0.75 | 0.71 | 1.40 |
| 59 | 0.39 | 0.21 | 0.01 | 1.10 | 1.22 | 0.85 | 1.66 | 1.34 | 0.92 | 1.95 | 0.87 | 1.38 | 1.31 | 0.08 | 0.04 | 0.01 | 1.08 | 0.56 | 0.42 | 1.32 |
| 60 | 0.34 | 0.31 | 0.00 | 1.59 | 1.72 | 1.21 | 2.10 | 1.80 | 0.87 | 2.08 | 1.22 | 1.08 | 1.46 | 0.00 | 0.08 | 0.00 | 0.66 | 0.90 | 0.48 | 1.61 |
| 61 | 0.65 | 0.19 | 0.00 | 1.15 | 1.27 | 0.88 | 2.23 | 1.69 | 0.64 | 2.12 | 0.90 | 1.01 | 1.60 | 0.14 | 0.03 | 0.00 | 0.58 | 0.53 | 1.02 | 1.63 |
| 62 | 0.60 | 0.20 | 0.00 | 1.17 | 1.28 | 0.89 | 2.14 | 1.83 | 0.66 | 2.20 | 0.91 | 1.12 | 1.69 | 0.00 | 0.02 | 0.00 | 0.61 | 0.78 | 0.56 | 1.70 |
| 63 | 1.22 | 0.09 | 0.00 | 0.62 | 0.65 | 0.45 | 1.69 | 1.62 | 0.21 | 1.60 | 0.47 | 1.07 | 1.91 | 0.00 | 0.00 | 0.00 | 0.22 | 0.89 | 1.10 | 1.53 |
| 64 | 1.10 | 0.20 | 0.00 | 1.15 | 1.27 | 0.89 | 2.40 | 1.40 | 0.71 | 2.18 | 0.91 | 1.32 | 1.75 | 0.29 | 0.04 | 0.00 | 0.77 | 0.62 | 1.97 | 1.79 |
| 65 | 0.26 | 0.18 | 0.00 | 0.96 | 1.05 | 0.73 | 1.96 | 1.17 | 0.71 | 1.87 | 0.75 | 1.90 | 1.40 | 0.05 | 0.04 | 0.00 | 1.11 | 0.42 | 0.63 | 1.40 |
| 66 | 1.26 | 0.10 | 0.00 | 0.69 | 0.73 | 0.51 | 1.89 | 1.66 | 0.29 | 1.64 | 0.53 | 1.60 | 1.76 | 0.03 | 0.01 | 0.00 | 0.38 | 0.77 | 0.77 | 1.55 |
| 67 | 0.49 | 1.11 | 0.00 | 2.96 | 3.09 | 2.35 | 1.06 | 0.98 | 1.09 | 1.07 | 2.33 | 2.33 | 0.81 | 0.03 | 0.32 | 0.00 | 0.39 | 0.84 | 0.20 | 0.75 |
| 68 | 0.57 | 1.06 | 0.00 | 2.68 | 2.97 | 2.26 | 0.97 | 0.71 | 1.63 | 1.12 | 2.24 | 1.46 | 0.61 | 0.12 | 0.28 | 0.00 | 0.98 | 1.56 | 0.15 | 0.71 |
| 69 | 0.48 | 0.32 | 0.01 | 1.37 | 1.70 | 1.20 | 1.36 | 1.54 | 1.82 | 1.80 | 1.20 | 1.40 | 0.86 | 0.13 | 0.14 | 0.00 | 1.24 | 0.38 | 0.16 | 1.08 |
| 70 | 0.23 | 0.31 | 0.02 | 1.38 | 1.64 | 1.16 | 1.13 | 1.11 | 1.61 | 1.70 | 1.16 | 1.33 | 0.80 | 0.20 | 0.17 | 0.00 | 1.12 | 0.44 | 0.21 | 0.90 |
| 71 | 0.29 | 0.29 | 0.01 | 1.25 | 1.39 | 0.99 | 1.34 | 1.08 | 1.16 | 1.69 | 1.00 | 1.46 | 0.99 | 0.15 | 0.16 | 0.00 | 1.54 | 0.55 | 0.51 | 1.00 |
| 72 | 0.15 | 0.30 | 0.01 | 1.27 | 1.49 | 1.06 | 1.25 | 1.29 | 1.67 | 1.75 | 1.06 | 1.44 | 0.93 | 0.07 | 0.13 | 0.00 | 2.02 | 0.33 | 0.44 | 1.05 |
| 73 | 0.38 | 0.41 | 0.07 | 1.31 | 1.32 | 0.99 | 0.75 | 0.93 | 1.26 | 1.04 | 0.98 | 1.54 | 0.53 | 1.04 | 0.41 | 0.00 | 1.90 | 1.51 | 0.18 | 0.62 |
| 74 | 0.17 | 1.46 | 0.00 | 2.06 | 2.06 | 1.80 | 0.42 | 0.55 | 1.13 | 0.57 | 1.78 | 1.02 | 0.36 | 22.88 | 15.71 | 0.00 | 0.73 | 0.83 | 0.10 | 0.34 |
| 75 | 0.06 | 1.03 | 0.00 | 1.95 | 1.75 | 1.46 | 0.29 | 0.42 | 1.09 | 0.46 | 1.44 | 1.01 | 0.24 | 0.83 | 0.47 | 0.00 | 4.23 | 4.47 | 0.04 | 0.23 |
| 76 | 0.02 | 4.67 | 0.01 | 3.51 | 2.90 | 3.37 | 0.10 | 0.08 | 1.85 | 0.15 | 3.31 | 0.39 | 0.09 | 0.92 | 4.87 | 0.00 | 4.32 | 3.02 | 0.03 | 0.10 |
| 77 | 0.03 | 1.35 | 0.00 | 1.27 | 1.13 | 1.16 | 0.14 | 0.17 | 1.09 | 0.21 | 1.15 | 0.80 | 0.12 | 6.32 | 3.27 | 0.02 | 2.26 | 1.20 | 0.02 | 0.12 |
| 78 | 0.00 | 2.73 | 0.01 | 2.29 | 2.11 | 2.24 | 0.18 | 0.17 | 1.93 | 0.26 | 2.20 | 0.81 | 0.12 | 3.90 | 5.14 | 0.00 | 2.51 | 2.71 | 0.08 | 0.16 |
| 79 | 0.00 | 4.14 | 4.97 | 2.01 | 1.86 | 2.69 | 0.06 | 0.12 | 2.26 | 0.15 | 2.65 | 0.52 | 0.07 | 4.08 | 4.70 | 0.05 | 1.91 | 1.15 | 0.05 | 0.07 |
| 80 | 0.06 | 0.51 | 0.38 | 0.97 | 1.10 | 0.89 | 0.32 | 0.39 | 1.28 | 0.56 | 0.88 | 1.61 | 0.26 | 0.26 | 0.33 | 0.02 | 1.39 | 1.42 | 0.02 | 0.28 |
| 81 | 0.08 | 1.58 | 1.32 | 1.12 | 1.20 | 1.33 | 0.18 | 0.29 | 1.88 | 0.43 | 1.31 | 1.38 | 0.17 | 2.63 | 1.48 | 0.02 | 1.45 | 1.38 | 0.00 | 0.23 |
| 82 | 0.05 | 0.50 | 0.44 | 0.19 | 0.20 | 0.31 | 0.03 | 0.04 | 0.26 | 0.11 | 0.30 | 4.31 | 0.10 | 0.03 | 0.34 | 0.13 | 0.71 | 0.05 | 0.00 | 0.09 |
| 83 | 0.00 | 2.81 | 0.63 | 2.18 | 1.99 | 2.20 | 0.09 | 0.10 | 2.81 | 0.14 | 2.17 | 0.46 | 0.08 | 5.19 | 4.89 | 0.01 | 2.32 | 0.86 | 0.22 | 0.11 |
| 84 | 0.00 | 0.50 | 0.12 | 0.92 | 1.04 | 0.84 | 0.24 | 0.36 | 1.97 | 0.62 | 0.83 | 1.56 | 0.32 | 1.57 | 0.20 | 0.01 | 1.51 | 1.20 | 0.01 | 0.45 |
| 85 | 0.00 | 2.41 | 3.00 | 1.02 | 1.00 | 1.52 | 0.02 | 0.01 | 2.05 | 0.14 | 1.50 | 0.97 | 0.07 | 1.86 | 2.17 | 0.07 | 1.19 | 0.32 | 0.05 | 0.08 |
| 86 | 0.00 | 2.39 | 2.98 | 1.01 | 1.04 | 1.54 | 0.03 | 0.10 | 3.62 | 0.07 | 1.52 | 0.71 | 0.03 | 0.01 | 0.87 | 0.10 | 1.23 | 0.47 | 0.00 | 0.02 |
| 87 | 0.02 | 2.41 | 2.71 | 1.38 | 1.33 | 1.72 | 0.12 | 0.14 | 1.59 | 0.23 | 1.69 | 1.33 | 0.11 | 2.22 | 2.63 | 0.06 | 1.48 | 1.38 | 0.07 | 0.12 |
| 88 | 0.07 | 2.45 | 0.01 | 2.38 | 2.06 | 2.12 | 0.20 | 0.24 | 1.27 | 0.30 | 2.09 | 0.81 | 0.17 | 4.07 | 4.41 | 0.00 | 2.96 | 3.15 | 0.04 | 0.17 |
| 89 | 0.38 | 0.33 | 0.01 | 1.35 | 1.52 | 1.09 | 1.39 | 1.27 | 1.07 | 1.63 | 1.09 | 1.42 | 1.03 | 0.07 | 0.10 | 0.00 | 0.85 | 0.57 | 0.45 | 1.08 |
| 90 | 0.01 | 2.63 | 0.01 | 2.21 | 2.02 | 2.14 | 0.14 | 0.09 | 2.30 | 0.22 | 2.11 | 0.96 | 0.12 | 4.98 | 4.01 | 0.00 | 2.35 | 1.48 | 0.03 | 0.21 |
| 91 | 0.24 | 0.34 | 0.32 | 0.79 | 0.91 | 0.70 | 0.52 | 0.38 | 1.73 | 0.73 | 0.70 | 2.80 | 0.41 | 0.01 | 0.23 | 0.20 | 0.41 | 0.32 | 0.02 | 0.41 |
| 92 | 0.01 | 3.76 | 12.53 | 0.87 | 0.95 | 2.24 | 0.05 | 0.08 | 2.94 | 0.12 | 2.21 | 0.85 | 0.07 | 0.07 | 1.73 | 0.07 | 0.88 | 0.90 | 0.08 | 0.05 |
| 93 | 0.00 | 4.70 | 13.07 | 0.88 | 0.96 | 2.57 | 0.08 | 0.09 | 3.01 | 0.13 | 2.53 | 0.82 | 0.07 | 0.05 | 1.82 | 0.15 | 1.55 | 0.14 | 0.03 | 0.06 |
| 94 | 0.01 | 4.01 | 9.68 | 0.55 | 0.63 | 2.02 | 0.10 | 0.05 | 3.57 | 0.12 | 1.99 | 0.84 | 0.08 | 0.03 | 1.05 | 16.38 | 0.89 | 1.05 | 0.02 | 0.12 |
| 95 | 0.14 | 2.48 | 3.57 | 0.56 | 0.63 | 1.32 | 0.12 | 0.20 | 1.99 | 0.19 | 1.30 | 3.77 | 0.13 | 0.01 | 0.45 | 4.96 | 0.59 | 0.42 | 0.07 | 0.23 |
| 96 | 0.00 | 0.55 | 0.95 | 0.04 | 0.04 | 0.24 | 0.00 | 0.00 | 0.36 | 0.01 | 0.23 | 0.16 | 0.00 | 0.01 | 0.04 | 7.20 | 0.03 | 0.04 | 0.00 | 0.00 |
| 97 | 0.00 | 3.95 | 9.00 | 0.39 | 0.44 | 1.86 | 0.02 | 0.04 | 3.40 | 0.07 | 1.83 | 0.79 | 0.04 | 4.57 | 0.48 | 16.75 | 0.45 | 0.09 | 0.12 | 0.05 |
| 98 | 0.00 | 4.40 | 8.43 | 0.52 | 0.58 | 2.07 | 0.02 | 0.07 | 2.90 | 0.07 | 2.04 | 1.17 | 0.05 | 0.00 | 1.71 | 0.37 | 1.00 | 0.00 | 0.00 | 0.07 |
| 99 | 0.00 | 2.98 | 6.09 | 0.60 | 0.62 | 1.56 | 0.03 | 0.00 | 2.92 | 0.09 | 1.54 | 1.06 | 0.06 | 4.58 | 0.91 | 0.07 | 1.11 | 0.30 | 0.00 | 0.01 |
| 100 | 0.00 | 0.02 | 0.00 | 0.00 | 0.00 | 0.01 | 0.00 | 0.00 | 0.00 | 0.01 | 0.01 | 0.74 | 0.00 | 0.00 | 0.00 | 0.00 | 0.02 | 0.00 | 1.08 | 0.18 |
| Base | 0.00 | 2.69 | 0.29 | 3.58 | 5.40 | 8.39 | 0.12 | 0.01 | 16.54 | 3.36 | 8.52 | 0.54 | 2.72 | 0.00 | 0.02 | 0.13 | 0.02 | 0.00 | 0.00 | 0.12 |

| | SINE family | Satellite family | Simple family | TcMar-Mariner family | TcMar-Tc2 family | TcMar-Tigger family | TcMar family | Unknown family | acro family | centr family | hAT-Blackjack family | hAT-Charlie family | hAT-Tip100 family | hAT family | rRNA family | scRNA family | snRNA family | srpRNA family | tRNA family | telo family |
|---|---|---|---|---|---|---|---|---|---|---|---|---|---|---|---|---|---|---|---|---|
| 1 | 0.00 | 0.00 | 0.12 | 0.00 | 0.00 | 0.00 | 0.00 | 0.00 | 0.00 | 0.00 | 0.00 | 0.00 | 0.00 | 0.00 | 0.09 | 0.00 | 0.04 | 0.00 | 0.16 | 0.00 |
| 2 | 0.01 | 0.00 | 0.16 | 0.00 | 0.01 | 0.00 | 0.01 | 1.38 | 0.00 | 0.00 | 0.00 | 0.00 | 0.00 | 0.00 | 0.04 | 0.00 | 0.05 | 0.00 | 0.05 | 0.00 |
| 3 | 0.01 | 0.00 | 0.15 | 0.00 | 0.02 | 0.00 | 0.01 | 0.73 | 0.00 | 0.00 | 0.00 | 0.00 | 0.00 | 0.00 | 0.07 | 0.00 | 0.02 | 0.00 | 0.03 | 0.00 |
| 4 | 0.02 | 0.01 | 0.19 | 0.00 | 0.02 | 0.00 | 0.02 | 1.76 | 0.00 | 0.00 | 0.01 | 0.01 | 0.01 | 0.01 | 0.00 | 0.00 | 0.05 | 0.08 | 0.11 | 0.00 |
| 5 | 0.07 | 0.00 | 0.15 | 0.11 | 0.03 | 0.07 | 0.19 | 4.82 | 0.00 | 0.00 | 0.03 | 0.03 | 0.02 | 0.03 | 0.09 | 0.02 | 0.05 | 0.00 | 2.51 | 0.00 |
| 6 | 0.45 | 0.00 | 0.17 | 0.85 | 1.05 | 0.34 | 0.47 | 2.81 | 0.00 | 0.00 | 0.89 | 1.09 | 0.69 | 1.04 | 0.25 | 0.00 | 0.17 | 0.05 | 4.33 | 0.00 |
| 7 | 0.12 | 0.01 | 0.31 | 0.19 | 0.37 | 0.07 | 0.10 | 0.68 | 0.00 | 0.00 | 0.16 | 0.22 | 0.17 | 0.22 | 0.05 | 0.00 | 0.08 | 0.08 | 0.46 | 0.00 |
| 8 | 0.07 | 0.01 | 0.17 | 0.16 | 0.05 | 0.06 | 0.11 | 1.96 | 0.00 | 0.00 | 0.05 | 0.05 | 0.03 | 0.05 | 0.02 | 0.01 | 0.02 | 0.01 | 1.71 | 0.00 |
| 9 | 0.47 | 0.00 | 0.16 | 0.75 | 0.98 | 0.28 | 0.39 | 2.12 | 0.00 | 0.00 | 0.96 | 1.15 | 0.75 | 1.10 | 0.24 | 0.00 | 0.09 | 0.02 | 3.13 | 0.00 |
| 10 | 0.45 | 0.01 | 0.26 | 0.58 | 1.08 | 0.26 | 0.35 | 1.73 | 0.00 | 0.00 | 1.10 | 1.20 | 0.90 | 1.17 | 0.23 | 0.00 | 0.09 | 0.02 | 2.12 | 0.00 |
| 11 | 0.46 | 0.01 | 0.46 | 0.49 | 1.16 | 0.25 | 0.34 | 1.70 | 0.00 | 0.00 | 1.26 | 1.37 | 1.13 | 1.35 | 0.14 | 0.00 | 0.08 | 0.02 | 1.49 | 0.00 |
| 12 | 0.43 | 0.01 | 0.30 | 0.60 | 1.02 | 0.26 | 0.35 | 2.01 | 0.00 | 0.00 | 1.25 | 1.33 | 1.15 | 1.32 | 0.22 | 0.00 | 0.09 | 0.02 | 1.86 | 0.01 |
| 13 | 0.45 | 0.01 | 0.21 | 0.66 | 1.47 | 0.29 | 0.41 | 1.99 | 0.00 | 0.00 | 1.22 | 1.41 | 1.03 | 1.36 | 0.32 | 0.00 | 0.07 | 0.00 | 1.86 | 0.00 |
| 14 | 0.47 | 0.01 | 0.26 | 0.54 | 1.09 | 0.25 | 0.35 | 1.51 | 0.00 | 0.00 | 1.12 | 1.26 | 1.01 | 1.24 | 0.22 | 0.00 | 0.13 | 0.02 | 1.88 | 0.00 |
| 15 | 0.46 | 0.01 | 0.20 | 0.64 | 0.89 | 0.25 | 0.34 | 1.80 | 0.00 | 0.00 | 0.97 | 1.14 | 0.78 | 1.10 | 0.23 | 0.00 | 0.07 | 0.02 | 2.74 | 0.00 |
| 16 | 0.50 | 0.00 | 0.40 | 0.62 | 1.81 | 0.37 | 0.49 | 1.76 | 0.00 | 0.00 | 1.73 | 1.80 | 1.62 | 1.79 | 0.38 | 0.00 | 0.17 | 0.02 | 1.20 | 0.00 |
| 17 | 0.51 | 0.00 | 0.39 | 0.72 | 2.03 | 0.40 | 0.53 | 2.00 | 0.00 | 0.00 | 1.72 | 1.83 | 1.60 | 1.81 | 0.40 | 0.00 | 0.17 | 0.03 | 1.58 | 0.00 |
| 18 | 0.49 | 0.01 | 0.42 | 0.58 | 1.56 | 0.32 | 0.43 | 1.70 | 0.00 | 0.00 | 1.64 | 1.68 | 1.51 | 1.67 | 0.33 | 0.00 | 0.13 | 0.02 | 1.25 | 0.00 |
| 19 | 0.37 | 0.01 | 0.18 | 0.56 | 0.70 | 0.21 | 0.30 | 2.00 | 0.00 | 0.00 | 0.75 | 0.87 | 0.58 | 0.84 | 0.16 | 0.01 | 0.06 | 0.02 | 2.81 | 0.00 |
| 20 | 0.47 | 0.01 | 0.31 | 0.59 | 1.32 | 0.27 | 0.38 | 1.79 | 0.00 | 0.00 | 1.32 | 1.44 | 1.09 | 1.40 | 0.24 | 0.00 | 0.05 | 0.00 | 1.81 | 0.00 |
| 21 | 0.42 | 0.01 | 0.42 | 0.56 | 1.18 | 0.23 | 0.33 | 2.15 | 0.00 | 0.00 | 1.28 | 1.29 | 1.20 | 1.30 | 0.25 | 0.01 | 0.04 | 0.01 | 1.86 | 0.00 |
| 22 | 0.47 | 0.02 | 0.53 | 0.49 | 1.73 | 0.28 | 0.39 | 1.81 | 0.00 | 0.00 | 1.78 | 1.68 | 1.69 | 1.70 | 0.17 | 0.00 | 0.17 | 0.01 | 1.13 | 0.00 |
| 23 | 0.42 | 0.00 | 0.61 | 0.43 | 2.63 | 0.39 | 0.51 | 1.50 | 0.00 | 0.00 | 2.21 | 2.07 | 2.35 | 2.12 | 0.50 | 0.08 | 0.25 | 0.05 | 0.63 | 0.00 |
| 24 | 0.40 | 0.00 | 0.71 | 0.48 | 2.22 | 0.37 | 0.48 | 1.49 | 0.00 | 0.00 | 2.22 | 1.85 | 2.42 | 1.94 | 0.44 | 0.08 | 0.11 | 0.04 | 0.51 | 0.00 |
| 25 | 0.40 | 0.00 | 0.71 | 0.42 | 2.58 | 0.58 | 0.68 | 1.40 | 0.00 | 0.00 | 2.28 | 1.99 | 2.48 | 2.07 | 0.32 | 0.02 | 0.09 | 0.01 | 0.69 | 0.00 |
| 26 | 0.48 | 0.00 | 0.29 | 0.67 | 2.19 | 0.37 | 0.50 | 1.88 | 0.00 | 0.00 | 1.76 | 1.99 | 1.89 | 1.97 | 0.49 | 0.00 | 0.09 | 0.02 | 0.99 | 0.00 |
| 27 | 0.46 | 0.00 | 0.35 | 0.57 | 1.82 | 0.33 | 0.45 | 1.73 | 0.00 | 0.00 | 1.60 | 1.77 | 1.67 | 1.76 | 0.42 | 0.01 | 0.11 | 0.02 | 1.20 | 0.00 |
| 28 | 0.10 | 0.23 | 2.45 | 0.10 | 0.17 | 0.05 | 0.06 | 0.13 | 0.00 | 0.00 | 0.13 | 0.19 | 0.15 | 0.18 | 0.15 | 0.08 | 0.21 | 0.05 | 0.40 | 0.03 |
| 29 | 0.41 | 0.01 | 0.65 | 0.42 | 2.22 | 0.36 | 0.47 | 1.41 | 0.00 | 0.00 | 2.17 | 1.78 | 2.07 | 1.85 | 0.17 | 0.02 | 0.13 | 0.05 | 0.95 | 0.00 |
| 30 | 0.38 | 0.00 | 0.38 | 0.47 | 2.33 | 0.36 | 0.47 | 1.69 | 0.00 | 0.00 | 2.31 | 1.88 | 2.45 | 1.98 | 0.30 | 0.02 | 0.09 | 0.04 | 0.69 | 0.00 |
| 31 | 0.39 | 0.01 | 0.49 | 0.54 | 2.30 | 0.30 | 0.44 | 1.16 | 0.00 | 0.00 | 1.96 | 1.86 | 2.25 | 1.91 | 0.41 | 0.06 | 0.11 | 0.05 | 0.87 | 0.01 |
| 32 | 0.38 | 0.01 | 0.88 | 0.33 | 2.60 | 0.48 | 0.56 | 0.60 | 0.21 | 0.00 | 2.95 | 1.77 | 2.75 | 1.95 | 0.28 | 0.38 | 0.46 | 0.19 | 0.95 | 0.01 |
| 33 | 0.37 | 0.04 | 2.28 | 0.33 | 2.05 | 0.31 | 0.40 | 1.01 | 0.00 | 0.00 | 2.53 | 1.72 | 2.26 | 1.82 | 0.80 | 0.10 | 0.27 | 0.16 | 0.28 | 0.00 |
| 34 | 0.42 | 0.01 | 0.99 | 0.66 | 1.46 | 0.28 | 0.39 | 1.50 | 0.00 | 0.00 | 1.60 | 1.51 | 1.57 | 1.52 | 0.38 | 0.01 | 0.32 | 0.03 | 1.13 | 0.00 |
| 35 | 0.46 | 0.01 | 0.39 | 0.59 | 1.70 | 0.27 | 0.39 | 1.54 | 0.00 | 0.00 | 1.42 | 1.65 | 1.37 | 1.61 | 0.27 | 0.01 | 0.14 | 0.04 | 1.55 | 0.00 |
| 36 | 0.43 | 0.00 | 0.46 | 0.49 | 2.66 | 0.49 | 0.61 | 1.77 | 0.00 | 0.00 | 2.15 | 2.08 | 2.42 | 2.13 | 0.70 | 0.15 | 0.24 | 0.02 | 0.79 | 0.00 |
| 37 | 0.49 | 0.00 | 0.20 | 0.81 | 1.69 | 0.33 | 0.46 | 2.21 | 0.00 | 0.00 | 1.27 | 1.61 | 1.23 | 1.55 | 0.31 | 0.03 | 0.12 | 0.02 | 1.95 | 0.00 |
| 38 | 0.45 | 0.00 | 0.24 | 0.65 | 2.25 | 0.32 | 0.46 | 1.97 | 0.00 | 0.00 | 1.60 | 1.86 | 1.63 | 1.82 | 0.30 | 0.02 | 0.08 | 0.13 | 1.26 | 0.00 |
| 39 | 0.40 | 0.00 | 0.31 | 0.61 | 2.04 | 0.29 | 0.42 | 1.88 | 0.00 | 0.00 | 1.74 | 1.78 | 1.71 | 1.78 | 0.47 | 0.01 | 0.15 | 0.04 | 0.96 | 0.01 |
| 40 | 0.33 | 0.00 | 0.48 | 0.43 | 2.42 | 0.51 | 0.60 | 1.34 | 0.00 | 0.00 | 2.13 | 1.87 | 2.81 | 2.00 | 0.56 | 0.06 | 0.25 | 0.03 | 0.51 | 0.00 |
| 41 | 0.42 | 0.00 | 0.59 | 0.47 | 2.09 | 0.48 | 0.57 | 1.63 | 0.00 | 0.00 | 2.17 | 1.82 | 1.96 | 1.87 | 0.21 | 0.03 | 0.21 | 0.04 | 0.75 | 0.00 |
| 42 | 0.45 | 0.02 | 0.61 | 0.49 | 1.81 | 0.34 | 0.44 | 1.66 | 0.00 | 0.00 | 1.96 | 1.82 | 2.12 | 1.87 | 0.29 | 0.02 | 0.21 | 0.04 | 0.89 | 0.00 |
| 43 | 0.40 | 0.00 | 0.42 | 0.42 | 2.25 | 0.32 | 0.44 | 1.43 | 0.00 | 0.00 | 1.80 | 1.81 | 1.97 | 1.84 | 0.49 | 0.03 | 0.17 | 0.03 | 1.87 | 0.00 |
| 44 | 0.37 | 0.01 | 1.31 | 0.36 | 2.99 | 0.39 | 0.51 | 1.04 | 0.00 | 0.00 | 2.65 | 1.93 | 2.50 | 2.03 | 0.67 | 0.32 | 0.53 | 0.20 | 1.05 | 0.01 |
| 45 | 0.39 | 0.01 | 0.89 | 0.44 | 2.01 | 0.34 | 0.44 | 1.26 | 0.00 | 0.00 | 1.94 | 1.77 | 2.42 | 1.87 | 0.34 | 0.10 | 0.33 | 0.24 | 0.79 | 0.00 |
| 46 | 0.45 | 0.00 | 0.62 | 0.50 | 2.19 | 0.36 | 0.47 | 1.46 | 0.00 | 0.00 | 1.83 | 1.99 | 2.12 | 2.00 | 0.42 | 0.08 | 0.20 | 0.02 | 0.72 | 0.00 |
| 47 | 0.42 | 0.01 | 0.57 | 0.46 | 1.36 | 0.27 | 0.36 | 1.32 | 0.00 | 0.00 | 1.48 | 1.51 | 1.46 | 1.51 | 0.30 | 0.01 | 0.13 | 0.01 | 1.05 | 0.00 |
| 48 | 0.39 | 0.00 | 0.40 | 0.46 | 1.99 | 0.34 | 0.45 | 1.52 | 0.00 | 0.00 | 1.79 | 1.78 | 1.97 | 1.81 | 0.54 | 0.04 | 0.20 | 0.03 | 0.63 | 0.00 |
| 49 | 0.44 | 0.00 | 0.26 | 0.56 | 1.39 | 0.26 | 0.36 | 1.52 | 0.00 | 0.00 | 1.29 | 1.49 | 1.25 | 1.46 | 0.25 | 0.02 | 0.07 | 0.03 | 1.23 | 0.00 |
| 50 | 0.33 | 0.01 | 0.16 | 0.48 | 0.60 | 0.18 | 0.26 | 1.66 | 0.00 | 0.00 | 0.65 | 0.77 | 0.52 | 0.74 | 0.14 | 0.01 | 0.06 | 0.02 | 2.10 | 0.00 |
| 51 | 0.45 | 0.00 | 0.30 | 0.51 | 1.37 | 0.28 | 0.38 | 1.52 | 0.00 | 0.00 | 1.47 | 1.51 | 1.39 | 1.51 | 0.30 | 0.02 | 0.11 | 0.01 | 1.06 | 0.00 |
| 52 | 0.45 | 0.01 | 0.53 | 0.49 | 1.35 | 0.29 | 0.38 | 1.45 | 0.00 | 0.00 | 1.57 | 1.55 | 1.47 | 1.55 | 0.28 | 0.00 | 0.11 | 0.01 | 1.06 | 0.00 |
| 53 | 0.34 | 0.01 | 0.27 | 0.47 | 0.64 | 0.18 | 0.25 | 1.53 | 0.00 | 0.00 | 0.75 | 0.82 | 0.57 | 0.80 | 0.12 | 0.00 | 0.07 | 0.01 | 2.17 | 0.00 |
| 54 | 0.03 | 0.01 | 0.26 | 0.04 | 0.08 | 0.01 | 0.02 | 0.47 | 0.00 | 0.00 | 0.03 | 0.04 | 0.04 | 0.04 | 0.04 | 0.00 | 0.03 | 0.01 | 0.22 | 0.00 |
| 55 | 0.33 | 0.01 | 0.18 | 0.46 | 0.66 | 0.19 | 0.27 | 1.83 | 0.00 | 0.00 | 0.66 | 0.77 | 0.54 | 0.74 | 0.13 | 0.00 | 0.06 | 0.02 | 2.25 | 0.00 |
| 56 | 0.44 | 0.01 | 0.47 | 0.48 | 1.48 | 0.29 | 0.38 | 1.54 | 0.00 | 0.00 | 1.54 | 1.52 | 1.46 | 1.52 | 0.23 | 0.00 | 0.12 | 0.02 | 1.03 | 0.00 |
| 57 | 0.40 | 0.00 | 0.39 | 0.45 | 1.89 | 0.31 | 0.42 | 1.44 | 0.00 | 0.00 | 1.68 | 1.70 | 1.85 | 1.72 | 0.40 | 0.01 | 0.17 | 0.03 | 0.82 | 0.00 |
| 58 | 0.39 | 0.01 | 0.80 | 0.42 | 1.51 | 0.29 | 0.38 | 1.17 | 0.00 | 0.00 | 1.61 | 1.58 | 1.89 | 1.63 | 0.31 | 0.08 | 0.24 | 0.08 | 0.69 | 0.00 |
| 59 | 0.36 | 0.01 | 1.12 | 0.40 | 2.05 | 0.34 | 0.44 | 0.93 | 0.00 | 0.00 | 1.99 | 1.63 | 2.08 | 1.71 | 0.67 | 0.43 | 0.43 | 0.15 | 0.61 | 0.01 |
| 60 | 0.33 | 0.00 | 0.70 | 0.35 | 1.90 | 0.47 | 0.54 | 1.11 | 0.00 | 0.00 | 1.97 | 1.63 | 2.37 | 1.75 | 0.32 | 0.12 | 0.28 | 0.06 | 0.38 | 0.00 |
| 61 | 0.36 | 0.00 | 0.54 | 0.38 | 1.87 | 0.31 | 0.40 | 1.22 | 0.00 | 0.00 | 1.99 | 1.65 | 2.18 | 1.74 | 0.30 | 0.02 | 0.12 | 0.06 | 0.57 | 0.00 |
| 62 | 0.38 | 0.00 | 0.60 | 0.42 | 1.99 | 0.37 | 0.47 | 1.28 | 0.00 | 0.00 | 1.99 | 1.80 | 2.17 | 1.86 | 0.44 | 0.13 | 0.16 | 0.05 | 0.50 | 0.00 |
| 63 | 0.41 | 0.01 | 0.45 | 0.42 | 1.15 | 0.24 | 0.32 | 1.43 | 0.00 | 0.00 | 1.37 | 1.35 | 1.28 | 1.36 | 0.16 | 0.01 | 0.08 | 0.00 | 1.15 | 0.00 |
| 64 | 0.42 | 0.01 | 0.88 | 0.36 | 1.89 | 0.31 | 0.41 | 1.34 | 0.00 | 0.00 | 2.25 | 1.80 | 2.13 | 1.87 | 0.54 | 0.09 | 0.17 | 0.07 | 0.63 | 0.00 |
| 65 | 0.55 | 0.02 | 1.44 | 0.67 | 1.75 | 0.45 | 0.54 | 1.14 | 0.00 | 0.00 | 1.74 | 1.54 | 1.98 | 1.61 | 1.01 | 0.88 | 1.07 | 0.31 | 0.67 | 0.03 |
| 66 | 0.39 | 0.03 | 1.16 | 0.47 | 1.07 | 0.23 | 0.31 | 1.32 | 0.00 | 0.00 | 1.54 | 1.37 | 1.31 | 1.38 | 0.22 | 0.10 | 0.17 | 0.04 | 0.75 | 0.00 |
| 67 | 0.55 | 0.02 | 2.64 | 0.46 | 0.86 | 0.32 | 0.36 | 0.75 | 0.00 | 0.00 | 1.06 | 0.89 | 1.13 | 0.94 | 1.50 | 0.94 | 1.38 | 0.37 | 0.70 | 0.00 |
| 68 | 1.04 | 0.00 | 1.39 | 0.49 | 1.14 | 0.56 | 0.58 | 0.42 | 0.00 | 0.00 | 1.25 | 0.96 | 1.44 | 1.04 | 0.90 | 1.21 | 1.45 | 0.82 | 0.38 | 0.00 |
| 69 | 0.62 | 0.02 | 1.33 | 0.41 | 2.03 | 0.58 | 0.63 | 0.77 | 0.62 | 0.00 | 2.76 | 1.55 | 2.71 | 1.75 | 0.88 | 0.99 | 0.84 | 0.58 | 0.49 | 0.00 |
| 70 | 0.60 | 0.01 | 1.34 | 0.47 | 1.83 | 0.60 | 0.65 | 0.53 | 0.37 | 0.00 | 2.35 | 1.42 | 2.38 | 1.59 | 0.90 | 1.31 | 0.98 | 0.54 | 0.38 | 0.01 |
| 71 | 0.85 | 0.01 | 1.27 | 0.49 | 1.46 | 0.78 | 0.79 | 0.70 | 0.00 | 0.00 | 2.02 | 1.46 | 2.07 | 1.56 | 1.25 | 1.39 | 1.21 | 0.98 | 0.42 | 0.00 |
| 72 | 0.78 | 0.01 | 1.23 | 0.41 | 1.91 | 0.72 | 0.75 | 0.78 | 0.00 | 0.00 | 2.39 | 1.68 | 2.41 | 1.80 | 0.91 | 0.92 | 0.93 | 0.81 | 0.54 | 0.00 |
| 73 | 2.52 | 0.01 | 1.94 | 1.01 | 1.52 | 1.29 | 1.26 | 0.31 | 0.00 | 0.00 | 1.61 | 1.24 | 1.43 | 1.27 | 2.54 | 3.49 | 3.75 | 3.16 | 0.97 | 0.04 |
| 74 | 1.80 | 0.00 | 0.95 | 3.43 | 0.58 | 14.38 | 12.68 | 0.23 | 0.00 | 0.00 | 0.75 | 0.60 | 0.86 | 0.64 | 2.47 | 3.60 | 3.66 | 5.27 | 0.69 | 0.06 |
| 75 | 3.71 | 0.01 | 1.01 | 1.86 | 0.54 | 1.15 | 1.15 | 0.18 | 0.04 | 0.01 | 0.63 | 1.06 | 0.68 | 0.97 | 4.07 | 8.12 | 6.29 | 6.21 | 0.86 | 0.14 |
| 76 | 2.14 | 0.02 | 0.45 | 5.57 | 0.19 | 3.77 | 3.66 | 0.07 | 0.31 | 0.01 | 0.24 | 0.95 | 0.22 | 0.79 | 2.78 | 3.90 | 3.74 | 5.13 | 0.57 | 0.39 |
| 77 | 4.52 | 0.05 | 1.29 | 2.08 | 0.23 | 2.28 | 2.13 | 0.07 | 0.00 | 0.00 | 0.29 | 0.48 | 0.39 | 0.45 | 2.15 | 3.21 | 2.49 | 4.19 | 0.38 | 0.03 |
| 78 | 2.47 | 0.03 | 0.98 | 3.59 | 0.40 | 3.87 | 3.62 | 0.09 | 0.02 | 0.01 | 0.45 | 0.70 | 0.51 | 0.65 | 2.66 | 5.20 | 4.21 | 4.92 | 0.67 | 0.09 |
| 79 | 2.20 | 0.49 | 0.74 | 2.93 | 0.21 | 3.26 | 3.04 | 0.08 | 0.31 | 0.08 | 0.31 | 0.44 | 0.31 | 0.40 | 1.77 | 2.19 | 2.22 | 2.56 | 0.48 | 1.11 |
| 80 | 3.86 | 0.02 | 2.44 | 0.65 | 0.71 | 0.62 | 0.62 | 0.19 | 1.31 | 0.01 | 0.88 | 0.67 | 0.88 | 0.70 | 1.95 | 2.52 | 2.09 | 1.34 | 0.40 | 0.02 |
| 81 | 2.79 | 0.34 | 2.28 | 1.07 | 0.63 | 1.33 | 1.25 | 0.14 | 0.00 | 0.05 | 0.75 | 0.63 | 0.76 | 0.64 | 3.05 | 2.68 | 2.50 | 1.74 | 0.77 | 2.50 |
| 82 | 0.66 | 0.57 | 4.38 | 0.65 | 0.14 | 0.72 | 0.68 | 0.04 | 7.75 | 0.04 | 0.09 | 0.16 | 0.13 | 0.15 | 3.16 | 2.85 | 2.51 | 2.58 | 1.14 | 2.38 |
| 83 | 1.74 | 0.28 | 0.69 | 2.77 | 0.16 | 3.27 | 3.02 | 0.03 | 0.59 | 0.11 | 0.23 | 0.48 | 0.27 | 0.43 | 2.12 | 1.76 | 2.64 | 2.75 | 0.30 | 1.01 |
| 84 | 2.30 | 0.08 | 2.22 | 0.61 | 0.67 | 0.69 | 0.67 | 0.13 | 10.79 | 0.03 | 1.08 | 0.87 | 1.10 | 0.90 | 1.16 | 2.27 | 2.80 | 2.46 | 1.24 | 0.24 |
| 85 | 3.33 | 1.24 | 1.70 | 1.29 | 0.18 | 1.55 | 1.44 | 0.06 | 1.26 | 0.21 | 0.27 | 0.34 | 0.22 | 0.32 | 1.36 | 2.62 | 1.62 | 2.61 | 0.31 | 5.85 |
| 86 | 1.61 | 0.66 | 1.15 | 1.42 | 0.05 | 1.14 | 1.09 | 0.01 | 0.00 | 0.23 | 0.21 | 0.19 | 0.17 | 0.18 | 0.51 | 1.23 | 0.91 | 1.02 | 0.18 | 0.90 |
| 87 | 3.34 | 0.38 | 2.20 | 1.80 | 0.29 | 1.92 | 1.80 | 0.08 | 1.18 | 0.08 | 0.37 | 0.47 | 0.40 | 0.44 | 2.05 | 2.25 | 2.22 | 2.27 | 0.50 | 1.91 |
| 88 | 3.09 | 0.02 | 1.10 | 3.29 | 0.32 | 3.90 | 3.61 | 0.12 | 0.14 | 0.01 | 0.39 | 0.86 | 0.44 | 0.76 | 3.30 | 3.95 | 3.86 | 4.88 | 0.60 | 0.19 |
| 89 | 0.63 | 0.03 | 1.59 | 0.37 | 1.58 | 0.44 | 0.49 | 0.73 | 0.45 | 0.00 | 1.86 | 1.40 | 2.00 | 1.50 | 0.72 | 0.59 | 0.69 | 0.41 | 0.53 | 0.01 |
| 90 | 2.04 | 0.23 | 1.79 | 3.00 | 0.30 | 3.18 | 2.97 | 0.07 | 0.00 | 0.11 | 0.33 | 0.55 | 0.38 | 0.51 | 1.42 | 2.48 | 2.80 | 2.55 | 0.46 | 0.33 |
| 91 | 2.09 | 0.07 | 5.87 | 0.34 | 0.88 | 0.40 | 0.41 | 0.24 | 0.60 | 0.01 | 1.32 | 0.85 | 1.22 | 0.91 | 1.08 | 1.69 | 1.68 | 0.74 | 0.71 | 0.06 |
| 92 | 1.78 | 0.94 | 1.62 | 1.30 | 0.20 | 1.08 | 1.03 | 0.04 | 8.59 | 0.38 | 0.22 | 0.25 | 0.24 | 0.24 | 1.30 | 0.94 | 1.27 | 1.04 | 0.37 | 3.84 |
| 93 | 1.44 | 2.10 | 1.70 | 1.18 | 0.28 | 1.01 | 0.97 | 0.08 | 8.78 | 1.46 | 0.30 | 0.22 | 0.23 | 0.22 | 3.35 | 0.72 | 1.51 | 1.11 | 0.46 | 7.79 |
| 94 | 0.78 | 4.07 | 2.34 | 0.49 | 0.16 | 0.59 | 0.55 | 0.20 | 2.41 | 2.78 | 0.18 | 0.18 | 0.28 | 0.19 | 3.88 | 0.67 | 0.81 | 0.51 | 0.42 | 11.88 |
| 95 | 1.44 | 4.54 | 9.49 | 0.55 | 0.25 | 0.45 | 0.44 | 0.06 | 39.31 | 2.21 | 0.24 | 0.28 | 0.26 | 0.28 | 6.24 | 0.83 | 1.29 | 1.38 | 0.38 | 21.02 |
| 96 | 0.10 | 7.52 | 0.77 | 0.01 | 0.01 | 0.03 | 0.03 | 0.01 | 0.80 | 9.30 | 0.01 | 0.02 | 0.02 | 0.02 | 0.22 | 0.06 | 0.09 | 0.02 | 0.02 | 1.79 |
| 97 | 0.84 | 10.79 | 3.26 | 0.43 | 0.12 | 0.40 | 0.38 | 0.16 | 0.02 | 7.01 | 0.07 | 0.14 | 0.11 | 0.13 | 2.20 | 0.86 | 0.43 | 0.21 | 0.11 | 10.14 |
| 98 | 1.95 | 4.32 | 2.81 | 0.47 | 0.13 | 0.71 | 0.65 | 0.00 | 5.36 | 2.78 | 0.09 | 0.20 | 0.12 | 0.18 | 2.60 | 0.89 | 0.99 | 1.57 | 0.14 | 6.27 |
| 99 | 2.26 | 1.82 | 2.55 | 0.69 | 0.18 | 0.74 | 0.69 | 0.04 | 3.27 | 1.11 | 0.15 | 0.21 | 0.24 | 0.21 | 0.50 | 1.01 | 0.83 | 1.04 | 0.22 | 5.30 |
| 100 | 0.01 | 0.00 | 0.44 | 0.02 | 0.00 | 0.00 | 0.00 | 0.00 | 0.00 | 0.00 | 0.00 | 0.00 | 0.00 | 0.00 | 2.51 | 0.00 | 0.14 | 0.00 | 5.39 | 0.12 |
| Base | 12.65 | 0.40 | 0.84 | 0.09 | 0.05 | 1.10 | 1.27 | 0.04 | 0.00 | 0.27 | 0.11 | 1.45 | 0.21 | 1.84 | 0.01 | 0.00 | 0.01 | 0.01 | 0.01 | 0.01 |

**Figure S14: Conservation state enrichments for RepeatMasker classes and families of repeats.** The rows correspond to different conservation states and columns correspond to different repeat classes or families. The first 16 columns are repeat classes, and the remaining are repeat families. The values correspond to fold enrichment for the repeat class or family for the conservation state. Values are shaded in a column specific manner. The last row gives the % of the genome the repeat class or family covers.

| state | 1_TssA | 2_PromU | 3_PromD1 | 4_PromD2 | 5_Tx5 | 6_Tx | 7_Tx3 | 8_TxWk | 9_TxReg | 10_TxEnh5 | 11_TxEnh3 | 12_TxEnhW | 13_EnhA1 | 14_EnhA2 | 15_EnhAF | 16_EnhW1 | 17_EnhW2 | 18_EnhAc | 19_DNase | 20_ZNF/Rpts | 21_Het | 22_PromP | 23_PromBiv | 24_ReprPC | 25_Quies |
|---|---|---|---|---|---|---|---|---|---|---|---|---|---|---|---|---|---|---|---|---|---|---|---|---|---|
| 1 | 2.1 | 3.4 | 5.4 | 3.8 | 0.3 | 8.6 | 8.7 | 1.6 | 6.7 | 1.3 | 8.5 | 0.8 | 1.4 | 1.6 | 1.0 | 1.2 | 1.0 | 0.5 | 1.0 | 4.0 | 0.3 | 1.6 | 6.3 | 1.6 | 0.5 |
| 2 | 1.0 | 2.5 | 1.1 | 1.7 | 0.3 | 2.0 | 2.8 | 1.2 | 2.7 | 1.6 | 4.2 | 1.1 | 3.5 | 4.7 | 3.1 | 3.3 | 3.4 | 1.2 | 1.9 | 0.4 | 0.1 | 0.7 | 3.4 | 1.6 | 0.8 |
| 3 | 1.3 | 2.7 | 2.3 | 2.5 | 0.3 | 4.2 | 4.9 | 1.4 | 3.7 | 1.5 | 5.8 | 1.0 | 2.6 | 3.1 | 2.2 | 2.4 | 2.3 | 1.0 | 1.5 | 3.2 | 0.3 | 1.2 | 4.1 | 1.6 | 0.7 |
| 4 | 4.7 | 4.0 | 5.7 | 3.0 | 0.4 | 2.8 | 3.0 | 1.1 | 4.7 | 2.0 | 3.7 | 1.3 | 2.5 | 3.6 | 2.3 | 3.0 | 2.4 | 1.1 | 2.1 | 0.8 | 0.2 | 1.4 | 4.8 | 1.7 | 0.8 |
| 5 | 3.6 | 3.5 | 3.8 | 2.7 | 0.6 | 1.1 | 1.3 | 1.0 | 3.9 | 2.3 | 2.6 | 1.7 | 3.3 | 4.2 | 2.9 | 3.1 | 2.9 | 1.6 | 2.5 | 0.2 | 0.1 | 1.0 | 3.8 | 2.0 | 0.9 |
| 6 | 1.4 | 2.2 | 2.2 | 2.7 | 1.0 | 0.9 | 1.0 | 1.0 | 2.3 | 2.1 | 1.6 | 2.1 | 2.0 | 2.3 | 2.0 | 2.1 | 2.2 | 1.2 | 2.0 | 0.2 | 0.1 | 0.8 | 2.1 | 1.9 | 1.0 |
| 7 | 5.4 | 3.9 | 5.9 | 2.3 | 0.5 | 2.9 | 4.5 | 1.8 | 3.6 | 1.7 | 4.6 | 1.0 | 2.1 | 2.0 | 1.7 | 1.8 | 1.7 | 1.0 | 1.2 | 1.3 | 0.2 | 1.5 | 5.6 | 1.6 | 0.7 |
| 8 | 2.6 | 2.6 | 2.9 | 2.2 | 0.6 | 1.4 | 1.9 | 1.1 | 3.1 | 2.0 | 2.9 | 1.5 | 2.7 | 3.2 | 2.5 | 2.4 | 2.5 | 1.3 | 2.0 | 0.4 | 0.1 | 0.9 | 3.0 | 1.7 | 0.9 |
| 9 | 1.3 | 2.0 | 2.1 | 2.4 | 1.0 | 0.9 | 1.0 | 1.0 | 2.1 | 2.0 | 1.6 | 2.0 | 1.8 | 2.1 | 2.0 | 2.0 | 2.1 | 1.1 | 1.9 | 0.2 | 0.1 | 0.7 | 2.1 | 1.9 | 1.0 |
| 10 | 1.4 | 2.1 | 2.0 | 2.1 | 1.0 | 1.1 | 1.3 | 1.1 | 2.1 | 2.0 | 1.8 | 1.9 | 1.9 | 2.2 | 1.9 | 1.9 | 2.0 | 1.1 | 1.8 | 0.3 | 0.2 | 0.7 | 2.0 | 1.7 | 1.0 |
| 11 | 1.1 | 1.6 | 1.4 | 1.6 | 1.0 | 1.0 | 1.3 | 1.1 | 1.4 | 1.6 | 1.4 | 1.6 | 1.5 | 1.5 | 1.4 | 1.6 | 1.6 | 1.0 | 1.5 | 0.4 | 0.2 | 0.6 | 1.4 | 1.5 | 1.0 |
| 12 | 0.8 | 1.2 | 1.1 | 1.3 | 0.9 | 0.7 | 0.8 | 0.9 | 1.1 | 1.2 | 1.0 | 1.3 | 1.3 | 1.4 | 1.3 | 1.3 | 1.5 | 0.9 | 1.4 | 0.3 | 0.3 | 0.6 | 1.3 | 1.3 | 1.0 |
| 13 | 0.9 | 1.4 | 1.4 | 1.3 | 1.0 | 0.8 | 0.9 | 1.0 | 1.0 | 1.3 | 1.0 | 1.5 | 1.2 | 1.4 | 1.4 | 1.3 | 1.6 | 0.9 | 1.4 | 0.3 | 0.2 | 0.6 | 1.2 | 1.3 | 1.0 |
| 14 | 1.7 | 3.2 | 2.7 | 2.3 | 1.3 | 1.4 | 1.5 | 1.2 | 2.0 | 2.4 | 1.9 | 2.2 | 2.4 | 2.3 | 1.9 | 2.4 | 1.9 | 1.3 | 2.0 | 0.3 | 0.2 | 0.8 | 2.5 | 1.7 | 0.9 |
| 15 | 1.6 | 2.5 | 2.5 | 2.5 | 1.0 | 1.1 | 1.2 | 1.0 | 2.5 | 2.2 | 1.8 | 2.0 | 2.0 | 2.4 | 2.0 | 2.2 | 2.2 | 1.1 | 2.0 | 0.3 | 0.1 | 0.8 | 2.5 | 2.0 | 0.9 |
| 16 | 0.8 | 1.3 | 1.1 | 1.3 | 1.1 | 1.0 | 0.9 | 1.0 | 1.3 | 1.6 | 1.1 | 1.7 | 1.2 | 1.2 | 1.3 | 1.3 | 1.4 | 0.8 | 1.4 | 0.3 | 0.3 | 0.7 | 1.1 | 1.3 | 1.0 |
| 17 | 0.7 | 1.2 | 0.9 | 1.4 | 1.2 | 1.0 | 0.9 | 1.0 | 1.1 | 1.5 | 1.0 | 1.7 | 1.1 | 1.1 | 1.2 | 1.2 | 1.4 | 0.8 | 1.3 | 0.4 | 0.3 | 0.7 | 0.9 | 1.2 | 1.0 |
| 18 | 0.9 | 1.4 | 1.2 | 1.3 | 1.1 | 1.0 | 1.0 | 1.0 | 1.3 | 1.7 | 1.2 | 1.7 | 1.3 | 1.4 | 1.4 | 1.4 | 1.5 | 0.9 | 1.5 | 0.4 | 0.3 | 0.7 | 1.2 | 1.3 | 1.0 |
| 19 | 1.8 | 2.4 | 2.4 | 2.4 | 0.9 | 1.1 | 1.3 | 1.0 | 2.5 | 2.2 | 2.1 | 2.0 | 2.2 | 2.7 | 2.2 | 2.2 | 2.3 | 1.2 | 2.1 | 0.3 | 0.1 | 0.8 | 2.6 | 1.9 | 0.9 |
| 20 | 3.7 | 1.9 | 3.1 | 1.4 | 0.9 | 0.9 | 1.1 | 1.0 | 1.4 | 1.4 | 1.3 | 1.5 | 1.4 | 1.5 | 1.5 | 1.4 | 1.6 | 0.8 | 1.4 | 0.3 | 0.2 | 0.9 | 2.2 | 1.3 | 1.0 |
| 21 | 1.0 | 1.4 | 1.4 | 1.4 | 0.9 | 1.0 | 1.2 | 1.0 | 1.5 | 1.5 | 1.6 | 1.5 | 1.5 | 1.7 | 1.5 | 1.5 | 1.6 | 0.8 | 1.4 | 0.4 | 0.3 | 0.6 | 1.4 | 1.4 | 1.0 |
| 22 | 2.5 | 1.9 | 2.2 | 1.1 | 1.2 | 1.2 | 1.2 | 1.1 | 1.0 | 1.7 | 1.4 | 1.5 | 1.2 | 1.3 | 1.3 | 1.5 | 1.3 | 0.8 | 1.4 | 0.4 | 0.3 | 1.0 | 1.8 | 1.2 | 1.0 |
| 23 | 0.8 | 1.1 | 0.7 | 0.8 | 1.6 | 1.6 | 1.2 | 1.2 | 1.1 | 1.4 | 1.3 | 1.4 | 1.0 | 0.8 | 1.0 | 1.1 | 0.9 | 0.8 | 1.1 | 1.0 | 0.5 | 0.7 | 0.7 | 0.9 | 1.0 |
| 24 | 0.5 | 0.7 | 0.5 | 0.7 | 1.2 | 1.2 | 1.0 | 1.0 | 0.9 | 1.3 | 1.0 | 1.2 | 0.9 | 0.8 | 0.9 | 0.9 | 0.9 | 0.5 | 1.1 | 0.7 | 0.6 | 0.6 | 0.5 | 0.9 | 1.0 |
| 25 | 0.5 | 0.6 | 0.4 | 0.6 | 1.1 | 1.0 | 0.9 | 0.9 | 0.7 | 1.0 | 0.7 | 1.1 | 0.7 | 0.9 | 0.9 | 0.9 | 1.0 | 0.6 | 1.0 | 0.7 | 0.5 | 0.6 | 0.5 | 0.9 | 1.1 |
| 26 | 0.7 | 0.8 | 0.8 | 0.9 | 1.3 | 1.2 | 1.2 | 1.1 | 0.9 | 1.2 | 1.2 | 1.4 | 0.9 | 0.9 | 1.0 | 1.0 | 1.1 | 0.7 | 1.1 | 0.7 | 0.4 | 0.6 | 0.6 | 0.9 | 1.0 |
| 27 | 0.9 | 1.3 | 1.1 | 1.3 | 1.4 | 1.5 | 1.3 | 1.2 | 1.5 | 1.8 | 1.4 | 1.9 | 1.2 | 1.3 | 1.3 | 1.4 | 1.4 | 0.8 | 1.4 | 0.7 | 0.3 | 0.7 | 0.9 | 1.2 | 1.0 |
| 28 | 35.1 | 4.5 | 19.9 | 2.4 | 0.5 | 2.2 | 2.4 | 1.0 | 3.6 | 1.3 | 2.0 | 0.6 | 1.3 | 1.3 | 0.8 | 1.3 | 0.7 | 0.9 | 1.1 | 5.4 | 1.4 | 8.0 | 13.1 | 1.9 | 0.6 |
| 29 | 0.6 | 0.5 | 0.5 | 0.5 | 0.9 | 0.6 | 0.7 | 0.8 | 0.6 | 0.8 | 0.6 | 1.0 | 0.7 | 0.8 | 1.0 | 0.8 | 1.1 | 0.5 | 1.0 | 0.5 | 0.5 | 0.5 | 0.5 | 0.9 | 1.1 |
| 30 | 0.3 | 0.4 | 0.2 | 0.4 | 1.1 | 0.8 | 0.7 | 0.9 | 0.4 | 0.7 | 0.5 | 0.9 | 0.7 | 0.6 | 0.7 | 0.7 | 0.9 | 0.6 | 0.9 | 0.7 | 0.6 | 0.4 | 0.3 | 0.7 | 1.1 |
| 31 | 0.8 | 0.8 | 0.6 | 0.5 | 1.2 | 1.0 | 0.9 | 1.1 | 0.4 | 1.0 | 0.7 | 1.1 | 0.9 | 0.7 | 0.8 | 0.9 | 0.9 | 0.8 | 1.1 | 0.9 | 0.7 | 0.7 | 0.6 | 0.9 | 1.0 |
| 32 | 0.5 | 0.5 | 0.3 | 0.4 | 1.3 | 1.0 | 0.7 | 1.1 | 0.5 | 0.9 | 0.6 | 0.8 | 0.6 | 0.6 | 0.6 | 0.7 | 0.6 | 0.6 | 0.8 | 1.5 | 1.1 | 0.7 | 0.5 | 0.9 | 1.0 |
| 33 | 1.1 | 0.4 | 0.4 | 0.4 | 1.0 | 0.8 | 0.7 | 1.0 | 0.3 | 0.4 | 0.5 | 0.6 | 0.5 | 0.4 | 0.5 | 0.6 | 0.6 | 0.4 | 0.7 | 1.1 | 1.5 | 1.2 | 0.5 | 0.8 | 1.0 |
| 34 | 0.7 | 0.8 | 0.6 | 1.1 | 1.3 | 1.0 | 1.0 | 1.1 | 0.9 | 1.0 | 1.0 | 1.2 | 0.6 | 0.6 | 0.8 | 1.0 | 0.9 | 0.5 | 1.0 | 0.6 | 0.7 | 0.7 | 0.7 | 1.0 | 1.0 |
| 35 | 1.3 | 2.1 | 1.7 | 1.7 | 1.5 | 1.8 | 1.6 | 1.4 | 1.9 | 2.1 | 1.7 | 2.0 | 1.7 | 1.6 | 1.5 | 1.7 | 1.4 | 1.1 | 1.6 | 0.6 | 0.3 | 0.8 | 1.4 | 1.4 | 0.9 |
| 36 | 0.4 | 0.7 | 0.4 | 0.8 | 1.5 | 1.3 | 1.0 | 1.1 | 0.9 | 1.3 | 0.9 | 1.4 | 0.8 | 0.8 | 0.9 | 1.0 | 1.0 | 0.7 | 1.1 | 0.7 | 0.5 | 0.7 | 0.4 | 0.9 | 1.0 |
| 37 | 0.8 | 1.2 | 1.1 | 1.6 | 1.4 | 1.4 | 1.3 | 1.2 | 1.5 | 1.7 | 1.4 | 1.9 | 1.3 | 1.3 | 1.4 | 1.4 | 1.5 | 0.8 | 1.4 | 0.5 | 0.2 | 0.6 | 0.9 | 1.2 | 1.0 |
| 38 | 0.6 | 0.8 | 0.7 | 0.9 | 1.4 | 1.2 | 1.1 | 1.2 | 0.8 | 1.2 | 0.9 | 1.6 | 0.8 | 0.8 | 1.0 | 1.0 | 1.2 | 0.7 | 1.1 | 0.8 | 0.3 | 0.5 | 0.5 | 0.9 | 1.0 |
| 39 | 0.4 | 0.7 | 0.5 | 0.7 | 1.1 | 0.8 | 0.8 | 1.0 | 0.5 | 0.8 | 0.7 | 1.2 | 0.8 | 0.7 | 0.9 | 0.8 | 1.1 | 0.6 | 1.0 | 0.8 | 0.6 | 0.6 | 0.5 | 0.9 | 1.0 |
| 40 | 0.3 | 0.4 | 0.2 | 0.4 | 1.2 | 0.9 | 0.7 | 0.9 | 0.4 | 0.8 | 0.5 | 1.1 | 0.6 | 0.5 | 0.7 | 0.7 | 0.9 | 0.5 | 0.9 | 0.8 | 0.6 | 0.4 | 0.2 | 0.8 | 1.1 |
| 41 | 0.4 | 0.6 | 0.4 | 0.7 | 1.0 | 0.7 | 0.6 | 0.9 | 0.5 | 1.0 | 0.6 | 1.1 | 0.7 | 0.7 | 0.9 | 0.9 | 1.1 | 0.6 | 1.1 | 0.4 | 0.4 | 0.5 | 0.5 | 1.1 | 1.1 |
| 42 | 0.8 | 1.4 | 0.8 | 1.0 | 1.4 | 1.1 | 1.0 | 1.1 | 0.9 | 1.4 | 1.0 | 1.5 | 1.1 | 1.1 | 1.1 | 1.4 | 1.2 | 1.0 | 1.4 | 0.5 | 0.5 | 0.7 | 1.0 | 1.3 | 1.0 |
| 43 | 0.7 | 0.9 | 0.6 | 0.7 | 1.5 | 1.2 | 1.0 | 1.2 | 0.6 | 1.3 | 1.1 | 1.4 | 0.9 | 0.9 | 1.0 | 1.1 | 1.0 | 0.9 | 1.2 | 1.0 | 0.5 | 0.7 | 0.6 | 1.0 | 1.0 |
| 44 | 0.8 | 0.7 | 0.5 | 0.6 | 1.5 | 1.2 | 0.9 | 1.1 | 0.6 | 0.7 | 0.6 | 0.9 | 0.7 | 0.6 | 0.7 | 0.8 | 0.6 | 0.7 | 1.0 | 1.5 | 1.2 | 1.1 | 0.7 | 1.1 | 1.0 |
| 45 | 0.5 | 0.7 | 0.4 | 0.6 | 1.1 | 0.9 | 0.8 | 1.0 | 0.5 | 0.8 | 0.7 | 0.9 | 0.6 | 0.7 | 0.8 | 0.8 | 0.9 | 0.6 | 1.0 | 0.7 | 0.6 | 0.5 | 0.5 | 1.0 | 1.1 |
| 46 | 0.5 | 0.8 | 0.5 | 0.9 | 1.3 | 1.2 | 1.2 | 1.2 | 0.8 | 1.0 | 1.1 | 1.3 | 0.9 | 0.7 | 0.9 | 1.0 | 1.0 | 0.9 | 1.1 | 0.6 | 0.5 | 0.6 | 0.5 | 1.0 | 1.0 |
| 47 | 1.4 | 1.8 | 1.6 | 1.4 | 1.2 | 1.4 | 1.3 | 1.1 | 1.9 | 2.1 | 1.6 | 1.9 | 1.6 | 1.7 | 1.5 | 1.7 | 1.6 | 1.0 | 1.7 | 0.5 | 0.3 | 0.8 | 1.6 | 1.4 | 1.0 |
| 48 | 0.6 | 0.9 | 0.6 | 0.9 | 1.3 | 1.2 | 0.9 | 1.0 | 1.0 | 1.3 | 1.0 | 1.5 | 0.9 | 0.9 | 1.0 | 1.0 | 1.1 | 0.7 | 1.2 | 0.8 | 0.5 | 0.7 | 0.6 | 1.0 | 1.0 |
| 49 | 1.3 | 1.9 | 1.7 | 1.6 | 1.3 | 1.6 | 1.5 | 1.2 | 2.0 | 2.1 | 1.7 | 2.0 | 1.6 | 1.6 | 1.5 | 1.7 | 1.6 | 0.9 | 1.6 | 0.6 | 0.2 | 0.7 | 1.4 | 1.4 | 0.9 |
| 50 | 2.2 | 2.7 | 2.9 | 2.5 | 0.8 | 1.1 | 1.4 | 1.0 | 2.7 | 2.1 | 2.1 | 1.8 | 2.2 | 2.6 | 2.1 | 2.2 | 2.2 | 1.2 | 2.0 | 0.3 | 0.1 | 0.9 | 3.0 | 2.0 | 0.9 |
| 51 | 1.1 | 1.7 | 1.4 | 1.4 | 1.1 | 1.0 | 1.0 | 1.0 | 1.4 | 1.7 | 1.2 | 1.7 | 1.4 | 1.5 | 1.4 | 1.5 | 1.5 | 0.9 | 1.5 | 0.4 | 0.3 | 0.7 | 1.5 | 1.4 | 1.0 |
| 52 | 1.2 | 1.8 | 1.5 | 1.5 | 1.1 | 1.1 | 1.1 | 1.0 | 1.6 | 1.9 | 1.3 | 1.8 | 1.5 | 1.6 | 1.5 | 1.6 | 1.6 | 1.0 | 1.6 | 0.4 | 0.3 | 0.8 | 1.6 | 1.4 | 1.0 |
| 53 | 2.3 | 2.9 | 3.0 | 2.5 | 0.8 | 1.3 | 1.6 | 1.1 | 2.9 | 2.3 | 2.4 | 1.9 | 2.3 | 2.8 | 2.2 | 2.4 | 2.3 | 1.2 | 2.1 | 0.4 | 0.1 | 0.9 | 3.2 | 2.0 | 0.9 |
| 54 | 2.5 | 3.0 | 3.3 | 2.2 | 0.3 | 3.8 | 4.8 | 1.4 | 3.7 | 1.5 | 5.2 | 0.9 | 2.3 | 2.6 | 1.8 | 2.2 | 1.9 | 1.0 | 1.5 | 2.8 | 0.4 | 1.3 | 4.4 | 1.6 | 0.7 |
| 55 | 2.4 | 2.5 | 2.8 | 2.3 | 0.8 | 1.2 | 1.4 | 1.0 | 2.5 | 2.0 | 2.1 | 1.8 | 2.1 | 2.5 | 2.1 | 2.2 | 2.2 | 1.1 | 2.0 | 0.3 | 0.1 | 0.8 | 2.8 | 1.9 | 0.9 |
| 56 | 1.4 | 1.4 | 1.4 | 1.2 | 1.0 | 0.9 | 0.9 | 0.9 | 1.2 | 1.5 | 1.1 | 1.5 | 1.2 | 1.3 | 1.3 | 1.4 | 1.4 | 0.8 | 1.4 | 0.4 | 0.4 | 0.7 | 1.3 | 1.3 | 1.0 |
| 57 | 1.1 | 1.2 | 1.0 | 1.0 | 1.3 | 1.3 | 1.1 | 1.1 | 1.2 | 1.5 | 1.1 | 1.5 | 1.1 | 1.0 | 1.1 | 1.2 | 1.2 | 0.8 | 1.3 | 0.7 | 0.5 | 0.7 | 0.9 | 1.1 | 1.0 |
| 58 | 1.0 | 1.2 | 1.0 | 0.9 | 1.1 | 1.1 | 1.1 | 1.0 | 1.2 | 1.3 | 1.2 | 1.3 | 1.1 | 1.1 | 1.1 | 1.2 | 1.1 | 0.8 | 1.3 | 0.6 | 0.5 | 0.7 | 1.0 | 1.2 | 1.0 |
| 59 | 1.3 | 1.3 | 1.0 | 0.8 | 1.4 | 1.4 | 1.0 | 1.1 | 1.0 | 1.4 | 1.1 | 1.3 | 1.0 | 1.0 | 0.9 | 1.2 | 0.9 | 0.9 | 1.3 | 1.3 | 0.9 | 1.1 | 1.1 | 1.2 | 1.0 |
| 60 | 0.5 | 0.7 | 0.4 | 0.5 | 1.1 | 0.9 | 0.7 | 0.9 | 0.6 | 1.1 | 0.6 | 1.2 | 0.8 | 0.8 | 0.9 | 0.9 | 1.0 | 0.6 | 1.1 | 0.7 | 0.6 | 0.5 | 0.5 | 1.1 | 1.1 |
| 61 | 0.7 | 0.7 | 0.5 | 0.5 | 1.0 | 0.9 | 0.8 | 0.9 | 0.6 | 1.0 | 0.7 | 1.1 | 0.9 | 0.9 | 0.9 | 0.9 | 1.0 | 0.6 | 1.1 | 0.6 | 0.6 | 0.5 | 0.6 | 0.9 | 1.1 |
| 62 | 0.9 | 1.0 | 0.8 | 0.8 | 1.3 | 1.5 | 1.2 | 1.0 | 1.2 | 1.6 | 1.2 | 1.4 | 1.0 | 1.0 | 1.0 | 1.1 | 1.0 | 0.7 | 1.2 | 0.8 | 0.5 | 0.7 | 0.7 | 1.0 | 1.0 |
| 63 | 3.2 | 2.0 | 2.8 | 1.3 | 0.9 | 1.1 | 1.3 | 1.0 | 1.6 | 1.7 | 1.6 | 1.5 | 1.5 | 1.7 | 1.5 | 1.6 | 1.6 | 0.8 | 1.5 | 0.4 | 0.3 | 0.9 | 2.4 | 1.4 | 1.0 |
| 64 | 0.8 | 0.6 | 0.5 | 0.6 | 1.0 | 0.8 | 0.9 | 0.9 | 0.5 | 0.9 | 0.7 | 0.9 | 0.8 | 0.7 | 0.9 | 0.8 | 1.0 | 0.7 | 1.0 | 0.6 | 0.5 | 0.6 | 0.5 | 0.9 | 1.1 |
| 65 | 0.9 | 0.9 | 0.7 | 0.7 | 1.2 | 1.1 | 1.0 | 1.0 | 0.8 | 1.2 | 0.9 | 1.2 | 0.8 | 0.8 | 0.9 | 1.0 | 0.9 | 0.8 | 1.0 | 0.7 | 0.6 | 0.7 | 0.7 | 1.0 | 1.0 |
| 66 | 1.4 | 1.5 | 1.3 | 1.4 | 1.0 | 1.1 | 1.2 | 1.1 | 1.5 | 1.6 | 1.3 | 1.5 | 1.3 | 1.4 | 1.3 | 1.4 | 1.4 | 0.7 | 1.4 | 0.4 | 0.3 | 0.7 | 1.4 | 1.4 | 1.0 |
| 67 | 1.3 | 1.1 | 1.0 | 1.0 | 1.0 | 1.0 | 1.1 | 1.0 | 1.1 | 1.2 | 1.1 | 1.1 | 1.1 | 1.2 | 1.1 | 1.0 | 1.1 | 0.9 | 1.1 | 0.5 | 0.5 | 0.9 | 1.2 | 1.2 | 1.0 |
| 68 | 0.4 | 0.4 | 0.3 | 0.6 | 1.2 | 1.0 | 0.9 | 1.0 | 0.6 | 0.8 | 0.6 | 0.9 | 0.6 | 0.6 | 0.7 | 0.6 | 0.8 | 0.7 | 0.7 | 0.8 | 0.7 | 0.6 | 0.3 | 0.8 | 1.1 |
| 69 | 0.4 | 0.4 | 0.2 | 0.4 | 1.1 | 0.9 | 0.7 | 1.0 | 0.4 | 0.6 | 0.5 | 0.7 | 0.6 | 0.4 | 0.6 | 0.6 | 0.7 | 0.7 | 0.8 | 1.0 | 1.1 | 0.7 | 0.3 | 0.9 | 1.1 |
| 70 | 0.8 | 0.7 | 0.5 | 0.5 | 1.2 | 1.1 | 0.8 | 1.0 | 0.6 | 1.0 | 0.7 | 0.9 | 0.7 | 0.6 | 0.7 | 0.9 | 0.7 | 0.7 | 1.0 | 1.2 | 1.1 | 1.0 | 0.7 | 1.0 | 1.0 |
| 71 | 1.1 | 0.8 | 0.8 | 0.7 | 1.3 | 1.3 | 1.1 | 1.1 | 0.8 | 1.2 | 0.8 | 1.0 | 0.8 | 0.7 | 0.8 | 0.9 | 0.8 | 0.9 | 1.0 | 1.0 | 0.8 | 0.9 | 0.7 | 1.0 | 1.0 |
| 72 | 0.6 | 0.6 | 0.4 | 0.6 | 1.3 | 1.1 | 0.9 | 1.1 | 0.5 | 0.8 | 0.6 | 0.8 | 0.6 | 0.5 | 0.6 | 0.7 | 0.6 | 0.8 | 0.8 | 1.2 | 1.0 | 0.8 | 0.4 | 0.9 | 1.0 |
| 73 | 0.6 | 0.5 | 0.3 | 0.8 | 1.6 | 1.2 | 1.1 | 1.4 | 0.4 | 0.7 | 0.6 | 0.9 | 0.6 | 0.4 | 0.7 | 0.6 | 0.6 | 1.2 | 0.7 | 1.2 | 1.0 | 0.9 | 0.5 | 1.2 | 1.0 |
| 74 | 0.4 | 0.6 | 0.5 | 0.9 | 1.6 | 1.4 | 1.4 | 1.3 | 0.8 | 0.9 | 0.8 | 1.2 | 0.8 | 0.7 | 0.9 | 0.7 | 0.9 | 0.9 | 0.8 | 1.1 | 0.6 | 0.7 | 0.5 | 1.0 | 1.0 |
| 75 | 0.3 | 0.4 | 0.2 | 0.9 | 1.6 | 1.2 | 1.3 | 1.5 | 0.5 | 0.8 | 0.6 | 0.9 | 0.6 | 0.4 | 0.7 | 0.6 | 0.7 | 1.2 | 0.6 | 1.0 | 0.6 | 0.8 | 0.3 | 1.0 | 1.0 |
| 76 | 0.1 | 0.3 | 0.1 | 0.4 | 1.1 | 0.8 | 0.7 | 1.0 | 0.4 | 0.5 | 0.5 | 0.6 | 1.0 | 0.7 | 1.0 | 0.7 | 0.8 | 1.5 | 0.7 | 1.8 | 2.0 | 0.6 | 0.1 | 0.6 | 1.0 |
| 77 | 0.5 | 0.4 | 0.3 | 0.8 | 2.1 | 1.2 | 1.4 | 2.0 | 0.2 | 0.4 | 0.5 | 0.6 | 0.6 | 0.3 | 0.6 | 0.4 | 0.4 | 1.6 | 0.4 | 1.1 | 1.0 | 1.2 | 0.2 | 0.8 | 0.9 |
| 78 | 0.4 | 0.5 | 0.3 | 0.7 | 1.4 | 1.1 | 0.9 | 1.2 | 0.4 | 0.4 | 0.7 | 0.6 | 0.8 | 0.5 | 0.8 | 0.7 | 0.6 | 1.3 | 0.7 | 2.1 | 1.8 | 1.0 | 0.3 | 0.7 | 1.0 |
| 79 | 0.2 | 0.3 | 0.1 | 0.4 | 1.1 | 0.8 | 0.6 | 1.0 | 0.4 | 0.4 | 0.5 | 0.4 | 0.9 | 0.5 | 0.8 | 0.7 | 0.5 | 1.6 | 0.7 | 2.7 | 2.8 | 0.9 | 0.2 | 0.7 | 1.0 |
| 80 | 0.5 | 0.4 | 0.2 | 0.8 | 1.9 | 1.2 | 1.3 | 1.7 | 0.5 | 0.6 | 0.4 | 0.7 | 0.5 | 0.3 | 0.5 | 0.4 | 0.5 | 1.5 | 0.5 | 1.0 | 0.7 | 1.0 | 0.2 | 0.8 | 1.0 |
| 81 | 0.5 | 0.5 | 0.2 | 0.4 | 1.3 | 1.0 | 0.8 | 1.3 | 0.4 | 0.4 | 0.5 | 0.4 | 0.4 | 0.3 | 0.6 | 0.5 | 0.4 | 1.3 | 0.6 | 2.3 | 2.0 | 1.2 | 0.5 | 1.0 | 1.0 |
| 82 | 4.2 | 1.2 | 1.7 | 0.9 | 0.9 | 1.6 | 1.6 | 1.3 | 0.9 | 0.7 | 1.0 | 0.5 | 0.4 | 0.6 | 0.5 | 0.8 | 0.4 | 1.1 | 1.1 | 6.4 | 3.5 | 4.8 | 2.1 | 2.1 | 0.8 |
| 83 | 0.2 | 0.1 | 0.1 | 0.4 | 0.7 | 0.4 | 0.4 | 0.8 | 0.1 | 0.2 | 0.2 | 0.2 | 0.4 | 0.3 | 0.4 | 0.2 | 0.3 | 1.1 | 0.4 | 2.0 | 2.5 | 0.6 | 0.1 | 0.4 | 1.1 |
| 84 | 0.4 | 0.2 | 0.1 | 0.6 | 1.1 | 0.7 | 0.7 | 1.1 | 0.1 | 0.2 | 0.2 | 0.3 | 0.3 | 0.2 | 0.3 | 0.2 | 0.2 | 0.7 | 0.3 | 1.0 | 1.6 | 1.1 | 0.3 | 0.8 | 1.0 |
| 85 | 0.2 | 0.1 | 0.1 | 0.3 | 1.3 | 0.7 | 0.6 | 1.4 | 0.1 | 0.2 | 0.2 | 0.2 | 0.4 | 0.2 | 0.3 | 0.2 | 0.2 | 1.1 | 0.3 | 1.6 | 2.1 | 1.0 | 0.1 | 0.5 | 1.0 |
| 86 | 0.1 | 0.2 | 0.0 | 0.2 | 0.7 | 0.4 | 0.3 | 0.9 | 0.1 | 0.1 | 0.1 | 0.1 | 0.3 | 0.2 | 0.4 | 0.3 | 0.2 | 1.0 | 0.3 | 1.9 | 2.5 | 0.7 | 0.1 | 0.4 | 1.1 |
| 87 | 0.4 | 0.4 | 0.2 | 0.6 | 1.5 | 1.0 | 0.9 | 1.3 | 0.4 | 0.4 | 0.5 | 0.5 | 0.6 | 0.4 | 0.6 | 0.6 | 0.4 | 1.5 | 0.6 | 2.0 | 2.0 | 1.1 | 0.3 | 0.8 | 1.0 |
| 88 | 0.3 | 0.4 | 0.3 | 0.7 | 1.5 | 1.1 | 1.1 | 1.3 | 0.5 | 0.7 | 0.6 | 0.8 | 0.8 | 0.6 | 0.8 | 0.6 | 0.7 | 1.3 | 0.7 | 1.3 | 1.3 | 0.7 | 0.3 | 0.8 | 1.0 |
| 89 | 1.4 | 0.8 | 0.9 | 0.7 | 1.1 | 1.1 | 1.0 | 1.0 | 0.8 | 1.1 | 0.8 | 1.0 | 0.8 | 0.8 | 0.8 | 0.9 | 0.8 | 0.8 | 1.0 | 0.9 | 0.9 | 1.0 | 0.8 | 1.1 | 1.0 |
| 90 | 0.2 | 0.3 | 0.1 | 0.4 | 1.1 | 0.7 | 0.6 | 1.0 | 0.2 | 0.3 | 0.4 | 0.4 | 0.6 | 0.3 | 0.6 | 0.5 | 0.5 | 0.9 | 0.5 | 1.8 | 2.6 | 0.7 | 0.2 | 0.7 | 1.0 |
| 91 | 0.8 | 0.3 | 0.2 | 0.5 | 1.3 | 0.8 | 0.8 | 1.3 | 0.2 | 0.4 | 0.3 | 0.5 | 0.4 | 0.3 | 0.4 | 0.5 | 0.4 | 1.1 | 0.6 | 1.0 | 1.4 | 1.5 | 0.4 | 1.0 | 1.0 |
| 92 | 0.2 | 0.2 | 0.1 | 0.3 | 0.7 | 0.4 | 0.4 | 0.9 | 0.3 | 0.2 | 0.3 | 0.2 | 0.7 | 0.4 | 0.6 | 0.4 | 0.4 | 3.0 | 0.5 | 1.9 | 3.1 | 1.2 | 0.2 | 0.5 | 1.1 |
| 93 | 0.2 | 0.3 | 0.1 | 0.1 | 0.5 | 0.2 | 0.3 | 0.7 | 0.1 | 0.2 | 0.2 | 0.1 | 0.7 | 0.3 | 0.6 | 0.4 | 0.4 | 2.5 | 0.5 | 2.0 | 3.9 | 1.4 | 0.3 | 0.8 | 1.1 |
| 94 | 0.7 | 0.2 | 0.1 | 0.3 | 0.3 | 0.2 | 0.2 | 0.7 | 0.2 | 0.1 | 0.3 | 0.1 | 0.7 | 0.2 | 0.5 | 0.3 | 0.3 | 3.0 | 0.5 | 1.1 | 3.3 | 5.2 | 0.1 | 0.4 | 1.1 |
| 95 | 0.8 | 0.3 | 0.2 | 0.5 | 0.5 | 0.3 | 0.4 | 0.9 | 0.1 | 0.1 | 0.2 | 0.2 | 0.4 | 0.2 | 0.3 | 0.3 | 0.2 | 1.7 | 0.5 | 1.1 | 3.0 | 2.9 | 0.6 | 0.9 | 1.0 |
| 96 | 0.0 | 0.0 | 0.0 | 0.2 | 0.0 | 0.0 | 0.0 | 0.1 | 0.0 | 0.0 | 0.0 | 0.0 | 0.0 | 0.0 | 0.0 | 0.0 | 0.0 | 0.2 | 0.0 | 0.3 | 1.1 | 1.3 | 0.0 | 0.0 | 1.1 |
| 97 | 0.3 | 0.1 | 0.1 | 0.7 | 0.2 | 0.1 | 0.1 | 0.5 | 0.1 | 0.1 | 0.1 | 0.1 | 0.3 | 0.1 | 0.2 | 0.2 | 0.1 | 1.2 | 0.2 | 0.7 | 2.2 | 4.1 | 0.0 | 0.2 | 0.9 |
| 98 | 0.3 | 0.2 | 0.1 | 0.5 | 0.7 | 0.3 | 0.3 | 0.8 | 0.0 | 0.1 | 0.2 | 0.1 | 0.9 | 0.2 | 0.5 | 0.3 | 0.2 | 1.9 | 0.4 | 1.4 | 2.9 | 2.2 | 0.1 | 0.6 | 1.1 |
| 99 | 0.1 | 0.1 | 0.0 | 0.2 | 0.8 | 0.3 | 0.4 | 1.0 | 0.1 | 0.1 | 0.1 | 0.1 | 0.4 | 0.1 | 0.3 | 0.2 | 0.2 | 1.2 | 0.3 | 1.5 | 2.2 | 1.1 | 0.1 | 0.4 | 1.1 |
| 100 | 0.4 | 0.4 | 0.2 | 0.2 | 0.7 | 1.5 | 0.9 | 0.7 | 0.1 | 0.3 | 0.4 | 0.5 | 0.3 | 0.7 | 0.6 | 0.5 | 0.4 | 0.5 | 1.4 | 8.4 | 2.8 | 1.9 | 1.2 | 1.5 | 1.0 |
| Base | 0.23 | 0.28 | 0.38 | 0.23 | 1.89 | 0.63 | 3.84 | 5.49 | 0.07 | 0.12 | 0.09 | 0.45 | 0.13 | 0.08 | 0.46 | 0.25 | 2.06 | 0.04 | 0.44 | 0.24 | 2.59 | 0.13 | 0.42 | 1.35 | 76.19 |

**Figure S15: Conservation state enrichments for chromatin states.** This is an extended version of **Figure 4A**. Each row corresponds to one of the 100 conservation states. Each column corresponds to one of 25 chromatin states from the Roadmap Epigenomics imputation based chromatin state model defined across 127 samples of diverse cell and tissue types.[39] The values reported are the median fold enrichment across the 127 samples of the chromatin state for the conservation state. Values that are boxed were the maximum value of the column.

A

| | GERP++ elements | PhastCons elements | SiPhy-Omega elements | SiPhy-Pi elements |
|---|---|---|---|---|
| 1 | 14.32 | 17.94 | 21.98 | 16.49 |
| 2 | 12.62 | 14.19 | 16.40 | 13.18 |
| 3 | 11.98 | 9.39 | 14.71 | 12.60 |
| 4 | 10.99 | 10.45 | 13.54 | 11.55 |
| 5 | 9.84 | 9.08 | 11.27 | 9.77 |
| 6 | 3.37 | 2.92 | 3.22 | 3.48 |
| 7 | 3.77 | 2.09 | 3.10 | 3.56 |
| 8 | 5.77 | 2.75 | 4.02 | 4.48 |
| 9 | 2.09 | 1.01 | 1.44 | 1.99 |
| 10 | 1.65 | 0.65 | 0.99 | 1.62 |
| 11 | 0.85 | 0.64 | 0.54 | 0.84 |
| 12 | 1.09 | 0.84 | 0.70 | 1.19 |
| 13 | 1.35 | 1.05 | 0.99 | 1.29 |
| 14 | 1.17 | 0.94 | 0.88 | 1.21 |
| 15 | 1.61 | 0.61 | 0.96 | 1.64 |
| 16 | 0.34 | 0.19 | 0.16 | 0.35 |
| 17 | 0.65 | 0.77 | 0.49 | 0.64 |
| 18 | 0.41 | 0.18 | 0.18 | 0.47 |
| 19 | 2.86 | 1.22 | 1.94 | 2.70 |
| 20 | 1.34 | 1.00 | 1.00 | 1.29 |
| 21 | 1.02 | 0.72 | 0.65 | 0.88 |
| 22 | 0.33 | 0.40 | 0.24 | 0.35 |
| 23 | 0.09 | 0.37 | 0.09 | 0.07 |
| 24 | 0.09 | 0.28 | 0.08 | 0.07 |
| 25 | 0.11 | 0.31 | 0.11 | 0.11 |
| 26 | 0.31 | 0.44 | 0.22 | 0.26 |
| 27 | 0.34 | 0.20 | 0.15 | 0.30 |
| 28 | 3.00 | 3.27 | 2.27 | 1.91 |
| 29 | 0.16 | 0.28 | 0.12 | 0.20 |
| 30 | 0.12 | 0.29 | 0.09 | 0.11 |
| 31 | 0.09 | 0.31 | 0.07 | 0.10 |
| 32 | 0.03 | 0.36 | 0.03 | 0.05 |
| 33 | 0.07 | 0.41 | 0.07 | 0.06 |
| 34 | 0.47 | 0.65 | 0.37 | 0.44 |
| 35 | 0.51 | 0.64 | 0.35 | 0.39 |
| 36 | 0.17 | 0.47 | 0.16 | 0.11 |
| 37 | 0.91 | 0.89 | 0.65 | 0.77 |
| 38 | 0.45 | 0.56 | 0.30 | 0.32 |
| 39 | 0.33 | 0.45 | 0.18 | 0.34 |
| 40 | 0.09 | 0.33 | 0.08 | 0.10 |
| 41 | 0.24 | 0.38 | 0.17 | 0.27 |
| 42 | 0.17 | 0.36 | 0.15 | 0.23 |
| 43 | 0.12 | 0.39 | 0.10 | 0.14 |
| 44 | 0.05 | 0.46 | 0.05 | 0.05 |
| 45 | 0.08 | 0.30 | 0.08 | 0.15 |
| 46 | 0.13 | 0.38 | 0.13 | 0.12 |
| 47 | 0.31 | 0.05 | 0.08 | 0.51 |
| 48 | 0.16 | 0.14 | 0.06 | 0.15 |
| 49 | 0.48 | 0.23 | 0.18 | 0.39 |
| 50 | 2.85 | 1.22 | 1.86 | 2.58 |
| 51 | 0.45 | 0.21 | 0.19 | 0.47 |
| 52 | 0.35 | 0.14 | 0.13 | 0.42 |
| 53 | 2.38 | 0.75 | 1.36 | 2.25 |
| 54 | 7.67 | 4.24 | 7.31 | 7.77 |
| 55 | 3.09 | 1.44 | 2.17 | 2.87 |
| 56 | 0.45 | 0.22 | 0.20 | 0.46 |
| 57 | 0.23 | 0.17 | 0.09 | 0.20 |
| 58 | 0.14 | 0.10 | 0.05 | 0.19 |
| 59 | 0.05 | 0.11 | 0.02 | 0.07 |
| 60 | 0.06 | 0.08 | 0.02 | 0.12 |
| 61 | 0.07 | 0.07 | 0.02 | 0.11 |
| 62 | 0.08 | 0.10 | 0.03 | 0.08 |
| 63 | 0.56 | 0.19 | 0.23 | 0.54 |
| 64 | 0.16 | 0.30 | 0.12 | 0.14 |
| 65 | 0.06 | 0.16 | 0.03 | 0.09 |
| 66 | 0.36 | 0.29 | 0.21 | 0.33 |
| 67 | 0.16 | 0.16 | 0.08 | 0.17 |
| 68 | 0.02 | 0.11 | 0.02 | 0.06 |
| 69 | 0.02 | 0.26 | 0.02 | 0.05 |
| 70 | 0.01 | 0.09 | 0.00 | 0.04 |
| 71 | 0.03 | 0.08 | 0.01 | 0.05 |
| 72 | 0.03 | 0.30 | 0.02 | 0.05 |
| 73 | 0.01 | 0.18 | 0.01 | 0.07 |
| 74 | 0.07 | 0.12 | 0.03 | 0.08 |
| 75 | 0.02 | 0.19 | 0.01 | 0.05 |
| 76 | 0.01 | 0.14 | 0.01 | 0.02 |
| 77 | 0.01 | 0.11 | 0.00 | 0.03 |
| 78 | 0.00 | 0.14 | 0.00 | 0.03 |
| 79 | 0.01 | 0.11 | 0.00 | 0.02 |
| 80 | 0.01 | 0.16 | 0.01 | 0.06 |
| 81 | 0.01 | 0.39 | 0.01 | 0.03 |
| 82 | 0.33 | 3.57 | 0.16 | 0.08 |
| 83 | 0.00 | 0.09 | 0.00 | 0.01 |
| 84 | 0.00 | 0.26 | 0.00 | 0.03 |
| 85 | 0.00 | 0.06 | 0.00 | 0.01 |
| 86 | 0.00 | 0.08 | 0.00 | 0.01 |
| 87 | 0.01 | 0.06 | 0.00 | 0.03 |
| 88 | 0.02 | 0.06 | 0.01 | 0.04 |
| 89 | 0.07 | 0.09 | 0.02 | 0.09 |
| 90 | 0.01 | 0.13 | 0.01 | 0.02 |
| 91 | 0.02 | 0.24 | 0.01 | 0.07 |
| 92 | 0.01 | 0.07 | 0.01 | 0.02 |
| 93 | 0.01 | 0.04 | 0.00 | 0.01 |
| 94 | 0.01 | 0.05 | 0.01 | 0.01 |
| 95 | 0.04 | 1.52 | 0.02 | 0.03 |
| 96 | 0.00 | 0.00 | 0.00 | 0.00 |
| 97 | 0.00 | 0.03 | 0.00 | 0.00 |
| 98 | 0.00 | 0.02 | 0.00 | 0.00 |
| 99 | 0.00 | 0.04 | 0.00 | 0.01 |
| 100 | 1.96 | 15.51 | 0.99 | 0.31 |
| Base | 6.775 | 5.174 | 4.042 | 5.565 |

B

| | mean phyloP score | mean PhastCons core | mean GERP++ score |
|---|---|---|---|
| 1 | 5.173 | 0.926 | 4.191 |
| 2 | 2.308 | 0.738 | 3.030 |
| 3 | 0.617 | 0.484 | 1.211 |
| 4 | 1.357 | 0.549 | 2.194 |
| 5 | 1.173 | 0.488 | 2.371 |
| 6 | 0.636 | 0.180 | 1.130 |
| 7 | 0.292 | 0.126 | 0.339 |
| 8 | 0.132 | 0.155 | 0.074 |
| 9 | -0.029 | 0.065 | -0.186 |
| 10 | -0.243 | 0.042 | -1.062 |
| 11 | 0.112 | 0.051 | -0.076 |
| 12 | 0.197 | 0.062 | 0.155 |
| 13 | 0.272 | 0.076 | 0.350 |
| 14 | 0.221 | 0.070 | 0.235 |
| 15 | -0.249 | 0.040 | -0.921 |
| 16 | -0.287 | 0.018 | -0.829 |
| 17 | 0.401 | 0.066 | 0.541 |
| 18 | -0.430 | 0.016 | -1.440 |
| 19 | -0.215 | 0.073 | -1.025 |
| 20 | 0.166 | 0.070 | 0.052 |
| 21 | 0.111 | 0.055 | -0.107 |
| 22 | 0.057 | 0.038 | -0.175 |
| 23 | 0.130 | 0.045 | -0.043 |
| 24 | 0.095 | 0.037 | -0.116 |
| 25 | 0.085 | 0.035 | -0.128 |
| 26 | 0.118 | 0.043 | -0.054 |
| 27 | -0.409 | 0.019 | -1.299 |
| 28 | 0.148 | 0.180 | -0.240 |
| 29 | 0.071 | 0.032 | -0.147 |
| 30 | 0.111 | 0.036 | -0.060 |
| 31 | 0.101 | 0.040 | -0.075 |
| 32 | 0.175 | 0.053 | 0.056 |
| 33 | 0.104 | 0.051 | -0.121 |
| 34 | 0.142 | 0.056 | -0.073 |
| 35 | 0.233 | 0.059 | 0.135 |
| 36 | 0.228 | 0.051 | 0.159 |
| 37 | 0.276 | 0.071 | 0.317 |
| 38 | 0.214 | 0.053 | 0.181 |
| 39 | 0.122 | 0.044 | -0.028 |
| 40 | 0.132 | 0.041 | -0.011 |
| 41 | 0.129 | 0.039 | -0.013 |
| 42 | 0.113 | 0.038 | -0.021 |
| 43 | 0.145 | 0.046 | -0.012 |
| 44 | 0.185 | 0.058 | 0.054 |
| 45 | 0.122 | 0.039 | -0.049 |
| 46 | 0.135 | 0.043 | -0.017 |
| 47 | -1.126 | 0.004 | -3.166 |
| 48 | -0.509 | 0.015 | -1.702 |
| 49 | -0.475 | 0.020 | -1.812 |
| 50 | -0.310 | 0.071 | -1.652 |
| 51 | -0.469 | 0.018 | -1.851 |
| 52 | -0.530 | 0.012 | -1.917 |
| 53 | -0.484 | 0.045 | -2.083 |
| 54 | -0.338 | 0.219 | -2.198 |
| 55 | -0.376 | 0.082 | -1.613 |
| 56 | -0.593 | 0.018 | -1.908 |
| 57 | -0.619 | 0.017 | -1.721 |
| 58 | -0.641 | 0.011 | -1.857 |
| 59 | -0.653 | 0.015 | -1.489 |
| 60 | -0.674 | 0.010 | -1.794 |
| 61 | -0.631 | 0.010 | -1.793 |
| 62 | -0.600 | 0.012 | -1.754 |
| 63 | -0.618 | 0.015 | -2.166 |
| 64 | 0.078 | 0.035 | -0.144 |
| 65 | -0.022 | 0.030 | -0.347 |
| 66 | -0.061 | 0.030 | -0.548 |
| 67 | -0.088 | 0.024 | -0.531 |
| 68 | -0.025 | 0.028 | -0.273 |
| 69 | 0.171 | 0.048 | 0.033 |
| 70 | -0.611 | 0.016 | -1.079 |
| 71 | -0.661 | 0.013 | -1.275 |
| 72 | 0.183 | 0.050 | 0.046 |
| 73 | -0.071 | 0.035 | -0.184 |
| 74 | 0.040 | 0.042 | -0.142 |
| 75 | 0.038 | 0.043 | -0.043 |
| 76 | 0.124 | 0.106 | 0.102 |
| 77 | 0.068 | 0.068 | 0.035 |
| 78 | 0.093 | 0.089 | 0.057 |
| 79 | 0.117 | 0.137 | 0.098 |
| 80 | 0.030 | 0.044 | -0.047 |
| 81 | 0.074 | 0.093 | 0.029 |
| 82 | 0.064 | 0.210 | -0.180 |
| 83 | 0.064 | 0.108 | 0.057 |
| 84 | 0.019 | 0.056 | -0.045 |
| 85 | 0.072 | 0.122 | 0.065 |
| 86 | 0.037 | 0.145 | 0.054 |
| 87 | -1.178 | 0.034 | -0.463 |
| 88 | -1.067 | 0.024 | -0.695 |
| 89 | -0.883 | 0.012 | -1.473 |
| 90 | 0.022 | 0.100 | -0.003 |
| 91 | -0.064 | 0.047 | -0.168 |
| 92 | 0.004 | 0.146 | -0.003 |
| 93 | 0.006 | 0.165 | 0.004 |
| 94 | 0.000 | 0.188 | -0.007 |
| 95 | 0.030 | 0.182 | -0.026 |
| 96 | 0.000 | 0.120 | 0.000 |
| 97 | -0.006 | 0.196 | -0.001 |
| 98 | 0.001 | 0.169 | 0.002 |
| 99 | -0.003 | 0.140 | -0.002 |
| 100 | 2.005 | 0.795 | 0.170 |

**Figure S16: Conservation states enrichments for evolutionary constrained element calls and average constraint scores. (A)** Each row corresponds to a conservation state and each column corresponds to a different constrained element set. The values correspond to the fold enrichment for bases in a constrained element set for the conservation state. The constrained element sets are from left to right GERP++[13], PhastCons[9], SiPhy-omega, and SiPhy-pi.[7,14] The bottom row gives the percentage of the genome of each constrained element set. **(B)** Each row corresponds to a conservation state and each column corresponds to a different score of constraint. The values correspond to the average constraint score in the conservation state. The constraint scores are from left to right: PhyloP[8], PhastCons[9], and GERP++[13].

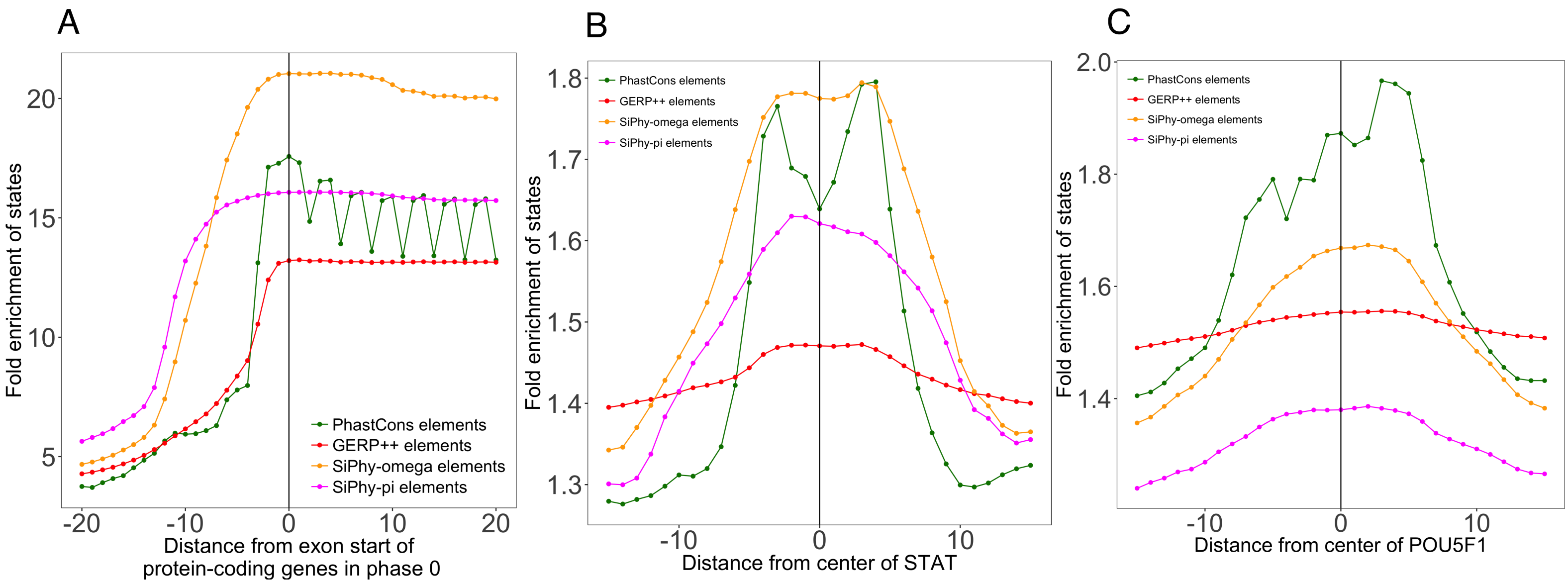


**Figure S17: Positional enrichment of constrained element sets.** Analogous to what was shown for conservation states in **Figure 3**, the graphs show the positional fold enrichments of GERP++, PhastCons, SiPhy-omega and SiPhy-pi constrained elements calls around **(A)** the start of exons of protein coding genes, **(B)** center of instances of a STAT motif, and **(C)** center of instances of a POU5F1 motif. Of these only PhastCons element calls are able to exhibit relevant single nucleotide enrichment variation, as was seen with the conservation states.

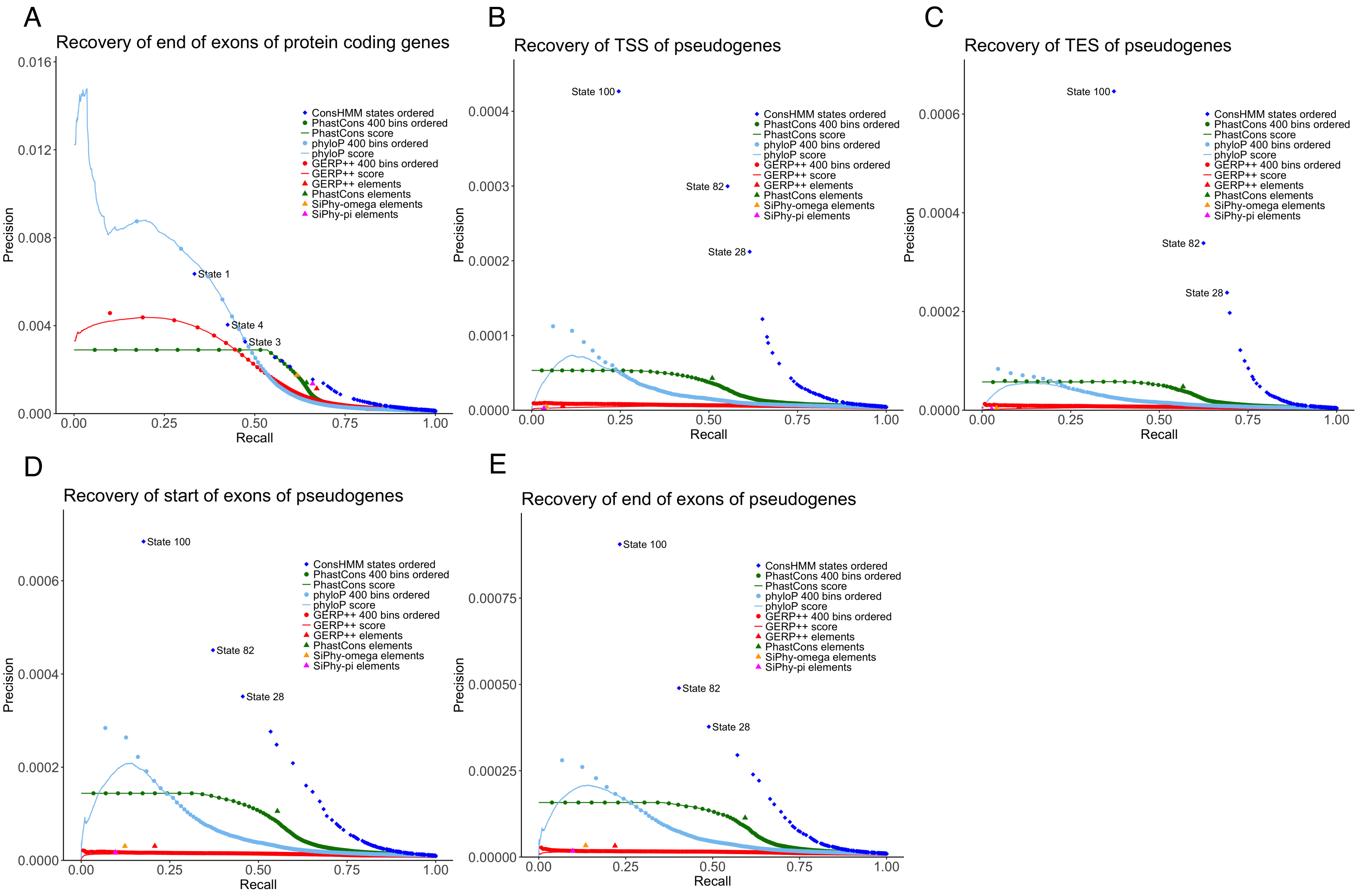


**Figure S18: Precision-recall recovery of conservation states and constrained element and scores for additional gene annotations.** Analogous precision-recall plots to those shown in **Figure 5A-C**, shown here for **(A)** ends of exons of protein coding genes, **(B)** TSS of pseudogenes, **(C)** TES of pseudogenes, **(D)** start of exons of pseudogenes, and **(E)** end of exons of pseudogenes. Precision-recall values were computed using the same procedure as for **Figure 5** (**Methods**). The first few conservation states added are labeled.

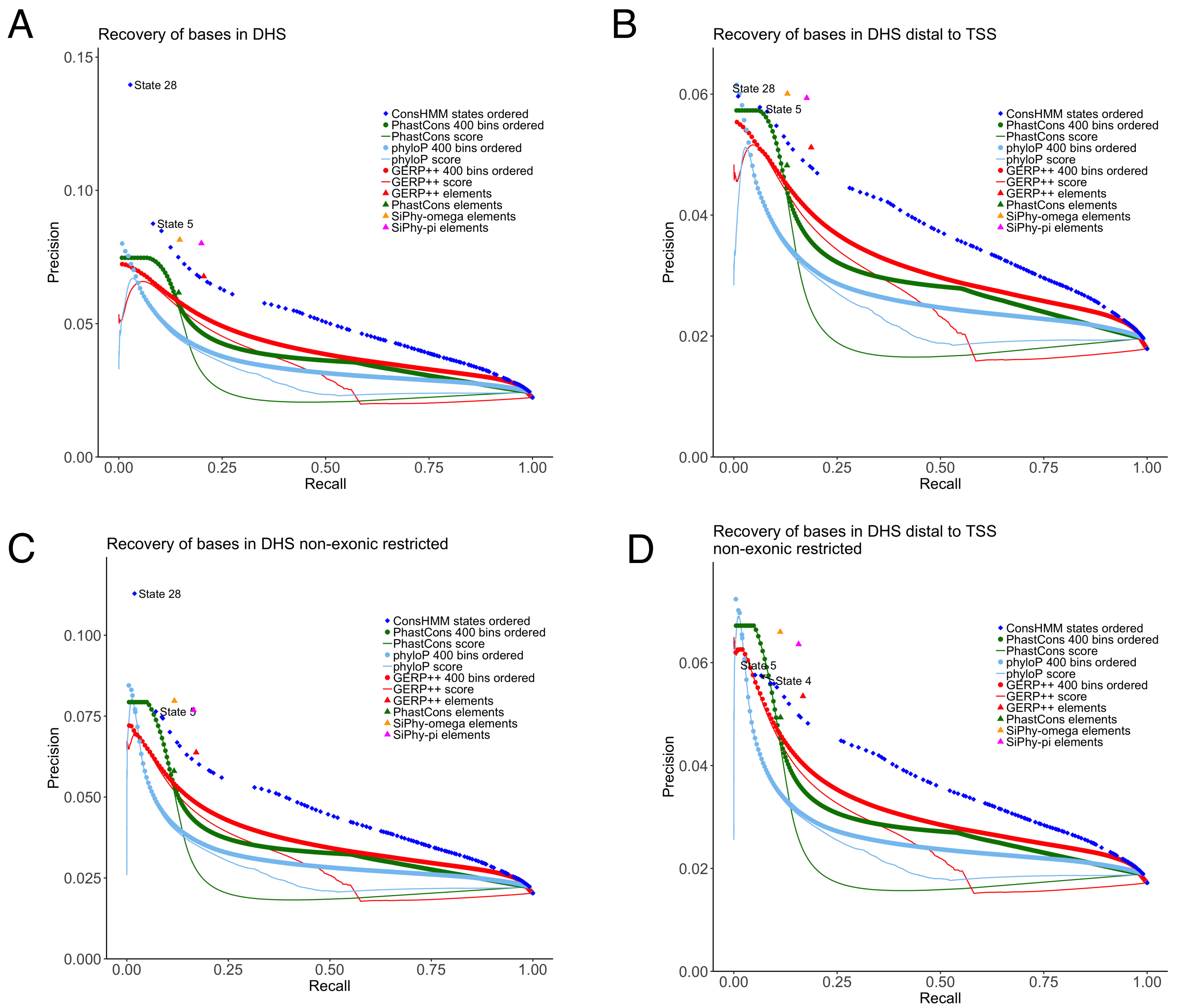


**Figure S19: Precision-recall recovery of conservation states and constrained element sets and scores for a concatenation of DHS bases in 53 cell and tissue types.** Analogous precision-recall plots to those shown in **Figures 5A-C** and **S18** shown here for DHS bases concatenated across experiments shown **(A)** without restriction and **(B-D)** with the following restrictions for the target and background **(B)** bases more than 2kb away from a TSS, **(C)** non-exonic bases, **(D)** non-exonic bases more than 2kb away from a TSS.

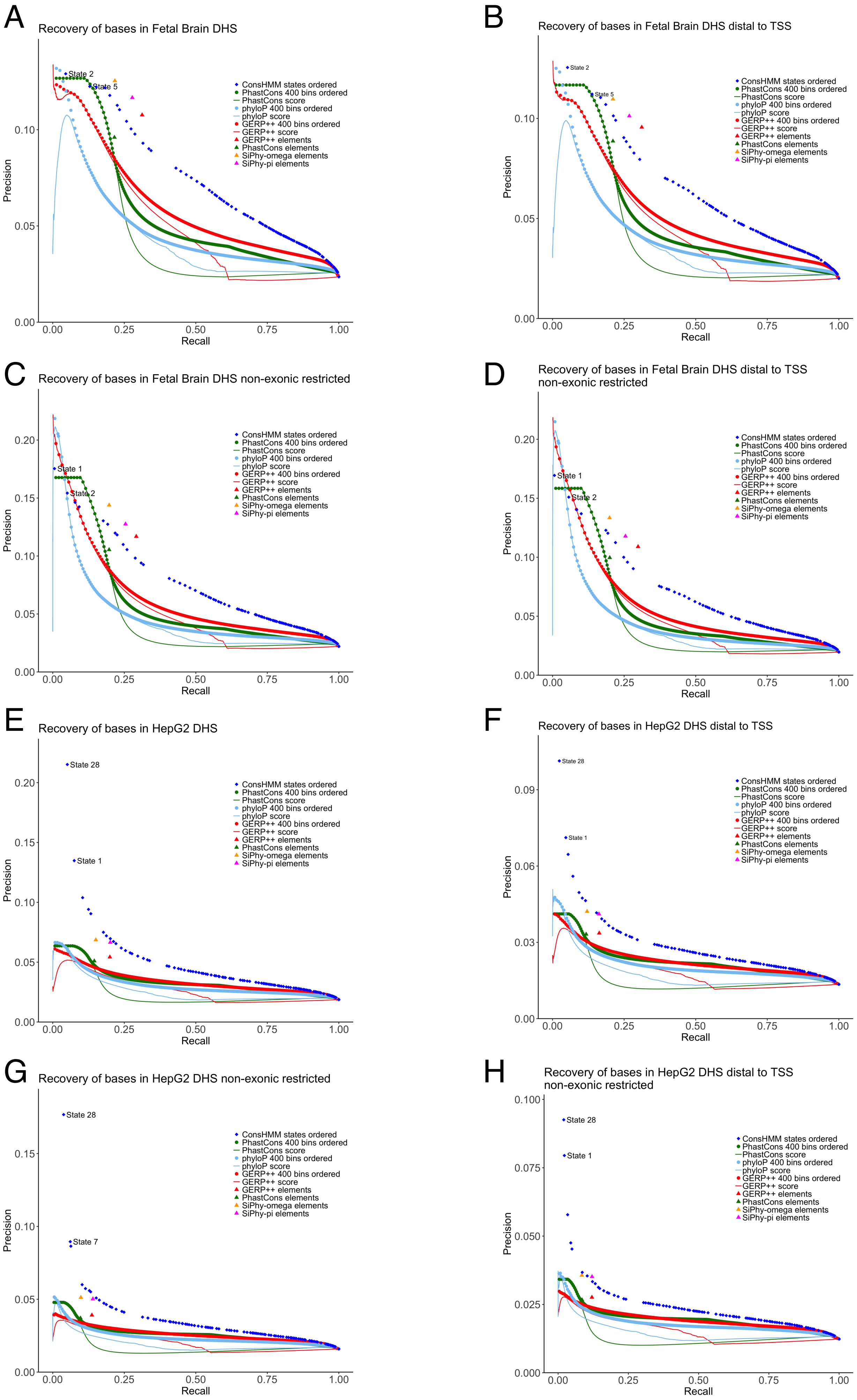


**Figure S20: Precision-recall recovery of conservation states and constrained element and scores for DHS in two cell types.** Analogous precision-recall plots to those shown in **Figures 5A-C, S18** and **S19** shown here for **(A)** Fetal Brain DHS, **(B)** Fetal Brain DHS when restricting target and background to bases more than 2kb away from a TSS, **(C)** Fetal Brain DHS when restricting target and background to non-exonic bases of the genome, **(D)** Fetal Brain DHS when restricting target and background to non-exonic bases of the genome that are more than 2kb away from a TSS, **(E)** HepG2 DHS, **(F)** HepG2 DHS when restricting target and background to bases more than 2kb away from a TSS, **(G)** HepG2 when restricting target and background to non-exonic bases of the genome, **(H)** HepG2 DHS when restricting target and background to non-exonic bases of the genome that are more than 2kb away from a TSS.

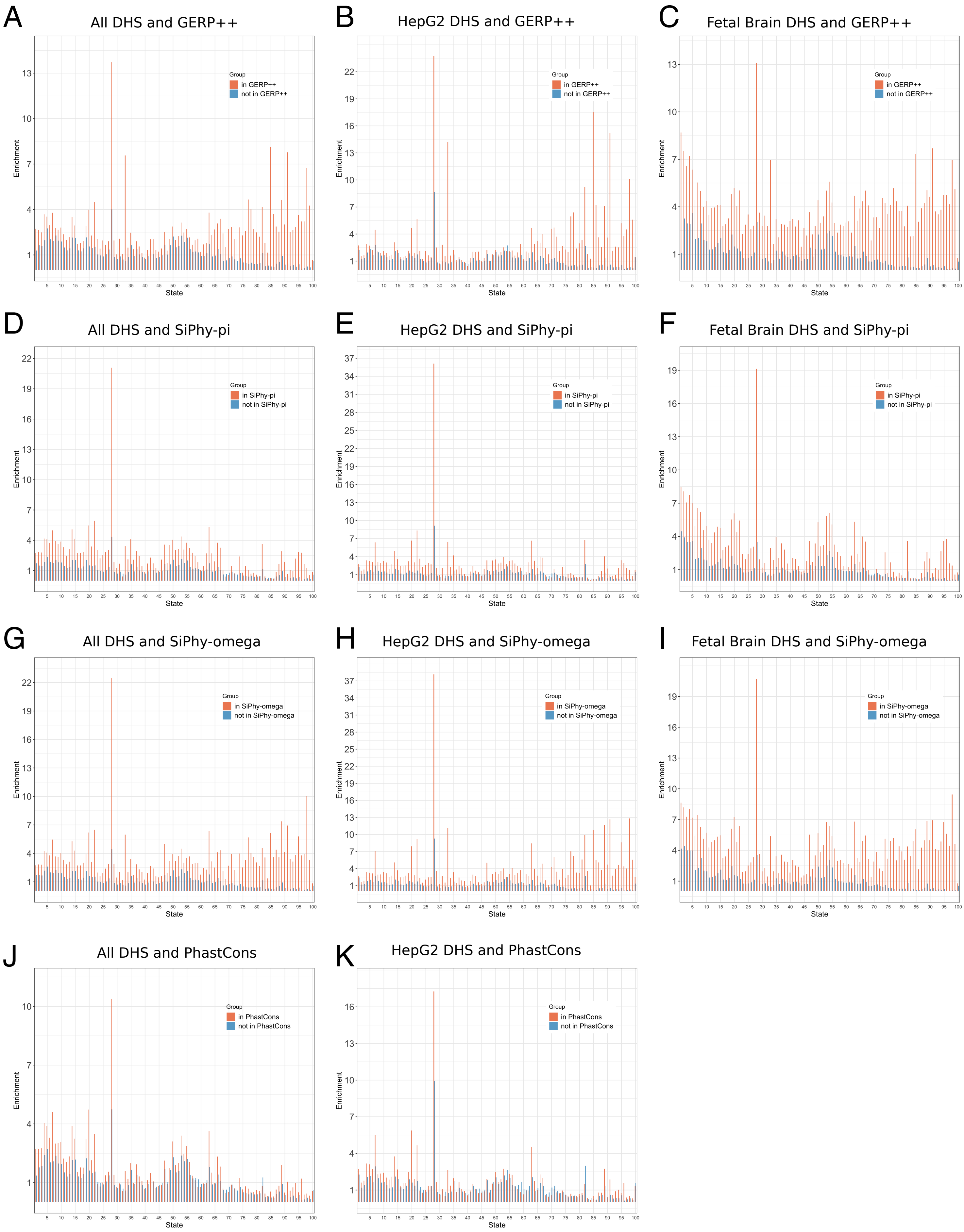


**Figure S21: Enrichment for non-exonic DHS conditioned on conservation state and constrained element sets.** Analogous graphs to **Figure 5D**, showing enrichments for bases in DHS in the non-exonic portion of each conservation state conditioned on whether it is in a constrained element or not. Enrichments are shown here for **(A-C)** bases in and out of GERP++ elements for **(A)** geometric mean over 53 cell and tissue types, **(B)** HepG2 DHS, **(C)** Fetal Brain DHS; **(D-F)** bases in and out of SiPhy-pi elements for **(D)** geometric mean over 53 cell and tissue types **(E)** HepG2 DHS, **(F)** Fetal Brain DHS; **(G-I)** bases in and out of SiPhy-omega elements for **(G)** geomtric mean over 53 cell and tissue types **(H)** HepG2 DHS, **(I)** Fetal Brain DHS; **(J-K)** bases in and out of PhastCons elements for **(J)** geometric mean over 53 cell and tissue types **(K)** HepG2 DHS.

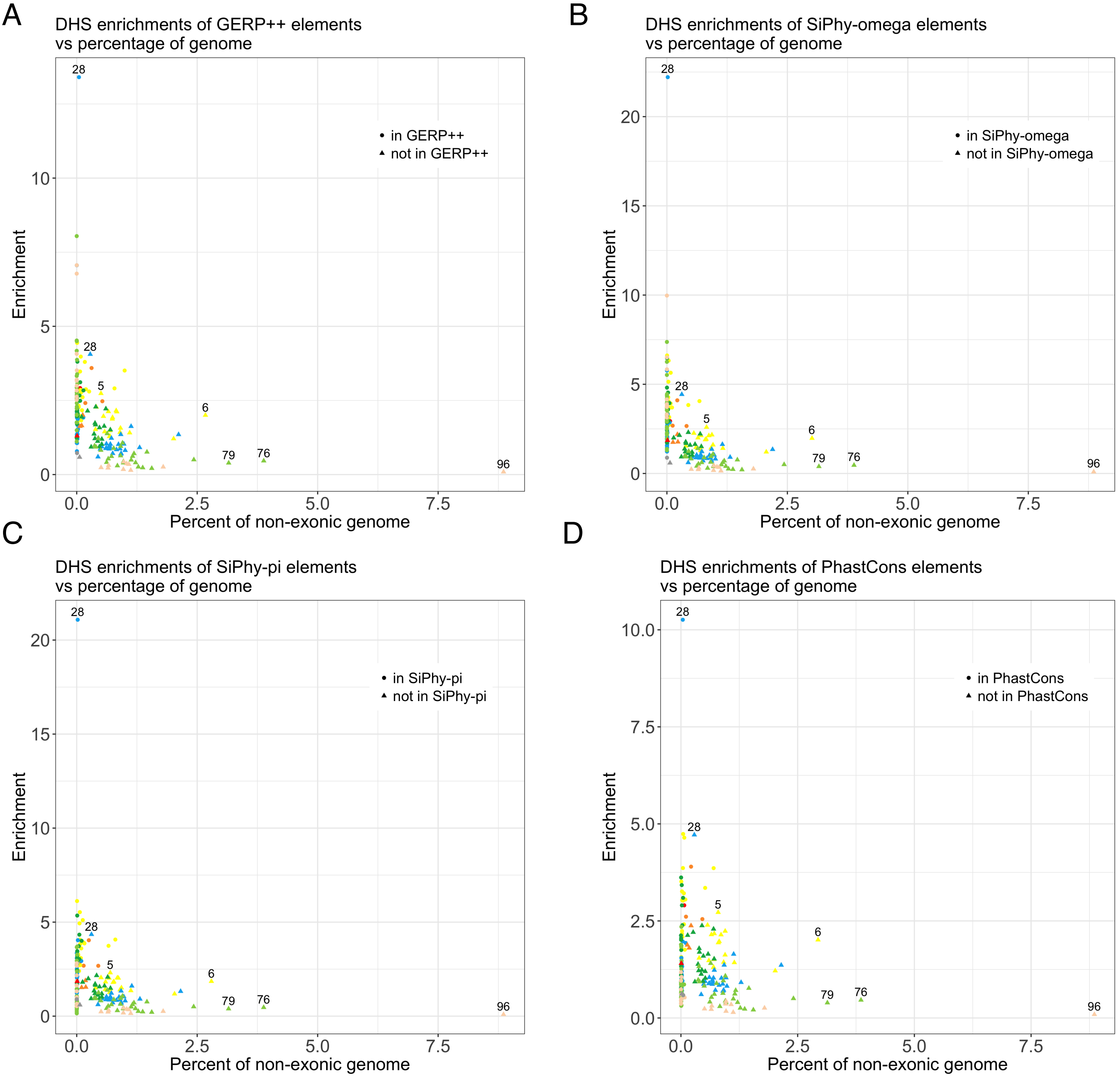


**Figure S22: Enrichment for non-exonic DHS conditioned on conservation state and constrained element sets versus percent of non-exonic genome covered.** The enrichments in **Figure S21A,D,G,J** are shown here on the y-axis (geometric mean over 53 cell and tissue types), with the x-axis corresponding to the median percentage of the non-exonic genome covered by the bases falling in each category across 53 cell and tissue types.The coloring of a point corresponds to the coloring of states in **Figure 2**. The shape of a point corresponds to whether it is inside or outside a constrained element set according to the legend shown. Labeled points either have an enrichment >10 fold, cover >2.5% of the non-exonic genome, or are subsets of states not in a constrained element set with an enrichment >2 fold. The figure shows substantial variation of enrichments for bases both inside and outside of constrained elements depending on the conservation state, including for subsets covering non-negligible portions of the non-exonic genome. The constrained element sets used for each panel are **(A)** GERP++ elements, **(B)** SiPhy-pi elements, **(C)** SiPhy-omega elements, and **(D)** PhastCons elements.

A

| | All CNEEs | Euteleostomi | Tetrapod | Amniote | Mammal | Theria | Eutheria | Boreoeutheria | Euarchontoglire | Hominini |
|---|---|---|---|---|---|---|---|---|---|---|
| 1 | 2.43 | 65.12 | 5.69 | 0.63 | 0.02 | 0.02 | 0.01 | 0.26 | 0.00 | 0.00 |
| 2 | 16.41 | 42.84 | 37.42 | 53.14 | 1.00 | 0.46 | 0.06 | 0.91 | 0.15 | 0.04 |
| 3 | 6.92 | 17.39 | 11.74 | 24.50 | 0.64 | 0.30 | 0.04 | 0.55 | 0.11 | 0.01 |
| 4 | 10.74 | 6.03 | 5.59 | 20.54 | 21.60 | 8.33 | 1.21 | 1.73 | 0.01 | 0.12 |
| 5 | 12.88 | 1.39 | 1.08 | 4.16 | 30.64 | 32.56 | 3.49 | 0.87 | 0.10 | 0.05 |
| 6 | 3.80 | 0.20 | 0.11 | 0.37 | 2.06 | 2.39 | 10.52 | 0.97 | 0.06 | 0.08 |
| 7 | 2.30 | 1.83 | 1.10 | 6.30 | 2.17 | 1.54 | 0.63 | 0.92 | 0.13 | 0.02 |
| 8 | 4.04 | 0.56 | 0.39 | 1.34 | 9.74 | 10.29 | 0.90 | 0.35 | 0.02 | 0.05 |
| 9 | 1.43 | 0.10 | 0.05 | 0.15 | 0.84 | 1.09 | 3.80 | 0.33 | 0.01 | 0.04 |
| 10 | 0.87 | 0.09 | 0.05 | 0.14 | 0.95 | 1.33 | 1.59 | 0.29 | 0.04 | 0.02 |
| 11 | 0.66 | 0.09 | 0.05 | 0.10 | 0.58 | 0.77 | 1.36 | 0.79 | 0.06 | 0.03 |
| 12 | 0.95 | 0.07 | 0.04 | 0.09 | 0.67 | 1.11 | 2.12 | 1.24 | 0.04 | 0.06 |
| 13 | 1.21 | 0.08 | 0.04 | 0.11 | 0.68 | 1.18 | 3.09 | 0.44 | 0.05 | 0.05 |
| 14 | 1.03 | 0.11 | 0.06 | 0.11 | 0.76 | 1.17 | 2.39 | 0.50 | 0.04 | 0.06 |
| 15 | 0.83 | 0.07 | 0.03 | 0.09 | 0.49 | 0.67 | 2.19 | 0.23 | 0.02 | 0.02 |
| 16 | 0.20 | 0.02 | 0.01 | 0.02 | 0.05 | 0.09 | 0.55 | 0.71 | 0.01 | 0.02 |
| 17 | 0.70 | 0.05 | 0.04 | 0.06 | 0.18 | 0.34 | 1.90 | 2.40 | 0.05 | 0.07 |
| 18 | 0.20 | 0.02 | 0.01 | 0.02 | 0.07 | 0.12 | 0.54 | 0.45 | 0.04 | 0.02 |
| 19 | 1.76 | 0.17 | 0.11 | 0.37 | 2.64 | 3.08 | 2.45 | 0.28 | 0.04 | 0.02 |
| 20 | 1.05 | 0.09 | 0.08 | 0.18 | 1.13 | 1.43 | 1.97 | 0.59 | 0.06 | 0.03 |
| 21 | 0.80 | 0.08 | 0.06 | 0.15 | 0.90 | 1.08 | 1.48 | 0.50 | 0.07 | 0.05 |
| 22 | 0.33 | 0.04 | 0.04 | 0.04 | 0.16 | 0.28 | 0.81 | 0.65 | 0.17 | 0.09 |
| 23 | 0.28 | 0.05 | 0.05 | 0.03 | 0.05 | 0.06 | 0.66 | 2.37 | 0.01 | 0.28 |
| 24 | 0.23 | 0.01 | 0.04 | 0.02 | 0.04 | 0.05 | 0.56 | 1.63 | 0.11 | 0.11 |
| 25 | 0.24 | 0.02 | 0.02 | 0.02 | 0.02 | 0.03 | 0.48 | 3.10 | 0.07 | 0.10 |
| 26 | 0.39 | 0.03 | 0.02 | 0.04 | 0.15 | 0.20 | 1.07 | 0.69 | 0.01 | 0.07 |
| 27 | 0.20 | 0.03 | 0.02 | 0.02 | 0.09 | 0.13 | 0.53 | 0.48 | 0.02 | 0.03 |
| 28 | 1.65 | 1.82 | 3.39 | 2.39 | 1.27 | 1.22 | 0.46 | 5.55 | 5.01 | 0.28 |
| 29 | 0.22 | 0.01 | 0.01 | 0.02 | 0.04 | 0.06 | 0.57 | 1.40 | 0.42 | 0.09 |
| 30 | 0.23 | 0.03 | 0.05 | 0.02 | 0.05 | 0.07 | 0.57 | 1.44 | 0.05 | 0.08 |
| 31 | 0.24 | 0.04 | 0.05 | 0.02 | 0.06 | 0.07 | 0.62 | 1.17 | 0.02 | 0.20 |
| 32 | 0.25 | 0.02 | 0.11 | 0.06 | 0.01 | 0.02 | 0.22 | 5.49 | 0.11 | 0.40 |
| 33 | 0.30 | 0.04 | 0.20 | 0.04 | 0.07 | 0.07 | 0.43 | 4.02 | 1.21 | 0.32 |
| 34 | 0.61 | 0.06 | 0.11 | 0.06 | 0.25 | 0.44 | 1.40 | 2.43 | 1.87 | 0.22 |
| 35 | 0.57 | 0.08 | 0.05 | 0.07 | 0.33 | 0.41 | 1.44 | 0.80 | 0.03 | 0.13 |
| 36 | 0.37 | 0.06 | 0.06 | 0.03 | 0.04 | 0.07 | 0.85 | 3.62 | 0.04 | 0.17 |
| 37 | 0.92 | 0.09 | 0.05 | 0.08 | 0.42 | 0.68 | 2.44 | 1.14 | 0.05 | 0.08 |
| 38 | 0.51 | 0.07 | 0.03 | 0.05 | 0.23 | 0.33 | 1.38 | 0.59 | 0.04 | 0.08 |
| 39 | 0.41 | 0.03 | 0.06 | 0.04 | 0.17 | 0.24 | 0.95 | 1.98 | 0.03 | 0.10 |
| 40 | 0.25 | 0.03 | 0.03 | 0.02 | 0.02 | 0.03 | 0.43 | 3.85 | 0.06 | 0.15 |
| 41 | 0.31 | 0.04 | 0.02 | 0.02 | 0.06 | 0.10 | 0.69 | 3.01 | 0.09 | 0.08 |
| 42 | 0.29 | 0.04 | 0.02 | 0.02 | 0.05 | 0.07 | 0.75 | 1.73 | 0.09 | 0.34 |
| 43 | 0.30 | 0.04 | 0.03 | 0.03 | 0.05 | 0.06 | 0.79 | 1.66 | 0.02 | 0.15 |
| 44 | 0.33 | 0.05 | 0.14 | 0.04 | 0.03 | 0.04 | 0.42 | 6.27 | 0.09 | 0.38 |
| 45 | 0.24 | 0.03 | 0.04 | 0.02 | 0.03 | 0.05 | 0.55 | 2.22 | 0.18 | 0.20 |
| 46 | 0.31 | 0.04 | 0.04 | 0.03 | 0.05 | 0.08 | 0.78 | 2.10 | 0.05 | 0.17 |
| 47 | 0.07 | 0.01 | 0.01 | 0.01 | 0.04 | 0.05 | 0.16 | 0.15 | 0.01 | 0.02 |
| 48 | 0.12 | 0.02 | 0.03 | 0.01 | 0.03 | 0.05 | 0.28 | 0.65 | 0.02 | 0.04 |
| 49 | 0.22 | 0.03 | 0.02 | 0.02 | 0.12 | 0.17 | 0.56 | 0.30 | 0.02 | 0.03 |
| 50 | 1.60 | 0.18 | 0.12 | 0.49 | 2.57 | 2.73 | 2.04 | 0.27 | 0.01 | 0.04 |
| 51 | 0.21 | 0.02 | 0.01 | 0.02 | 0.08 | 0.14 | 0.55 | 0.43 | 0.01 | 0.04 |
| 52 | 0.14 | 0.01 | 0.01 | 0.01 | 0.05 | 0.09 | 0.36 | 0.33 | 0.06 | 0.02 |
| 53 | 1.01 | 0.16 | 0.10 | 0.38 | 1.58 | 1.65 | 1.27 | 0.20 | 0.03 | 0.02 |
| 54 | 3.08 | 8.06 | 5.55 | 9.78 | 0.87 | 0.37 | 0.07 | 0.93 | 0.14 | 0.03 |
| 55 | 2.01 | 0.23 | 0.20 | 0.79 | 3.35 | 3.34 | 2.36 | 0.53 | 0.02 | 0.02 |
| 56 | 0.23 | 0.02 | 0.02 | 0.02 | 0.11 | 0.18 | 0.54 | 0.70 | 0.17 | 0.03 |
| 57 | 0.15 | 0.02 | 0.03 | 0.02 | 0.06 | 0.08 | 0.36 | 0.77 | 0.01 | 0.03 |
| 58 | 0.09 | 0.02 | 0.02 | 0.01 | 0.03 | 0.05 | 0.20 | 0.49 | 0.06 | 0.04 |
| 59 | 0.08 | 0.02 | 0.05 | 0.01 | 0.02 | 0.01 | 0.14 | 1.08 | 0.03 | 0.10 |
| 60 | 0.06 | 0.01 | 0.01 | 0.01 | 0.01 | 0.01 | 0.11 | 0.86 | 0.02 | 0.03 |
| 61 | 0.06 | 0.01 | 0.02 | 0.01 | 0.01 | 0.02 | 0.14 | 0.34 | 0.05 | 0.02 |
| 62 | 0.08 | 0.01 | 0.02 | 0.01 | 0.02 | 0.02 | 0.19 | 0.53 | 0.03 | 0.03 |
| 63 | 0.20 | 0.03 | 0.02 | 0.04 | 0.20 | 0.23 | 0.38 | 0.25 | 0.10 | 0.01 |
| 64 | 0.25 | 0.01 | 0.03 | 0.03 | 0.10 | 0.12 | 0.59 | 1.38 | 0.66 | 0.18 |
| 65 | 0.13 | 0.02 | 0.06 | 0.03 | 0.05 | 0.04 | 0.32 | 0.18 | 1.97 | 0.61 |
| 66 | 0.27 | 0.06 | 0.04 | 0.08 | 0.23 | 0.26 | 0.57 | 0.12 | 1.72 | 0.08 |
| 67 | 0.14 | 0.06 | 0.07 | 0.05 | 0.06 | 0.07 | 0.17 | 0.48 | 13.81 | 0.41 |
| 68 | 0.11 | 0.03 | 0.16 | 0.05 | 0.01 | 0.01 | 0.08 | 0.54 | 13.31 | 0.60 |
| 69 | 0.19 | 0.04 | 0.21 | 0.06 | 0.01 | 0.01 | 0.19 | 3.40 | 0.76 | 0.59 |
| 70 | 0.08 | 0.02 | 0.15 | 0.03 | 0.01 | 0.01 | 0.06 | 1.18 | 0.19 | 0.25 |
| 71 | 0.07 | 0.01 | 0.09 | 0.02 | 0.02 | 0.02 | 0.09 | 0.81 | 0.34 | 0.21 |
| 72 | 0.23 | 0.02 | 0.16 | 0.05 | 0.02 | 0.03 | 0.26 | 3.96 | 0.99 | 0.60 |
| 73 | 0.15 | 0.06 | 0.22 | 0.05 | 0.01 | 0.01 | 0.10 | 2.71 | 0.88 | 0.97 |
| 74 | 0.15 | 0.05 | 0.25 | 0.21 | 0.04 | 0.04 | 0.08 | 0.33 | 2.51 | 7.05 |
| 75 | 0.16 | 0.16 | 0.48 | 0.12 | 0.03 | 0.02 | 0.09 | 0.69 | 2.63 | 3.11 |
| 76 | 0.14 | 0.02 | 0.41 | 0.25 | 0.02 | 0.02 | 0.04 | 0.17 | 0.45 | 5.17 |
| 77 | 0.16 | 0.04 | 0.81 | 0.13 | 0.01 | 0.01 | 0.03 | 0.30 | 1.30 | 3.44 |
| 78 | 0.17 | 0.02 | 0.53 | 0.31 | 0.01 | 0.00 | 0.04 | 0.48 | 1.42 | 3.82 |
| 79 | 0.17 | 0.01 | 0.47 | 0.40 | 0.01 | 0.01 | 0.03 | 0.25 | 0.42 | 3.40 |
| 80 | 0.14 | 0.06 | 0.47 | 0.12 | 0.02 | 0.01 | 0.06 | 0.61 | 5.97 | 1.83 |
| 81 | 0.26 | 0.08 | 0.97 | 0.28 | 0.05 | 0.01 | 0.08 | 1.61 | 0.92 | 2.10 |
| 82 | 3.31 | 0.94 | 22.57 | 0.63 | 0.54 | 0.07 | 1.05 | 3.13 | 11.88 | 4.50 |
| 83 | 0.20 | 0.01 | 0.79 | 0.41 | 0.00 | 0.00 | 0.01 | 0.14 | 0.28 | 4.12 |
| 84 | 0.25 | 0.05 | 1.07 | 0.21 | 0.02 | 0.01 | 0.05 | 1.59 | 4.02 | 2.60 |
| 85 | 0.19 | 0.01 | 0.93 | 0.28 | 0.00 | 0.00 | 0.01 | 0.25 | 0.68 | 2.10 |
| 86 | 0.16 | 0.01 | 0.37 | 0.50 | 0.00 | 0.00 | 0.01 | 0.14 | 0.23 | 2.04 |
| 87 | 0.12 | 0.02 | 0.61 | 0.16 | 0.01 | 0.01 | 0.02 | 0.15 | 0.51 | 0.69 |
| 88 | 0.10 | 0.05 | 0.42 | 0.11 | 0.02 | 0.01 | 0.03 | 0.12 | 0.49 | 1.16 |
| 89 | 0.07 | 0.02 | 0.09 | 0.02 | 0.03 | 0.03 | 0.09 | 0.68 | 0.85 | 0.11 |
| 90 | 0.22 | 0.06 | 0.66 | 0.49 | 0.02 | 0.01 | 0.03 | 0.36 | 1.14 | 2.55 |
| 91 | 0.21 | 0.06 | 0.56 | 0.13 | 0.03 | 0.03 | 0.10 | 2.18 | 4.96 | 0.90 |
| 92 | 0.15 | 0.01 | 0.42 | 0.40 | 0.02 | 0.01 | 0.03 | 0.22 | 0.59 | 1.16 |
| 93 | 0.18 | 0.01 | 0.46 | 0.51 | 0.01 | 0.01 | 0.02 | 0.17 | 0.48 | 1.07 |
| 94 | 0.15 | 0.01 | 0.45 | 0.40 | 0.02 | 0.02 | 0.03 | 0.11 | 0.26 | 0.65 |
| 95 | 0.78 | 0.12 | 5.00 | 0.43 | 0.17 | 0.04 | 0.13 | 0.58 | 4.27 | 1.15 |
| 96 | 0.02 | 0.00 | 0.09 | 0.02 | 0.00 | 0.00 | 0.00 | 0.01 | 0.05 | 0.02 |
| 97 | 0.11 | 0.00 | 0.48 | 0.22 | 0.00 | 0.00 | 0.01 | 0.07 | 0.09 | 0.35 |
| 98 | 0.13 | 0.01 | 0.46 | 0.35 | 0.01 | 0.00 | 0.00 | 0.14 | 0.10 | 0.37 |
| 99 | 0.16 | 0.01 | 0.64 | 0.32 | 0.00 | 0.00 | 0.01 | 0.22 | 0.73 | 0.92 |
| 100 | 6.87 | 3.73 | 50.95 | 0.41 | 0.20 | 0.07 | 1.69 | 5.16 | 28.81 | 11.10 |
| Base | 2.70 | 0.07 | 0.31 | 0.54 | 0.52 | 0.42 | 0.75 | 0.08 | 0.01 | 0.01 |

Enrichments of Lowe et al.[22] CNEEs

B

| | All CNEEs | Euteleostomi | Tetrapod | Amniote | Mammal | Theria | Eutheria | Boreoeutheria | Euarchontoglire | Hominini |
|---|---|---|---|---|---|---|---|---|---|---|
| 1 | 3.20 | 70.28 | 8.59 | 0.76 | 0.02 | 0.02 | 0.01 | 0.55 | 0.01 | 0.00 |
| 2 | 20.88 | 45.36 | 56.45 | 62.31 | 1.08 | 0.51 | 0.08 | 1.87 | 0.36 | 0.15 |
| 3 | 7.98 | 16.85 | 15.96 | 26.21 | 0.58 | 0.28 | 0.04 | 0.98 | 0.25 | 0.02 |
| 4 | 12.65 | 5.43 | 7.68 | 22.61 | 22.67 | 8.90 | 1.55 | 3.36 | 0.03 | 0.39 |
| 5 | 15.37 | 1.26 | 1.47 | 4.61 | 32.90 | 35.38 | 4.53 | 1.60 | 0.16 | 0.16 |
| 6 | 4.09 | 0.14 | 0.12 | 0.37 | 2.04 | 2.30 | 12.55 | 1.54 | 0.09 | 0.23 |
| 7 | 2.10 | 0.96 | 1.00 | 5.63 | 1.75 | 1.23 | 0.57 | 1.33 | 0.22 | 0.03 |
| 8 | 3.92 | 0.36 | 0.41 | 1.20 | 8.60 | 9.06 | 0.91 | 0.49 | 0.01 | 0.10 |
| 9 | 1.24 | 0.05 | 0.04 | 0.12 | 0.70 | 0.89 | 3.62 | 0.39 | 0.01 | 0.08 |
| 10 | 0.73 | 0.04 | 0.04 | 0.12 | 0.77 | 1.08 | 1.40 | 0.29 | 0.02 | 0.04 |
| 11 | 0.53 | 0.04 | 0.04 | 0.07 | 0.50 | 0.63 | 1.14 | 0.86 | 0.09 | 0.05 |
| 12 | 0.84 | 0.03 | 0.04 | 0.08 | 0.58 | 0.98 | 1.98 | 1.71 | 0.04 | 0.17 |
| 13 | 1.11 | 0.04 | 0.04 | 0.10 | 0.60 | 1.05 | 3.09 | 0.56 | 0.03 | 0.10 |
| 14 | 0.89 | 0.06 | 0.05 | 0.09 | 0.65 | 1.01 | 2.18 | 0.57 | 0.03 | 0.17 |
| 15 | 0.67 | 0.03 | 0.02 | 0.07 | 0.38 | 0.49 | 1.95 | 0.24 | 0.01 | 0.05 |
| 16 | 0.11 | 0.01 | 0.01 | 0.01 | 0.03 | 0.05 | 0.33 | 0.56 | 0.01 | 0.03 |
| 17 | 0.57 | 0.02 | 0.03 | 0.05 | 0.15 | 0.29 | 1.69 | 3.04 | 0.04 | 0.15 |
| 18 | 0.13 | 0.01 | 0.00 | 0.01 | 0.05 | 0.08 | 0.37 | 0.39 | 0.02 | 0.03 |
| 19 | 1.62 | 0.10 | 0.11 | 0.32 | 2.29 | 2.67 | 2.35 | 0.33 | 0.05 | 0.03 |
| 20 | 0.96 | 0.05 | 0.08 | 0.16 | 1.03 | 1.30 | 1.84 | 0.80 | 0.07 | 0.08 |
| 21 | 0.70 | 0.04 | 0.05 | 0.13 | 0.79 | 0.94 | 1.30 | 0.59 | 0.08 | 0.09 |
| 22 | 0.22 | 0.01 | 0.02 | 0.03 | 0.12 | 0.22 | 0.55 | 0.58 | 0.20 | 0.22 |
| 23 | 0.15 | 0.01 | 0.01 | 0.01 | 0.03 | 0.04 | 0.39 | 1.90 | 0.00 | 0.56 |
| 24 | 0.11 | 0.01 | 0.01 | 0.01 | 0.02 | 0.03 | 0.32 | 1.26 | 0.08 | 0.18 |
| 25 | 0.14 | 0.01 | 0.01 | 0.01 | 0.01 | 0.02 | 0.31 | 3.03 | 0.09 | 0.18 |
| 26 | 0.26 | 0.01 | 0.01 | 0.03 | 0.12 | 0.16 | 0.75 | 0.67 | 0.01 | 0.14 |
| 27 | 0.12 | 0.01 | 0.01 | 0.01 | 0.06 | 0.08 | 0.32 | 0.37 | 0.01 | 0.06 |
| 28 | 1.56 | 1.25 | 3.57 | 2.21 | 1.06 | 1.07 | 0.46 | 8.95 | 9.58 | 0.30 |
| 29 | 0.13 | 0.01 | 0.01 | 0.01 | 0.02 | 0.04 | 0.38 | 1.28 | 0.61 | 0.19 |
| 30 | 0.12 | 0.00 | 0.01 | 0.01 | 0.04 | 0.05 | 0.33 | 1.15 | 0.05 | 0.13 |
| 31 | 0.12 | 0.00 | 0.01 | 0.01 | 0.04 | 0.04 | 0.34 | 0.90 | 0.01 | 0.19 |
| 32 | 0.12 | 0.00 | 0.02 | 0.01 | 0.01 | 0.01 | 0.13 | 4.60 | 0.14 | 0.26 |
| 33 | 0.16 | 0.01 | 0.05 | 0.02 | 0.05 | 0.07 | 0.26 | 3.59 | 1.45 | 0.62 |
| 34 | 0.47 | 0.03 | 0.06 | 0.05 | 0.21 | 0.38 | 1.15 | 3.17 | 3.59 | 0.60 |
| 35 | 0.41 | 0.04 | 0.04 | 0.05 | 0.27 | 0.33 | 1.07 | 0.74 | 0.02 | 0.19 |
| 36 | 0.23 | 0.02 | 0.02 | 0.02 | 0.02 | 0.05 | 0.59 | 3.60 | 0.03 | 0.36 |
| 37 | 0.77 | 0.05 | 0.03 | 0.06 | 0.35 | 0.57 | 2.20 | 1.50 | 0.04 | 0.19 |
| 38 | 0.36 | 0.02 | 0.02 | 0.04 | 0.18 | 0.27 | 1.03 | 0.55 | 0.04 | 0.12 |
| 39 | 0.27 | 0.01 | 0.03 | 0.03 | 0.13 | 0.18 | 0.64 | 2.41 | 0.02 | 0.19 |
| 40 | 0.13 | 0.01 | 0.01 | 0.01 | 0.01 | 0.02 | 0.27 | 3.31 | 0.07 | 0.32 |
| 41 | 0.21 | 0.01 | 0.01 | 0.01 | 0.04 | 0.07 | 0.52 | 3.22 | 0.03 | 0.21 |
| 42 | 0.16 | 0.01 | 0.01 | 0.01 | 0.03 | 0.05 | 0.48 | 1.51 | 0.08 | 0.88 |
| 43 | 0.17 | 0.01 | 0.01 | 0.02 | 0.03 | 0.04 | 0.50 | 1.45 | 0.00 | 0.22 |
| 44 | 0.17 | 0.01 | 0.04 | 0.01 | 0.02 | 0.03 | 0.25 | 5.38 | 0.05 | 0.55 |
| 45 | 0.12 | 0.01 | 0.01 | 0.01 | 0.02 | 0.03 | 0.31 | 1.74 | 0.13 | 0.30 |
| 46 | 0.17 | 0.01 | 0.02 | 0.01 | 0.03 | 0.05 | 0.46 | 1.81 | 0.05 | 0.29 |
| 47 | 0.03 | 0.00 | 0.00 | 0.00 | 0.02 | 0.03 | 0.08 | 0.08 | 0.00 | 0.02 |
| 48 | 0.06 | 0.00 | 0.01 | 0.00 | 0.02 | 0.03 | 0.16 | 0.53 | 0.00 | 0.07 |
| 49 | 0.14 | 0.01 | 0.01 | 0.01 | 0.08 | 0.11 | 0.39 | 0.28 | 0.01 | 0.03 |
| 50 | 1.54 | 0.11 | 0.13 | 0.45 | 2.31 | 2.46 | 2.03 | 0.34 | 0.01 | 0.07 |
| 51 | 0.15 | 0.01 | 0.01 | 0.01 | 0.06 | 0.11 | 0.42 | 0.42 | 0.00 | 0.07 |
| 52 | 0.08 | 0.00 | 0.00 | 0.01 | 0.03 | 0.06 | 0.24 | 0.28 | 0.03 | 0.04 |
| 53 | 0.88 | 0.09 | 0.10 | 0.32 | 1.31 | 1.36 | 1.15 | 0.19 | 0.02 | 0.03 |
| 54 | 3.30 | 7.22 | 7.14 | 9.80 | 0.70 | 0.29 | 0.06 | 1.48 | 0.31 | 0.08 |
| 55 | 2.02 | 0.16 | 0.22 | 0.77 | 3.12 | 3.13 | 2.46 | 0.80 | 0.02 | 0.04 |
| 56 | 0.17 | 0.01 | 0.01 | 0.01 | 0.09 | 0.14 | 0.43 | 0.87 | 0.27 | 0.06 |
| 57 | 0.09 | 0.00 | 0.01 | 0.01 | 0.04 | 0.06 | 0.23 | 0.81 | 0.00 | 0.05 |
| 58 | 0.04 | 0.00 | 0.01 | 0.00 | 0.02 | 0.03 | 0.10 | 0.32 | 0.02 | 0.03 |
| 59 | 0.03 | 0.00 | 0.01 | 0.00 | 0.01 | 0.01 | 0.07 | 0.72 | 0.00 | 0.07 |
| 60 | 0.02 | 0.00 | 0.00 | 0.00 | 0.00 | 0.00 | 0.05 | 0.58 | 0.01 | 0.05 |
| 61 | 0.02 | 0.00 | 0.00 | 0.00 | 0.01 | 0.01 | 0.07 | 0.23 | 0.04 | 0.02 |
| 62 | 0.03 | 0.00 | 0.00 | 0.00 | 0.01 | 0.01 | 0.09 | 0.35 | 0.01 | 0.03 |
| 63 | 0.14 | 0.01 | 0.01 | 0.03 | 0.14 | 0.16 | 0.27 | 0.24 | 0.08 | 0.01 |
| 64 | 0.15 | 0.00 | 0.01 | 0.02 | 0.08 | 0.10 | 0.36 | 1.08 | 0.90 | 0.23 |
| 65 | 0.05 | 0.01 | 0.01 | 0.01 | 0.03 | 0.02 | 0.13 | 0.09 | 1.05 | 0.33 |
| 66 | 0.19 | 0.03 | 0.03 | 0.06 | 0.17 | 0.20 | 0.39 | 0.09 | 1.07 | 0.09 |
| 67 | 0.08 | 0.03 | 0.03 | 0.03 | 0.04 | 0.05 | 0.10 | 0.42 | 12.89 | 0.21 |
| 68 | 0.03 | 0.00 | 0.00 | 0.01 | 0.00 | 0.01 | 0.03 | 0.34 | 9.65 | 0.55 |
| 69 | 0.07 | 0.00 | 0.01 | 0.01 | 0.01 | 0.01 | 0.09 | 2.47 | 0.51 | 0.36 |
| 70 | 0.02 | 0.00 | 0.01 | 0.00 | 0.00 | 0.00 | 0.02 | 0.72 | 0.06 | 0.07 |
| 71 | 0.02 | 0.00 | 0.00 | 0.00 | 0.01 | 0.01 | 0.04 | 0.45 | 0.22 | 0.11 |
| 72 | 0.10 | 0.00 | 0.02 | 0.01 | 0.01 | 0.02 | 0.14 | 2.86 | 1.05 | 0.59 |
| 73 | 0.05 | 0.01 | 0.01 | 0.01 | 0.00 | 0.00 | 0.05 | 2.04 | 0.96 | 0.50 |
| 74 | 0.05 | 0.02 | 0.02 | 0.02 | 0.03 | 0.03 | 0.04 | 0.19 | 1.59 | 8.74 |
| 75 | 0.04 | 0.06 | 0.09 | 0.02 | 0.02 | 0.01 | 0.05 | 0.46 | 1.88 | 1.50 |
| 76 | 0.02 | 0.01 | 0.02 | 0.01 | 0.01 | 0.01 | 0.02 | 0.06 | 0.18 | 3.70 |
| 77 | 0.02 | 0.01 | 0.04 | 0.01 | 0.00 | 0.00 | 0.01 | 0.16 | 1.29 | 2.40 |
| 78 | 0.02 | 0.01 | 0.04 | 0.01 | 0.00 | 0.00 | 0.01 | 0.32 | 1.10 | 3.36 |
| 79 | 0.02 | 0.00 | 0.02 | 0.01 | 0.00 | 0.01 | 0.01 | 0.08 | 0.25 | 3.82 |
| 80 | 0.03 | 0.03 | 0.05 | 0.01 | 0.01 | 0.01 | 0.03 | 0.41 | 5.25 | 0.95 |
| 81 | 0.10 | 0.03 | 0.40 | 0.04 | 0.03 | 0.00 | 0.06 | 1.34 | 1.33 | 2.58 |
| 82 | 1.29 | 0.32 | 7.41 | 0.35 | 0.27 | 0.05 | 1.11 | 4.77 | 20.72 | 11.44 |
| 83 | 0.01 | 0.00 | 0.01 | 0.01 | 0.00 | 0.00 | 0.00 | 0.06 | 0.08 | 3.78 |
| 84 | 0.06 | 0.03 | 0.15 | 0.02 | 0.01 | 0.00 | 0.03 | 1.17 | 2.62 | 1.96 |
| 85 | 0.01 | 0.00 | 0.01 | 0.00 | 0.00 | 0.00 | 0.00 | 0.09 | 0.27 | 2.05 |
| 86 | 0.00 | 0.00 | 0.01 | 0.01 | 0.00 | 0.00 | 0.00 | 0.02 | 0.01 | 1.25 |
| 87 | 0.01 | 0.01 | 0.05 | 0.00 | 0.00 | 0.00 | 0.01 | 0.08 | 0.30 | 0.38 |
| 88 | 0.01 | 0.01 | 0.03 | 0.01 | 0.01 | 0.01 | 0.02 | 0.06 | 0.27 | 0.60 |
| 89 | 0.03 | 0.00 | 0.01 | 0.01 | 0.02 | 0.02 | 0.05 | 0.49 | 0.62 | 0.08 |
| 90 | 0.02 | 0.01 | 0.04 | 0.02 | 0.01 | 0.01 | 0.02 | 0.15 | 0.74 | 1.06 |
| 91 | 0.06 | 0.01 | 0.05 | 0.02 | 0.01 | 0.02 | 0.05 | 1.76 | 3.19 | 0.76 |
| 92 | 0.02 | 0.00 | 0.02 | 0.01 | 0.01 | 0.01 | 0.02 | 0.09 | 0.28 | 0.70 |
| 93 | 0.01 | 0.00 | 0.02 | 0.00 | 0.00 | 0.01 | 0.02 | 0.08 | 0.29 | 1.13 |
| 94 | 0.02 | 0.01 | 0.02 | 0.01 | 0.02 | 0.02 | 0.02 | 0.05 | 0.40 | 0.08 |
| 95 | 0.44 | 0.06 | 3.37 | 0.13 | 0.08 | 0.03 | 0.10 | 0.62 | 4.75 | 1.34 |
| 96 | 0.00 | 0.00 | 0.01 | 0.00 | 0.00 | 0.00 | 0.00 | 0.00 | 0.00 | 0.00 |
| 97 | 0.01 | 0.00 | 0.02 | 0.00 | 0.00 | 0.00 | 0.00 | 0.02 | 0.04 | 0.14 |
| 98 | 0.00 | 0.00 | 0.01 | 0.00 | 0.00 | 0.00 | 0.00 | 0.02 | 0.09 | 0.04 |
| 99 | 0.01 | 0.00 | 0.02 | 0.00 | 0.00 | 0.00 | 0.01 | 0.08 | 0.27 | 0.06 |
| 100 | 8.23 | 2.97 | 71.01 | 0.45 | 0.21 | 0.08 | 2.30 | 10.60 | 68.10 | 44.34 |
| Base | 2.03 | 0.06 | 0.20 | 0.44 | 0.44 | 0.34 | 0.51 | 0.04 | 0.003 | 0.003 |

Enrichments of Lowe et al.[22] CNEEs overlapping PhastCons elements called on 100-way alignment

C

| CNEE | Non-Exonic % | CpG Island |
|---|---|---|
| Euteleostomi | 0.07 | 4.8 |
| Tetrapod | 0.29 | 6.7 |
| Amniote | 0.56 | 2.8 |
| Mammal | 0.53 | 2.8 |
| Theria | 0.43 | 3.8 |
| Eutheria | 0.77 | 1.7 |
| Boreoeutheria | 0.08 | 2.5 |
| Euarchontoglire | 0.01 | 1.4 |
| Hominini | 0.01 | 0.2 |
| No assignment | 97.28 | 0.9 |
| Base | 100 | 0.43 |

Enrichments of Lowe et al.[22] CNEEs

D

| CNEE | Non-Exonic % | CpG Island |
|---|---|---|
| Euteleostomi | 0.06 | 3.0 |
| Tetrapod | 0.18 | 3.5 |
| Amniote | 0.45 | 1.7 |
| Mammal | 0.45 | 1.6 |
| Theria | 0.35 | 2.4 |
| Eutheria | 0.52 | 1.2 |
| Boreoeutheria | 0.03 | 2.2 |
| Euarchontoglire | 0.00 | 0.9 |
| Hominini | 0.00 | 0.4 |
| No assignment | 97.96 | 1.0 |
| Base | 100 | 0.43 |

Enrichments of Lowe et al.[22] CNEEs overlapping PhastCons elements called on 100-way alignment

**Figure S23: Relationship between conservation states and CNEEs from Lowe et al.[22]** **(A,B)** The conservation state enrichments for **(A)** the set of CNEEs defined in Lowe et al.[22] and **(B)** the subset restricted to those that overlap PhastCons element bases called on the 100-way alignment in which the conservation states were defined. The first column of each panel gives the enrichment of all CNEEs included, followed by the enrichments of subset of CNEEs, based on the branch of origin of the CNEE. The second through last columns are sorted in order of distance of branch point to human. The last row gives the % of the genome covered by the annotation in the column. **(C,D)** The enrichments within non-exonic regions of bases in CpG islands of **(C)** the set of CNEEs defined in Lowe et al.[22] and **(D)** the subset restricted to those that overlap PhastCons element bases called on the 100-way alignment in which the conservation states were defined. The first column of each panel gives the percentage of non-exonic bases found in each subset, and the second columns gives the enrichment for bases within non-exonic CpG islands. The last row gives the % of the non-exonic genome covered by the annotation in the column.

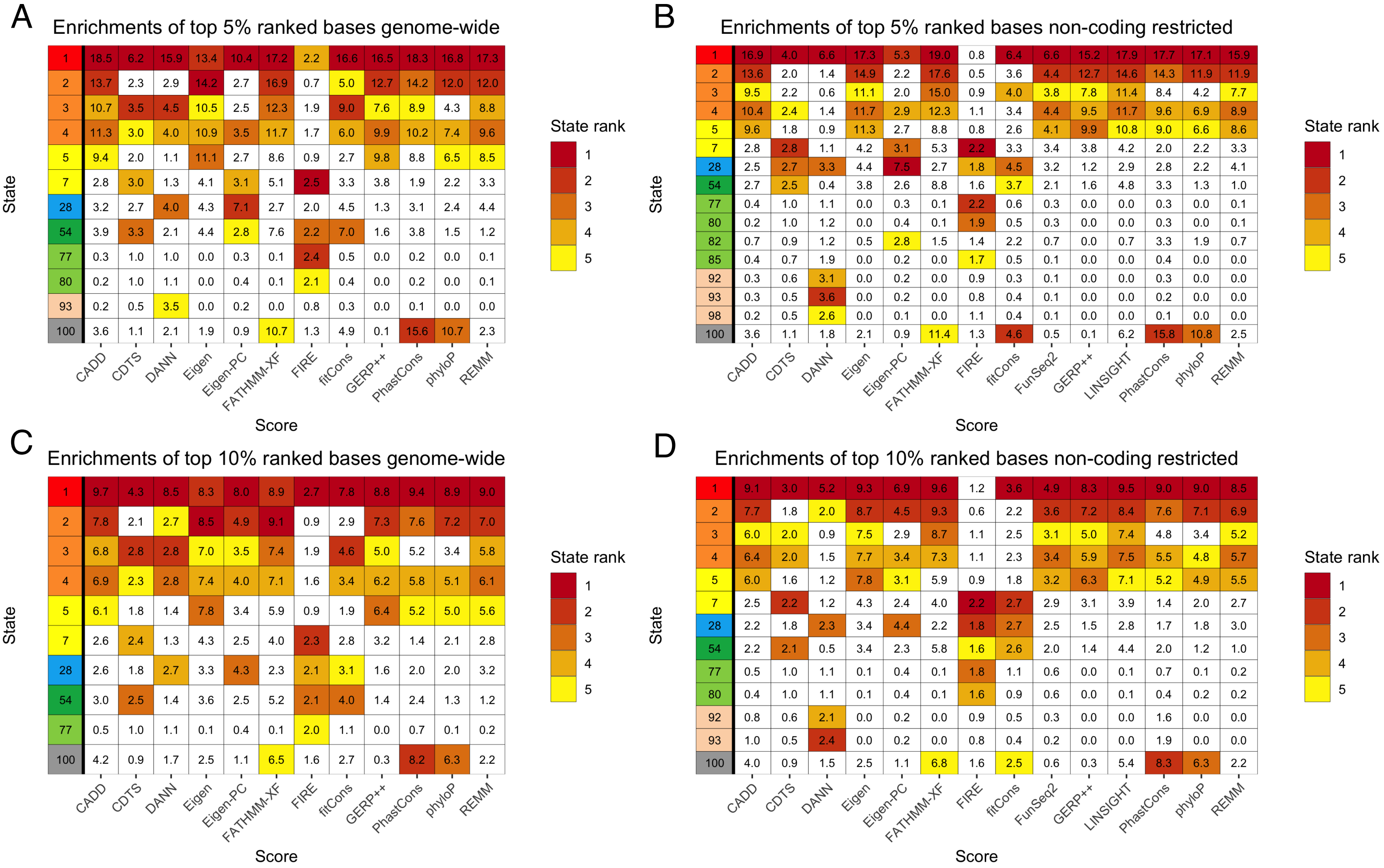


**Figure S24: Enrichments of selected conservation states for bases prioritized by variant prioritization scores.** Analogous figures to those shown in **Figure 6A** and **6B** of conservation state enrichments of top 1% bases of variant prioritization scores except shown here for **(A)** bases ranked in top 5% genome-wide **(B)** bases ranked in the top 5% of the genome restricted to non-coding bases **(C)** bases ranked in the top 10% genome-wide **(D)** bases ranked in the top 10% of the genome restricted to non-coding bases. Only states which were one of the five most enriched states by at least one variant prioritization score are shown. Coloring of enrichments is based on the rank of the state for the score as indicated in the color legend.

**A**

Enrichments of top 1% ranked bases genome-wide

| State | Genome % | CADD 1.4 | CADD 1.0 | Eigen | Eigen-PC | DANN | GERP++ | phyloP | PhastCons | REMM | fitCons | FIRE | FATHMM-XF | CDTS |
|---|---|---|---|---|---|---|---|---|---|---|---|---|---|---|
| 1 | 0.60 | 77.2 | 43.7 | 17.5 | 1.9 | 72.6 | 54.5 | 68.0 | 37.7 | 56.0 | 73.7 | 0.8 | 58.4 | 12.6 |
| 2 | 0.84 | 19.7 | 21.5 | 32.9 | 1.1 | 8.4 | 30.5 | 26.7 | 24.6 | 21.0 | 9.5 | 0.5 | 36.1 | 2.5 |
| 3 | 0.46 | 14.6 | 8.0 | 13.6 | 2.0 | 16.7 | 10.3 | 3.3 | 10.6 | 9.0 | 31.0 | 1.7 | 10.8 | 5.8 |
| 4 | 0.60 | 17.3 | 13.7 | 16.2 | 3.2 | 13.9 | 16.8 | 11.9 | 13.8 | 15.3 | 15.9 | 1.8 | 12.8 | 5.5 |
| 5 | 1.66 | 5.9 | 10.8 | 14.8 | 3.3 | 1.7 | 12.3 | 6.6 | 10.6 | 10.6 | 1.7 | 0.9 | 4.7 | 2.9 |
| 6 | 3.59 | 0.6 | 2.1 | 1.6 | 1.9 | 0.2 | 0.5 | 1.5 | 2.1 | 2.2 | 0.2 | 0.8 | 0.4 | 1.5 |
| 7 | 0.85 | 1.7 | 1.9 | 3.6 | 5.8 | 1.7 | 2.0 | 1.1 | 1.4 | 2.6 | 2.1 | 3.2 | 1.7 | 5.7 |
| 8 | 0.73 | 0.5 | 2.6 | 1.9 | 2.4 | 0.6 | 1.2 | 0.4 | 1.5 | 1.0 | 1.1 | 1.5 | 0.6 | 2.3 |
| 9 | 0.90 | 0.1 | 0.8 | 0.4 | 1.7 | 0.1 | 0.1 | 0.0 | 0.4 | 0.2 | 0.2 | 1.0 | 0.1 | 1.4 |
| 10 | 0.86 | 0.1 | 0.6 | 0.3 | 1.8 | 0.1 | 0.0 | 0.0 | 0.2 | 0.1 | 0.3 | 1.1 | 0.1 | 1.5 |
| 11 | 0.75 | 0.1 | 0.5 | 0.3 | 1.5 | 0.1 | 0.1 | 0.1 | 0.2 | 0.2 | 0.2 | 1.0 | 0.1 | 1.2 |
| 12 | 0.75 | 0.1 | 0.6 | 0.2 | 1.2 | 0.1 | 0.1 | 0.2 | 0.3 | 0.3 | 0.1 | 0.6 | 0.1 | 0.9 |
| 13 | 1.24 | 0.1 | 0.7 | 0.4 | 1.3 | 0.1 | 0.1 | 0.3 | 0.4 | 0.5 | 0.1 | 0.6 | 0.1 | 1.0 |
| 14 | 0.94 | 0.1 | 0.6 | 0.6 | 2.5 | 0.1 | 0.1 | 0.2 | 0.3 | 0.5 | 0.2 | 1.4 | 0.2 | 1.9 |
| 15 | 0.77 | 0.0 | 0.5 | 0.3 | 2.2 | 0.1 | 0.0 | 0.0 | 0.2 | 0.1 | 0.2 | 1.2 | 0.1 | 1.7 |
| 16 | 0.72 | 0.0 | 0.2 | 0.1 | 1.2 | 0.0 | 0.0 | 0.0 | 0.0 | 0.0 | 0.1 | 0.7 | 0.1 | 0.9 |
| 17 | 2.14 | 0.0 | 0.4 | 0.1 | 1.0 | 0.1 | 0.0 | 0.1 | 0.2 | 0.2 | 0.1 | 0.6 | 0.1 | 0.8 |
| 18 | 0.94 | 0.0 | 0.2 | 0.1 | 1.3 | 0.0 | 0.0 | 0.0 | 0.0 | 0.0 | 0.1 | 0.8 | 0.1 | 1.0 |
| 19 | 1.07 | 0.1 | 1.1 | 0.6 | 2.1 | 0.2 | 0.1 | 0.0 | 0.5 | 0.2 | 0.5 | 1.3 | 0.2 | 1.8 |
| 20 | 1.03 | 0.4 | 0.7 | 1.1 | 2.4 | 0.4 | 0.3 | 0.3 | 0.5 | 0.8 | 0.2 | 0.9 | 0.2 | 2.1 |
| 21 | 0.64 | 0.1 | 0.6 | 0.3 | 1.5 | 0.1 | 0.2 | 0.2 | 0.3 | 0.3 | 0.2 | 0.9 | 0.1 | 1.2 |
| 22 | 0.41 | 0.1 | 0.2 | 0.3 | 2.1 | 0.2 | 0.0 | 0.0 | 0.1 | 0.2 | 0.1 | 1.0 | 0.1 | 1.7 |
| 23 | 0.63 | 0.0 | 0.1 | 0.0 | 1.0 | 0.1 | 0.0 | 0.0 | 0.0 | 0.0 | 0.1 | 1.1 | 0.0 | 0.8 |
| 24 | 0.53 | 0.0 | 0.1 | 0.0 | 0.7 | 0.0 | 0.0 | 0.0 | 0.0 | 0.0 | 0.0 | 0.8 | 0.0 | 0.6 |
| 25 | 0.61 | 0.0 | 0.1 | 0.0 | 0.7 | 0.0 | 0.0 | 0.0 | 0.0 | 0.0 | 0.0 | 0.6 | 0.0 | 0.6 |
| 26 | 0.72 | 0.0 | 0.2 | 0.1 | 0.8 | 0.1 | 0.0 | 0.0 | 0.1 | 0.1 | 0.0 | 0.9 | 0.1 | 0.7 |
| 27 | 0.96 | 0.0 | 0.1 | 0.1 | 1.1 | 0.0 | 0.0 | 0.0 | 0.0 | 0.0 | 0.1 | 1.1 | 0.1 | 0.9 |
| 28 | 0.44 | 4.5 | 0.9 | 6.3 | 18.4 | 8.5 | 0.1 | 2.4 | 3.1 | 3.7 | 5.9 | 1.9 | 1.8 | 8.3 |
| 29 | 0.92 | 0.0 | 0.1 | 0.1 | 0.7 | 0.1 | 0.0 | 0.0 | 0.0 | 0.0 | 0.0 | 0.4 | 0.0 | 0.5 |
| 30 | 0.94 | 0.0 | 0.1 | 0.0 | 0.3 | 0.0 | 0.0 | 0.0 | 0.0 | 0.0 | 0.0 | 0.5 | 0.0 | 0.4 |
| 31 | 0.63 | 0.0 | 0.1 | 0.0 | 0.7 | 0.1 | 0.0 | 0.0 | 0.0 | 0.0 | 0.1 | 0.7 | 0.0 | 0.6 |
| 32 | 0.93 | 0.0 | 0.0 | 0.0 | 0.6 | 0.1 | 0.0 | 0.0 | 0.0 | 0.0 | 0.0 | 0.6 | 0.0 | 0.6 |
| 33 | 0.45 | 0.0 | 0.1 | 0.1 | 0.6 | 0.2 | 0.0 | 0.0 | 0.1 | 0.0 | 0.1 | 0.7 | 0.0 | 0.5 |
| 34 | 0.51 | 0.0 | 0.4 | 0.1 | 0.7 | 0.1 | 0.0 | 0.1 | 0.2 | 0.1 | 0.1 | 0.9 | 0.1 | 0.6 |
| 35 | 1.20 | 0.0 | 0.3 | 0.2 | 1.7 | 0.1 | 0.0 | 0.0 | 0.1 | 0.1 | 0.1 | 1.5 | 0.1 | 1.3 |
| 36 | 1.34 | 0.0 | 0.2 | 0.0 | 0.6 | 0.0 | 0.0 | 0.0 | 0.0 | 0.0 | 0.0 | 0.8 | 0.0 | 0.5 |
| 37 | 2.30 | 0.0 | 0.5 | 0.2 | 1.0 | 0.1 | 0.0 | 0.1 | 0.3 | 0.2 | 0.1 | 1.0 | 0.1 | 0.8 |
| 38 | 0.91 | 0.0 | 0.3 | 0.1 | 0.7 | 0.1 | 0.0 | 0.0 | 0.1 | 0.1 | 0.1 | 0.7 | 0.1 | 0.7 |
| 39 | 0.70 | 0.0 | 0.2 | 0.0 | 0.5 | 0.0 | 0.0 | 0.0 | 0.1 | 0.0 | 0.1 | 0.6 | 0.1 | 0.5 |
| 40 | 0.76 | 0.0 | 0.1 | 0.0 | 0.4 | 0.0 | 0.0 | 0.0 | 0.0 | 0.0 | 0.0 | 0.5 | 0.0 | 0.4 |
| 41 | 0.70 | 0.0 | 0.2 | 0.0 | 0.7 | 0.0 | 0.0 | 0.0 | 0.1 | 0.0 | 0.0 | 0.4 | 0.0 | 0.5 |
| 42 | 0.74 | 0.0 | 0.2 | 0.1 | 1.3 | 0.1 | 0.0 | 0.0 | 0.0 | 0.0 | 0.1 | 0.8 | 0.1 | 0.9 |
| 43 | 1.00 | 0.0 | 0.1 | 0.0 | 0.7 | 0.0 | 0.0 | 0.0 | 0.0 | 0.0 | 0.1 | 0.8 | 0.1 | 0.7 |
| 44 | 1.02 | 0.0 | 0.1 | 0.0 | 0.8 | 0.1 | 0.0 | 0.0 | 0.0 | 0.0 | 0.1 | 0.9 | 0.1 | 0.6 |
| 45 | 0.84 | 0.0 | 0.1 | 0.0 | 0.6 | 0.1 | 0.0 | 0.0 | 0.0 | 0.0 | 0.1 | 0.6 | 0.0 | 0.5 |
| 46 | 0.70 | 0.0 | 0.1 | 0.0 | 0.7 | 0.1 | 0.0 | 0.0 | 0.0 | 0.0 | 0.1 | 1.1 | 0.1 | 0.6 |
| 47 | 0.41 | 0.0 | 0.0 | 0.1 | 1.9 | 0.0 | 0.0 | 0.0 | 0.0 | 0.0 | 0.2 | 1.0 | 0.1 | 1.3 |
| 48 | 0.53 | 0.0 | 0.0 | 0.0 | 0.8 | 0.0 | 0.0 | 0.0 | 0.0 | 0.0 | 0.1 | 0.8 | 0.0 | 0.6 |
| 49 | 0.61 | 0.0 | 0.1 | 0.1 | 1.6 | 0.0 | 0.0 | 0.0 | 0.0 | 0.0 | 0.1 | 1.4 | 0.1 | 1.3 |
| 50 | 0.79 | 0.1 | 0.4 | 0.5 | 2.5 | 0.2 | 0.0 | 0.0 | 0.5 | 0.0 | 0.7 | 1.5 | 0.3 | 2.2 |
| 51 | 0.74 | 0.0 | 0.0 | 0.1 | 1.6 | 0.0 | 0.0 | 0.0 | 0.0 | 0.0 | 0.1 | 0.9 | 0.1 | 1.1 |
| 52 | 0.50 | 0.0 | 0.1 | 0.1 | 1.8 | 0.1 | 0.0 | 0.0 | 0.0 | 0.0 | 0.1 | 0.9 | 0.1 | 1.2 |
| 53 | 0.50 | 0.1 | 0.6 | 0.5 | 2.7 | 0.3 | 0.0 | 0.0 | 0.3 | 0.1 | 0.8 | 1.6 | 0.2 | 2.4 |
| 54 | 0.27 | 4.2 | 2.6 | 2.0 | 3.9 | 6.3 | 0.8 | 1.1 | 3.7 | 0.2 | 21.1 | 2.8 | 4.7 | 5.8 |
| 55 | 0.30 | 0.2 | 0.8 | 0.5 | 2.4 | 0.2 | 0.3 | 0.0 | 0.8 | 0.0 | 0.7 | 1.3 | 0.3 | 2.1 |
| 56 | 0.38 | 0.0 | 0.1 | 0.1 | 1.5 | 0.1 | 0.0 | 0.0 | 0.1 | 0.0 | 0.1 | 0.7 | 0.1 | 1.1 |
| 57 | 0.39 | 0.0 | 0.1 | 0.0 | 1.1 | 0.1 | 0.0 | 0.0 | 0.0 | 0.0 | 0.1 | 0.9 | 0.1 | 0.9 |
| 58 | 0.46 | 0.0 | 0.0 | 0.0 | 1.3 | 0.1 | 0.0 | 0.0 | 0.0 | 0.0 | 0.2 | 0.9 | 0.1 | 0.9 |
| 59 | 0.53 | 0.0 | 0.0 | 0.0 | 1.5 | 0.1 | 0.0 | 0.0 | 0.0 | 0.0 | 0.2 | 0.9 | 0.0 | 1.0 |
| 60 | 0.30 | 0.0 | 0.0 | 0.0 | 0.9 | 0.0 | 0.0 | 0.0 | 0.0 | 0.0 | 0.1 | 0.5 | 0.0 | 0.6 |
| 61 | 0.48 | 0.0 | 0.0 | 0.0 | 0.8 | 0.0 | 0.0 | 0.0 | 0.0 | 0.0 | 0.1 | 0.5 | 0.0 | 0.6 |
| 62 | 0.44 | 0.0 | 0.0 | 0.0 | 1.1 | 0.0 | 0.0 | 0.0 | 0.0 | 0.0 | 0.1 | 0.9 | 0.0 | 0.8 |
| 63 | 0.41 | 0.0 | 0.1 | 0.2 | 2.8 | 0.1 | 0.0 | 0.0 | 0.0 | 0.0 | 0.2 | 1.1 | 0.1 | 2.1 |
| 64 | 0.79 | 0.0 | 0.1 | 0.1 | 0.7 | 0.1 | 0.0 | 0.0 | 0.0 | 0.0 | 0.1 | 0.7 | 0.0 | 0.6 |
| 65 | 0.93 | 0.0 | 0.1 | 0.0 | 1.0 | 0.1 | 0.0 | 0.0 | 0.0 | 0.0 | 0.1 | 0.9 | 0.0 | 0.7 |
| 66 | 0.67 | 0.0 | 0.2 | 0.1 | 1.8 | 0.1 | 0.0 | 0.0 | 0.1 | 0.1 | 0.1 | 1.1 | 0.1 | 1.2 |
| 67 | 0.98 | 0.0 | 0.1 | 0.1 | 1.6 | 0.1 | 0.0 | 0.0 | 0.0 | 0.0 | 0.1 | 1.0 | 0.1 | 1.1 |
| 68 | 1.17 | 0.0 | 0.0 | 0.0 | 0.5 | 0.1 | 0.0 | 0.0 | 0.0 | 0.0 | 0.0 | 1.1 | 0.0 | 0.5 |
| 69 | 0.92 | 0.0 | 0.0 | 0.0 | 0.5 | 0.1 | 0.0 | 0.0 | 0.0 | 0.0 | 0.0 | 0.7 | 0.0 | 0.5 |
| 70 | 0.51 | 0.0 | 0.0 | 0.0 | 1.0 | 0.1 | 0.0 | 0.0 | 0.0 | 0.0 | 0.1 | 0.7 | 0.0 | 0.7 |
| 71 | 0.41 | 0.0 | 0.0 | 0.0 | 1.2 | 0.1 | 0.0 | 0.0 | 0.0 | 0.0 | 0.1 | 1.1 | 0.0 | 0.8 |
| 72 | 1.21 | 0.0 | 0.0 | 0.0 | 0.7 | 0.1 | 0.0 | 0.0 | 0.0 | 0.0 | 0.1 | 0.9 | 0.0 | 0.6 |
| 73 | 0.74 | 0.0 | 0.0 | 0.0 | 0.5 | 0.2 | 0.0 | 0.0 | 0.0 | 0.0 | 0.2 | 1.5 | 0.0 | 0.7 |
| 74 | 1.49 | 0.0 | 0.0 | 0.0 | 0.6 | 0.1 | 0.0 | 0.0 | 0.0 | 0.0 | 0.0 | 1.6 | 0.0 | 0.7 |
| 75 | 2.47 | 0.0 | 0.0 | 0.0 | 0.3 | 0.1 | 0.0 | 0.0 | 0.0 | 0.0 | 0.0 | 2.0 | 0.0 | 0.6 |
| 76 | 3.92 | 0.0 | 0.0 | 0.0 | 0.1 | 0.2 | 0.0 | 0.0 | 0.0 | 0.0 | 0.0 | 1.1 | 0.0 | 0.4 |
| 77 | 0.98 | 0.0 | 0.0 | 0.0 | 0.3 | 0.2 | 0.0 | 0.0 | 0.0 | 0.0 | 0.0 | 3.0 | 0.0 | 0.6 |
| 78 | 1.14 | 0.0 | 0.0 | 0.0 | 0.3 | 0.2 | 0.0 | 0.0 | 0.0 | 0.0 | 0.0 | 1.7 | 0.0 | 0.6 |
| 79 | 3.18 | 0.0 | 0.0 | 0.0 | 0.1 | 0.6 | 0.0 | 0.0 | 0.0 | 0.0 | 0.0 | 1.2 | 0.0 | 0.5 |
| 80 | 1.24 | 0.0 | 0.0 | 0.0 | 0.3 | 0.2 | 0.0 | 0.0 | 0.0 | 0.0 | 0.0 | 2.5 | 0.0 | 0.7 |
| 81 | 0.98 | 0.0 | 0.0 | 0.0 | 0.4 | 0.4 | 0.0 | 0.0 | 0.0 | 0.0 | 0.0 | 1.4 | 0.0 | 0.6 |
| 82 | 0.54 | 0.3 | 0.1 | 0.2 | 3.6 | 1.0 | 0.0 | 0.3 | 1.8 | 0.1 | 1.1 | 1.4 | 0.5 | 1.6 |
| 83 | 1.29 | 0.0 | 0.0 | 0.0 | 0.1 | 0.5 | 0.0 | 0.0 | 0.0 | 0.0 | 0.0 | 1.0 | 0.0 | 0.3 |
| 84 | 0.86 | 0.0 | 0.0 | 0.0 | 0.2 | 0.4 | 0.0 | 0.0 | 0.0 | 0.0 | 0.2 | 1.5 | 0.0 | 0.4 |
| 85 | 1.38 | 0.0 | 0.0 | 0.0 | 0.1 | 0.5 | 0.0 | 0.0 | 0.0 | 0.0 | 0.0 | 2.1 | 0.0 | 0.4 |
| 86 | 1.56 | 0.0 | 0.0 | 0.0 | 0.1 | 0.9 | 0.0 | 0.0 | 0.0 | 0.0 | 0.0 | 1.0 | 0.0 | 0.3 |
| 87 | 0.63 | 0.0 | 0.0 | 0.0 | 0.3 | 0.1 | 0.0 | 0.0 | 0.0 | 0.0 | 0.0 | 1.5 | 0.0 | 0.6 |
| 88 | 0.66 | 0.0 | 0.0 | 0.0 | 0.3 | 0.1 | 0.0 | 0.0 | 0.0 | 0.0 | 0.0 | 1.4 | 0.0 | 0.5 |
| 89 | 0.38 | 0.0 | 0.0 | 0.0 | 1.3 | 0.1 | 0.0 | 0.0 | 0.0 | 0.0 | 0.1 | 0.8 | 0.0 | 0.8 |
| 90 | 1.09 | 0.0 | 0.1 | 0.0 | 0.2 | 0.7 | 0.0 | 0.0 | 0.0 | 0.0 | 0.0 | 1.2 | 0.0 | 0.4 |
| 91 | 0.99 | 0.0 | 0.0 | 0.0 | 0.5 | 0.4 | 0.0 | 0.0 | 0.0 | 0.0 | 0.1 | 1.1 | 0.0 | 0.6 |
| 92 | 1.81 | 0.0 | 0.0 | 0.0 | 0.1 | 1.1 | 0.0 | 0.0 | 0.0 | 0.0 | 0.0 | 0.9 | 0.0 | 0.4 |
| 93 | 0.67 | 0.0 | 0.0 | 0.0 | 0.1 | 1.3 | 0.0 | 0.0 | 0.0 | 0.0 | 0.0 | 0.8 | 0.0 | 0.4 |
| 94 | 0.65 | 0.0 | 0.0 | 0.0 | 0.1 | 0.4 | 0.0 | 0.0 | 0.0 | 0.0 | 0.0 | 0.5 | 0.0 | 0.2 |
| 95 | 1.06 | 0.0 | 0.0 | 0.0 | 0.5 | 0.7 | 0.0 | 0.0 | 0.2 | 0.0 | 0.2 | 0.8 | 0.0 | 0.4 |
| 96 | 8.86 | 0.0 | 0.0 | 0.0 | 0.0 | 0.3 | 0.0 | 0.0 | 0.0 | 0.0 | 0.0 | 0.0 | 0.0 | 0.0 |
| 97 | 1.13 | 0.0 | 0.0 | 0.0 | 0.0 | 0.1 | 0.0 | 0.0 | 0.0 | 0.0 | 0.0 | 0.4 | 0.0 | 0.1 |
| 98 | 0.51 | 0.0 | 0.0 | 0.0 | 0.1 | 0.9 | 0.0 | 0.0 | 0.0 | 0.0 | 0.0 | 1.2 | 0.0 | 0.4 |
| 99 | 0.97 | 0.0 | 0.0 | 0.0 | 0.1 | 0.8 | 0.0 | 0.0 | 0.0 | 0.0 | 0.0 | 1.2 | 0.0 | 0.3 |
| 100 | 0.25 | 1.9 | 0.7 | 0.1 | 0.3 | 2.9 | 0.0 | 28.9 | 26.1 | 0.7 | 3.5 | 0.5 | 13.1 | 1.6 |
| Base | 100 | 1.00 | 1.00 | 1.00 | 1.00 | 1.00 | 1.00 | 1.00 | 2.13 | 1.02 | 1.02 | 1.02 | 1.00 | 1.00 |

**B**

Enrichments of top 1% ranked bases non-coding restricted

| State | Non coding region % | CADD 1.4 | CADD 1.0 | Eigen | Eigen-PC | DANN | GERP++ | phyloP | PhastCons | REMM | fitCons | FIRE | FATHMM-XF | CDTS | LINSIGHT | FunSeq2 |
|---|---|---|---|---|---|---|---|---|---|---|---|---|---|---|---|---|
| 1 | 0.13 | 57.8 | 30.0 | 50.8 | 2.6 | 16.3 | 56.5 | 71.7 | 45.9 | 53.9 | 15.2 | 0.5 | 80.7 | 6.8 | 50.5 | 6.3 |
| 2 | 0.83 | 33.3 | 25.3 | 35.8 | 1.0 | 0.8 | 36.2 | 32.9 | 30.4 | 27.8 | 8.3 | 0.3 | 47.2 | 2.1 | 24.9 | 3.5 |
| 3 | 0.33 | 13.3 | 10.2 | 15.0 | 1.3 | 0.4 | 14.7 | 4.2 | 12.3 | 10.7 | 9.3 | 0.9 | 20.4 | 2.7 | 6.6 | 3.1 |
| 4 | 0.54 | 19.2 | 14.8 | 18.1 | 3.1 | 1.7 | 19.5 | 11.9 | 15.4 | 16.8 | 5.9 | 1.3 | 17.7 | 4.1 | 13.1 | 5.4 |
| 5 | 1.80 | 15.8 | 13.4 | 14.9 | 3.1 | 0.6 | 17.2 | 9.3 | 13.0 | 15.0 | 3.9 | 0.8 | 6.9 | 2.8 | 13.1 | 5.0 |
| 6 | 3.95 | 2.5 | 2.9 | 1.6 | 1.8 | 0.2 | 1.8 | 3.7 | 2.6 | 3.3 | 1.5 | 0.8 | 0.8 | 1.6 | 5.4 | 2.8 |
| 7 | 0.91 | 2.5 | 2.4 | 3.6 | 5.5 | 1.0 | 3.3 | 1.8 | 1.6 | 3.5 | 5.0 | 2.8 | 2.6 | 5.5 | 2.5 | 3.7 |
| 8 | 0.79 | 2.0 | 3.6 | 1.8 | 2.2 | 0.2 | 2.3 | 0.8 | 1.8 | 1.8 | 3.5 | 1.4 | 0.9 | 2.3 | 1.2 | 2.7 |
| 9 | 0.99 | 0.4 | 1.2 | 0.4 | 1.6 | 0.1 | 0.2 | 0.0 | 0.4 | 0.5 | 1.3 | 0.9 | 0.2 | 1.5 | 0.8 | 1.9 |
| 10 | 0.95 | 0.2 | 0.8 | 0.3 | 1.7 | 0.1 | 0.1 | 0.0 | 0.2 | 0.3 | 1.5 | 1.0 | 0.2 | 1.6 | 0.3 | 1.7 |
| 11 | 0.83 | 0.2 | 0.7 | 0.3 | 1.4 | 0.1 | 0.3 | 0.6 | 0.2 | 0.4 | 1.2 | 1.0 | 0.2 | 1.2 | 0.8 | 1.5 |
| 12 | 0.83 | 0.3 | 0.8 | 0.2 | 1.1 | 0.1 | 0.2 | 0.8 | 0.4 | 0.5 | 0.8 | 0.5 | 0.2 | 0.9 | 0.8 | 1.2 |
| 13 | 1.37 | 0.5 | 1.1 | 0.4 | 1.2 | 0.1 | 0.4 | 1.2 | 0.5 | 0.7 | 0.8 | 0.5 | 0.2 | 1.1 | 1.5 | 1.4 |
| 14 | 1.03 | 0.4 | 0.9 | 0.6 | 2.3 | 0.2 | 0.4 | 0.9 | 0.4 | 0.8 | 1.5 | 1.4 | 0.3 | 1.9 | 1.3 | 2.5 |
| 15 | 0.85 | 0.2 | 0.8 | 0.3 | 2.0 | 0.1 | 0.1 | 0.0 | 0.2 | 0.3 | 1.5 | 1.1 | 0.2 | 1.8 | 0.4 | 1.9 |
| 16 | 0.79 | 0.0 | 0.3 | 0.1 | 1.1 | 0.1 | 0.0 | 0.0 | 0.0 | 0.1 | 0.7 | 0.7 | 0.1 | 0.9 | 0.1 | 1.0 |
| 17 | 2.36 | 0.2 | 0.6 | 0.1 | 0.9 | 0.1 | 0.1 | 1.0 | 0.2 | 0.3 | 0.7 | 0.6 | 0.2 | 0.8 | 0.9 | 1.2 |
| 18 | 1.04 | 0.0 | 0.3 | 0.1 | 1.2 | 0.1 | 0.0 | 0.0 | 0.0 | 0.1 | 0.8 | 0.7 | 0.1 | 1.0 | 0.1 | 1.0 |
| 19 | 1.17 | 0.6 | 1.5 | 0.6 | 2.0 | 0.1 | 0.3 | 0.0 | 0.6 | 0.5 | 2.0 | 1.1 | 0.3 | 1.8 | 0.4 | 2.0 |
| 20 | 1.13 | 0.7 | 1.0 | 1.0 | 2.3 | 0.6 | 0.5 | 1.1 | 0.6 | 1.2 | 1.1 | 0.8 | 0.3 | 2.1 | 1.3 | 1.9 |
| 21 | 0.71 | 0.3 | 0.8 | 0.3 | 1.4 | 0.1 | 0.3 | 0.7 | 0.3 | 0.5 | 1.2 | 0.8 | 0.2 | 1.2 | 0.8 | 1.4 |
| 22 | 0.45 | 0.1 | 0.3 | 0.3 | 1.9 | 0.4 | 0.0 | 0.2 | 0.1 | 0.4 | 0.9 | 1.0 | 0.1 | 1.7 | 0.3 | 1.4 |
| 23 | 0.69 | 0.0 | 0.2 | 0.0 | 0.9 | 0.2 | 0.0 | 0.0 | 0.0 | 0.1 | 0.6 | 1.0 | 0.1 | 0.8 | 0.0 | 0.8 |
| 24 | 0.59 | 0.0 | 0.2 | 0.0 | 0.6 | 0.2 | 0.0 | 0.0 | 0.0 | 0.0 | 0.6 | 0.7 | 0.1 | 0.6 | 0.0 | 0.7 |
| 25 | 0.68 | 0.0 | 0.2 | 0.0 | 0.6 | 0.1 | 0.0 | 0.0 | 0.0 | 0.1 | 0.5 | 0.5 | 0.1 | 0.6 | 0.1 | 0.7 |
| 26 | 0.80 | 0.1 | 0.4 | 0.1 | 0.7 | 0.1 | 0.0 | 0.2 | 0.1 | 0.1 | 0.6 | 0.8 | 0.1 | 0.7 | 0.2 | 0.9 |
| 27 | 1.05 | 0.0 | 0.2 | 0.1 | 1.1 | 0.1 | 0.0 | 0.0 | 0.0 | 0.0 | 0.9 | 1.0 | 0.1 | 0.9 | 0.0 | 1.0 |
| 28 | 0.44 | 3.1 | 0.4 | 6.2 | 18.6 | 7.6 | 0.3 | 2.9 | 3.1 | 5.0 | 3.7 | 1.8 | 2.4 | 8.0 | 0.6 | 5.0 |
| 29 | 1.01 | 0.0 | 0.2 | 0.1 | 0.7 | 0.2 | 0.0 | 0.0 | 0.0 | 0.1 | 0.4 | 0.4 | 0.1 | 0.5 | 0.1 | 0.6 |
| 30 | 1.03 | 0.0 | 0.2 | 0.0 | 0.3 | 0.1 | 0.0 | 0.0 | 0.0 | 0.0 | 0.3 | 0.4 | 0.1 | 0.4 | 0.0 | 0.5 |
| 31 | 0.70 | 0.0 | 0.1 | 0.0 | 0.6 | 0.2 | 0.0 | 0.0 | 0.0 | 0.1 | 0.4 | 0.7 | 0.1 | 0.7 | 0.0 | 0.6 |
| 32 | 1.00 | 0.0 | 0.1 | 0.0 | 0.5 | 0.6 | 0.0 | 0.0 | 0.0 | 0.0 | 0.4 | 0.6 | 0.0 | 0.6 | 0.0 | 0.4 |
| 33 | 0.49 | 0.0 | 0.2 | 0.1 | 0.6 | 0.8 | 0.0 | 0.0 | 0.1 | 0.1 | 0.4 | 0.7 | 0.1 | 0.5 | 0.0 | 0.4 |
| 34 | 0.56 | 0.1 | 0.5 | 0.1 | 0.7 | 0.2 | 0.1 | 0.4 | 0.2 | 0.2 | 0.6 | 0.8 | 0.1 | 0.6 | 0.4 | 0.8 |
| 35 | 1.33 | 0.1 | 0.4 | 0.2 | 1.6 | 0.2 | 0.1 | 0.2 | 0.1 | 0.3 | 1.2 | 1.3 | 0.2 | 1.4 | 0.3 | 1.6 |
| 36 | 1.47 | 0.0 | 0.3 | 0.0 | 0.5 | 0.1 | 0.0 | 0.0 | 0.1 | 0.1 | 0.5 | 0.7 | 0.1 | 0.6 | 0.1 | 0.7 |
| 37 | 2.54 | 0.2 | 0.7 | 0.2 | 0.9 | 0.1 | 0.2 | 0.8 | 0.3 | 0.4 | 0.9 | 0.9 | 0.2 | 0.9 | 0.9 | 1.4 |
| 38 | 1.00 | 0.1 | 0.4 | 0.1 | 0.6 | 0.1 | 0.0 | 0.2 | 0.1 | 0.1 | 0.6 | 0.7 | 0.1 | 0.7 | 0.3 | 0.9 |
| 39 | 0.76 | 0.0 | 0.3 | 0.0 | 0.5 | 0.1 | 0.0 | 0.1 | 0.1 | 0.1 | 0.5 | 0.5 | 0.1 | 0.5 | 0.1 | 0.6 |
| 40 | 0.83 | 0.0 | 0.2 | 0.0 | 0.3 | 0.2 | 0.0 | 0.0 | 0.0 | 0.0 | 0.3 | 0.4 | 0.0 | 0.4 | 0.0 | 0.4 |
| 41 | 0.78 | 0.0 | 0.3 | 0.0 | 0.7 | 0.1 | 0.0 | 0.1 | 0.1 | 0.1 | 0.4 | 0.4 | 0.1 | 0.6 | 0.1 | 0.7 |
| 42 | 0.82 | 0.0 | 0.3 | 0.1 | 1.2 | 0.1 | 0.0 | 0.1 | 0.0 | 0.1 | 0.6 | 0.8 | 0.1 | 0.9 | 0.1 | 1.0 |
| 43 | 1.10 | 0.0 | 0.2 | 0.0 | 0.7 | 0.2 | 0.0 | 0.0 | 0.0 | 0.1 | 0.5 | 0.7 | 0.1 | 0.7 | 0.0 | 0.8 |
| 44 | 1.12 | 0.0 | 0.1 | 0.0 | 0.7 | 0.6 | 0.0 | 0.0 | 0.0 | 0.0 | 0.5 | 0.9 | 0.1 | 0.7 | 0.0 | 0.5 |
| 45 | 0.93 | 0.0 | 0.2 | 0.0 | 0.6 | 0.2 | 0.0 | 0.0 | 0.0 | 0.0 | 0.5 | 0.6 | 0.1 | 0.6 | 0.0 | 0.6 |
| 46 | 0.77 | 0.0 | 0.2 | 0.0 | 0.6 | 0.2 | 0.0 | 0.0 | 0.0 | 0.1 | 0.7 | 1.0 | 0.1 | 0.7 | 0.1 | 0.8 |
| 47 | 0.45 | 0.0 | 0.1 | 0.1 | 1.7 | 0.1 | 0.0 | 0.0 | 0.0 | 0.0 | 1.2 | 0.9 | 0.1 | 1.4 | 0.0 | 1.2 |
| 48 | 0.59 | 0.0 | 0.1 | 0.0 | 0.7 | 0.0 | 0.0 | 0.0 | 0.0 | 0.0 | 0.6 | 0.7 | 0.1 | 0.7 | 0.0 | 0.6 |
| 49 | 0.67 | 0.0 | 0.1 | 0.1 | 1.5 | 0.1 | 0.0 | 0.0 | 0.0 | 0.0 | 1.1 | 1.3 | 0.1 | 1.3 | 0.0 | 1.2 |
| 50 | 0.87 | 0.2 | 0.6 | 0.4 | 2.3 | 0.1 | 0.1 | 0.0 | 0.6 | 0.1 | 2.1 | 1.3 | 0.4 | 2.2 | 0.0 | 2.0 |
| 51 | 0.81 | 0.0 | 0.1 | 0.1 | 1.5 | 0.1 | 0.0 | 0.0 | 0.0 | 0.0 | 0.9 | 0.8 | 0.1 | 1.2 | 0.0 | 1.1 |
| 52 | 0.55 | 0.0 | 0.2 | 0.1 | 1.7 | 0.1 | 0.0 | 0.0 | 0.0 | 0.1 | 1.0 | 0.8 | 0.1 | 1.3 | 0.0 | 1.1 |
| 53 | 0.55 | 0.3 | 0.9 | 0.5 | 2.5 | 0.1 | 0.1 | 0.0 | 0.3 | 0.3 | 2.4 | 1.4 | 0.3 | 2.4 | 0.1 | 2.0 |
| 54 | 0.23 | 1.2 | 2.2 | 1.5 | 3.4 | 0.3 | 1.4 | 1.3 | 3.8 | 0.3 | 7.5 | 2.2 | 7.9 | 4.0 | 0.1 | 2.2 |
| 55 | 0.33 | 0.4 | 1.0 | 0.4 | 2.3 | 0.3 | 0.6 | 0.0 | 1.0 | 0.0 | 2.0 | 1.2 | 0.5 | 2.1 | 0.0 | 2.0 |
| 56 | 0.42 | 0.0 | 0.1 | 0.1 | 1.4 | 0.4 | 0.0 | 0.0 | 0.1 | 0.0 | 0.8 | 0.7 | 0.1 | 1.1 | 0.0 | 1.0 |
| 57 | 0.43 | 0.0 | 0.1 | 0.0 | 1.1 | 0.6 | 0.0 | 0.0 | 0.0 | 0.0 | 0.7 | 0.8 | 0.1 | 0.9 | 0.0 | 0.8 |
| 58 | 0.51 | 0.0 | 0.1 | 0.0 | 1.2 | 0.1 | 0.0 | 0.0 | 0.0 | 0.0 | 0.8 | 0.9 | 0.1 | 1.0 | 0.0 | 0.8 |
| 59 | 0.58 | 0.0 | 0.0 | 0.0 | 1.4 | 0.2 | 0.0 | 0.0 | 0.0 | 0.0 | 0.7 | 0.9 | 0.1 | 1.0 | 0.0 | 0.8 |
| 60 | 0.34 | 0.0 | 0.0 | 0.0 | 0.8 | 0.1 | 0.0 | 0.0 | 0.0 | 0.0 | 0.4 | 0.5 | 0.0 | 0.6 | 0.0 | 0.5 |
| 61 | 0.53 | 0.0 | 0.0 | 0.0 | 0.7 | 0.1 | 0.0 | 0.0 | 0.0 | 0.0 | 0.5 | 0.5 | 0.0 | 0.6 | 0.0 | 0.5 |
| 62 | 0.49 | 0.0 | 0.0 | 0.0 | 1.0 | 0.1 | 0.0 | 0.0 | 0.0 | 0.0 | 0.7 | 0.8 | 0.1 | 0.8 | 0.0 | 0.7 |
| 63 | 0.45 | 0.0 | 0.1 | 0.2 | 2.6 | 0.2 | 0.0 | 0.0 | 0.0 | 0.1 | 1.3 | 1.0 | 0.1 | 2.1 | 0.0 | 1.3 |
| 64 | 0.87 | 0.0 | 0.2 | 0.1 | 0.7 | 0.2 | 0.0 | 0.0 | 0.0 | 0.1 | 0.5 | 0.6 | 0.1 | 0.6 | 0.1 | 0.6 |
| 65 | 1.03 | 0.0 | 0.1 | 0.0 | 0.9 | 0.3 | 0.0 | 0.0 | 0.0 | 0.0 | 0.6 | 0.8 | 0.1 | 0.8 | 0.0 | 0.6 |
| 66 | 0.73 | 0.1 | 0.3 | 0.1 | 1.6 | 0.2 | 0.1 | 0.0 | 0.1 | 0.2 | 1.1 | 1.0 | 0.1 | 1.3 | 0.1 | 1.2 |
| 67 | 1.08 | 0.0 | 0.1 | 0.1 | 1.4 | 0.3 | 0.0 | 0.0 | 0.0 | 0.1 | 0.8 | 0.9 | 0.1 | 1.1 | 0.1 | 0.8 |
| 68 | 1.30 | 0.0 | 0.1 | 0.0 | 0.4 | 0.3 | 0.0 | 0.0 | 0.0 | 0.0 | 0.4 | 1.0 | 0.0 | 0.6 | 0.0 | 0.3 |
| 69 | 1.01 | 0.0 | 0.1 | 0.0 | 0.4 | 0.6 | 0.0 | 0.0 | 0.0 | 0.0 | 0.3 | 0.6 | 0.0 | 0.5 | 0.0 | 0.3 |
| 70 | 0.55 | 0.0 | 0.0 | 0.0 | 0.9 | 0.3 | 0.0 | 0.0 | 0.0 | 0.0 | 0.4 | 0.7 | 0.0 | 0.7 | 0.0 | 0.5 |
| 71 | 0.45 | 0.0 | 0.0 | 0.0 | 1.1 | 0.2 | 0.0 | 0.0 | 0.0 | 0.0 | 0.6 | 1.0 | 0.0 | 0.8 | 0.0 | 0.6 |
| 72 | 1.31 | 0.0 | 0.1 | 0.0 | 0.6 | 0.6 | 0.0 | 0.0 | 0.0 | 0.0 | 0.4 | 0.9 | 0.1 | 0.6 | 0.0 | 0.4 |
| 73 | 0.81 | 0.0 | 0.0 | 0.0 | 0.5 | 0.6 | 0.0 | 0.0 | 0.0 | 0.0 | 0.6 | 1.4 | 0.1 | 0.7 | 0.0 | 0.4 |
| 74 | 1.64 | 0.0 | 0.1 | 0.0 | 0.6 | 0.3 | 0.0 | 0.0 | 0.0 | 0.0 | 0.6 | 1.5 | 0.1 | 0.7 | 0.0 | 0.5 |
| 75 | 2.73 | 0.0 | 0.0 | 0.0 | 0.3 | 0.7 | 0.0 | 0.0 | 0.0 | 0.0 | 0.4 | 1.8 | 0.0 | 0.6 | 0.0 | 0.3 |
| 76 | 4.33 | 0.0 | 0.1 | 0.0 | 0.1 | 1.5 | 0.0 | 0.0 | 0.0 | 0.0 | 0.3 | 1.0 | 0.0 | 0.4 | 0.0 | 0.2 |
| 77 | 1.08 | 0.0 | 0.0 | 0.0 | 0.2 | 1.0 | 0.0 | 0.0 | 0.0 | 0.0 | 0.4 | 2.8 | 0.0 | 0.7 | 0.0 | 0.2 |
| 78 | 1.26 | 0.0 | 0.1 | 0.0 | 0.3 | 1.0 | 0.0 | 0.0 | 0.0 | 0.0 | 0.3 | 1.6 | 0.0 | 0.6 | 0.0 | 0.3 |
| 79 | 3.51 | 0.0 | 0.0 | 0.0 | 0.1 | 3.0 | 0.0 | 0.0 | 0.0 | 0.0 | 0.2 | 1.1 | 0.0 | 0.5 | 0.0 | 0.2 |
| 80 | 1.37 | 0.0 | 0.0 | 0.0 | 0.3 | 1.0 | 0.0 | 0.0 | 0.0 | 0.0 | 0.3 | 2.3 | 0.0 | 0.8 | 0.0 | 0.3 |
| 81 | 1.08 | 0.0 | 0.0 | 0.0 | 0.3 | 2.0 | 0.0 | 0.0 | 0.0 | 0.0 | 0.3 | 1.3 | 0.0 | 0.7 | 0.0 | 0.3 |
| 82 | 0.57 | 0.1 | 0.1 | 0.2 | 3.4 | 1.4 | 0.0 | 0.8 | 2.2 | 0.2 | 3.0 | 1.3 | 0.7 | 1.6 | 0.0 | 0.8 |
| 83 | 1.38 | 0.0 | 0.1 | 0.0 | 0.1 | 2.6 | 0.0 | 0.0 | 0.0 | 0.0 | 0.3 | 1.0 | 0.0 | 0.3 | 0.0 | 0.1 |
| 84 | 0.91 | 0.0 | 0.0 | 0.0 | 0.2 | 1.6 | 0.0 | 0.0 | 0.0 | 0.0 | 0.5 | 1.5 | 0.0 | 0.4 | 0.0 | 0.2 |
| 85 | 1.46 | 0.0 | 0.0 | 0.0 | 0.1 | 2.6 | 0.0 | 0.0 | 0.0 | 0.0 | 0.3 | 2.1 | 0.0 | 0.5 | 0.0 | 0.2 |
| 86 | 1.73 | 0.0 | 0.0 | 0.0 | 0.0 | 3.9 | 0.0 | 0.0 | 0.0 | 0.0 | 0.2 | 1.0 | 0.0 | 0.3 | 0.0 | 0.1 |
| 87 | 0.69 | 0.0 | 0.0 | 0.0 | 0.2 | 0.4 | 0.0 | 0.0 | 0.0 | 0.0 | 0.3 | 1.4 | 0.0 | 0.6 | 0.0 | 0.3 |
| 88 | 0.73 | 0.0 | 0.0 | 0.0 | 0.3 | 0.3 | 0.0 | 0.0 | 0.0 | 0.0 | 0.4 | 1.3 | 0.0 | 0.6 | 0.0 | 0.3 |
| 89 | 0.41 | 0.0 | 0.0 | 0.0 | 1.2 | 0.6 | 0.0 | 0.0 | 0.0 | 0.0 | 0.6 | 0.7 | 0.1 | 0.8 | 0.0 | 0.6 |
| 90 | 1.19 | 0.0 | 0.2 | 0.0 | 0.2 | 3.6 | 0.0 | 0.0 | 0.0 | 0.0 | 0.3 | 1.1 | 0.0 | 0.5 | 0.0 | 0.2 |
| 91 | 1.07 | 0.0 | 0.1 | 0.0 | 0.5 | 1.7 | 0.0 | 0.0 | 0.0 | 0.0 | 0.5 | 1.1 | 0.1 | 0.7 | 0.0 | 0.3 |
| 92 | 2.00 | 0.0 | 0.0 | 0.0 | 0.1 | 4.9 | 0.0 | 0.0 | 0.0 | 0.0 | 0.2 | 0.8 | 0.0 | 0.4 | 0.0 | 0.1 |
| 93 | 0.74 | 0.0 | 0.0 | 0.0 | 0.1 | 5.8 | 0.0 | 0.0 | 0.0 | 0.0 | 0.2 | 0.7 | 0.0 | 0.4 | 0.0 | 0.2 |
| 94 | 0.72 | 0.0 | 0.0 | 0.0 | 0.1 | 1.7 | 0.0 | 0.0 | 0.0 | 0.0 | 0.3 | 0.5 | 0.0 | 0.2 | 0.0 | 0.1 |
| 95 | 1.08 | 0.0 | 0.0 | 0.0 | 0.4 | 2.0 | 0.0 | 0.0 | 0.1 | 0.0 | 0.9 | 0.8 | 0.1 | 0.5 | 0.0 | 0.2 |
| 96 | 1.33 | 0.0 | 0.0 | 0.1 | 0.2 | 4.2 | 0.0 | 0.0 | 0.0 | 0.0 | 0.5 | 0.3 | 0.4 | 0.2 | 0.0 | 0.1 |
| 97 | 0.96 | 0.0 | 0.0 | 0.0 | 0.0 | 0.4 | 0.0 | 0.0 | 0.0 | 0.0 | 0.5 | 0.5 | 0.0 | 0.1 | 0.0 | 0.1 |
| 98 | 0.56 | 0.0 | 0.0 | 0.0 | 0.1 | 4.1 | 0.0 | 0.0 | 0.0 | 0.0 | 0.4 | 1.1 | 0.0 | 0.4 | 0.0 | 0.1 |
| 99 | 1.05 | 0.0 | 0.0 | 0.0 | 0.0 | 3.5 | 0.0 | 0.0 | 0.0 | 0.0 | 0.2 | 1.1 | 0.0 | 0.3 | 0.0 | 0.1 |
| 100 | 0.26 | 0.7 | 0.6 | 0.1 | 0.3 | 2.0 | 0.0 | 34.5 | 32.8 | 1.3 | 14.7 | 0.5 | 17.9 | 1.6 | 0.0 | 0.3 |
| Base | 90.44 | 0.99 | 0.99 | 0.99 | 0.99 | 0.99 | 1.00 | 0.99 | 1.71 | 1.01 | 1.01 | 1.01 | 0.99 | 0.99 | 0.99 | 0.99 |

**Figure S25: Enrichment of all conservation states for top 1% of bases prioritized by variant prioritization scores.** The figure displays for top 1% prioritized bases **(A)** genome-wide and **(B)** in non-coding regions by variant prioritization scores the same enrichments as shown in **Figure 6A** and **6B** except here enrichments for all conservation states are shown and also included is the original version of the CADD score in addition to v1.4. Coloring of enrichments is based on their value in a column specific manner. The second columns gives the percentage of the background region used to compute the enrichments falling in each state, which is for **(A)** the whole genome and for **(B)** bases scored by both LINSIGHT and FunSeq2. The last line in both heatmaps gives the actual percentage of the background set covered by each set of prioritized bases, which can differ from 1.00% because of how ties were handled.

A

Enrichments of top 5% ranked bases genome-wide

| State | Genome % | CADD 1.4 | CADD 1.0 | Eigen | Eigen-PC | DANN | GERP++ | phyloP | PhastCons | REMM | fitCons | FIRE | FATHMM-XF | CDTS |
|---|---|---|---|---|---|---|---|---|---|---|---|---|---|---|
| 1 | 0.60 | 18.5 | 15.7 | 13.4 | 10.4 | 15.9 | 16.5 | 16.8 | 18.3 | 17.3 | 16.6 | 2.2 | 17.2 | 6.2 |
| 2 | 0.84 | 13.7 | 10.6 | 14.2 | 2.7 | 2.9 | 12.7 | 12.0 | 14.2 | 12.0 | 5.0 | 0.7 | 16.9 | 2.3 |
| 3 | 0.46 | 10.7 | 7.0 | 10.5 | 2.5 | 4.5 | 7.6 | 4.3 | 8.9 | 8.8 | 9.0 | 1.9 | 12.3 | 3.5 |
| 4 | 0.60 | 11.3 | 8.5 | 10.9 | 3.5 | 4.0 | 9.9 | 7.4 | 10.2 | 9.6 | 6.0 | 1.7 | 11.7 | 3.0 |
| 5 | 1.66 | 9.4 | 7.1 | 11.1 | 2.7 | 1.1 | 9.8 | 6.5 | 8.8 | 8.5 | 2.7 | 0.9 | 8.6 | 2.0 |
| 6 | 3.59 | 2.8 | 2.6 | 3.6 | 1.8 | 0.6 | 4.5 | 3.2 | 2.8 | 3.2 | 1.4 | 0.8 | 2.0 | 1.4 |
| 7 | 0.85 | 2.8 | 2.7 | 4.1 | 3.1 | 1.3 | 3.8 | 2.2 | 1.9 | 3.3 | 3.3 | 2.5 | 5.1 | 3.0 |
| 8 | 0.73 | 3.3 | 3.2 | 4.4 | 2.1 | 0.6 | 3.3 | 1.2 | 2.4 | 3.0 | 2.4 | 1.2 | 3.3 | 1.8 |
| 9 | 0.90 | 1.2 | 1.6 | 1.5 | 1.7 | 0.4 | 1.4 | 0.4 | 0.9 | 1.6 | 1.3 | 0.9 | 0.7 | 1.4 |
| 10 | 0.86 | 0.8 | 1.3 | 1.1 | 1.7 | 0.3 | 0.7 | 0.2 | 0.5 | 1.1 | 1.4 | 1.0 | 0.7 | 1.4 |
| 11 | 0.75 | 0.7 | 1.1 | 0.9 | 1.5 | 0.5 | 1.5 | 1.1 | 0.6 | 1.2 | 1.1 | 1.0 | 0.7 | 1.2 |
| 12 | 0.75 | 0.8 | 1.2 | 0.9 | 1.2 | 0.5 | 1.6 | 1.3 | 0.8 | 1.2 | 0.9 | 0.6 | 0.8 | 1.0 |
| 13 | 1.24 | 1.0 | 1.4 | 1.3 | 1.3 | 0.5 | 2.2 | 1.6 | 1.0 | 1.4 | 1.0 | 0.7 | 0.8 | 1.1 |
| 14 | 0.94 | 1.0 | 1.3 | 1.5 | 2.0 | 0.5 | 2.0 | 1.4 | 0.9 | 1.8 | 1.5 | 1.3 | 0.8 | 1.6 |
| 15 | 0.77 | 0.7 | 1.3 | 1.2 | 1.9 | 0.3 | 0.8 | 0.2 | 0.5 | 1.2 | 1.4 | 1.0 | 0.6 | 1.5 |
| 16 | 0.72 | 0.2 | 0.6 | 0.3 | 1.2 | 0.3 | 0.4 | 0.1 | 0.2 | 0.6 | 0.9 | 0.8 | 0.2 | 1.1 |
| 17 | 2.14 | 0.6 | 1.0 | 0.8 | 1.1 | 0.5 | 1.9 | 1.8 | 0.7 | 1.1 | 0.8 | 0.7 | 0.8 | 1.0 |
| 18 | 0.94 | 0.2 | 0.6 | 0.3 | 1.3 | 0.3 | 0.2 | 0.1 | 0.1 | 0.5 | 0.9 | 0.8 | 0.2 | 1.1 |
| 19 | 1.07 | 1.5 | 1.9 | 2.0 | 1.9 | 0.4 | 1.1 | 0.3 | 1.0 | 1.6 | 1.7 | 1.0 | 1.3 | 1.6 |
| 20 | 1.03 | 1.0 | 1.3 | 1.4 | 1.6 | 0.6 | 2.0 | 1.4 | 0.9 | 1.6 | 1.2 | 0.9 | 0.9 | 1.4 |
| 21 | 0.64 | 0.7 | 1.1 | 1.0 | 1.4 | 0.5 | 1.5 | 1.1 | 0.7 | 1.2 | 1.1 | 0.9 | 0.8 | 1.2 |
| 22 | 0.41 | 0.4 | 0.7 | 0.6 | 1.6 | 0.6 | 0.8 | 1.0 | 0.4 | 1.0 | 1.1 | 1.1 | 0.5 | 1.3 |
| 23 | 0.63 | 0.3 | 0.5 | 0.2 | 1.0 | 0.7 | 0.1 | 1.5 | 0.4 | 0.6 | 0.8 | 1.2 | 0.4 | 1.0 |
| 24 | 0.53 | 0.2 | 0.5 | 0.2 | 0.8 | 0.6 | 0.2 | 1.3 | 0.3 | 0.5 | 0.7 | 0.9 | 0.4 | 0.9 |
| 25 | 0.61 | 0.2 | 0.6 | 0.2 | 0.8 | 0.5 | 0.3 | 1.3 | 0.3 | 0.5 | 0.6 | 0.7 | 0.4 | 0.9 |
| 26 | 0.72 | 0.4 | 0.7 | 0.4 | 0.9 | 0.5 | 0.8 | 1.1 | 0.4 | 0.7 | 0.7 | 1.0 | 0.5 | 1.0 |
| 27 | 0.96 | 0.2 | 0.6 | 0.3 | 1.2 | 0.3 | 0.1 | 0.0 | 0.2 | 0.5 | 0.9 | 1.1 | 0.2 | 1.1 |
| 28 | 0.44 | 3.2 | 1.7 | 4.3 | 7.1 | 4.0 | 1.3 | 2.4 | 3.1 | 4.4 | 4.5 | 2.0 | 2.7 | 2.7 |
| 29 | 0.92 | 0.2 | 0.6 | 0.2 | 0.9 | 0.5 | 0.4 | 1.1 | 0.3 | 0.5 | 0.6 | 0.5 | 0.4 | 0.8 |
| 30 | 0.94 | 0.2 | 0.5 | 0.1 | 0.6 | 0.5 | 0.2 | 1.5 | 0.3 | 0.4 | 0.5 | 0.7 | 0.4 | 0.8 |
| 31 | 0.63 | 0.2 | 0.5 | 0.2 | 0.8 | 0.6 | 0.1 | 1.4 | 0.3 | 0.5 | 0.7 | 0.9 | 0.3 | 0.9 |
| 32 | 0.93 | 0.3 | 0.4 | 0.1 | 0.7 | 1.0 | 0.0 | 0.5 | 0.4 | 0.3 | 0.5 | 0.9 | 0.2 | 0.9 |
| 33 | 0.45 | 0.4 | 0.7 | 0.1 | 0.6 | 1.3 | 0.0 | 1.0 | 0.4 | 0.4 | 0.5 | 0.9 | 0.3 | 0.7 |
| 34 | 0.51 | 0.6 | 1.1 | 0.5 | 0.8 | 0.8 | 0.8 | 1.4 | 0.6 | 0.7 | 0.7 | 1.0 | 0.6 | 0.9 |
| 35 | 1.20 | 0.5 | 0.8 | 0.7 | 1.5 | 0.6 | 0.9 | 1.6 | 0.6 | 1.1 | 1.2 | 1.4 | 0.7 | 1.3 |
| 36 | 1.34 | 0.3 | 0.6 | 0.3 | 0.8 | 0.6 | 0.4 | 1.7 | 0.5 | 0.6 | 0.7 | 0.9 | 0.5 | 0.9 |
| 37 | 2.30 | 0.8 | 1.1 | 0.9 | 1.1 | 0.5 | 1.7 | 1.5 | 0.8 | 1.2 | 1.0 | 1.0 | 0.7 | 1.1 |
| 38 | 0.91 | 0.4 | 0.8 | 0.4 | 0.8 | 0.5 | 1.0 | 1.4 | 0.5 | 0.7 | 0.7 | 0.9 | 0.6 | 1.0 |
| 39 | 0.70 | 0.3 | 0.7 | 0.3 | 0.7 | 0.5 | 0.5 | 1.3 | 0.4 | 0.6 | 0.6 | 0.8 | 0.5 | 0.8 |
| 40 | 0.76 | 0.2 | 0.5 | 0.1 | 0.6 | 0.6 | 0.1 | 1.6 | 0.3 | 0.4 | 0.5 | 0.7 | 0.4 | 0.8 |
| 41 | 0.70 | 0.3 | 0.7 | 0.3 | 0.9 | 0.5 | 0.6 | 1.1 | 0.4 | 0.6 | 0.6 | 0.5 | 0.5 | 0.9 |
| 42 | 0.74 | 0.3 | 0.6 | 0.3 | 1.2 | 0.5 | 0.5 | 1.2 | 0.3 | 0.8 | 0.8 | 0.9 | 0.4 | 1.1 |
| 43 | 1.00 | 0.3 | 0.5 | 0.2 | 0.9 | 0.6 | 0.1 | 1.6 | 0.4 | 0.6 | 0.7 | 1.0 | 0.4 | 1.0 |
| 44 | 1.02 | 0.3 | 0.5 | 0.1 | 0.9 | 1.0 | 0.0 | 1.1 | 0.5 | 0.4 | 0.7 | 1.2 | 0.4 | 0.9 |
| 45 | 0.84 | 0.2 | 0.5 | 0.2 | 0.8 | 0.6 | 0.1 | 1.3 | 0.3 | 0.5 | 0.6 | 0.9 | 0.4 | 0.9 |
| 46 | 0.70 | 0.3 | 0.6 | 0.2 | 0.9 | 0.6 | 0.3 | 1.4 | 0.4 | 0.6 | 0.8 | 1.2 | 0.4 | 1.0 |
| 47 | 0.41 | 0.0 | 0.3 | 0.2 | 1.6 | 0.2 | 0.0 | 0.0 | 0.0 | 0.2 | 1.2 | 1.0 | 0.2 | 1.2 |
| 48 | 0.53 | 0.1 | 0.3 | 0.1 | 0.9 | 0.2 | 0.0 | 0.0 | 0.1 | 0.1 | 0.7 | 0.9 | 0.2 | 0.9 |
| 49 | 0.61 | 0.1 | 0.3 | 0.3 | 1.4 | 0.2 | 0.1 | 0.0 | 0.2 | 0.3 | 1.1 | 1.3 | 0.3 | 1.3 |
| 50 | 0.79 | 0.8 | 0.9 | 1.5 | 2.0 | 0.2 | 0.6 | 0.3 | 1.0 | 0.3 | 1.8 | 1.1 | 1.3 | 1.7 |
| 51 | 0.74 | 0.1 | 0.3 | 0.3 | 1.4 | 0.2 | 0.1 | 0.1 | 0.2 | 0.2 | 1.0 | 0.8 | 0.3 | 1.2 |
| 52 | 0.50 | 0.1 | 0.5 | 0.4 | 1.6 | 0.3 | 0.1 | 0.0 | 0.1 | 0.5 | 1.1 | 0.9 | 0.2 | 1.2 |
| 53 | 0.50 | 0.9 | 1.4 | 1.4 | 2.2 | 0.4 | 0.5 | 0.2 | 0.6 | 1.2 | 1.9 | 1.2 | 1.2 | 1.8 |
| 54 | 0.27 | 3.9 | 2.7 | 4.4 | 2.8 | 2.1 | 1.6 | 1.5 | 3.8 | 1.2 | 7.0 | 2.2 | 7.6 | 3.3 |
| 55 | 0.30 | 1.0 | 1.3 | 1.5 | 2.0 | 0.5 | 1.0 | 0.4 | 1.3 | 0.1 | 1.7 | 1.1 | 1.4 | 1.6 |
| 56 | 0.38 | 0.1 | 0.4 | 0.2 | 1.3 | 0.5 | 0.2 | 0.1 | 0.2 | 0.0 | 0.9 | 0.7 | 0.3 | 1.1 |
| 57 | 0.39 | 0.0 | 0.2 | 0.1 | 1.1 | 0.5 | 0.1 | 0.0 | 0.1 | 0.0 | 0.8 | 0.9 | 0.2 | 1.0 |
| 58 | 0.46 | 0.1 | 0.3 | 0.2 | 1.2 | 0.3 | 0.0 | 0.0 | 0.1 | 0.2 | 0.9 | 1.0 | 0.2 | 1.1 |
| 59 | 0.53 | 0.0 | 0.2 | 0.1 | 1.4 | 0.4 | 0.0 | 0.0 | 0.1 | 0.2 | 0.9 | 1.1 | 0.1 | 1.0 |
| 60 | 0.30 | 0.0 | 0.2 | 0.1 | 1.0 | 0.2 | 0.0 | 0.0 | 0.1 | 0.1 | 0.6 | 0.6 | 0.1 | 0.9 |
| 61 | 0.48 | 0.0 | 0.2 | 0.1 | 0.9 | 0.2 | 0.0 | 0.0 | 0.1 | 0.1 | 0.7 | 0.7 | 0.1 | 0.8 |
| 62 | 0.44 | 0.0 | 0.2 | 0.1 | 1.1 | 0.3 | 0.0 | 0.0 | 0.1 | 0.2 | 0.8 | 1.0 | 0.1 | 1.0 |
| 63 | 0.41 | 0.2 | 0.4 | 0.5 | 1.9 | 0.3 | 0.1 | 0.1 | 0.2 | 0.4 | 1.3 | 1.0 | 0.3 | 1.4 |
| 64 | 0.79 | 0.3 | 0.6 | 0.2 | 0.8 | 0.6 | 0.3 | 1.4 | 0.3 | 0.5 | 0.6 | 0.8 | 0.3 | 0.9 |
| 65 | 0.93 | 0.2 | 0.4 | 0.1 | 1.0 | 0.6 | 0.0 | 0.4 | 0.2 | 0.4 | 0.7 | 1.0 | 0.2 | 0.9 |
| 66 | 0.67 | 0.3 | 0.6 | 0.5 | 1.5 | 0.5 | 0.3 | 1.1 | 0.3 | 0.8 | 1.1 | 1.0 | 0.4 | 1.2 |
| 67 | 0.98 | 0.2 | 0.4 | 0.3 | 1.3 | 0.6 | 0.1 | 0.6 | 0.2 | 0.5 | 0.8 | 1.0 | 0.3 | 1.1 |
| 68 | 1.17 | 0.2 | 0.3 | 0.1 | 0.6 | 0.7 | 0.0 | 0.1 | 0.1 | 0.2 | 0.5 | 1.1 | 0.1 | 0.9 |
| 69 | 0.92 | 0.2 | 0.4 | 0.1 | 0.7 | 1.0 | 0.0 | 0.4 | 0.3 | 0.3 | 0.5 | 0.9 | 0.2 | 0.8 |
| 70 | 0.51 | 0.1 | 0.2 | 0.1 | 1.0 | 0.5 | 0.0 | 0.0 | 0.1 | 0.1 | 0.6 | 0.9 | 0.1 | 0.9 |
| 71 | 0.41 | 0.1 | 0.2 | 0.1 | 1.1 | 0.4 | 0.0 | 0.0 | 0.1 | 0.1 | 0.7 | 1.1 | 0.1 | 0.9 |
| 72 | 1.21 | 0.3 | 0.4 | 0.1 | 0.8 | 0.9 | 0.0 | 0.3 | 0.3 | 0.3 | 0.6 | 1.1 | 0.2 | 0.9 |
| 73 | 0.74 | 0.2 | 0.3 | 0.1 | 0.6 | 0.9 | 0.0 | 0.1 | 0.2 | 0.2 | 0.6 | 1.5 | 0.1 | 1.0 |
| 74 | 1.49 | 0.2 | 0.3 | 0.1 | 0.7 | 0.6 | 0.0 | 0.0 | 0.2 | 0.3 | 0.6 | 1.5 | 0.1 | 1.0 |
| 75 | 2.47 | 0.2 | 0.3 | 0.0 | 0.5 | 0.9 | 0.0 | 0.0 | 0.2 | 0.1 | 0.5 | 1.7 | 0.1 | 1.0 |
| 76 | 3.92 | 0.5 | 0.5 | 0.0 | 0.3 | 1.2 | 0.0 | 0.0 | 0.4 | 0.1 | 0.5 | 1.1 | 0.0 | 0.8 |
| 77 | 0.98 | 0.3 | 0.3 | 0.0 | 0.3 | 1.0 | 0.0 | 0.0 | 0.2 | 0.1 | 0.5 | 2.4 | 0.1 | 1.0 |
| 78 | 1.14 | 0.4 | 0.4 | 0.0 | 0.4 | 1.0 | 0.0 | 0.0 | 0.3 | 0.1 | 0.5 | 1.5 | 0.1 | 0.9 |
| 79 | 3.18 | 0.4 | 0.4 | 0.0 | 0.3 | 2.0 | 0.0 | 0.0 | 0.4 | 0.0 | 0.4 | 1.2 | 0.0 | 0.8 |
| 80 | 1.24 | 0.2 | 0.2 | 0.0 | 0.4 | 1.1 | 0.0 | 0.0 | 0.2 | 0.1 | 0.4 | 2.1 | 0.1 | 1.0 |
| 81 | 0.98 | 0.3 | 0.3 | 0.0 | 0.5 | 1.6 | 0.0 | 0.0 | 0.5 | 0.1 | 0.4 | 1.4 | 0.1 | 0.9 |
| 82 | 0.54 | 0.7 | 0.4 | 0.5 | 2.7 | 1.2 | 0.0 | 1.9 | 3.3 | 0.6 | 2.0 | 1.5 | 1.4 | 0.9 |
| 83 | 1.29 | 0.6 | 0.5 | 0.0 | 0.2 | 1.7 | 0.0 | 0.0 | 0.3 | 0.0 | 0.4 | 1.0 | 0.0 | 0.5 |
| 84 | 0.86 | 0.3 | 0.3 | 0.0 | 0.3 | 1.4 | 0.0 | 0.0 | 0.3 | 0.1 | 0.5 | 1.4 | 0.1 | 0.6 |
| 85 | 1.38 | 0.3 | 0.2 | 0.0 | 0.2 | 1.7 | 0.0 | 0.0 | 0.2 | 0.0 | 0.4 | 1.8 | 0.0 | 0.7 |
| 86 | 1.56 | 0.4 | 0.3 | 0.0 | 0.1 | 2.4 | 0.0 | 0.0 | 0.3 | 0.0 | 0.2 | 1.0 | 0.0 | 0.5 |
| 87 | 0.63 | 0.0 | 0.0 | 0.0 | 0.3 | 0.4 | 0.0 | 0.0 | 0.1 | 0.0 | 0.4 | 1.4 | 0.0 | 0.8 |
| 88 | 0.66 | 0.0 | 0.0 | 0.0 | 0.4 | 0.3 | 0.0 | 0.0 | 0.1 | 0.0 | 0.5 | 1.3 | 0.1 | 0.9 |
| 89 | 0.38 | 0.0 | 0.1 | 0.1 | 1.1 | 0.6 | 0.0 | 0.0 | 0.1 | 0.0 | 0.7 | 0.9 | 0.1 | 0.9 |
| 90 | 1.09 | 1.0 | 1.0 | 0.0 | 0.3 | 2.3 | 0.0 | 0.0 | 0.3 | 0.1 | 0.4 | 1.2 | 0.0 | 0.7 |
| 91 | 0.99 | 0.3 | 0.3 | 0.1 | 0.5 | 1.6 | 0.0 | 0.1 | 0.3 | 0.1 | 0.5 | 1.2 | 0.1 | 0.8 |
| 92 | 1.81 | 0.2 | 0.2 | 0.0 | 0.2 | 3.0 | 0.0 | 0.0 | 0.2 | 0.0 | 0.3 | 0.9 | 0.0 | 0.6 |
| 93 | 0.67 | 0.2 | 0.2 | 0.0 | 0.2 | 3.5 | 0.0 | 0.0 | 0.1 | 0.0 | 0.3 | 0.8 | 0.0 | 0.5 |
| 94 | 0.65 | 0.0 | 0.0 | 0.0 | 0.1 | 1.3 | 0.0 | 0.0 | 0.0 | 0.0 | 0.3 | 0.6 | 0.0 | 0.3 |
| 95 | 1.06 | 0.1 | 0.1 | 0.1 | 0.6 | 1.1 | 0.0 | 0.1 | 1.4 | 0.1 | 0.6 | 0.8 | 0.2 | 0.4 |
| 96 | 8.86 | 0.0 | 0.0 | 0.0 | 0.0 | 0.2 | 0.0 | 0.0 | 0.0 | 0.0 | 0.0 | 0.0 | 0.3 | 0.0 |
| 97 | 1.13 | 0.0 | 0.0 | 0.0 | 0.1 | 0.2 | 0.0 | 0.0 | 0.0 | 0.0 | 0.3 | 0.4 | 0.0 | 0.2 |
| 98 | 0.51 | 0.2 | 0.1 | 0.0 | 0.1 | 2.5 | 0.0 | 0.0 | 0.0 | 0.0 | 0.4 | 1.1 | 0.0 | 0.5 |
| 99 | 0.97 | 0.1 | 0.1 | 0.0 | 0.1 | 2.2 | 0.0 | 0.0 | 0.1 | 0.0 | 0.3 | 1.1 | 0.0 | 0.5 |
| 100 | 0.25 | 3.6 | 1.4 | 1.9 | 0.9 | 2.1 | 0.1 | 10.7 | 15.6 | 2.3 | 4.9 | 1.3 | 10.7 | 1.1 |
| Base | 100 | 5.00 | 5.00 | 4.98 | 5.00 | 5.00 | 5.02 | 5.00 | 5.00 | 5.03 | 5.03 | 5.01 | 5.00 | 5.00 |

B

Enrichments of top 5% ranked bases non-coding restricted

| State | Non coding region % | CADD 1.4 | CADD 1.0 | Eigen | Eigen-PC | DANN | GERP++ | phyloP | PhastCons | REMM | fitCons | FIRE | FATHMM-XF | CDTS | LINSIGHT | FunSeq2 |
|---|---|---|---|---|---|---|---|---|---|---|---|---|---|---|---|---|
| 1 | 0.13 | 16.9 | 12.2 | 17.3 | 5.3 | 6.6 | 15.2 | 17.1 | 17.7 | 15.9 | 6.4 | 0.8 | 19.0 | 4.0 | 17.9 | 6.6 |
| 2 | 0.83 | 13.6 | 10.5 | 14.9 | 2.2 | 1.4 | 12.7 | 11.9 | 14.3 | 11.9 | 3.6 | 0.5 | 17.6 | 2.0 | 14.6 | 4.4 |
| 3 | 0.33 | 9.5 | 6.8 | 11.1 | 2.0 | 0.6 | 7.8 | 4.2 | 8.4 | 7.7 | 4.0 | 0.9 | 15.0 | 2.2 | 11.4 | 3.8 |
| 4 | 0.54 | 10.4 | 7.9 | 11.7 | 2.9 | 1.4 | 9.5 | 6.9 | 9.6 | 8.9 | 3.4 | 1.1 | 12.3 | 2.4 | 11.7 | 4.4 |
| 5 | 1.80 | 9.6 | 7.2 | 11.3 | 2.7 | 0.9 | 9.9 | 6.6 | 9.0 | 8.6 | 2.6 | 0.8 | 8.8 | 1.8 | 10.8 | 4.1 |
| 6 | 3.95 | 3.0 | 2.7 | 3.6 | 1.8 | 0.7 | 4.7 | 3.3 | 2.9 | 3.4 | 1.5 | 0.8 | 2.1 | 1.4 | 3.9 | 2.8 |
| 7 | 0.91 | 2.8 | 2.7 | 4.2 | 3.1 | 1.1 | 3.8 | 2.2 | 2.0 | 3.3 | 3.3 | 2.2 | 5.3 | 2.8 | 4.2 | 3.4 |
| 8 | 0.79 | 3.5 | 3.3 | 4.4 | 2.1 | 0.5 | 3.4 | 1.3 | 2.5 | 3.1 | 2.4 | 1.1 | 3.5 | 1.7 | 4.5 | 2.5 |
| 9 | 0.99 | 1.3 | 1.7 | 1.6 | 1.7 | 0.4 | 1.5 | 0.4 | 0.9 | 1.7 | 1.5 | 0.8 | 0.7 | 1.3 | 1.7 | 1.8 |
| 10 | 0.95 | 0.9 | 1.3 | 1.2 | 1.7 | 0.4 | 0.8 | 0.2 | 0.6 | 1.2 | 1.5 | 0.9 | 0.8 | 1.4 | 1.4 | 1.5 |
| 11 | 0.83 | 0.7 | 1.1 | 0.9 | 1.5 | 0.5 | 1.6 | 1.2 | 0.6 | 1.3 | 1.3 | 0.9 | 0.7 | 1.2 | 1.0 | 1.6 |
| 12 | 0.83 | 0.9 | 1.2 | 1.0 | 1.2 | 0.6 | 1.7 | 1.4 | 0.8 | 1.3 | 1.0 | 0.6 | 0.8 | 1.0 | 1.1 | 1.4 |
| 13 | 1.37 | 1.1 | 1.4 | 1.3 | 1.2 | 0.5 | 2.3 | 1.7 | 1.0 | 1.5 | 1.1 | 0.6 | 0.9 | 1.1 | 1.5 | 1.7 |
| 14 | 1.03 | 1.1 | 1.3 | 1.5 | 2.0 | 0.6 | 2.1 | 1.5 | 0.9 | 1.9 | 1.7 | 1.3 | 0.8 | 1.5 | 1.4 | 2.3 |
| 15 | 0.85 | 0.9 | 1.3 | 1.2 | 1.9 | 0.4 | 0.8 | 0.2 | 0.5 | 1.3 | 1.6 | 0.9 | 0.6 | 1.4 | 1.3 | 1.7 |
| 16 | 0.79 | 0.2 | 0.7 | 0.3 | 1.2 | 0.4 | 0.4 | 0.1 | 0.2 | 0.6 | 1.0 | 0.7 | 0.3 | 1.0 | 0.3 | 1.0 |
| 17 | 2.36 | 0.7 | 1.1 | 0.8 | 1.1 | 0.6 | 2.0 | 1.9 | 0.8 | 1.2 | 0.9 | 0.7 | 0.8 | 1.0 | 0.9 | 1.6 |
| 18 | 1.04 | 0.2 | 0.7 | 0.3 | 1.3 | 0.3 | 0.2 | 0.1 | 0.2 | 0.6 | 1.1 | 0.7 | 0.3 | 1.1 | 0.4 | 1.0 |
| 19 | 1.17 | 1.6 | 1.9 | 2.0 | 1.9 | 0.4 | 1.1 | 0.4 | 1.1 | 1.8 | 1.8 | 0.9 | 1.3 | 1.5 | 2.4 | 1.8 |
| 20 | 1.13 | 1.1 | 1.4 | 1.5 | 1.6 | 0.7 | 2.1 | 1.4 | 1.0 | 1.6 | 1.3 | 0.8 | 0.9 | 1.3 | 1.5 | 1.8 |
| 21 | 0.71 | 0.8 | 1.2 | 1.0 | 1.4 | 0.5 | 1.6 | 1.2 | 0.7 | 1.3 | 1.3 | 0.8 | 0.8 | 1.1 | 1.1 | 1.6 |
| 22 | 0.45 | 0.5 | 0.8 | 0.6 | 1.5 | 0.7 | 0.8 | 1.0 | 0.4 | 1.1 | 1.2 | 1.0 | 0.5 | 1.3 | 0.4 | 1.4 |
| 23 | 0.69 | 0.3 | 0.5 | 0.2 | 1.0 | 0.7 | 0.2 | 1.6 | 0.4 | 0.6 | 0.9 | 1.1 | 0.4 | 1.0 | 0.1 | 0.9 |
| 24 | 0.59 | 0.3 | 0.5 | 0.2 | 0.8 | 0.6 | 0.2 | 1.4 | 0.3 | 0.6 | 0.8 | 0.8 | 0.4 | 0.8 | 0.1 | 0.8 |
| 25 | 0.68 | 0.3 | 0.6 | 0.2 | 0.8 | 0.6 | 0.4 | 1.3 | 0.3 | 0.6 | 0.7 | 0.7 | 0.4 | 0.8 | 0.1 | 0.9 |
| 26 | 0.80 | 0.4 | 0.8 | 0.4 | 0.8 | 0.6 | 0.9 | 1.2 | 0.5 | 0.7 | 0.8 | 0.9 | 0.5 | 0.9 | 0.3 | 1.2 |
| 27 | 1.05 | 0.2 | 0.6 | 0.3 | 1.2 | 0.4 | 0.2 | 0.1 | 0.2 | 0.5 | 1.1 | 1.0 | 0.3 | 1.1 | 0.2 | 0.9 |
| 28 | 0.44 | 2.5 | 1.3 | 4.4 | 7.5 | 3.3 | 1.2 | 2.2 | 2.8 | 4.1 | 4.5 | 1.8 | 2.7 | 2.7 | 2.9 | 3.2 |
| 29 | 1.01 | 0.3 | 0.6 | 0.2 | 0.9 | 0.6 | 0.4 | 1.2 | 0.3 | 0.6 | 0.7 | 0.5 | 0.4 | 0.8 | 0.2 | 0.8 |
| 30 | 1.03 | 0.3 | 0.6 | 0.1 | 0.6 | 0.6 | 0.2 | 1.5 | 0.3 | 0.5 | 0.6 | 0.6 | 0.4 | 0.7 | 0.1 | 0.7 |
| 31 | 0.70 | 0.3 | 0.5 | 0.2 | 0.8 | 0.7 | 0.1 | 1.5 | 0.3 | 0.6 | 0.8 | 0.9 | 0.4 | 0.9 | 0.1 | 0.7 |
| 32 | 1.00 | 0.3 | 0.5 | 0.1 | 0.7 | 1.1 | 0.0 | 0.6 | 0.5 | 0.3 | 0.6 | 0.8 | 0.2 | 0.8 | 0.0 | 0.4 |
| 33 | 0.49 | 0.5 | 0.8 | 0.1 | 0.6 | 1.5 | 0.0 | 1.1 | 0.5 | 0.4 | 0.6 | 0.9 | 0.3 | 0.7 | 0.1 | 0.4 |
| 34 | 0.56 | 0.7 | 1.2 | 0.5 | 0.8 | 0.9 | 0.9 | 1.4 | 0.6 | 0.8 | 0.8 | 0.9 | 0.6 | 0.9 | 0.5 | 1.0 |
| 35 | 1.33 | 0.6 | 0.9 | 0.7 | 1.5 | 0.7 | 0.9 | 1.6 | 0.7 | 1.2 | 1.4 | 1.3 | 0.7 | 1.3 | 0.5 | 1.7 |
| 36 | 1.47 | 0.4 | 0.7 | 0.3 | 0.7 | 0.6 | 0.4 | 1.7 | 0.5 | 0.7 | 0.7 | 0.9 | 0.6 | 0.9 | 0.1 | 1.0 |
| 37 | 2.54 | 0.9 | 1.2 | 0.9 | 1.1 | 0.6 | 1.8 | 1.6 | 0.9 | 1.3 | 1.1 | 1.0 | 0.8 | 1.1 | 0.9 | 1.7 |
| 38 | 1.00 | 0.5 | 0.9 | 0.4 | 0.8 | 0.6 | 1.1 | 1.4 | 0.6 | 0.8 | 0.8 | 0.9 | 0.6 | 1.0 | 0.4 | 1.2 |
| 39 | 0.76 | 0.4 | 0.7 | 0.3 | 0.7 | 0.6 | 0.5 | 1.3 | 0.5 | 0.6 | 0.7 | 0.7 | 0.5 | 0.8 | 0.2 | 0.9 |
| 40 | 0.83 | 0.3 | 0.5 | 0.1 | 0.6 | 0.7 | 0.1 | 1.7 | 0.4 | 0.4 | 0.6 | 0.7 | 0.4 | 0.8 | 0.0 | 0.6 |
| 41 | 0.78 | 0.4 | 0.7 | 0.3 | 0.9 | 0.6 | 0.6 | 1.2 | 0.4 | 0.7 | 0.7 | 0.5 | 0.5 | 0.8 | 0.3 | 0.9 |
| 42 | 0.82 | 0.4 | 0.7 | 0.4 | 1.2 | 0.6 | 0.5 | 1.2 | 0.4 | 0.8 | 1.0 | 0.9 | 0.5 | 1.0 | 0.2 | 1.1 |
| 43 | 1.10 | 0.3 | 0.6 | 0.2 | 0.9 | 0.7 | 0.2 | 1.7 | 0.4 | 0.7 | 0.8 | 0.9 | 0.4 | 1.0 | 0.1 | 0.8 |
| 44 | 1.12 | 0.4 | 0.5 | 0.1 | 0.9 | 1.1 | 0.0 | 1.2 | 0.5 | 0.4 | 0.8 | 1.1 | 0.4 | 0.9 | 0.0 | 0.5 |
| 45 | 0.93 | 0.3 | 0.6 | 0.2 | 0.8 | 0.7 | 0.2 | 1.4 | 0.3 | 0.5 | 0.7 | 0.8 | 0.4 | 0.9 | 0.1 | 0.7 |
| 46 | 0.77 | 0.3 | 0.6 | 0.2 | 0.9 | 0.7 | 0.4 | 1.5 | 0.4 | 0.7 | 0.9 | 1.1 | 0.5 | 0.9 | 0.1 | 1.0 |
| 47 | 0.45 | 0.1 | 0.3 | 0.2 | 1.6 | 0.2 | 0.0 | 0.0 | 0.0 | 0.3 | 1.3 | 0.9 | 0.3 | 1.2 | 0.2 | 1.0 |
| 48 | 0.59 | 0.1 | 0.3 | 0.1 | 0.9 | 0.3 | 0.0 | 0.0 | 0.1 | 0.2 | 0.8 | 0.8 | 0.2 | 0.9 | 0.1 | 0.6 |
| 49 | 0.67 | 0.1 | 0.4 | 0.3 | 1.4 | 0.2 | 0.1 | 0.1 | 0.2 | 0.3 | 1.3 | 1.2 | 0.3 | 1.2 | 0.2 | 1.0 |
| 50 | 0.87 | 0.9 | 0.9 | 1.5 | 2.0 | 0.2 | 0.7 | 0.3 | 1.1 | 0.4 | 1.9 | 1.0 | 1.3 | 1.6 | 1.8 | 1.6 |
| 51 | 0.81 | 0.1 | 0.3 | 0.3 | 1.4 | 0.2 | 0.1 | 0.1 | 0.2 | 0.2 | 1.1 | 0.8 | 0.3 | 1.1 | 0.2 | 0.9 |
| 52 | 0.55 | 0.2 | 0.5 | 0.4 | 1.5 | 0.3 | 0.1 | 0.1 | 0.1 | 0.5 | 1.2 | 0.8 | 0.3 | 1.2 | 0.3 | 1.0 |
| 53 | 0.55 | 1.0 | 1.4 | 1.5 | 2.2 | 0.3 | 0.5 | 0.2 | 0.6 | 1.3 | 2.0 | 1.1 | 1.2 | 1.7 | 1.8 | 1.6 |
| 54 | 0.23 | 2.7 | 2.2 | 3.8 | 2.6 | 0.4 | 1.6 | 1.3 | 3.3 | 1.0 | 3.7 | 1.6 | 8.8 | 2.5 | 4.8 | 2.1 |
| 55 | 0.33 | 1.0 | 1.3 | 1.5 | 1.9 | 0.4 | 1.1 | 0.4 | 1.3 | 0.1 | 1.8 | 1.0 | 1.5 | 1.5 | 1.9 | 1.7 |
| 56 | 0.42 | 0.1 | 0.4 | 0.2 | 1.3 | 0.5 | 0.2 | 0.1 | 0.2 | 0.0 | 1.1 | 0.7 | 0.3 | 1.0 | 0.2 | 0.9 |
| 57 | 0.43 | 0.0 | 0.3 | 0.1 | 1.1 | 0.6 | 0.1 | 0.0 | 0.2 | 0.0 | 1.0 | 0.9 | 0.3 | 1.0 | 0.1 | 0.7 |
| 58 | 0.51 | 0.1 | 0.3 | 0.2 | 1.2 | 0.3 | 0.0 | 0.0 | 0.1 | 0.2 | 1.0 | 0.9 | 0.2 | 1.0 | 0.1 | 0.7 |
| 59 | 0.58 | 0.1 | 0.2 | 0.1 | 1.3 | 0.5 | 0.0 | 0.0 | 0.1 | 0.2 | 1.0 | 1.0 | 0.1 | 1.0 | 0.0 | 0.6 |
| 60 | 0.34 | 0.0 | 0.2 | 0.1 | 1.0 | 0.3 | 0.0 | 0.0 | 0.1 | 0.1 | 0.7 | 0.6 | 0.1 | 0.8 | 0.0 | 0.5 |
| 61 | 0.53 | 0.0 | 0.2 | 0.1 | 0.9 | 0.3 | 0.0 | 0.0 | 0.1 | 0.2 | 0.7 | 0.6 | 0.1 | 0.8 | 0.0 | 0.5 |
| 62 | 0.49 | 0.1 | 0.2 | 0.1 | 1.1 | 0.3 | 0.0 | 0.0 | 0.1 | 0.2 | 0.9 | 0.9 | 0.1 | 0.9 | 0.0 | 0.7 |
| 63 | 0.45 | 0.2 | 0.4 | 0.5 | 1.8 | 0.3 | 0.1 | 0.1 | 0.2 | 0.4 | 1.5 | 0.9 | 0.3 | 1.3 | 0.4 | 1.1 |
| 64 | 0.87 | 0.3 | 0.6 | 0.2 | 0.8 | 0.7 | 0.3 | 1.4 | 0.3 | 0.6 | 0.7 | 0.8 | 0.4 | 0.9 | 0.1 | 0.8 |
| 65 | 1.03 | 0.2 | 0.4 | 0.2 | 1.0 | 0.7 | 0.0 | 0.5 | 0.2 | 0.5 | 0.8 | 1.0 | 0.2 | 0.9 | 0.1 | 0.6 |
| 66 | 0.73 | 0.4 | 0.6 | 0.5 | 1.5 | 0.6 | 0.4 | 1.1 | 0.3 | 0.9 | 1.2 | 1.0 | 0.4 | 1.1 | 0.4 | 1.2 |
| 67 | 1.08 | 0.2 | 0.4 | 0.3 | 1.3 | 0.6 | 0.1 | 0.7 | 0.2 | 0.5 | 1.0 | 0.9 | 0.3 | 1.0 | 0.2 | 0.7 |
| 68 | 1.30 | 0.2 | 0.3 | 0.1 | 0.6 | 0.7 | 0.0 | 0.1 | 0.2 | 0.3 | 0.6 | 1.0 | 0.2 | 0.8 | 0.0 | 0.4 |
| 69 | 1.01 | 0.3 | 0.4 | 0.1 | 0.7 | 1.1 | 0.0 | 0.4 | 0.4 | 0.3 | 0.6 | 0.8 | 0.2 | 0.8 | 0.0 | 0.3 |
| 70 | 0.55 | 0.1 | 0.2 | 0.1 | 1.0 | 0.6 | 0.0 | 0.0 | 0.1 | 0.1 | 0.7 | 0.8 | 0.1 | 0.9 | 0.0 | 0.4 |
| 71 | 0.45 | 0.1 | 0.2 | 0.1 | 1.1 | 0.5 | 0.0 | 0.0 | 0.1 | 0.2 | 0.8 | 1.0 | 0.1 | 0.9 | 0.0 | 0.5 |
| 72 | 1.31 | 0.3 | 0.4 | 0.1 | 0.8 | 1.1 | 0.0 | 0.4 | 0.4 | 0.3 | 0.7 | 1.1 | 0.2 | 0.9 | 0.0 | 0.4 |
| 73 | 0.81 | 0.2 | 0.3 | 0.1 | 0.6 | 1.0 | 0.0 | 0.1 | 0.3 | 0.2 | 0.7 | 1.4 | 0.1 | 0.9 | 0.0 | 0.4 |
| 74 | 1.64 | 0.3 | 0.3 | 0.1 | 0.7 | 0.7 | 0.0 | 0.0 | 0.2 | 0.3 | 0.8 | 1.4 | 0.1 | 1.0 | 0.1 | 0.5 |
| 75 | 2.73 | 0.3 | 0.4 | 0.0 | 0.5 | 1.0 | 0.0 | 0.0 | 0.3 | 0.2 | 0.6 | 1.6 | 0.1 | 1.0 | 0.0 | 0.4 |
| 76 | 4.33 | 0.6 | 0.5 | 0.0 | 0.3 | 1.3 | 0.0 | 0.0 | 0.5 | 0.1 | 0.5 | 1.0 | 0.1 | 0.8 | 0.0 | 0.3 |
| 77 | 1.08 | 0.4 | 0.3 | 0.0 | 0.3 | 1.1 | 0.0 | 0.0 | 0.3 | 0.1 | 0.6 | 2.2 | 0.1 | 1.0 | 0.0 | 0.3 |
| 78 | 1.26 | 0.5 | 0.5 | 0.0 | 0.4 | 1.1 | 0.0 | 0.0 | 0.4 | 0.1 | 0.5 | 1.4 | 0.1 | 0.9 | 0.0 | 0.3 |
| 79 | 3.51 | 0.6 | 0.4 | 0.0 | 0.3 | 2.1 | 0.0 | 0.0 | 0.5 | 0.0 | 0.5 | 1.1 | 0.0 | 0.7 | 0.0 | 0.2 |
| 80 | 1.37 | 0.2 | 0.2 | 0.0 | 0.4 | 1.2 | 0.0 | 0.0 | 0.3 | 0.1 | 0.5 | 1.9 | 0.1 | 1.0 | 0.0 | 0.3 |
| 81 | 1.08 | 0.4 | 0.3 | 0.0 | 0.5 | 1.7 | 0.0 | 0.0 | 0.6 | 0.1 | 0.5 | 1.3 | 0.1 | 0.9 | 0.0 | 0.2 |
| 82 | 0.57 | 0.7 | 0.4 | 0.5 | 2.8 | 1.2 | 0.0 | 1.9 | 3.3 | 0.7 | 2.2 | 1.4 | 1.5 | 0.9 | 0.7 | 0.7 |
| 83 | 1.38 | 0.7 | 0.6 | 0.0 | 0.2 | 1.8 | 0.0 | 0.0 | 0.4 | 0.0 | 0.4 | 0.9 | 0.0 | 0.5 | 0.0 | 0.1 |
| 84 | 0.91 | 0.4 | 0.3 | 0.0 | 0.3 | 1.6 | 0.0 | 0.0 | 0.4 | 0.1 | 0.6 | 1.3 | 0.1 | 0.6 | 0.0 | 0.2 |
| 85 | 1.46 | 0.4 | 0.3 | 0.0 | 0.2 | 1.9 | 0.0 | 0.0 | 0.4 | 0.0 | 0.5 | 1.7 | 0.0 | 0.7 | 0.0 | 0.1 |
| 86 | 1.73 | 0.5 | 0.3 | 0.0 | 0.1 | 2.5 | 0.0 | 0.0 | 0.5 | 0.0 | 0.3 | 1.0 | 0.0 | 0.5 | 0.0 | 0.1 |
| 87 | 0.69 | 0.0 | 0.0 | 0.0 | 0.3 | 0.4 | 0.0 | 0.0 | 0.1 | 0.0 | 0.5 | 1.3 | 0.0 | 0.8 | 0.0 | 0.2 |
| 88 | 0.73 | 0.0 | 0.0 | 0.0 | 0.4 | 0.3 | 0.0 | 0.0 | 0.1 | 0.0 | 0.6 | 1.2 | 0.1 | 0.9 | 0.0 | 0.3 |
| 89 | 0.41 | 0.0 | 0.1 | 0.1 | 1.1 | 0.6 | 0.0 | 0.0 | 0.1 | 0.0 | 0.9 | 0.8 | 0.1 | 0.9 | 0.0 | 0.5 |
| 90 | 1.19 | 1.2 | 1.0 | 0.0 | 0.3 | 2.4 | 0.0 | 0.0 | 0.4 | 0.1 | 0.4 | 1.1 | 0.1 | 0.7 | 0.1 | 0.2 |
| 91 | 1.07 | 0.3 | 0.4 | 0.1 | 0.5 | 1.8 | 0.0 | 0.1 | 0.3 | 0.1 | 0.6 | 1.2 | 0.1 | 0.8 | 0.0 | 0.2 |
| 92 | 2.00 | 0.3 | 0.2 | 0.0 | 0.2 | 3.1 | 0.0 | 0.0 | 0.3 | 0.0 | 0.3 | 0.9 | 0.0 | 0.6 | 0.0 | 0.1 |
| 93 | 0.74 | 0.3 | 0.2 | 0.0 | 0.2 | 3.6 | 0.0 | 0.0 | 0.2 | 0.0 | 0.4 | 0.8 | 0.0 | 0.5 | 0.0 | 0.1 |
| 94 | 0.72 | 0.0 | 0.0 | 0.0 | 0.1 | 1.4 | 0.0 | 0.0 | 0.0 | 0.0 | 0.4 | 0.6 | 0.0 | 0.3 | 0.0 | 0.1 |
| 95 | 1.08 | 0.2 | 0.1 | 0.1 | 0.6 | 1.2 | 0.0 | 0.1 | 1.4 | 0.1 | 0.7 | 0.8 | 0.2 | 0.4 | 0.1 | 0.2 |
| 96 | 1.33 | 0.0 | 0.0 | 0.0 | 0.2 | 1.4 | 0.0 | 0.0 | 0.0 | 0.0 | 0.4 | 0.4 | 2.1 | 0.1 | 0.0 | 0.0 |
| 97 | 0.96 | 0.0 | 0.0 | 0.0 | 0.1 | 0.3 | 0.0 | 0.0 | 0.0 | 0.0 | 0.4 | 0.5 | 0.0 | 0.2 | 0.0 | 0.1 |
| 98 | 0.56 | 0.2 | 0.1 | 0.0 | 0.1 | 2.6 | 0.0 | 0.0 | 0.1 | 0.0 | 0.4 | 1.1 | 0.0 | 0.5 | 0.0 | 0.1 |
| 99 | 1.05 | 0.2 | 0.1 | 0.0 | 0.1 | 2.3 | 0.0 | 0.0 | 0.2 | 0.0 | 0.3 | 1.0 | 0.0 | 0.5 | 0.0 | 0.1 |
| 100 | 0.26 | 3.6 | 1.2 | 2.1 | 0.9 | 1.8 | 0.1 | 10.8 | 15.8 | 2.5 | 4.6 | 1.3 | 11.4 | 1.1 | 6.2 | 0.5 |
| Base | 90.44 | 4.96 | 4.97 | 4.97 | 4.96 | 4.95 | 4.98 | 4.97 | 4.96 | 5.00 | 5.02 | 4.97 | 4.96 | 4.97 | 4.97 | 4.96 |

**Figure S26: Enrichment of all conservation states for top 5% of bases prioritized by variant prioritization scores.** The figure displays for top 5% prioritized bases **(A)** genome-wide and **(B)** in non-coding regions by variant prioritization scores the same enrichments as shown in **Figure S24A** and **S24B** except here enrichments for all conservation states are shown and also included is the original version of the CADD score in addition to v1.4. Coloring of enrichments is based on their value in a column specific manner. The second columns gives the percentage of the background region used to compute the enrichments falling in each state, which is for **(A)** the whole genome and for **(B)** bases scored by both LINSIGHT and FunSeq2. The last line in both heatmaps gives the actual percentage of the background set covered by each set of prioritized bases, which can differ from 5.00% because of how ties were handled.

## A

#### Enrichments of top 10% ranked bases genome-wide

| State | Genome % | CADD 1.4 | CADD 1.0 | Eigen | Eigen-PC | DANN | GERP++ | phyloP | PhastCons | REMM | fitCons | FIRE | FATHMM-XF | CDTS |
|---|---|---|---|---|---|---|---|---|---|---|---|---|---|---|
| 1 | 0.60 | 9.7 | 8.9 | 8.3 | 8.0 | 8.5 | 8.8 | 8.9 | 9.4 | 9.0 | 7.8 | 2.7 | 8.9 | 4.3 |
| 2 | 0.84 | 7.8 | 6.9 | 8.5 | 4.9 | 2.7 | 7.3 | 7.2 | 7.6 | 7.0 | 2.9 | 0.9 | 9.1 | 2.1 |
| 3 | 0.46 | 6.8 | 5.4 | 7.0 | 3.5 | 2.8 | 5.0 | 3.4 | 5.2 | 5.8 | 4.6 | 1.9 | 7.4 | 2.8 |
| 4 | 0.60 | 6.9 | 6.0 | 7.4 | 4.0 | 2.8 | 6.2 | 5.1 | 5.8 | 6.1 | 3.4 | 1.6 | 7.1 | 2.3 |
| 5 | 1.66 | 6.1 | 5.3 | 7.8 | 3.4 | 1.4 | 6.4 | 5.0 | 5.2 | 5.6 | 1.9 | 0.9 | 5.9 | 1.8 |
| 6 | 3.59 | 2.5 | 2.5 | 3.9 | 1.9 | 1.0 | 4.0 | 3.0 | 2.0 | 2.9 | 1.3 | 0.9 | 2.4 | 1.4 |
| 7 | 0.85 | 2.6 | 2.8 | 4.3 | 2.5 | 1.3 | 3.2 | 2.1 | 1.4 | 2.8 | 2.8 | 2.3 | 4.0 | 2.4 |
| 8 | 0.73 | 2.9 | 3.0 | 4.8 | 2.1 | 0.8 | 2.9 | 1.4 | 1.7 | 2.8 | 1.9 | 1.2 | 3.4 | 1.6 |
| 9 | 0.90 | 1.4 | 1.7 | 2.2 | 1.6 | 0.7 | 1.8 | 0.7 | 0.7 | 1.9 | 1.3 | 0.9 | 1.5 | 1.4 |
| 10 | 0.86 | 1.0 | 1.4 | 1.6 | 1.6 | 0.6 | 1.0 | 0.4 | 0.5 | 1.6 | 1.3 | 1.0 | 1.4 | 1.4 |
| 11 | 0.75 | 0.9 | 1.2 | 1.4 | 1.4 | 0.8 | 1.9 | 1.4 | 0.6 | 1.6 | 1.2 | 1.0 | 1.2 | 1.3 |
| 12 | 0.75 | 1.0 | 1.3 | 1.4 | 1.2 | 0.9 | 2.1 | 1.6 | 0.7 | 1.5 | 1.0 | 0.7 | 1.3 | 1.1 |
| 13 | 1.24 | 1.2 | 1.5 | 1.9 | 1.3 | 0.9 | 2.6 | 1.8 | 0.8 | 1.7 | 1.0 | 0.8 | 1.4 | 1.2 |
| 14 | 0.94 | 1.3 | 1.5 | 2.1 | 1.8 | 0.9 | 2.4 | 1.6 | 0.8 | 2.1 | 1.5 | 1.3 | 1.3 | 1.5 |
| 15 | 0.77 | 1.0 | 1.4 | 1.7 | 1.7 | 0.6 | 1.1 | 0.4 | 0.4 | 1.7 | 1.4 | 1.0 | 1.4 | 1.4 |
| 16 | 0.72 | 0.5 | 0.8 | 0.6 | 1.1 | 0.6 | 0.8 | 0.3 | 0.2 | 1.0 | 1.0 | 0.8 | 0.8 | 1.1 |
| 17 | 2.14 | 1.0 | 1.2 | 1.4 | 1.1 | 0.9 | 2.6 | 2.1 | 0.7 | 1.5 | 1.0 | 0.8 | 1.4 | 1.1 |
| 18 | 0.94 | 0.4 | 0.8 | 0.6 | 1.2 | 0.6 | 0.5 | 0.2 | 0.2 | 1.0 | 1.1 | 0.8 | 0.8 | 1.2 |
| 19 | 1.07 | 1.6 | 1.9 | 2.5 | 1.8 | 0.7 | 1.3 | 0.6 | 0.8 | 2.0 | 1.5 | 1.0 | 1.9 | 1.5 |
| 20 | 1.03 | 1.2 | 1.5 | 1.9 | 1.5 | 0.9 | 2.3 | 1.5 | 0.8 | 1.8 | 1.2 | 0.9 | 1.4 | 1.3 |
| 21 | 0.64 | 1.0 | 1.2 | 1.5 | 1.4 | 0.8 | 1.9 | 1.4 | 0.6 | 1.6 | 1.2 | 0.9 | 1.3 | 1.2 |
| 22 | 0.41 | 0.7 | 0.9 | 0.9 | 1.4 | 0.9 | 1.5 | 1.2 | 0.4 | 1.4 | 1.2 | 1.1 | 0.9 | 1.3 |
| 23 | 0.63 | 0.6 | 0.7 | 0.4 | 1.0 | 0.9 | 0.7 | 1.4 | 0.5 | 1.0 | 1.1 | 1.2 | 0.8 | 1.1 |
| 24 | 0.53 | 0.5 | 0.6 | 0.4 | 0.9 | 0.9 | 0.8 | 1.3 | 0.4 | 0.9 | 0.9 | 0.9 | 0.8 | 1.0 |
| 25 | 0.61 | 0.5 | 0.7 | 0.4 | 0.8 | 0.8 | 1.1 | 1.2 | 0.4 | 0.9 | 0.8 | 0.8 | 0.9 | 1.0 |
| 26 | 0.72 | 0.7 | 0.9 | 0.7 | 0.9 | 0.8 | 1.5 | 1.3 | 0.5 | 1.1 | 1.0 | 1.0 | 1.1 | 1.1 |
| 27 | 0.96 | 0.4 | 0.7 | 0.5 | 1.1 | 0.6 | 0.4 | 0.2 | 0.2 | 0.9 | 1.2 | 1.1 | 0.7 | 1.2 |
| 28 | 0.44 | 2.6 | 2.0 | 3.3 | 4.3 | 2.7 | 1.6 | 2.0 | 2.0 | 3.2 | 3.1 | 2.1 | 2.3 | 1.8 |
| 29 | 0.92 | 0.5 | 0.7 | 0.4 | 0.9 | 0.8 | 1.1 | 1.1 | 0.3 | 0.9 | 0.7 | 0.6 | 0.9 | 0.9 |
| 30 | 0.94 | 0.5 | 0.6 | 0.3 | 0.7 | 0.8 | 0.9 | 1.2 | 0.4 | 0.8 | 0.7 | 0.7 | 0.8 | 0.9 |
| 31 | 0.63 | 0.5 | 0.6 | 0.3 | 0.9 | 0.9 | 0.6 | 1.3 | 0.4 | 0.9 | 0.9 | 1.0 | 0.7 | 1.0 |
| 32 | 0.93 | 0.6 | 0.6 | 0.2 | 0.7 | 1.1 | 0.1 | 2.0 | 0.6 | 0.6 | 0.8 | 1.0 | 0.5 | 0.9 |
| 33 | 0.45 | 0.8 | 1.0 | 0.3 | 0.6 | 1.5 | 0.3 | 1.6 | 0.6 | 0.6 | 0.7 | 1.0 | 0.5 | 0.8 |
| 34 | 0.51 | 1.0 | 1.3 | 0.9 | 0.9 | 1.1 | 1.5 | 1.4 | 0.6 | 1.1 | 0.9 | 1.1 | 1.1 | 1.0 |
| 35 | 1.20 | 0.8 | 1.0 | 1.1 | 1.4 | 0.9 | 1.7 | 1.6 | 0.7 | 1.5 | 1.4 | 1.4 | 1.2 | 1.3 |
| 36 | 1.34 | 0.6 | 0.8 | 0.5 | 0.8 | 0.9 | 1.3 | 1.5 | 0.6 | 1.0 | 0.9 | 1.0 | 1.1 | 1.0 |
| 37 | 2.30 | 1.1 | 1.3 | 1.5 | 1.2 | 0.9 | 2.3 | 1.8 | 0.8 | 1.6 | 1.2 | 1.1 | 1.3 | 1.2 |
| 38 | 0.91 | 0.8 | 1.0 | 0.8 | 0.9 | 0.8 | 1.8 | 1.6 | 0.6 | 1.2 | 1.0 | 1.0 | 1.1 | 1.1 |
| 39 | 0.70 | 0.6 | 0.8 | 0.5 | 0.8 | 0.8 | 1.3 | 1.3 | 0.5 | 1.0 | 0.8 | 0.8 | 1.0 | 1.0 |
| 40 | 0.76 | 0.5 | 0.6 | 0.3 | 0.7 | 0.9 | 0.7 | 1.3 | 0.4 | 0.8 | 0.7 | 0.8 | 0.8 | 0.9 |
| 41 | 0.70 | 0.6 | 0.8 | 0.6 | 0.9 | 0.8 | 1.4 | 1.3 | 0.4 | 1.0 | 0.7 | 0.6 | 1.0 | 1.0 |
| 42 | 0.74 | 0.6 | 0.8 | 0.6 | 1.2 | 0.9 | 1.3 | 1.3 | 0.4 | 1.2 | 1.0 | 1.0 | 0.9 | 1.1 |
| 43 | 1.00 | 0.6 | 0.7 | 0.4 | 1.0 | 0.9 | 0.8 | 1.4 | 0.5 | 1.0 | 1.0 | 1.0 | 0.8 | 1.1 |
| 44 | 1.02 | 0.7 | 0.7 | 0.3 | 0.8 | 1.2 | 0.2 | 2.0 | 0.6 | 0.7 | 0.9 | 1.2 | 0.7 | 1.0 |
| 45 | 0.84 | 0.5 | 0.7 | 0.3 | 0.8 | 0.9 | 0.7 | 1.4 | 0.4 | 0.9 | 0.8 | 0.9 | 0.8 | 1.0 |
| 46 | 0.70 | 0.6 | 0.8 | 0.5 | 0.9 | 0.9 | 1.1 | 1.3 | 0.5 | 1.0 | 1.0 | 1.2 | 0.9 | 1.1 |
| 47 | 0.41 | 0.2 | 0.4 | 0.3 | 1.4 | 0.4 | 0.1 | 0.0 | 0.0 | 0.5 | 1.3 | 1.0 | 0.7 | 1.2 |
| 48 | 0.53 | 0.2 | 0.4 | 0.2 | 0.9 | 0.4 | 0.1 | 0.1 | 0.2 | 0.3 | 0.9 | 0.9 | 0.6 | 1.0 |
| 49 | 0.61 | 0.3 | 0.5 | 0.5 | 1.3 | 0.4 | 0.3 | 0.2 | 0.2 | 0.5 | 1.3 | 1.2 | 0.7 | 1.3 |
| 50 | 0.79 | 0.9 | 0.9 | 1.6 | 1.8 | 0.3 | 0.9 | 0.5 | 0.8 | 0.5 | 1.5 | 1.1 | 1.8 | 1.5 |
| 51 | 0.74 | 0.2 | 0.4 | 0.4 | 1.3 | 0.3 | 0.3 | 0.2 | 0.2 | 0.3 | 1.1 | 0.9 | 0.7 | 1.2 |
| 52 | 0.50 | 0.3 | 0.6 | 0.5 | 1.4 | 0.5 | 0.3 | 0.1 | 0.1 | 0.9 | 1.1 | 0.9 | 0.8 | 1.2 |
| 53 | 0.50 | 1.1 | 1.4 | 1.9 | 1.9 | 0.6 | 0.7 | 0.3 | 0.5 | 1.6 | 1.7 | 1.2 | 1.8 | 1.6 |
| 54 | 0.27 | 3.0 | 2.5 | 3.6 | 2.5 | 1.4 | 1.4 | 1.3 | 2.4 | 1.2 | 4.0 | 2.1 | 5.2 | 2.5 |
| 55 | 0.30 | 0.9 | 1.3 | 1.5 | 1.7 | 0.5 | 1.1 | 0.6 | 0.9 | 0.1 | 1.5 | 1.1 | 1.7 | 1.5 |
| 56 | 0.38 | 0.2 | 0.7 | 0.3 | 1.2 | 0.5 | 0.4 | 0.2 | 0.2 | 0.0 | 1.0 | 0.8 | 0.6 | 1.1 |
| 57 | 0.39 | 0.1 | 0.6 | 0.2 | 1.0 | 0.5 | 0.2 | 0.1 | 0.2 | 0.0 | 1.0 | 1.0 | 0.5 | 1.1 |
| 58 | 0.46 | 0.2 | 0.4 | 0.2 | 1.1 | 0.4 | 0.1 | 0.1 | 0.1 | 0.4 | 1.0 | 1.0 | 0.4 | 1.1 |
| 59 | 0.53 | 0.2 | 0.3 | 0.2 | 1.2 | 0.6 | 0.0 | 0.0 | 0.2 | 0.4 | 1.1 | 1.1 | 0.3 | 1.0 |
| 60 | 0.30 | 0.1 | 0.3 | 0.1 | 0.9 | 0.4 | 0.0 | 0.0 | 0.1 | 0.3 | 0.8 | 0.7 | 0.4 | 1.0 |
| 61 | 0.48 | 0.1 | 0.3 | 0.1 | 0.9 | 0.4 | 0.1 | 0.1 | 0.1 | 0.3 | 0.8 | 0.7 | 0.3 | 0.9 |
| 62 | 0.44 | 0.1 | 0.3 | 0.2 | 1.0 | 0.4 | 0.1 | 0.1 | 0.1 | 0.3 | 1.0 | 1.0 | 0.4 | 1.0 |
| 63 | 0.41 | 0.3 | 0.5 | 0.6 | 1.5 | 0.4 | 0.3 | 0.2 | 0.2 | 0.6 | 1.3 | 1.0 | 0.7 | 1.3 |
| 64 | 0.79 | 0.5 | 0.7 | 0.4 | 0.9 | 0.9 | 0.9 | 1.2 | 0.4 | 0.9 | 0.8 | 0.9 | 0.8 | 1.0 |
| 65 | 0.93 | 0.4 | 0.5 | 0.2 | 1.0 | 0.8 | 0.2 | 1.3 | 0.3 | 0.7 | 0.9 | 1.0 | 0.4 | 1.0 |
| 66 | 0.67 | 0.6 | 0.7 | 0.7 | 1.4 | 0.8 | 0.9 | 0.9 | 0.3 | 1.2 | 1.1 | 1.1 | 0.8 | 1.2 |
| 67 | 0.98 | 0.4 | 0.5 | 0.4 | 1.1 | 0.8 | 0.3 | 1.2 | 0.2 | 0.8 | 1.0 | 1.0 | 0.5 | 1.1 |
| 68 | 1.17 | 0.4 | 0.4 | 0.1 | 0.6 | 0.9 | 0.1 | 1.2 | 0.3 | 0.5 | 0.8 | 1.0 | 0.3 | 0.9 |
| 69 | 0.92 | 0.5 | 0.6 | 0.2 | 0.7 | 1.1 | 0.1 | 1.8 | 0.5 | 0.6 | 0.7 | 0.9 | 0.4 | 0.9 |
| 70 | 0.51 | 0.2 | 0.3 | 0.1 | 0.9 | 0.6 | 0.0 | 0.0 | 0.2 | 0.3 | 0.8 | 0.9 | 0.1 | 0.9 |
| 71 | 0.41 | 0.2 | 0.3 | 0.1 | 1.0 | 0.6 | 0.0 | 0.0 | 0.1 | 0.3 | 0.9 | 1.1 | 0.2 | 1.0 |
| 72 | 1.21 | 0.6 | 0.6 | 0.2 | 0.8 | 1.1 | 0.1 | 1.8 | 0.5 | 0.6 | 0.8 | 1.1 | 0.3 | 0.9 |
| 73 | 0.74 | 0.4 | 0.5 | 0.1 | 0.7 | 1.0 | 0.0 | 0.9 | 0.4 | 0.4 | 0.9 | 1.4 | 0.2 | 1.0 |
| 74 | 1.49 | 0.4 | 0.3 | 0.2 | 0.7 | 0.8 | 0.1 | 0.2 | 0.4 | 0.5 | 1.0 | 1.4 | 0.2 | 1.1 |
| 75 | 2.47 | 0.5 | 0.5 | 0.1 | 0.5 | 1.0 | 0.0 | 0.3 | 0.4 | 0.3 | 1.0 | 1.5 | 0.2 | 1.1 |
| 76 | 3.92 | 0.8 | 0.6 | 0.1 | 0.4 | 1.2 | 0.0 | 0.0 | 1.1 | 0.2 | 0.7 | 1.0 | 0.1 | 0.9 |
| 77 | 0.98 | 0.5 | 0.4 | 0.1 | 0.4 | 1.1 | 0.0 | 0.1 | 0.7 | 0.2 | 1.1 | 2.0 | 0.1 | 1.0 |
| 78 | 1.14 | 0.7 | 0.5 | 0.1 | 0.4 | 1.0 | 0.0 | 0.0 | 0.9 | 0.3 | 0.8 | 1.4 | 0.1 | 0.9 |
| 79 | 3.18 | 0.9 | 0.7 | 0.1 | 0.4 | 1.6 | 0.0 | 0.0 | 1.5 | 0.1 | 0.7 | 1.1 | 0.1 | 0.8 |
| 80 | 1.24 | 0.4 | 0.4 | 0.1 | 0.4 | 1.1 | 0.0 | 0.2 | 0.4 | 0.2 | 0.9 | 1.8 | 0.1 | 1.0 |
| 81 | 0.98 | 0.7 | 0.5 | 0.1 | 0.5 | 1.4 | 0.0 | 0.2 | 1.0 | 0.1 | 0.8 | 1.3 | 0.1 | 0.9 |
| 82 | 0.54 | 1.1 | 0.6 | 0.7 | 2.1 | 1.1 | 0.1 | 1.9 | 2.4 | 0.8 | 1.7 | 1.5 | 1.3 | 0.8 |
| 83 | 1.29 | 1.0 | 0.7 | 0.1 | 0.3 | 1.4 | 0.0 | 0.0 | 1.1 | 0.1 | 0.5 | 0.9 | 0.0 | 0.6 |
| 84 | 0.86 | 0.6 | 0.5 | 0.1 | 0.4 | 1.3 | 0.0 | 0.3 | 0.6 | 0.2 | 0.7 | 1.3 | 0.1 | 0.6 |
| 85 | 1.38 | 0.7 | 0.5 | 0.1 | 0.2 | 1.4 | 0.0 | 0.0 | 1.3 | 0.0 | 0.7 | 1.6 | 0.0 | 0.7 |
| 86 | 1.56 | 0.9 | 0.6 | 0.1 | 0.2 | 1.8 | 0.0 | 0.0 | 1.6 | 0.0 | 0.5 | 1.0 | 0.0 | 0.6 |
| 87 | 0.63 | 0.0 | 0.1 | 0.0 | 0.3 | 0.3 | 0.0 | 0.0 | 0.2 | 0.0 | 0.8 | 1.3 | 0.1 | 0.9 |
| 88 | 0.66 | 0.0 | 0.1 | 0.0 | 0.4 | 0.3 | 0.0 | 0.0 | 0.2 | 0.0 | 0.9 | 1.2 | 0.1 | 0.9 |
| 89 | 0.38 | 0.1 | 0.4 | 0.1 | 1.0 | 0.5 | 0.1 | 0.0 | 0.1 | 0.0 | 0.9 | 0.9 | 0.2 | 0.9 |
| 90 | 1.09 | 1.4 | 1.1 | 0.2 | 0.3 | 1.8 | 0.0 | 0.0 | 1.0 | 0.2 | 0.6 | 1.1 | 0.1 | 0.8 |
| 91 | 0.99 | 0.6 | 0.6 | 0.1 | 0.5 | 1.5 | 0.0 | 0.5 | 0.5 | 0.2 | 0.7 | 1.2 | 0.1 | 0.8 |
| 92 | 1.81 | 0.8 | 0.5 | 0.0 | 0.2 | 2.1 | 0.0 | 0.0 | 1.7 | 0.0 | 0.5 | 0.9 | 0.0 | 0.6 |
| 93 | 0.67 | 1.0 | 0.7 | 0.0 | 0.3 | 2.4 | 0.0 | 0.0 | 1.9 | 0.0 | 0.4 | 0.9 | 0.1 | 0.6 |
| 94 | 0.65 | 0.2 | 0.1 | 0.0 | 0.2 | 0.9 | 0.0 | 0.0 | 2.3 | 0.0 | 0.4 | 0.7 | 0.0 | 0.3 |
| 95 | 1.06 | 0.3 | 0.2 | 0.1 | 0.5 | 0.8 | 0.0 | 0.4 | 2.1 | 0.1 | 0.6 | 0.8 | 0.3 | 0.4 |
| 96 | 8.86 | 0.0 | 0.0 | 0.0 | 0.0 | 0.1 | 0.0 | 0.0 | 0.0 | 0.0 | 0.0 | 0.1 | 0.2 | 0.0 |
| 97 | 1.13 | 0.1 | 0.0 | 0.0 | 0.1 | 0.2 | 0.0 | 0.0 | 2.5 | 0.0 | 0.3 | 0.5 | 0.0 | 0.2 |
| 98 | 0.51 | 0.7 | 0.4 | 0.0 | 0.2 | 1.8 | 0.0 | 0.0 | 1.9 | 0.0 | 0.5 | 1.1 | 0.0 | 0.5 |
| 99 | 0.97 | 0.6 | 0.4 | 0.0 | 0.2 | 1.5 | 0.0 | 0.0 | 1.5 | 0.0 | 0.5 | 1.1 | 0.0 | 0.5 |
| 100 | 0.25 | 4.2 | 1.6 | 2.5 | 1.1 | 1.7 | 0.3 | 6.3 | 8.2 | 2.2 | 2.7 | 1.6 | 6.5 | 0.9 |
| Base | 100 | 10.00 | 10.00 | 9.95 | 9.95 | 10.00 | 10.02 | 10.00 | 10.00 | 10.03 | 10.88 | 10.03 | 10.00 | 10.00 |

## B

#### Enrichments of top 10% ranked bases non-coding restricted

| State | Non coding region % | CADD 1.4 | CADD 1.0 | Eigen | Eigen-PC | DANN | GERP++ | phyloP | PhastCons | REMM | fitCons | FIRE | FATHMM-XF | CDTS | LINSIGHT | FunSeq2 |
|---|---|---|---|---|---|---|---|---|---|---|---|---|---|---|---|---|
| 1 | 0.13 | 9.1 | 7.7 | 9.3 | 6.9 | 5.2 | 8.3 | 9.0 | 9.0 | 8.5 | 3.6 | 1.2 | 9.6 | 3.0 | 9.5 | 4.9 |
| 2 | 0.83 | 7.7 | 6.8 | 8.7 | 4.5 | 2.0 | 7.2 | 7.1 | 7.6 | 6.9 | 2.2 | 0.6 | 9.3 | 1.8 | 8.4 | 3.6 |
| 3 | 0.33 | 6.0 | 5.0 | 7.5 | 2.9 | 0.9 | 5.0 | 3.4 | 4.8 | 5.2 | 2.5 | 1.1 | 8.7 | 2.0 | 7.4 | 3.1 |
| 4 | 0.54 | 6.4 | 5.5 | 7.7 | 3.4 | 1.5 | 5.9 | 4.8 | 5.5 | 5.7 | 2.3 | 1.1 | 7.3 | 2.0 | 7.5 | 3.4 |
| 5 | 1.80 | 6.0 | 5.3 | 7.8 | 3.1 | 1.2 | 6.3 | 4.9 | 5.2 | 5.5 | 1.8 | 0.9 | 5.9 | 1.6 | 7.1 | 3.2 |
| 6 | 3.95 | 2.5 | 2.5 | 3.8 | 1.8 | 1.0 | 3.9 | 2.9 | 2.0 | 2.9 | 1.3 | 0.8 | 2.4 | 1.3 | 3.5 | 2.3 |
| 7 | 0.91 | 2.5 | 2.7 | 4.3 | 2.4 | 1.2 | 3.1 | 2.0 | 1.4 | 2.7 | 2.7 | 2.2 | 4.0 | 2.2 | 3.9 | 2.9 |
| 8 | 0.79 | 2.9 | 3.0 | 4.7 | 2.0 | 0.7 | 2.8 | 1.4 | 1.7 | 2.8 | 1.8 | 1.1 | 3.3 | 1.5 | 4.3 | 2.2 |
| 9 | 0.99 | 1.4 | 1.7 | 2.1 | 1.6 | 0.7 | 1.8 | 0.7 | 0.7 | 1.9 | 1.3 | 0.9 | 1.5 | 1.3 | 2.2 | 1.6 |
| 10 | 0.95 | 1.0 | 1.4 | 1.6 | 1.6 | 0.6 | 1.0 | 0.4 | 0.5 | 1.6 | 1.3 | 0.9 | 1.3 | 1.3 | 2.0 | 1.4 |
| 11 | 0.83 | 0.9 | 1.2 | 1.4 | 1.4 | 0.8 | 1.9 | 1.3 | 0.6 | 1.6 | 1.2 | 1.0 | 1.2 | 1.2 | 1.4 | 1.5 |
| 12 | 0.83 | 1.0 | 1.2 | 1.4 | 1.2 | 0.9 | 2.1 | 1.5 | 0.7 | 1.5 | 1.0 | 0.7 | 1.3 | 1.0 | 1.5 | 1.3 |
| 13 | 1.37 | 1.2 | 1.5 | 1.8 | 1.2 | 0.8 | 2.5 | 1.7 | 0.8 | 1.7 | 1.0 | 0.7 | 1.3 | 1.1 | 1.8 | 1.6 |
| 14 | 1.03 | 1.3 | 1.5 | 2.0 | 1.8 | 0.9 | 2.3 | 1.6 | 0.8 | 2.1 | 1.5 | 1.3 | 1.2 | 1.4 | 1.8 | 2.0 |
| 15 | 0.85 | 1.0 | 1.4 | 1.6 | 1.7 | 0.6 | 1.1 | 0.4 | 0.4 | 1.7 | 1.4 | 1.0 | 1.3 | 1.4 | 2.0 | 1.5 |
| 16 | 0.79 | 0.5 | 0.8 | 0.6 | 1.1 | 0.6 | 0.8 | 0.3 | 0.2 | 1.0 | 1.0 | 0.8 | 0.8 | 1.1 | 0.7 | 1.0 |
| 17 | 2.36 | 1.0 | 1.2 | 1.3 | 1.1 | 0.9 | 2.5 | 2.1 | 0.7 | 1.5 | 1.0 | 0.8 | 1.3 | 1.1 | 1.3 | 1.5 |
| 18 | 1.04 | 0.4 | 0.7 | 0.6 | 1.2 | 0.6 | 0.5 | 0.2 | 0.2 | 1.0 | 1.0 | 0.8 | 0.7 | 1.1 | 0.9 | 1.0 |
| 19 | 1.17 | 1.5 | 1.8 | 2.4 | 1.8 | 0.6 | 1.3 | 0.6 | 0.8 | 2.0 | 1.5 | 1.0 | 1.8 | 1.4 | 2.8 | 1.6 |
| 20 | 1.13 | 1.2 | 1.4 | 1.8 | 1.4 | 0.9 | 2.2 | 1.5 | 0.8 | 1.7 | 1.2 | 0.8 | 1.3 | 1.2 | 1.8 | 1.6 |
| 21 | 0.71 | 1.0 | 1.2 | 1.4 | 1.3 | 0.8 | 1.9 | 1.3 | 0.6 | 1.6 | 1.2 | 0.9 | 1.3 | 1.2 | 1.6 | 1.4 |
| 22 | 0.45 | 0.7 | 0.9 | 0.9 | 1.3 | 0.9 | 1.4 | 1.2 | 0.4 | 1.4 | 1.2 | 1.0 | 0.9 | 1.2 | 0.9 | 1.3 |
| 23 | 0.69 | 0.6 | 0.6 | 0.4 | 1.0 | 0.9 | 0.7 | 1.4 | 0.5 | 1.0 | 1.1 | 1.1 | 0.8 | 1.0 | 0.3 | 1.0 |
| 24 | 0.59 | 0.5 | 0.6 | 0.3 | 0.8 | 0.9 | 0.7 | 1.3 | 0.4 | 0.9 | 0.9 | 0.9 | 0.8 | 0.9 | 0.3 | 0.8 |
| 25 | 0.68 | 0.5 | 0.7 | 0.4 | 0.8 | 0.8 | 1.1 | 1.1 | 0.4 | 0.9 | 0.8 | 0.7 | 0.9 | 0.9 | 0.4 | 0.9 |
| 26 | 0.80 | 0.7 | 0.9 | 0.7 | 0.9 | 0.8 | 1.5 | 1.3 | 0.5 | 1.1 | 1.0 | 0.9 | 1.0 | 1.0 | 0.5 | 1.2 |
| 27 | 1.05 | 0.4 | 0.7 | 0.5 | 1.1 | 0.6 | 0.4 | 0.2 | 0.2 | 0.9 | 1.2 | 1.0 | 0.7 | 1.1 | 0.6 | 1.0 |
| 28 | 0.44 | 2.2 | 1.6 | 3.4 | 4.4 | 2.3 | 1.5 | 1.8 | 1.7 | 3.0 | 2.7 | 1.8 | 2.2 | 1.8 | 2.8 | 2.5 |
| 29 | 1.01 | 0.5 | 0.7 | 0.4 | 0.9 | 0.8 | 1.1 | 1.1 | 0.3 | 0.9 | 0.7 | 0.6 | 0.8 | 0.9 | 0.6 | 0.8 |
| 30 | 1.03 | 0.5 | 0.6 | 0.3 | 0.7 | 0.8 | 0.9 | 1.2 | 0.4 | 0.8 | 0.7 | 0.7 | 0.8 | 0.9 | 0.2 | 0.7 |
| 31 | 0.70 | 0.5 | 0.6 | 0.3 | 0.8 | 0.9 | 0.6 | 1.3 | 0.4 | 0.9 | 0.9 | 0.9 | 0.7 | 0.9 | 0.2 | 0.7 |
| 32 | 1.00 | 0.6 | 0.6 | 0.2 | 0.7 | 1.1 | 0.1 | 2.0 | 0.6 | 0.6 | 0.8 | 0.9 | 0.5 | 0.9 | 0.1 | 0.6 |
| 33 | 0.49 | 0.8 | 1.0 | 0.3 | 0.6 | 1.5 | 0.3 | 1.6 | 0.6 | 0.6 | 0.7 | 0.9 | 0.5 | 0.8 | 0.3 | 0.5 |
| 34 | 0.56 | 1.0 | 1.2 | 0.9 | 0.8 | 1.1 | 1.4 | 1.4 | 0.6 | 1.1 | 0.9 | 1.0 | 1.1 | 1.0 | 0.9 | 1.0 |
| 35 | 1.33 | 0.8 | 1.0 | 1.1 | 1.4 | 0.9 | 1.6 | 1.5 | 0.7 | 1.5 | 1.4 | 1.3 | 1.1 | 1.3 | 0.8 | 1.6 |
| 36 | 1.47 | 0.6 | 0.8 | 0.5 | 0.8 | 0.9 | 1.3 | 1.5 | 0.6 | 1.0 | 0.9 | 0.9 | 1.1 | 1.0 | 0.4 | 1.0 |
| 37 | 2.54 | 1.1 | 1.2 | 1.5 | 1.2 | 0.8 | 2.2 | 1.7 | 0.8 | 1.5 | 1.2 | 1.0 | 1.3 | 1.1 | 1.2 | 1.6 |
| 38 | 1.00 | 0.8 | 0.9 | 0.8 | 0.9 | 0.8 | 1.8 | 1.5 | 0.6 | 1.1 | 1.0 | 0.9 | 1.1 | 1.0 | 0.6 | 1.2 |
| 39 | 0.76 | 0.6 | 0.8 | 0.5 | 0.8 | 0.8 | 1.2 | 1.3 | 0.5 | 0.9 | 0.8 | 0.8 | 1.0 | 0.9 | 0.5 | 0.9 |
| 40 | 0.83 | 0.5 | 0.6 | 0.3 | 0.7 | 0.9 | 0.7 | 1.2 | 0.4 | 0.7 | 0.7 | 0.7 | 0.8 | 0.9 | 0.2 | 0.7 |
| 41 | 0.78 | 0.6 | 0.8 | 0.6 | 0.9 | 0.8 | 1.4 | 1.3 | 0.4 | 1.0 | 0.7 | 0.6 | 1.0 | 0.9 | 0.6 | 0.9 |
| 42 | 0.82 | 0.6 | 0.8 | 0.6 | 1.2 | 0.9 | 1.3 | 1.2 | 0.4 | 1.2 | 1.0 | 1.0 | 0.9 | 1.1 | 0.6 | 1.1 |
| 43 | 1.10 | 0.6 | 0.7 | 0.4 | 0.9 | 0.9 | 0.8 | 1.3 | 0.5 | 1.0 | 1.0 | 1.0 | 0.8 | 1.0 | 0.3 | 0.9 |
| 44 | 1.12 | 0.7 | 0.7 | 0.3 | 0.8 | 1.2 | 0.1 | 1.9 | 0.6 | 0.7 | 0.9 | 1.1 | 0.7 | 0.9 | 0.2 | 0.6 |
| 45 | 0.93 | 0.5 | 0.7 | 0.3 | 0.8 | 0.9 | 0.7 | 1.4 | 0.4 | 0.8 | 0.8 | 0.9 | 0.7 | 0.9 | 0.3 | 0.7 |
| 46 | 0.77 | 0.6 | 0.7 | 0.5 | 0.9 | 0.9 | 1.0 | 1.3 | 0.5 | 1.0 | 1.0 | 1.1 | 0.9 | 1.0 | 0.4 | 1.0 |
| 47 | 0.45 | 0.2 | 0.4 | 0.3 | 1.4 | 0.4 | 0.1 | 0.0 | 0.0 | 0.5 | 1.2 | 0.9 | 0.7 | 1.2 | 0.7 | 1.0 |
| 48 | 0.59 | 0.2 | 0.3 | 0.2 | 0.9 | 0.4 | 0.1 | 0.1 | 0.2 | 0.3 | 0.9 | 0.9 | 0.5 | 1.0 | 0.3 | 0.7 |
| 49 | 0.67 | 0.3 | 0.4 | 0.4 | 1.3 | 0.4 | 0.2 | 0.2 | 0.2 | 0.5 | 1.3 | 1.1 | 0.7 | 1.2 | 0.6 | 1.1 |
| 50 | 0.87 | 0.9 | 0.9 | 1.6 | 1.8 | 0.3 | 0.9 | 0.5 | 0.8 | 0.5 | 1.5 | 1.0 | 1.7 | 1.5 | 2.4 | 1.5 |
| 51 | 0.81 | 0.2 | 0.4 | 0.4 | 1.3 | 0.3 | 0.3 | 0.2 | 0.2 | 0.3 | 1.1 | 0.8 | 0.6 | 1.1 | 0.8 | 0.9 |
| 52 | 0.55 | 0.3 | 0.6 | 0.5 | 1.4 | 0.5 | 0.3 | 0.1 | 0.1 | 0.9 | 1.1 | 0.9 | 0.7 | 1.1 | 0.9 | 1.0 |
| 53 | 0.55 | 1.1 | 1.4 | 1.8 | 1.9 | 0.6 | 0.6 | 0.3 | 0.5 | 1.6 | 1.6 | 1.1 | 1.7 | 1.5 | 2.5 | 1.5 |
| 54 | 0.23 | 2.2 | 1.9 | 3.4 | 2.3 | 0.5 | 1.4 | 1.2 | 2.0 | 1.0 | 2.6 | 1.6 | 5.8 | 2.1 | 4.4 | 2.0 |
| 55 | 0.33 | 0.9 | 1.3 | 1.5 | 1.7 | 0.4 | 1.1 | 0.5 | 0.9 | 0.1 | 1.5 | 1.0 | 1.7 | 1.4 | 2.4 | 1.5 |
| 56 | 0.42 | 0.2 | 0.7 | 0.3 | 1.1 | 0.5 | 0.4 | 0.2 | 0.2 | 0.0 | 1.0 | 0.8 | 0.6 | 1.0 | 0.7 | 0.9 |
| 57 | 0.43 | 0.1 | 0.6 | 0.2 | 1.0 | 0.5 | 0.2 | 0.1 | 0.2 | 0.0 | 1.0 | 0.9 | 0.5 | 1.0 | 0.3 | 0.8 |
| 58 | 0.51 | 0.2 | 0.4 | 0.2 | 1.1 | 0.4 | 0.1 | 0.1 | 0.1 | 0.4 | 1.0 | 1.0 | 0.4 | 1.0 | 0.4 | 0.8 |
| 59 | 0.58 | 0.2 | 0.3 | 0.2 | 1.2 | 0.5 | 0.0 | 0.0 | 0.2 | 0.4 | 1.1 | 1.0 | 0.3 | 1.0 | 0.2 | 0.7 |
| 60 | 0.34 | 0.1 | 0.3 | 0.1 | 0.9 | 0.4 | 0.0 | 0.0 | 0.1 | 0.3 | 0.8 | 0.7 | 0.3 | 0.9 | 0.2 | 0.6 |
| 61 | 0.53 | 0.1 | 0.3 | 0.1 | 0.9 | 0.4 | 0.1 | 0.0 | 0.1 | 0.3 | 0.8 | 0.7 | 0.3 | 0.9 | 0.2 | 0.6 |
| 62 | 0.49 | 0.1 | 0.3 | 0.2 | 1.0 | 0.4 | 0.1 | 0.1 | 0.1 | 0.3 | 1.0 | 0.9 | 0.4 | 1.0 | 0.2 | 0.8 |
| 63 | 0.45 | 0.3 | 0.5 | 0.6 | 1.5 | 0.4 | 0.3 | 0.2 | 0.2 | 0.6 | 1.3 | 0.9 | 0.7 | 1.2 | 0.9 | 1.1 |
| 64 | 0.87 | 0.5 | 0.7 | 0.4 | 0.8 | 0.9 | 0.9 | 1.1 | 0.4 | 0.9 | 0.8 | 0.8 | 0.7 | 0.9 | 0.4 | 0.8 |
| 65 | 1.03 | 0.4 | 0.5 | 0.2 | 1.0 | 0.8 | 0.2 | 1.3 | 0.3 | 0.7 | 0.9 | 1.0 | 0.4 | 1.0 | 0.2 | 0.7 |
| 66 | 0.73 | 0.6 | 0.7 | 0.6 | 1.3 | 0.8 | 0.9 | 0.9 | 0.3 | 1.2 | 1.1 | 1.0 | 0.7 | 1.1 | 0.9 | 1.1 |
| 67 | 1.08 | 0.4 | 0.5 | 0.4 | 1.1 | 0.8 | 0.3 | 1.2 | 0.2 | 0.8 | 1.0 | 0.9 | 0.5 | 1.0 | 0.5 | 0.8 |
| 68 | 1.30 | 0.4 | 0.4 | 0.1 | 0.6 | 0.9 | 0.1 | 1.2 | 0.3 | 0.5 | 0.8 | 1.0 | 0.2 | 0.9 | 0.1 | 0.5 |
| 69 | 1.01 | 0.5 | 0.6 | 0.2 | 0.7 | 1.1 | 0.1 | 1.7 | 0.5 | 0.5 | 0.7 | 0.8 | 0.4 | 0.8 | 0.1 | 0.5 |
| 70 | 0.55 | 0.2 | 0.3 | 0.1 | 0.9 | 0.6 | 0.0 | 0.0 | 0.2 | 0.3 | 0.8 | 0.8 | 0.1 | 0.9 | 0.1 | 0.6 |
| 71 | 0.45 | 0.2 | 0.3 | 0.1 | 1.0 | 0.6 | 0.0 | 0.0 | 0.1 | 0.3 | 0.9 | 1.0 | 0.2 | 0.9 | 0.1 | 0.6 |
| 72 | 1.31 | 0.5 | 0.6 | 0.2 | 0.8 | 1.1 | 0.1 | 1.7 | 0.5 | 0.6 | 0.8 | 1.1 | 0.3 | 0.9 | 0.1 | 0.6 |
| 73 | 0.81 | 0.4 | 0.4 | 0.1 | 0.6 | 1.0 | 0.0 | 0.9 | 0.4 | 0.4 | 0.9 | 1.3 | 0.2 | 1.0 | 0.1 | 0.6 |
| 74 | 1.64 | 0.4 | 0.3 | 0.2 | 0.7 | 0.8 | 0.1 | 0.2 | 0.4 | 0.5 | 1.0 | 1.3 | 0.2 | 1.0 | 0.1 | 0.7 |
| 75 | 2.73 | 0.5 | 0.4 | 0.1 | 0.5 | 1.0 | 0.0 | 0.2 | 0.4 | 0.3 | 1.0 | 1.4 | 0.1 | 1.0 | 0.1 | 0.6 |
| 76 | 4.33 | 0.8 | 0.6 | 0.1 | 0.4 | 1.2 | 0.0 | 0.0 | 1.1 | 0.2 | 0.7 | 1.0 | 0.1 | 0.8 | 0.1 | 0.4 |
| 77 | 1.08 | 0.5 | 0.4 | 0.1 | 0.4 | 1.1 | 0.0 | 0.1 | 0.7 | 0.2 | 1.1 | 1.8 | 0.1 | 1.0 | 0.1 | 0.6 |
| 78 | 1.26 | 0.7 | 0.5 | 0.1 | 0.4 | 1.0 | 0.0 | 0.0 | 0.9 | 0.2 | 0.8 | 1.3 | 0.1 | 0.9 | 0.1 | 0.5 |
| 79 | 3.51 | 0.9 | 0.6 | 0.1 | 0.3 | 1.6 | 0.0 | 0.0 | 1.5 | 0.1 | 0.7 | 1.0 | 0.1 | 0.8 | 0.0 | 0.4 |
| 80 | 1.37 | 0.4 | 0.4 | 0.1 | 0.4 | 1.1 | 0.0 | 0.2 | 0.4 | 0.2 | 0.9 | 1.6 | 0.1 | 1.0 | 0.1 | 0.6 |
| 81 | 1.08 | 0.7 | 0.5 | 0.1 | 0.5 | 1.4 | 0.0 | 0.2 | 1.0 | 0.1 | 0.8 | 1.2 | 0.1 | 0.9 | 0.1 | 0.4 |
| 82 | 0.57 | 1.0 | 0.5 | 0.7 | 2.1 | 1.0 | 0.1 | 1.8 | 2.3 | 0.8 | 1.6 | 1.4 | 1.3 | 0.8 | 1.3 | 0.8 |
| 83 | 1.38 | 1.0 | 0.7 | 0.1 | 0.2 | 1.4 | 0.0 | 0.0 | 1.1 | 0.1 | 0.6 | 0.9 | 0.0 | 0.6 | 0.0 | 0.3 |
| 84 | 0.91 | 0.6 | 0.5 | 0.1 | 0.4 | 1.3 | 0.0 | 0.2 | 0.6 | 0.2 | 0.7 | 1.2 | 0.1 | 0.6 | 0.1 | 0.4 |
| 85 | 1.46 | 0.7 | 0.5 | 0.0 | 0.2 | 1.4 | 0.0 | 0.0 | 1.3 | 0.0 | 0.7 | 1.5 | 0.0 | 0.7 | 0.0 | 0.4 |
| 86 | 1.73 | 0.9 | 0.6 | 0.0 | 0.2 | 1.8 | 0.0 | 0.0 | 1.6 | 0.0 | 0.5 | 0.9 | 0.0 | 0.6 | 0.0 | 0.3 |
| 87 | 0.69 | 0.0 | 0.1 | 0.0 | 0.3 | 0.3 | 0.0 | 0.0 | 0.2 | 0.0 | 0.8 | 1.2 | 0.1 | 0.8 | 0.0 | 0.5 |
| 88 | 0.73 | 0.0 | 0.1 | 0.0 | 0.4 | 0.3 | 0.0 | 0.0 | 0.2 | 0.0 | 0.9 | 1.1 | 0.1 | 0.9 | 0.1 | 0.5 |
| 89 | 0.41 | 0.1 | 0.4 | 0.1 | 1.0 | 0.5 | 0.0 | 0.0 | 0.1 | 0.0 | 0.9 | 0.9 | 0.2 | 0.9 | 0.2 | 0.6 |
| 90 | 1.19 | 1.4 | 1.1 | 0.2 | 0.3 | 1.8 | 0.0 | 0.0 | 1.0 | 0.2 | 0.6 | 1.0 | 0.1 | 0.8 | 0.1 | 0.4 |
| 91 | 1.07 | 0.6 | 0.6 | 0.1 | 0.5 | 1.5 | 0.0 | 0.5 | 0.5 | 0.2 | 0.7 | 1.1 | 0.1 | 0.8 | 0.1 | 0.4 |
| 92 | 2.00 | 0.8 | 0.5 | 0.0 | 0.2 | 2.1 | 0.0 | 0.0 | 1.6 | 0.0 | 0.5 | 0.9 | 0.0 | 0.6 | 0.0 | 0.3 |
| 93 | 0.74 | 1.0 | 0.6 | 0.0 | 0.2 | 2.4 | 0.0 | 0.0 | 1.9 | 0.0 | 0.4 | 0.8 | 0.0 | 0.5 | 0.0 | 0.2 |
| 94 | 0.72 | 0.2 | 0.1 | 0.0 | 0.2 | 0.9 | 0.0 | 0.0 | 2.3 | 0.0 | 0.4 | 0.6 | 0.0 | 0.3 | 0.0 | 0.2 |
| 95 | 1.08 | 0.3 | 0.2 | 0.1 | 0.5 | 0.9 | 0.0 | 0.3 | 2.1 | 0.1 | 0.6 | 0.8 | 0.3 | 0.4 | 0.2 | 0.3 |
| 96 | 1.33 | 0.0 | 0.0 | 0.0 | 0.2 | 1.0 | 0.0 | 0.0 | 0.1 | 0.0 | 0.3 | 0.4 | 1.5 | 0.1 | 0.0 | 0.2 |
| 97 | 0.96 | 0.1 | 0.0 | 0.0 | 0.1 | 0.2 | 0.0 | 0.0 | 2.5 | 0.0 | 0.4 | 0.6 | 0.0 | 0.2 | 0.0 | 0.2 |
| 98 | 0.56 | 0.7 | 0.4 | 0.0 | 0.2 | 1.8 | 0.0 | 0.0 | 1.9 | 0.0 | 0.5 | 1.1 | 0.0 | 0.5 | 0.0 | 0.2 |
| 99 | 1.05 | 0.5 | 0.4 | 0.0 | 0.2 | 1.5 | 0.0 | 0.0 | 1.5 | 0.0 | 0.5 | 1.0 | 0.0 | 0.4 | 0.0 | 0.3 |
| 100 | 0.26 | 4.0 | 1.5 | 2.5 | 1.1 | 1.5 | 0.3 | 6.3 | 8.3 | 2.2 | 2.5 | 1.6 | 6.8 | 0.9 | 5.4 | 0.6 |
| Base | 90.44 | 9.92 | 9.93 | 9.93 | 9.91 | 9.91 | 10.01 | 9.93 | 9.93 | 9.96 | 10.74 | 9.91 | 9.93 | 9.93 | 9.93 | 9.91 |

**Figure S27: Enrichment of all conservation states for top 10% of bases prioritized by variant prioritization scores.** The figure displays for top 10% prioritized bases **(A)** genome-wide and **(B)** in non-coding regions by variant prioritization scores the same enrichments as shown in **Figure S24C** and **S24D** except here enrichments for all conservation states are shown and also included is the original version of the CADD score in addition to v1.4. Coloring of enrichments is based on their value in a column specific manner. The second columns gives the percentage of the background region used to compute the enrichments falling in each state, which is for **(A)** the whole genome and for **(B)** bases scored by both LINSIGHT and FunSeq2. The last line in both heatmaps gives the actual percentage of the background set covered by each set of prioritized bases, which can differ from 10.00% because of how ties were handled.

| | MAF < 0.001 | 0.001 <= MAF < 0.01 | 0.01 <= MAF < 0.1 | 0.1 <= MAF < 0.2 | 0.2 <= MAF < 0.3 | 0.3 <= MAF < 0.4 | 0.4 <= MAF < 0.5 |
|---|---|---|---|---|---|---|---|
| 1 | -0.52 | -1.05 | -1.64 | -2.26 | -2.53 | -2.66 | -2.80 |
| 2 | -0.20 | -0.31 | -0.63 | -1.04 | -1.26 | -1.43 | -1.52 |
| 3 | 0.22 | 0.08 | -0.21 | -0.60 | -0.75 | -0.92 | -1.03 |
| 4 | -0.10 | -0.21 | -0.50 | -0.88 | -1.07 | -1.26 | -1.30 |
| 5 | -0.19 | -0.24 | -0.48 | -0.81 | -1.04 | -1.17 | -1.27 |
| 6 | -0.17 | -0.20 | -0.35 | -0.64 | -0.80 | -0.95 | -1.00 |
| 7 | -0.06 | -0.09 | -0.31 | -0.64 | -0.82 | -0.98 | -1.00 |
| 8 | 0.06 | 0.05 | -0.14 | -0.40 | -0.61 | -0.68 | -0.76 |
| 9 | 0.01 | 0.00 | -0.16 | -0.42 | -0.55 | -0.69 | -0.80 |
| 10 | 0.10 | 0.11 | -0.04 | -0.29 | -0.45 | -0.58 | -0.65 |
| 11 | -0.06 | -0.04 | -0.17 | -0.36 | -0.51 | -0.68 | -0.74 |
| 12 | -0.11 | -0.06 | -0.20 | -0.41 | -0.55 | -0.72 | -0.74 |
| 13 | -0.10 | -0.06 | -0.20 | -0.42 | -0.59 | -0.75 | -0.81 |
| 14 | -0.11 | -0.08 | -0.22 | -0.49 | -0.64 | -0.77 | -0.82 |
| 15 | 0.16 | 0.15 | 0.01 | -0.19 | -0.32 | -0.39 | -0.45 |
| 16 | 0.12 | 0.14 | 0.04 | -0.16 | -0.26 | -0.37 | -0.38 |
| 17 | -0.16 | -0.15 | -0.27 | -0.52 | -0.64 | -0.78 | -0.85 |
| 18 | 0.14 | 0.17 | 0.05 | -0.21 | -0.34 | -0.45 | -0.50 |
| 19 | 0.13 | 0.11 | -0.04 | -0.33 | -0.48 | -0.59 | -0.69 |
| 20 | -0.04 | -0.04 | -0.17 | -0.44 | -0.63 | -0.70 | -0.78 |
| 21 | -0.06 | -0.02 | -0.13 | -0.37 | -0.54 | -0.68 | -0.70 |
| 22 | -0.03 | -0.01 | -0.13 | -0.39 | -0.50 | -0.63 | -0.70 |
| 23 | -0.03 | 0.00 | -0.11 | -0.32 | -0.43 | -0.58 | -0.59 |
| 24 | -0.08 | -0.01 | -0.12 | -0.32 | -0.43 | -0.55 | -0.59 |
| 25 | -0.03 | 0.00 | -0.11 | -0.31 | -0.43 | -0.57 | -0.63 |
| 26 | -0.09 | -0.05 | -0.17 | -0.40 | -0.55 | -0.64 | -0.71 |
| 27 | 0.12 | 0.15 | 0.04 | -0.21 | -0.36 | -0.50 | -0.55 |
| 28 | -0.04 | -0.13 | -0.28 | -0.56 | -0.71 | -0.74 | -0.85 |
| 29 | -0.04 | 0.01 | -0.08 | -0.29 | -0.46 | -0.58 | -0.56 |
| 30 | -0.07 | -0.02 | -0.11 | -0.34 | -0.47 | -0.57 | -0.65 |
| 31 | -0.04 | 0.01 | -0.09 | -0.29 | -0.44 | -0.57 | -0.60 |
| 32 | -0.07 | -0.08 | -0.17 | -0.41 | -0.54 | -0.64 | -0.72 |
| 33 | -0.13 | -0.09 | -0.16 | -0.31 | -0.39 | -0.53 | -0.62 |
| 34 | -0.09 | -0.06 | -0.16 | -0.36 | -0.50 | -0.55 | -0.65 |
| 35 | -0.08 | -0.05 | -0.17 | -0.39 | -0.54 | -0.70 | -0.71 |
| 36 | -0.10 | -0.07 | -0.18 | -0.38 | -0.53 | -0.63 | -0.67 |
| 37 | -0.14 | -0.10 | -0.25 | -0.50 | -0.64 | -0.76 | -0.83 |
| 38 | -0.09 | -0.05 | -0.17 | -0.40 | -0.60 | -0.68 | -0.71 |
| 39 | -0.14 | -0.09 | -0.19 | -0.41 | -0.55 | -0.64 | -0.69 |
| 40 | -0.05 | -0.01 | -0.11 | -0.33 | -0.42 | -0.55 | -0.61 |
| 41 | -0.06 | 0.00 | -0.11 | -0.33 | -0.45 | -0.53 | -0.65 |
| 42 | -0.05 | -0.02 | -0.13 | -0.36 | -0.46 | -0.62 | -0.67 |
| 43 | -0.03 | 0.01 | -0.07 | -0.30 | -0.42 | -0.52 | -0.61 |
| 44 | -0.20 | -0.18 | -0.27 | -0.43 | -0.60 | -0.66 | -0.74 |
| 45 | -0.06 | -0.03 | -0.11 | -0.32 | -0.47 | -0.59 | -0.66 |
| 46 | -0.10 | -0.09 | -0.22 | -0.42 | -0.55 | -0.65 | -0.70 |
| 47 | 0.42 | 0.41 | 0.33 | 0.18 | 0.02 | -0.05 | -0.09 |
| 48 | 0.31 | 0.31 | 0.20 | 0.00 | -0.10 | -0.27 | -0.37 |
| 49 | 0.34 | 0.33 | 0.20 | -0.04 | -0.19 | -0.30 | -0.32 |
| 50 | 0.50 | 0.43 | 0.27 | 0.04 | -0.12 | -0.26 | -0.27 |
| 51 | 0.46 | 0.43 | 0.31 | 0.12 | -0.03 | -0.12 | -0.17 |
| 52 | 0.14 | 0.16 | 0.02 | -0.22 | -0.38 | -0.50 | -0.61 |
| 53 | 0.21 | 0.17 | 0.00 | -0.30 | -0.48 | -0.60 | -0.68 |
| 54 | 0.87 | 0.83 | 1.08 | 1.44 | 1.65 | 1.79 | 1.81 |
| 55 | 1.50 | 1.69 | 2.70 | 3.59 | 3.92 | 4.12 | 4.21 |
| 56 | 1.46 | 1.65 | 2.62 | 3.50 | 3.83 | 4.02 | 4.12 |
| 57 | 1.47 | 1.69 | 2.72 | 3.61 | 3.95 | 4.14 | 4.24 |
| 58 | 0.38 | 0.37 | 0.27 | 0.11 | 0.02 | -0.10 | -0.15 |
| 59 | 0.34 | 0.35 | 0.30 | 0.14 | 0.11 | 0.04 | -0.02 |
| 60 | 0.36 | 0.37 | 0.26 | 0.07 | -0.03 | -0.11 | -0.19 |
| 61 | 0.38 | 0.39 | 0.28 | 0.11 | -0.01 | -0.10 | -0.18 |
| 62 | 0.39 | 0.38 | 0.29 | 0.11 | 0.01 | -0.08 | -0.13 |
| 63 | 0.39 | 0.37 | 0.26 | 0.03 | -0.12 | -0.23 | -0.28 |
| 64 | -0.04 | -0.02 | -0.13 | -0.36 | -0.54 | -0.62 | -0.75 |
| 65 | 0.06 | 0.09 | -0.01 | -0.25 | -0.40 | -0.53 | -0.60 |
| 66 | 0.08 | 0.10 | -0.04 | -0.26 | -0.37 | -0.52 | -0.54 |
| 67 | 0.13 | 0.14 | 0.03 | -0.19 | -0.34 | -0.46 | -0.56 |
| 68 | 0.08 | 0.10 | -0.02 | -0.24 | -0.39 | -0.54 | -0.60 |
| 69 | -0.06 | -0.01 | -0.11 | -0.32 | -0.49 | -0.57 | -0.66 |
| 70 | 0.32 | 0.32 | 0.25 | 0.10 | 0.03 | -0.10 | -0.11 |
| 71 | 0.35 | 0.32 | 0.21 | 0.00 | -0.14 | -0.23 | -0.24 |
| 72 | -0.05 | -0.07 | -0.18 | -0.41 | -0.56 | -0.71 | -0.72 |
| 73 | 0.08 | 0.10 | 0.00 | -0.16 | -0.31 | -0.43 | -0.46 |
| 74 | 0.11 | 0.12 | -0.01 | -0.27 | -0.43 | -0.54 | -0.60 |
| 75 | 0.18 | 0.17 | 0.04 | -0.21 | -0.36 | -0.50 | -0.57 |
| 76 | 0.18 | 0.15 | 0.04 | -0.21 | -0.35 | -0.47 | -0.56 |
| 77 | 0.09 | 0.08 | -0.06 | -0.33 | -0.50 | -0.62 | -0.69 |
| 78 | 0.13 | 0.13 | 0.01 | -0.23 | -0.37 | -0.49 | -0.58 |
| 79 | 0.22 | 0.17 | 0.04 | -0.19 | -0.34 | -0.46 | -0.53 |
| 80 | 0.26 | 0.26 | 0.13 | -0.12 | -0.31 | -0.45 | -0.49 |
| 81 | 0.19 | 0.16 | 0.05 | -0.16 | -0.36 | -0.48 | -0.56 |
| 82 | -0.23 | -0.20 | -0.23 | -0.32 | -0.41 | -0.41 | -0.41 |
| 83 | -0.13 | -0.21 | -0.33 | -0.58 | -0.72 | -0.79 | -0.90 |
| 84 | -0.43 | -0.46 | -0.57 | -0.81 | -0.96 | -1.12 | -1.11 |
| 85 | -0.05 | -0.10 | -0.22 | -0.47 | -0.63 | -0.73 | -0.83 |
| 86 | 0.26 | 0.16 | 0.09 | -0.02 | -0.12 | -0.18 | -0.24 |
| 87 | 1.41 | 1.49 | 2.10 | 2.76 | 3.00 | 3.17 | 3.24 |
| 88 | 1.37 | 1.45 | 2.08 | 2.74 | 3.03 | 3.18 | 3.25 |
| 89 | 1.47 | 1.69 | 2.71 | 3.60 | 3.92 | 4.12 | 4.22 |
| 90 | 0.20 | 0.17 | 0.19 | 0.19 | 0.19 | 0.19 | 0.18 |
| 91 | 0.23 | 0.28 | 0.29 | 0.29 | 0.27 | 0.28 | 0.22 |
| 92 | 0.42 | 0.35 | 0.37 | 0.34 | 0.32 | 0.35 | 0.27 |
| 93 | 0.38 | 0.29 | 0.32 | 0.35 | 0.31 | 0.34 | 0.34 |
| 94 | 0.16 | 0.09 | 0.09 | 0.07 | 0.04 | 0.01 | 0.03 |
| 95 | -0.10 | -0.18 | -0.15 | -0.19 | -0.25 | -0.25 | -0.31 |
| 96 | -3.18 | -3.34 | -3.38 | -3.51 | -3.70 | -3.60 | -3.48 |
| 97 | -0.70 | -0.78 | -0.79 | -0.86 | -0.93 | -0.98 | -1.05 |
| 98 | 0.16 | 0.07 | 0.07 | 0.03 | 0.03 | -0.07 | -0.05 |
| 99 | 0.09 | 0.03 | 0.02 | -0.02 | -0.05 | -0.09 | -0.13 |
| 100 | -0.52 | -0.57 | -0.52 | -0.55 | -0.56 | -0.63 | -0.56 |
| % genome | 1.46 | 0.39 | 0.18 | 0.05 | 0.04 | 0.03 | 0.03 |

**Figure S28: Conservation state enrichments for single nucleotide variants from Iulio et al.[37]** Rows 1-100 corresponds to states, color coded based on their group, and the last line represents the percentage of the genome covered by each annotation in the columns. The table displays the log2 fold enrichments of all the single nucleotide variants from whole genome sequencing data of 7794 unrelated individuals that were used to generate the context dependent tolerance score. The variants are grouped into disjoint sets according to minor allele frequency (MAF). Depletions are shown in shades of blue and enrichments in shades of red. The large depletions in state 96 are due to the state capturing assembly gaps.
